# FASTCAR: Rapid alignment-free prediction of sequence alignment identity scores using self-supervised general linear models

**DOI:** 10.1101/380824

**Authors:** Benjamin T. James, Brian B. Luczak, Hani Z. Girgis

## Abstract

**Motivation:** Pairwise alignment is a predominant algorithm in the field of bioinformatics. This algorithm is quadratic — slow especially on long sequences. Many applications utilize identity scores without the corresponding alignments. For these applications, we propose FASTCAR. It produces identity scores for pairs of DNA sequences using alignment-free methods and two self-supervised general linear models.

**Results:** For the first time, the new tool can predict the pair-wise identity score in linear time and space. On two large-scale sequence databases, FASTCAR provided the best compromise between sensitivity and precision while being faster than BLAST by 40% and faster than USEARCH by 6–10 times. Further, FASTCAR is capable of producing the pair-wise identity scores of long DNA sequences — millions-of-nucleotides-long bacterial genomes; this task cannot be accomplished by any alignment-based tool.

**Availability:** FASTCAR is available at https://github.com/TulsaBioinformaticsToolsmith/FASTCAR and as the Supplementary Dataset 1.

**Supplementary information:** Supplementary data are available online.

## 1 Introduction

We live in an era when sequences are generated at an unprecedented rate. Analyzing these countless sequences requires efficient computational methods. Algorithms for comparing sequence similarity are among the most fundamental tools for analyzing DNA, RNA, and protein sequences.

Alignment algorithms (Needleman and Wunsch, 1970; Gotoh, 1982) have been the standard methods for assessing sequence similarity over the past 40 years. Multiple software tools for alignment are available (Rice *et al.*, 2000; Rizk and Lavenier, 2010). Applications include gene finding (Korf, 2004), genome assembly (Butler *et al.*, 2008; Luo *et al.*, 2012; Eaton, 2014), function prediction (Peled *et al.*, 2016; Carradec *et al.*, 2018), phylogenetic trees (Costello *et al.*, 2009), and other applications.

Many advancements have been made since the Needleman-Wunsch alignment algorithm was devised (Edgar, 2010; Gotoh, 1982; Altschul *et al.*, 1990; Loh *et al.*, 2012), but these new algorithms still depend on slow, quadratic, dynamic programming. This limitation is well manifested when comparing two very long sequences or scanning a very large sequence database. Almost all of the speed-ups are based on heuristics methods.

This shortcoming of alignment algorithms has led the field to develop plenty of faster, alignment-free methods (Blaisdell, 1986; Zharkikh and Rzhetsky, 1993; Wu *et al.*, 2001; Almeida and Vinga, 2002; Lippert *et al.*, 2002; Pham and Zuegg, 2004; Kantorovitz *et al.*, 2007; Dai *et al.*, 2008; Reinert *et al.*, 2010; Sims *et al.*, 2009; Costa *et al.*, 2011; Liu *et al.*, 2011; Zhang and Chen, 2011; Göke *et al.*, 2012; Ren *et al.*, 2013; Ghandi *et al.*, 2014; Haubold, 2014; Leimeister *et al.*, 2014; Pinello *et al.*, 2014; Borozan *et al.*, 2015; Liao *et al.*, 2016). Multiple reviews of alignment-free methods have been published (Vinga and Almeida, 2003; Vinga *et al.*, 2012; Bonham-Carter *et al.*, 2014; Song *et al.*, 2014; Vinga, 2014; Chattopadhyay *et al.*, 2015; Luczak *et al.*, 2017; Zielezinski *et al.*, 2017), indicating the importance and the abundance of such methods. One particular class of these methods depends on comparing two histograms of short words called k-mers, i.e. words of fixed length k. Building the histograms and comparing them can be done very efficiently. Although these alignment-free methods are very efficient, they have not been widely adapted by the field because their scores are not as intuitive or biologically relevant as the identity scores generated by alignment algorithms. However, k-mer statistics are often used in alignment tools as heuristics, such as in BLAST and USEARCH.

Often times, the identity score alone is enough; generating the alignment itself is not needed. For example, consider scanning GenBank for similar sequences to a particular gene. For another example, consider the task of clustering a large number of sequences. In these two applications there is no need to generate the alignment; only the identity score is used.

We propose a Fast and Accurate Search Tool for Classification And Regression (FASTCAR) to predict global similarity of two DNA sequences. FASTCAR (Supplementary Dataset 1) can predict global identity scores in linear time and space for the first time. The tool utilizes self-supervised machine learning algorithms in predicting global identity scores using a small number of alignment-free, k-mer statistics. FASTCAR overcomes the weaknesses of alignment algorithms and those of alignment-free methods. It produces identity scores, which are intuitive, biologically relevant, and the standard metric in the field. Because calculating the k-mer statistics and predicting the identity score require linear time, FASTCAR is more efficient than alignment algorithms.

The core of FASTCAR is an adaptive, hierarchical, linear model for predicting the identity scores above a user provided threshold. This design was inspired by our earlier research. We have successfully implemented adaptive software tools using self-supervised learning algorithms for locating cis-regulatory modules (Girgis and Ovcharenko, 2012), identifying DNA repeats (Girgis, 2015; Velasco II *et al.*, 2018), and for clustering DNA sequences (James *et al.*, 2018; James and Girgis, 2018). Hierarchical models were reported to perform very well in ranking the quality of predicted protein structures (Girgis and Corso, 2008; Girgis, 2008; Girgis *et al.*, 2009). Multiple software tools we developed earlier utilize General Linear Models (GLMs) (Girgis and Corso, 2008; Girgis, 2008; Girgis *et al.*, 2009; Girgis and Sheetlin, 2013; James *et al.*, 2018; James and Girgis, 2018; Velasco II *et al.*, 2018). Alignment-assisted methods that can classify similar and dissimilar sequences and predict identity scores were developed (James *et al.*, 2018; Velasco II *et al.*, 2018); such methods use alignment-free statistics to predict the alignment identity scores on which they are trained, i.e. they are not alignment-free completely. These earlier tools justify our design choice of the adaptive, hierarchical, linear model as the core of FASTCAR.

The main contributions of this research are: (i) calculating identity scores in linear time and space for the first time, (ii) calculating the identity scores for pairs of very long sequences for the first time, and (iii) the FASTCAR software tool.

## 2 Methods

### Method overview

FASTCAR is an instance of self-supervised learning; such learning algorithms generate their own training data. To generate labeled data for training and testing, FASTCAR first selects randomly a small number of the input sequences — 300 by default. The selected sequences are uniformly distributed with respect to length. A few semi-synthetic sequences — 15 by default — are generated by mutating each of these sequences to generate identity scores to simulate the actual data. Since the mutated data has a known mutation rate, the identity score can be easily calculated; therefore, alignment algorithms are avoided. After that, a k-mer histogram is obtained from each sequence. A model can be trained to predict identity scores from a few statistics calculated on pairs of k-mer histograms. The advantage of this tool over the traditional alignment algorithm is that it uses a limited number of efficient, linear, k-mer statistics rather than the slow, quadratic dynamic programming utilized in alignment algorithms. Two components comprise this new predictive model. The first component is a classifier that recognizes whether the similarity between two sequences is above the desired threshold or below it. The second component is a regression model, which estimates the identity score of two sequences if they are above the threshold. In some applications, the user would be interested in finding similar sequences to a query sequence; however, the user is not interested in the identity scores themselves. For this reason, the user will also have the option to use the classifier only rather than the classifier followed by the regression model (James *et al.*, 2018; James and Girgis, 2018). In other situations, the value of the threshold may not have a biological meaning; thus, the user may select to use regression without the preceding classification step (Velasco II *et al.*, 2018). Next, we explain how the semi-synthetic data are generated.

### Semi-synthetic data generation

Semi-synthetic data are generated by mutating real sequences taken from the input database. This data are then mutated using the following mutation types: (i) single-point mutation or (ii) block mutation. In single-point mutation, a single nucleotide is mismatched, deleted, or inserted. In block mutation, a block of random nucleotides is inserted; or a block of consecutive nucleotides is deleted; or a block of nucleotides is duplicated and placed in tandem to the original block. The size of the block is chosen at random at a minimum of 2 and a maximum of 50 nucleotides. To ensure that the original nucleotide composition is conserved, random nucleotides to be inserted or to be changed are generated from the same distribution of nucleotides in the original sequence. For example, suppose that the original sequence has the following nucleotide distribution: A: 0.4, C: 0.1, G: 0.1, T: 0.4. When a random nucleotide to be inserted, A or T has the highest probability of 0.4 each and C or G has the lowest probability of 0.1 each. If the single-point and the block mutation models are applied together, the ratio of each is determined randomly for each mutated sequence.

We invented this generative process to avoid using alignment algorithms for three reasons. First, alignment algorithms are slow. Second, the training dataset may not have enough sequence pairs with specific identity scores to train the classifier or the regression model. Third, alignment algorithms are almost infeasible on very long sequences.

Because mutated sequences are generated with specific mutation types and rates, the identity scores can be calculated without using any alignment algorithms. To calculate an identity score, we need to know the length of the alignment and the number of matches — identical nucleotides — between two sequences. Each mutation type affects these two numbers in a unique way. If we keep track of the mutations applied and update the alignment length and the number of matches accordingly, the corresponding identity score can be obtained without actually aligning the two sequences. For a very simple example, consider a 10-nucleotides-long sequence. We wish to mutate 30% of this sequence. For simplicity, assume that the three mutations are mismatch, insertion, and deletion. Initially, the length of the alignment and the number of matches are equal to 10 — the length of the original sequence. A mismatch does not affect the length of the alignment; however, it decreases the number of matches. After this mismatch, the length of the alignment is 10, and the number of matches is 9. An insertion results in a gap in the original sequence if the two sequences were to be aligned versus each other, i.e. it increases the length of the alignment by 1 and does not affect the number of matches. After this insertion, the length of the alignment is 11, and the number of matches is 9. Deleting a nucleotide results in a gap in the mutated sequence when it is aligned versus the original sequence. This gap does not affect the alignment length; but the number of matches is decreased by 1. After this deletion, the length of the alignment is 11, and the number of matches is 8. These three mutations lead to an identity score of 0.73. Table 1 lists the mutation types used in our study and their effects on the alignment length and the number of matches. This procedure is applied to generating two datasets discussed next.

**Table 1.** The effects of each mutation type on the identity score. The identity score is the ratio of identical nucleotides between two sequences to the total length of the alignment, which may include gaps. Initially, the alignment length and the number of matches are equal to the length of the original sequence that is the template to be mutated. As the mutation process proceeds, a loaded coin is flipped to decide whether a single nucleotide or a block of nucleotides will be mutated. Next, a mutation sub-type — e.g. mismatch, insertion, or deletion if single-point mutation — is selected randomly. Each mutation type affects the alignment length and the number of matches in a unique way. For example, a mismatch has no effect on the length of the alignment; it decreases the number of matches by 1.

| Mutation Type | Alignment Length | Number of Matches |
| --- | --- | --- |
| Mismatch | No effect | Subtract 1 |
| Deletion | No effect | Subtract the number of the deleted nucleotide(s) |
| Insertion | Add the number of the inserted nucleotide(s) | No effect |
| Duplication | Add the number of the duplicated nucleotides | No effect |

### Training and testing sets

When a database of DNA sequences is given, the first step is to sample 300 sequences to generate semi-synthetic data (4500 original-mutated sequence pairs). The number of sequence pairs with identity scores above the threshold is half of the number of pairs with identity scores below the threshold. Specifically, for every original sequence, 5 sequences above the threshold and 10 below the threshold are generated in order to more accurately match real distributions of alignment identities. If the user choose not to provide an identity threshold, sequence pairs with identity scores are generated randomly between 35% and 100%. To ensure that the model is not biased towards a particular segment in the identity scores, each 5% segment of the identity scores is equally represented. Balancing the pairs with identity scores above the threshold is done separately from balancing those with identity scores below the threshold because of the different sizes of these two sets. When a threshold is not provided, the entire dataset is balanced together. Finally, the dataset is divided into two mutually exclusive sets — the training set and the testing set. Next, we illustrate how these datasets are represented to the classifier and the regression model as few statistics calculated on pairs of k-mer histograms.

### Calculating the k-mer statistics

Each sequence is represented as a k-mer histogram. Then statistics are calculated on each pair of histograms. The choice of *k* — the size of k-mers — guarantees that the histogram size is linear with respect to an average input sequence. We calculate *k* according to Equation 1 (Luczak *et al*., 2017; James *et al*., 2018; James and Girgis, 2018).

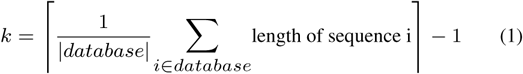

Using our survey of alignment-free methods as a resource (Luczak *et al.*, 2017), we chose the following statistics: Earth Mover’s Distance, Euclidean, Intersection, Kulczynski_2_, Length Difference, Manhattan, Normalized Vectors, Pearson Correlation Coefficient, and Similarity Ratio. These statistics are chosen because they are much faster while still maintaining strong predictive power. We compute these 9 statistics then normalize each of them between 0 and 1. Some statistics represent distances and others represent similarities. We convert each distance to a similarity by subtracting the normalized distance from 1. We call these 9 statistics single statistics. One of the primary results of our evaluation study was that squared versions or multiplicative combinations can often times outperform single statistics. For this reason, we square each of the single statistics to create 9 additional statistics. Finally, the paired statistics are generated by multiplying each unique combination of the 18 single and squared statistics. These statistics are the features, on which the self-supervised GLMs are trained. Next, we illustrate GLMs briefly.

### GLMs

The general form of the linear model is ***y*** = ***F w*** where ***y*** is the target we wish to predict. For classification, ***y*** is a vector of 1’s (the sequence pair has similarity above the threshold) and 0’s (the sequence pair has similarity below the threshold). For regression, ***y*** represents the identity scores. ***F*** is the feature matrix; each of its columns represents a particular statistic except the first column is all ones. The coefficients in the ***w*** vector are found using the pseudoinverse solution (Equation 2).

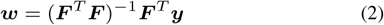

Now that we have the coefficients of the GLM, Equation 3 is used for making predictions.

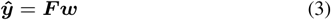

Here, *ŷ* represents the predicted labels, above or below the identity threshold, or the predicted identity score for a given sequence pair. The output of the GLM — *ŷ* — is processed further. For classification, Equation 4 is used for assigning a label to a pair of sequences (1: above the threshold and 0: below the threshold).

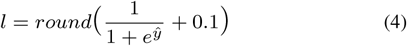

Here, *l* is the final predicted label and *ŷ* is the row output of the GLM for one sequence pair. According to this equation a sequence pair with a value of 0.6 or greater is classified as having an identity score above the threshold. We used 0.6 instead of 0.5 to compensate for training on semi-synthetic data — not on completely real sequences. For regression, if *ŷ* is greater than 1 (less than 0), it is set to 1 (0).

Using a small number of features is necessary to the success of a predictive model because it prevents the learning algorithm from overfitting the training data. Therefore, we utilize a greedy featureselection procedure, which depends on GLMs. In the next step, we discuss how to select the best four or five features, i.e. statistics. a predictive model because it prevents the learning algorithm from overfitting the training data. Therefore, we utilize a greedy feature-selection procedure, which depends on GLMs. In the next step, we discuss how to select the best four or five features, i.e. statistics.

### Greedy feature selection

Features are selected automatically on each input dataset. A greedy algorithm is used for selecting a strong group of features without trying every possible combination. FASTCAR utilizes a GLM in selecting features that maximize the testing accuracy for classification or minimize the mean absolute error for regression. Two datasets are utilized in this step: (i) the training set is used for training the GLM and (ii) the testing set is used for measuring the accuracy or the mean absolute error. We assess the performance on the testing set to guard against over-fitting.

First, we discuss how to select features for the classifier; this algorithm is based on forward stepwise selection, which is often used in statistical learning. The algorithm selects 4 or 5 features. In each iteration of the algorithm, the classifier is trained on the training set to recognize if a pair of sequences have an identity score above or below a given threshold. The testing accuracy is the number of correct classifications divided by the number of sequence pairs in the testing set. The first step involves training the classifier using one feature on the training set, then evaluating the classifier due to this feature on the testing set to find the best performing feature. Once found, it is added to the best-features set and is excluded in the next iterations of the algorithm. After that, the tool goes through the remaining features one by one, attempting to combine each of these features with the best performing feature(s) found in the previous iteration(s). Once this step is finished, the feature that results in the best testing accuracy is added to the best-features set and is excluded in the subsequent iterations. After selecting the minimum number of features — 4 — the fifth feature is added if it improves the testing accuracy. If no improvement is attained or five features are selected, the algorithm stops. Our research has shown that adding additional features beyond this point does not increase the accuracy enough to warrant a speed reduction (James *et al.*, 2018). The end result is a set of 4 or 5 features that have very high testing accuracy. A similar process is used for selecting the best 4 or 5 features for the regression model. The performance is measured according to the mean absolute error, i.e. the average absolute difference between the true identity score and the predicted one — the lower, the better.

Up to this point, we discussed how the training and the testing datasets are generated and how the features are extracted and selected. Next, we discuss three modes, in which the trained models can be applied.

### Prediction modes

The classifier determines whether a sequence pair falls above or below the identity score threshold, which is provided by the user. Pairs classified below the threshold are removed; the remaining sequence pairs are sent to the regression model to predict their identity scores. Alternatively, there are cases where classification only is desired (James *et al.*, 2018; James and Girgis, 2018); it is possible to disable regression. If identity scores throughout the entire range of sequences are desired, regression alone can be performed, not excluding any sequence pairs (Velasco II *et al.*, 2018). At this point, the description of FASTCAR’s method is complete.

### Executing FASTCAR and the related tools

We chose 2 widely-used tools, USEARCH (Edgar, 2010) and BLAST (Altschul *et al.*, 1990), to compare to FASTCAR. Ground truth sets, on which the three tools can be evaluated, were assembled. For this purpose, we chose needleall from EMBOSS (Rice *et al.*, 2000) because of its ability to do all-versus-all global alignments. The program needleall does not use heuristics like the ones used in the other two tools; it is much slower than USEARCH and BLAST.

While BLAST is designed for local alignment, it also generates global alignments, as they are a special case of local alignments. It is possible to get global alignment scores by manipulating parameters and filtering out BLAST results. To obtain the global identity score, we multiply the identity score due to local alignment by the query coverage and by the subject coverage (Equation 5).

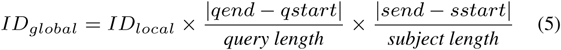

Here, *ID_global_* and *ID_local_* are BLAST’s global and local identity scores; qstart and qend are the start and the end of the aligned region in the query sequence; sstart and send are the start and the end of the aligned region in the subject sequence. Table 2 shows few examples of local identity scores and their corresponding global scores due to Equation 5 and the global alignment algorithm. The adjusted scores produced by Equation 5 are very close to the scores calculated by the global alignment algorithm. To get BLAST to print many alignments, the parameter “num_alignments” can coax BLAST into printing out more alignments (1000000) than just the few best local alignments. The maximum adjusted alignment score of several alignments between the same sequence pair is considered as the global identity score.

**Table 2.**
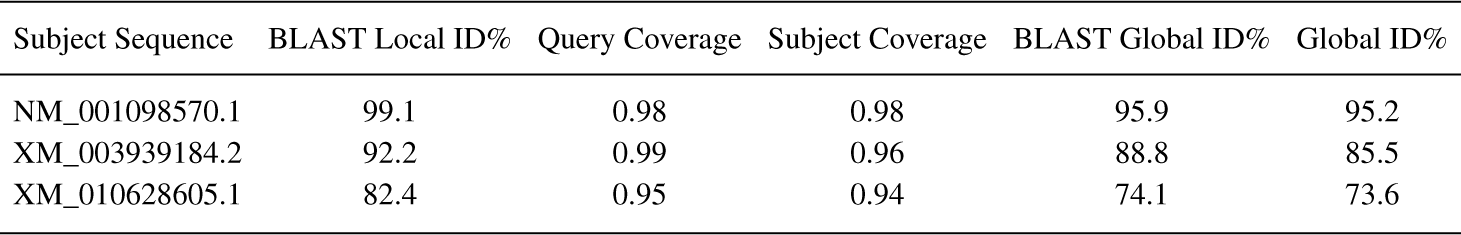
Conversion from BLAST’s local identity scores to global identity scores. These samples come from the Keratin dataset. The query sequence is NM_002283.3. Equation 5 is applied to BLAST’s local identity score, the query coverage, and the subject coverage to produce the corresponding global identity score, which is very similar to the one produced by the global alignment algorithm (Global ID).

| Subject Sequence | BLAST Local ID% | Query Coverage | Subject Coverage | BLAST Global ID% | Global ID% |
| --- | --- | --- | --- | --- | --- |
| NM_001098570.1 | 99.1 | 0.98 | 0.98 | 95.9 | 95.2 |
| XM_003939184.2 | 92.2 | 0.99 | 0.96 | 88.8 | 85.5 |
| XM_010628605.1 | 82.4 | 0.95 | 0.94 | 74.1 | 73.6 |

Additionally, we even created a multi-process module around both BLAST and USEARCH to allow it to use 20 threads instead of the maximum, 8 and 1 threads, respectively, to asses the speed-up due to the linear algorithm of FASTCAR. Thus, we divided the database sequences into 20 files, each of which was scanned using BLAST or USEARCH. This means for BLAST, makeblastdb was run on each of the 20 files once. The parameters used for BLAST were “-task blastn -strand plus -perc_identity $cutoff -num_threads 1 -num_alignments 1000000 -reward 1 -penalty -1 - gapopen 2 -gapextend 1”, where $cutoff is the alignment identity threshold provided by the user. This command was executed in parallel on 20 cores using the Unix Parallel utility. For USEARCH, the parameters were “-search_global -strand plus -id $cutoff -threads 1 -blast6out”. Similarly, this command was executed in parallel on 20 cores. FASTCAR was run in its default mode, using classification followed by regression, using 20 threads. All tools were run on the same computer, Dell Precision Tower 5810, 10-core Xeon E5-2630 CPU, and 32 GB RAM.

We have just finished discussing how FASTCAR and the related tools were executed. Next, we explain how we constructed 3 datasets, on which FASTCAR, USEARCH, and BLAST were evaluated.

### Datasets

Three datasets were used in evaluating the tools. The first set is the Keratin set, which consists of 5220536 sequences with sequence lengths ranging between 1353 base pairs (bp) and 3250 bp. The second set — the P27 — consists of 7990947 sequences between 1500 bp and 4000 bp. The third set consists of 3613 bacterial genomes; the lengths of these genomes range from 112,031 bp to 14,782,125 bp. This dataset comes from the Ensembl Bacteria release 40 (Zerbino *et al.*, 2017). All genomes containing only one contig were selected. Statistics about these sets are displayed in Table 3.

**Table 3.** Statistics of the datasets used in evaluating FASTCAR and the related tools.

| Dataset | Sequence Count | Nucleotide Count | Maximum Length | Minimum Length | Mean | Median |
| --- | --- | --- | --- | --- | --- | --- |
| Keratin | 5220536 | 12517803991 | 3250 | 1353 | 2398 | 2169 |
| P27 | 7990947 | 19919189291 | 4000 | 1500 | 2493 | 3412 |
| Bacterial | 3613 | 12412294439 | 14782125 | 112031 | 3435454 | 1607556 |

Searching a sequence database for similar sequences to a query sequence is an important and common application of alignment tools and the proposed tool. We utilized two datasets as databases to be searched. Query sequences were selected for the Keratin and the P27 sets. The Keratin query sequence is the NM_002283.3 — *Homo sapiens* keratin 85 (KRT85), transcript variant 1, mRNA sequence. The P27 query sequence is the NM_004064.4 — *Homo sapiens* cyclin dependent kinase inhibitor 1B. Both the P27 and the Keratin ground truth sets, i.e. similar sequences to the query sequence, were found by searching the NCBI database (O’Leary *et al.*, 2016). The search parameters for Keratin were: “srcdb_refseq[PROP] AND Keratin NOT Homo Sapiens”. This search was restricted to animal sequences between 1700 bp and 3250 bp in length. When gathered on 26 June 2018, this query resulted in 6669 sequences. To find similar sequences to the P27 query, we searched: “srcdb_refseq[PROP] AND cyclin dependent kinase inhibitor 1B NOT Homo Sapiens”. This search was restricted to animal sequences 2000– 3000 bp long, resulting in 131 sequences when gathered on 26 June 2018. After that, sequences that have less than 70% identity with the query sequence was removed from the ground truth sets, resulting in 56 and 66 sequences similar to the Keratin and the P27 query sequences.

We utilized the bacterial set in a hierarchical clustering application requiring all-versus-all identity scores; therefore, no query or ground truth set are needed. Additionally, the parameters passed to FASTCAR included a 60% identity threshold and 150 as the number of samples — the default is 300 — to reduce the time required to generate the semi-synthetic sequences. Hierarchical clustering was carried out using the ward2 algorithm in the R function hclust. The tree was generated using the ete3 package (http://etetoolkit.org/) in Python.

Up to here, we described the computational principles behind FASTCAR. Then the details of the related tools and the evaluation datasets were illustrated. After that, the performances of the 3 tools on the 3 datasets are reported and discussed.

## 3 Results

### Evaluation measures

We evaluated the 3 tools using the following 6 measures:
1. **Sensitivity:** Sensitivity is the rate of True Positives (TP) to the combined TP and false negatives. It measures the ability of a tool to identify TP is a large dataset — the more TP found, the better.
2. **Precision:** Precision is the ratio of TP to TP and false positives (FP). Precision measures the relevancy of returned results, since it rates TP to the total predicted positive labels. This measure is very important when experimental validations of the results are considered.
3. **F-measure:** F-measure combines sensitivity and precision by taking the harmonic mean between them (Equation 6).

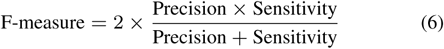
4. **Mean absolute error:** This measure is used in regression analysis to measure how close, on average, the predicted value is to the actual value. The mean absolute error is the average absolute difference between the predicted value and the actual value. Using this metric allows a comparable benchmark for the error in estimating the identity scores in relation to the ones due to global alignment algorithms. Additionally, it can be used as an expected margin of error.
5. **Time:** Time reported is the wall clock time, as multi-threaded applications are best estimated using real time.
6. **Memory:** Max memory used by a tool is measured, as the memory requirement is set by the maximum amount.

We report sensitivity, precision, F-measure, and mean absolute error as percentages. The time is measured in seconds and the memory requirements in Gigabytes. Next, we discuss the performances of the tools.

### Evaluations on large-scale datasets

Using the 2 datasets described earlier, FASTCAR, BLAST, and USEARCH were evaluated by searching for one query sequence in a database of 5–8 millions of sequences (Table 4).

**Table 4.**
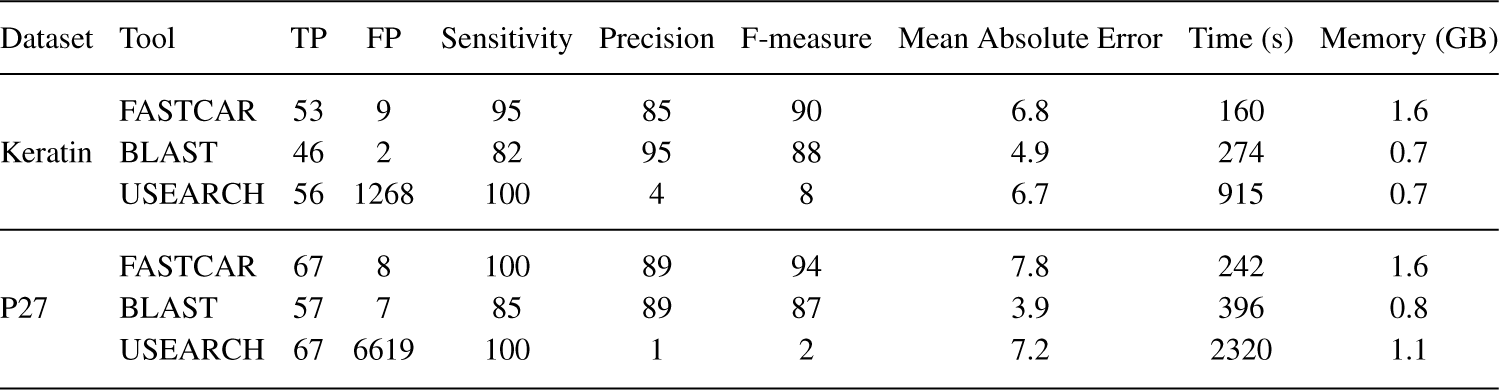
Evaluations of FASTCAR, BLAST, and USEARCH. We tested the tools’ abilities to search large databases for a query sequence. Alignment scores were generated with needleall, which is a tool for global alignment. These evaluations were conducted on two datasets: Keratin and P27. The Keratin dataset has 5,220,536 sequences, ranging between 1353 bp and 3250 bp. There is only one query sequence for the Keratin dataset. We searched for sequences that at least 70% identical to the query sequence, resulting in a ground truth set consisting of 56 sequences. The P27 set consists of 7,990,947 sequences between 1500 bp and 4000 bp. Similar to the Keratin set, there is one query sequence. Sequences with at least 70% identity to the Keratin query sequence — 67 sequences — are considered True Positives (TP). Sequences detected by a tool and do not belong to the ground truth set are considered False Positives (FP). Sensitivity and precision are reported as percentages. F-measure could be thought of as percentage. The mean absolute error is displayed as percentage; it is measured on the TP only. The time is reported in seconds (s), and the memory requirement in Gigabytes (GB).

| Dataset | Tool | TP | FP | Sensitivity | Precision | F-measure | Mean Absolute Error | Time (s) | Memory (GB) |
| --- | --- | --- | --- | --- | --- | --- | --- | --- | --- |
| Keratin | FASTCAR | 53 | 9 | 95 | 85 | 90 | 6.8 | 160 | 1.6 |
|  | BLAST | 46 | 2 | 82 | 95 | 88 | 4.9 | 274 | 0.7 |
|  | USEARCH | 56 | 1268 | 100 | 4 | 8 | 6.7 | 915 | 0.7 |
| P27 | FASTCAR | 67 | 8 | 100 | 89 | 94 | 7.8 | 242 | 1.6 |
|  | BLAST | 57 | 7 | 85 | 89 | 87 | 3.9 | 396 | 0.8 |
|  | USEARCH | 67 | 6619 | 100 | 1 | 2 | 7.2 | 2320 | 1.1 |

We start by looking at the sensitivities of the 3 tools. On the Keratin set, USEARCH was the most sensitive tool (100%), followed by FASTCAR (95%), followed by BLAST (82%). On the P27 set, USEARCH and FASTCAR achieved perfect sensitivities of 100%, whereas BLAST achieved 85%. The number of FP is quite important because of the large number of sequences to be searched. BLAST had the lowest number of FP on the Keratin and the P27 sets (2 and 7), followed by FASTCAR (9 and 8). USEARCH found a large number of FP on these two datasets (1268 and 6619). These numbers represent very small percentages of the databases (0.02% – 0.08%), but they are much larger than the numbers of similar sequences in each set (56 and 67). For this reason, we discuss the precision metric next. BLAST was the most precise tool (95% and 89%) on the two sets, followed by FASTCAR (85% and 89%). Because of the large numbers of FP detected by USEARCH, it was the least precise tool (4% and 1%). On the two datasets, FASTCAR had the highest F-measure. On the Keratin dataset, FASTCAR’s F-measure was better than BLAST’s (90 versus 88) and much better than USEARCH’s (90 versus 8). Similar results were obtained on the P27 dataset (FASTCAR: 94, BLAST: 87, and USEARCH: 2). In addition, we evaluated how close the identity scores due to the three tools to those calculated by the global alignment algorithm. BLAST had the lowest error (4–5%). FASTCAR and USEARCH had comparable errors of 7%–8%. Keep in mind that these errors were calculated on the TP only. One of the main advantages of the proposed method is its speed. FASTCAR was the fastest tool on the two sets. It required about 40% less time than BLAST. FASTCAR was faster than USEARCH by 6–10 times. Regarding memory, FASTCAR required more memory than BLAST and USEARCH. However, the max amount of memory used — 1.6 GB — is readily available on average personal computers. Moreover, once the classifier and the regression model are trained, memory is kept low as sequences are processed as they are read.

When it comes to searching large databases, USEARCH is the most sensitive tool, whereas BLAST is the most precise tool. FASTCAR provides the best compromise between sensitivity and precision while being the fastest tool. Next, we report the results on another application.

### Evaluations on long sequences

In this experiment, we show that it is possible to use entire genomes and their identity scores to provide hierarchical clusterings/phylogenetic trees with high accuracy. Comparing long sequences — millions of nucleotides long — using alignment algorithms require prohibitive long time. Because the bacterial genomes are too long (3.4 mega bp on average) to be aligned with the other methods feasibly, we were able to apply FASTCAR only. Often times, all-versus-all comparisons can generate useful information such as phylogenetic trees. To generate these all-versus-all comparisons, an external script was run (Supplementary Data 2). This script generated the upper triangular matrix, which is the input to the hierarchical clustering algorithm. The clustering produced match most subspecies within the same cluster. Additionally, the same is true for matching species, as well as extending out to higher taxonomies such as genus. The full tree is provided as Supplementary Data 3. A subset of that tree is shown in Fig. 3, which displays all of *Listeria* and *Pseudomonas* as complete subtrees not mixed with other genus. For another example, the species *L. monocytogenes* is all in one cluster, as well as *L. ivanovii*, *P. protegens*, and *P. chlororaphis*.

In sum, aligning long sequences such as bacterial genomes cannot be accomplished by any of the currently available alignment-based tools. Our results demonstrate that FASTCAR is the first tool capable of assessing the similarity between very long DNA sequences via identity scores.

## 4 Discussion

In this section we discuss few points. We start by giving the rational of using GLMs as the core of FASTCAR. Then we analyze its time and memory requirements. After that we give the details of similar tools that were not evaluated. Finally, we outline some directions for future research.

### Rationale of choosing GLMs

A number of machine learning algorithms for classification and regression are available. These algorithms include Support Vector Machines (SVMs) and Artificial Neural Networks (ANNs). We are attracted to GLMs because they are parameter-free models, which are well suited to the idea of adaptive training. We chose GLMs primarily because of their time efficiency, which is due to the absence of parameters to be optimized. Both the SVMs and the ANNs can be highly accurate if given enough time to train. However, they require several variables that need to be optimized which does not fit well into our adaptive training idea. On the other hand, GLMs only require calculating the pseudo-inverse solution to find the linear coefficients. This operation is much cheaper than searching for optimal parameters required by the other algorithms. Our experiments show that the hierarchical GLM can obtain comparable results to SVMs and ANNs — without parameter optimization however.

### Runtime and space analysis

The algorithm behind FASTCAR is a linear algorithm. We start by analyzing the time required by the training stage. Then we analyze the time required by the prediction stage. First, the size of k-mers and the histogram data type are determined in constant time by scanning a fixed number of sequences — 10000. The training stage involves (i) generating semi-synthetic sequences, (ii) selecting features, and (iii) training two final models. Generating a fixed number of semi-synthetic sequences takes constant time. There are 162 features, from which 5 features at most are selected using the greedy selection algorithm. Therefore, 800 GLMs are trained as classifiers and 800 GLMs are trained as regression models. Because the number of the features is fixed, this operation takes constant time too. Training the final models are done while selecting the features. Thus, the entire training process takes constant time. Next, we discuss the time requirement for the prediction stage. The training process results in at most 6 parameters for the classifier and 6 parameters for the regression model because the maximum number of features is 5 and each model uses an additional parameter representing bias. To predict the identity score of a pair of sequences, the two sequences are read (linear time). Then the histograms are generated (linear time). Assuming that each of the 10 features are unique and each is a multiplicative combination of two features, the algorithm scans the two histograms 20 times in the worst case scenario. Because the histogram size is guaranteed to be linear with respect to an average sequence in the database, calculating the features is done in linear time. In sum, the identity score of a pair of sequences is produced in linear time.

With regard to the space requirement, the training stage requires loading 10000 sequences in memory. Once these are read, 300 of them are selected and the rest are discarded. Then 4500 semi-synthetic sequences are generated and kept in memory. Therefore, the entire training process requires constant space to store a fixed number of sequences. To predict the pair-wise identity score, the space requirement is linear because only two histograms representing the two sequences are needed.

**Fig. 1.**
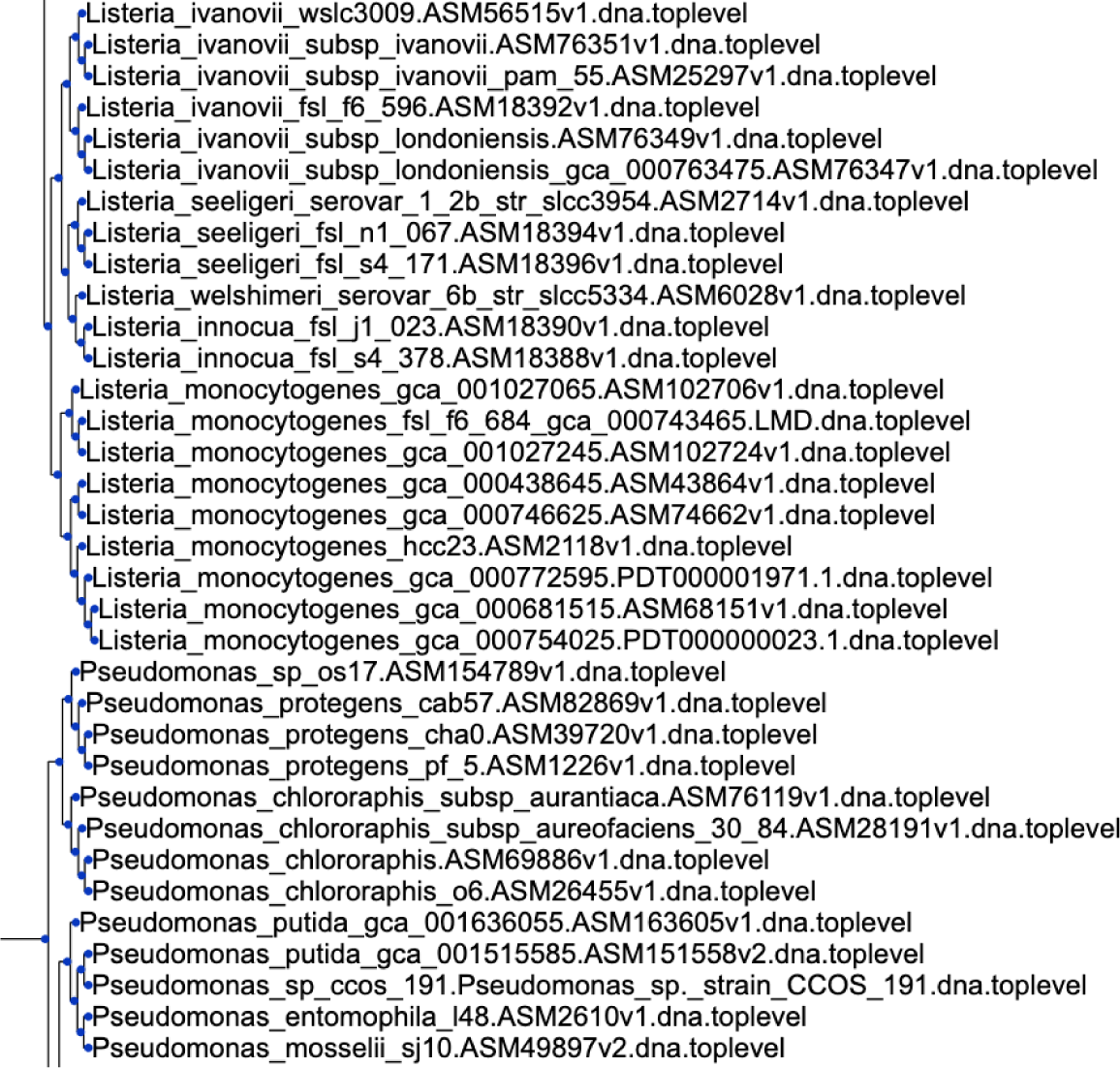
FASTCAR was applied to about 3600 bacterial genomes to generate all-versus-all identity scores, which were used for building a phylogenetic tree using hierarchical clustering. Here, we show only a part of the tree. All of Listeria and Pseudomonas are in complete subtrees with no other genus. The species L. monocytogenes is all in one cluster, as well as L. ivanovii, P. protegens, and P. chlororaphis. Even the subspecies londoniensis and ivanovii are distinctly clustered.

### Other methods

MASH (Ondov *et al.*, 2016) is a program, which can accurately estimate genomic distances using MinHash and Jaccard similarity to estimate pair-wise similarity. However, MASH does not report the identity scores, instead opting for a novel distance metric based on Jaccard similarity, which the authors of MASH argue may correlate with other similarity metrics such as average nucleotide identity. We could not evaluate MASH because it does not output identity scores. Local alignment tools (other than BLAST) were not compared. Tools that are purely computational improvements such as SWIPE (a SIMD-parallel optimized version of BLAST) (Rognes, 2011) were not considered since the results other than time should be very similar to BLAST. Our rationale is that these improvements — using specialized instructions or hardware — could be applied to other tools (such as FASTCAR and USEARCH) to similarly speed up these tools with no change in output. In this way, our comparisons are focused on highlighting improvements due to algorithmic advancements. We could not compare to CaBLAST (Loh *et al.*, 2012); although novel, the currently available proof-of-concept is too slow. CaBLAST applies BLAST to compressed representations of the sequences rather than to the original sequences themselves.

## 5 Conclusion

A very important algorithm in bioinformatics, pairwise alignment, is slow. Fast alternatives such as k-mer distances produce scores that do not have relevant biological meanings as the identity scores produced by alignment algorithms. We developed a novel software tools, FASTCAR, for estimating identity scores of DNA sequence pairs. On an input database, FASTCAR trains a self-supervised classifier and a self-supervised regression model to predict identity scores using few, efficient, k-mer statistics. Training these models is done with a novel method of generating sequences with known identity scores, allowing for alignment-free prediction of alignment identity scores. This is the first time identity scores are obtained in linear time using linear space.

## Acknowledgements

The authors would like to thank Alexander Baumgartner for his help with data processing and coding the module for generating mutated sequences.

## References

Almeida, J. S. and Vinga, S. (2002). Universal sequence map (USM) of arbitrary discrete sequences. BMC Bioinformatics, 3(1), 6.

Altschul, S. F., Gish, W., Miller, W., Myers, E. W., and Lipman, D. J. (1990). Basic local alignment search tool. J Mol Biol, 215(3), 403–410.

Blaisdell, B. E. (1986). A measure of the similarity of sets of sequences not requiring sequence alignment. Proc Natl Acad Sci U S A, 83(14), 5155–5159.

Bonham-Carter, O., Steele, J., and Bastola, D. (2014). Alignment-free genetic sequence comparisons: a review of recent approaches by word analysis. Brief Bioinform, 15(6), 890.

Borozan, I., Watt, S., and Ferretti, V. (2015). Integrating alignment-based and alignment-free sequence similarity measures for biological sequence classification. Bioinformatics, 31(9), 1396–1404.

Butler, J., MacCallum, I., Kleber, M., Shlyakhter, I. A., Belmonte, M. K., Lander, E. S., Nusbaum, C., and Jaffe, D. B. (2008). ALLPATHS: de novo assembly of whole-genome shotgun microreads. Genome Res, 18(5), 810–820.

Carradec, Q., Pelletier, E., Da Silva, C., Alberti, A., Seeleuthner, Y., Blanc-Mathieu, R., Lima-Mendez, G., Rocha, F., Tirichine, L., Labadie, K., Kirilovsky, A., Bertrand, A., Engelen, S., Madoui, M.-A., Méheust, R., Poulain, J., Romac, S., Richter, D. J., Yoshikawa, G., Dimier, C., Kandels-Lewis, S., Picheral, M., Searson, S., Jaillon, O., Aury, J.-M., Karsenti, E., Sullivan, M. B., Sunagawa, S., Bork, P., Not, F., Hingamp, P., Raes, J., Guidi, L., Ogata, H., de Vargas, C., Iudicone, D., Bowler, C., and Wincker, P. (2018). A global ocean atlas of eukaryotic genes. Nat Commun, 9(1), 373.

Chattopadhyay, A. K., Nasiev, D., and Flower, D. R. (2015). A statistical physics perspective on alignment-independent protein sequence comparison. Bioinformatics, 31(15), 2469–2474.

Costa, A. M., Machado, J. T., and Quelhas, M. D. (2011). Histogram-based DNA analysis for the visualization of chromosome, genome and species information. Bioinformatics, 27(9), 1207.

Costello, E. K., Lauber, C. L., Hamady, M., Fierer, N., Gordon, J. I., and Knight, R. (2009). Bacterial community variation in human body habitats across space and time. Science, 326(5960), 1694–1697.

Dai, Q., Yang, Y., and Wang, T. (2008). Markov model plus k-word distributions: a synergy that produces novel statistical measures for sequence comparison. Bioinformatics, 24(20), 2296.

Eaton, D. A. R. (2014). Pyrad: assembly of de novo radseq loci for phylogenetic analyses. Bioinformatics, 30(13), 1844–1849.

Edgar, R. C. (2010). Search and clustering orders of magnitude faster than BLAST. Bioinformatics, 26(19), 2460–2461.

Ghandi, M., Lee, D., Mohammad-Noori, M., and Beer, M. A. (2014). Enhanced regulatory sequence prediction using gapped k-mer features. PLoS Comput Biol, 10(12), e1004035.

Girgis, H. Z. (2008). Machine-learning-based meta approaches to protein structure prediction. Ph.D. thesis, The State University of New York at Buffalo.

Girgis, H. Z. (2015). Red: an intelligent, rapid, accurate tool for detecting repeats de-novo on the genomic scale. BMC Bioinformatics, 16(1).

Girgis, H. Z. and Corso, J. J. (2008). Stp: the sample-train-predict algorithm and its application to protein structure meta-selection. Technical Report 16, The State University of New York at Buffalo.

Girgis, H. Z. and Ovcharenko, I. (2012). Predicting tissue specific cis-regulatory modules in the human genome using pairs of co-occurring motifs. BMC Bioinformatics, 13(1), 25.

Girgis, H. Z. and Sheetlin, S. L. (2013). MsDetector: toward a standard computational tool for DNA microsatellites detection. Nucleic Acids Res, 41(1), e22.

Girgis, H. Z., Corso, J. J., and Fischer, D. (2009). On-line hierarchy of general linear models for selecting and ranking the best predicted protein structures. In Conf Proc IEEE Eng Med Biol Soc, pages 4949–4953.

Göke, J., Schulz, M. H., Lasserre, J., and Vingron, M. (2012). Estimation of pairwise sequence similarity of mammalian enhancers with word neighbourhood counts. Bioinformatics, 28(5), 656.

Gotoh, O. (1982). An improved algorithm for matching biological sequences. J Mol Biol, 162, 705–708.

Haubold, B. (2014). Alignment-free phylogenetics and population genetics. Brief Bioinform, 15(3), 407.

James, B. T. and Girgis, H. Z. (2018). MeShClust2: Application of alignment-free identity scores in clustering long DNA sequences. BioRxiv, page 451278.

James, B. T., Luczak, B. B., and Girgis, H. Z. (2018). MeShClust: an intelligent tool for clustering DNA sequences. Nucleic Acids Res, page gky315.

Kantorovitz, M. R., Robinson, G. E., and Sinha, S. (2007). A statistical method for alignment-free comparison of regulatory sequences. Bioinformatics, 23(13), i249.

Korf, I. (2004). Gene finding in novel genomes. BMC Bioinformatics, 5(1), 59.

Leimeister, C.-A., Boden, M., Horwege, S., Lindner, S., and Morgenstern, B. (2014). Fast alignment-free sequence comparison using spaced-word frequencies. Bioinformatics, 30(14), 1991.

Liao, W., Ren, J., Wang, K., Wang, S., Zeng, F., Wang, Y., and Sun, F. (2016). Alignment-free transcriptomic and metatranscriptomic comparison using sequencing signatures with variable length markov chains. Sci Rep, 6(37243).

Lippert, R. A., Huang, H., and Waterman, M. S. (2002). Distributional regimes for the number of k-word matches between two random sequences. Proc Natl Acad Sci U S A, 99(22), 13980–13989.

Liu, X., Wan, L., Li, J., Reinert, G., Waterman, M. S., and Sun, F. (2011). New powerful statistics for alignment-free sequence comparison under a pattern transfer model. J Theor Biol, 284(1), 106–116.

Loh, P.-R., Baym, M., and Berger, B. (2012). Compressive genomics. Nat Biotechnol, 30(7), 627.

Luczak, B. B., James, B. T., and Girgis, H. Z. (2017). A survey and evaluations of histogram-based statistics in alignment-free sequence comparison. Brief Bioinform, page bbx161.

Luo, R., Liu, B., Xie, Y., Li, Z., Huang, W., Yuan, J., He, G., Chen, Y., Pan, Q., Liu, Y., Tang, J., Wu, G., Zhang, H., Shi, Y., Liu, Y., Yu, C., Wang, B., Lu, Y., Han, C., Cheung, D. W., Yiu, S.-M., Peng, S., Xiaoqian, Z., Liu, G., Liao, X., Li, Y., Yang, H., Wang, J., Lam, T.-W., and Wang, J. (2012). SOAPdenovo2: an empirically improved memory-efficient short-read de novo assembler. Gigascience, 1(1), 1–6.

Needleman, S. B. and Wunsch, C. D. (1970). A general method applicable to the search for similarities in the amino acid sequence of two proteins. J Mol Biol, 48, 443–453.

O’Leary, N. A., Wright, M. W., Brister, J. R., Ciufo, S., Haddad, D., McVeigh, R., Rajput, B., Robbertse, B., Smith-White, B., Ako-Adjei, D., Astashyn, A., Badretdin, A., Bao, Y., Blinkova, O., Brover, V., Chetvernin, V., Choi, J., Cox, E., Ermolaeva, O., Farrell, C. M., Goldfarb, T., Gupta, T., Haft, D., Hatcher, E., Hlavina, W., Joardar, V. S., Kodali, V. K., Li, W., Maglott, D., Masterson, P., McGarvey, K. M., Murphy, M. R., O’Neill, K., Pujar, S., Rangwala, S. H., Rausch, D., Riddick, L. D., Schoch, C., Shkeda, A., Storz, S. S., Sun, H., Thibaud-Nissen, F., Tolstoy, I., Tully, R. E., Vatsan, A. R., Wallin, C., Webb, D., Wu, W., Landrum, M. J., Kimchi, A., Tatusova, T., DiCuccio, M., Kitts, P., Murphy, T. D., and Pruitt, K. D. (2016). Reference sequence (RefSeq) database at NCBI: current status, taxonomic expansion, and functional annotation. Nucleic Acids Res, 44(D1), D733–D745.

Ondov, B. D., Treangen, T. J., Melsted, P., Mallonee, A. B., Bergman, N. H., Koren, S., and Phillippy, A. M. (2016). Mash: fast genome and metagenome distance estimation using minhash. Genome Biol, 17(1), 132.

Peled, S., Leiderman, O., Charar, R., Efroni, G., Shav-Tal, Y., and Ofran, Y. (2016). De-novo protein function prediction using dna binding and rna binding proteins as a test case. Nat Commun, 7(13424).

Pham, T. D. and Zuegg, J. (2004). A probabilistic measure for alignment-free sequence comparison. Bioinformatics, 20(18), 3455.

Pinello, L., Lo Bosco, G., and Yuan, G.-C. (2014). Applications of alignment-free methods in epigenomics. Brief Bioinform, 15(3), 419.

Reinert, G., Chew, D., Sun, F., and Waterman, M. S. (2010). Alignment-free sequence comparison (i): Statistics and power. J Comput Biol, 16(12), 1615–1634.

Ren, J., Song, K., Sun, F., Deng, M., and Reinert, G. (2013). Multiple alignment-free sequence comparison. Bioinformatics, 29(21), 2690.

Rice, P., Longden, I., and Bleasby, A. (2000). EMBOSS: the european molecular biology open software suite. Trends Genet, 16(6), 276–277.

Rizk, G. and Lavenier, D. (2010). GASSST: global alignment short sequence search tool. Bioinformatics, 26(20), 2534–2540.

Rognes, T. (2011). Faster smith-waterman database searches with inter-sequence simd parallelisation. BMC Bioinformatics, 12(1), 221.

Sims, G. E., Jun, S.-R., Wu, G. A., and Kim, S.-H. (2009). Alignment-free genome comparison with feature frequency profiles (FFP) and optimal resolutions. Proc Natl Acad Sci U S A, 106(8), 2677–2682.

Song, K., Ren, J., Reinert, G., Deng, M., Waterman, M. S., and Sun, F. (2014). New developments of alignment-free sequence comparison: measures, statistics and next-generation sequencing. Brief Bioinform, 15(3), 343.

Velasco II A., James, B. T., Wells, V. D., and Girgis, H. Z. (2018). Look4TRs: A de-novo tool for detecting simple tandem repeats using self-supervised hidden Markov models. BioRxiv, page 449801.

Vinga, S. (2014). Editorial: Alignment-free methods in computational biology. Brief Bioinform, 15(3), 341–342.

Vinga, S. and Almeida, J. (2003). Alignment-free sequence comparison–a review. Bioinformatics, 19(4), 513.

Vinga, S., Carvalho, A. M., Francisco, A. P., Russo, L. M., and Almeida, J. S. (2012). Pattern matching through Chaos Game Representation: bridging numerical and discrete data structures for biological sequence analysis. Algorithms Mol Biol, 7(1), 10.

Wu, T.-J., Hsieh, Y.-C., and Li, L.-A. (2001). Statistical measures of dna sequence dissimilarity under markov chain models of base composition. Biometrics, 57, 441–448.

Zerbino, D. R., Achuthan, P., Akanni, W., Amode, M. R., Barrell, D., Bhai, J., Billis, K., Cummins, C., Gall, A., Girón, C. G., et al. (2017). Ensembl 2018. Nucleic Acids Res, 46(D1), D754–D761.

Zhang, Y. and Chen, W. (2011). A new measure for similarity searching in dna sequences. MATCH Commun. Math. Comput. Chem., 65(2), 477–488.

Zharkikh, A. A. and Rzhetsky, A. Y. (1993). Quick assessment of similarity of two sequences by comparison of their l-tuple frequencies. Biosystems, 30, 93–111.

Zielezinski, A., Vinga, S., Almeida, J., and Karlowski, W. M. (2017). Alignment-free sequence comparison: benefits, applications, and tools. Genome Biol, 18(1), 186.

