## Supplementary Data 1 for "FASTCAR: Rapid alignment-free prediction of sequence alignment identity scores using self-supervised general linear models": Makefile

```
all: bin/Red.o bin/fastcar

bin/Red.o:
	mkdir -p bin
	mkdir -p bin/exception
	mkdir -p bin/nonltr
	mkdir -p bin/utility
	$(MAKE) -C src
bin/fastcar: bin/Red.o
	$(MAKE) -C src/cluster
	cp src/cluster/fastcar bin

clean:
	$(MAKE) clean -C src
	$(MAKE) clean -C src/cluster
	$(RM) -r bin

rebuild: clean all
.PHONY: all clean
```
