## Supplementary Data 1 for "FASTCAR: Rapid alignment-free prediction of sequence alignment identity scores using self-supervised general linear models": README

FASTCAR - Fast Alignment-free Search Tool for Classification and Regression
Release version

Requirements: g++ 4.9.1 or later, requires Homebrew on Mac OS X

Compilation using g++ (homebrew) and GNU Make on Mac OS X
CXX=g++-7 make

see: https://stackoverflow.com/questions/29057437/compile-openmp-programs-with-gcc-compiler-on-os-x-yosemite


Linux/Unix compilation:
make

Usage: bin/fastcar *.fasta --query queryFile.fasta [--id 0.90] [--mode rc] [--feat fast|slow] [--kmer 3] [--chunk 10000] [--output output_first_string] [--sample 3000] [--threads 4]

All arguments except for id can be shortened to one dash followed by the first letter of the argument,
for example: --query => -q

The most important parameter, --id, controls the identity of the sequences.

--query is a required parameter which takes in a query file, which searches against the database FASTA files

--mode specifies the mode[s], which can be "r" for regression and "c" for classification.
       They can be combined to form the default mode, which regresses identity scores above a cutoff.

--feat specifies which features are used. By default, "fast" (9 features) are used. However, for added
       potential accuracy, "slow" can be used, utilizing 2 more features which are slower.

--output specifies the output file prefix, to which .searchXXX is appended.

--chunk specifies chunk size, or, how many sequences are read in at once and processed. The larger
	the size, the more memory is used. However, this may lead to faster processing as more
	sequences can be processed at once.

--kmer decides the size of the kmers. It is by default automatically decided by average sequence length,
       but if provided, MeShClust can speed up a little by not having to find the largest sequence length.
       Increasing kmer size can increase accuracy, but increases memory consumption fourfold.


--threads sets the number of threads to be used. By default OpenMP uses the number of available cores
	  on your machine, but this parameter overwrites that.

--sample selects the total number of sample pairs of sequences used for both training and testing.
	 1500 is the default value.


If the argument is not listed here, it is interpreted as an input file.
