## Supplementary Data 1 for "FASTCAR: Rapid alignment-free prediction of sequence alignment identity scores using self-supervised general linear models": Makefile

```
TARGET ?= fastcar
VERSION ?= 0.7.1
CXX ?= g++
ifeq ($(debug),yes)
	CXXFLAGS += -ggdb -fno-omit-frame-pointer -fopenmp -DMUTDEBUG
#	LDFLAGS += -fsanitize=thread
else
	CXXFLAGS += -fopenmp -O3 -march=native -g
endif
CXXFLAGS += -std=c++11 -DVERSION=\"$(VERSION)\"
LDFLAGS += -lm -lstdc++

SOURCES := $(shell find ./src -name '*.cpp')
OBJECTS = $(SOURCES:%.cpp=bin/%.o) ../train/Trainer.o ../train/Predictor.o ../train/HandleSeq.o ../train/MultiMute.o ../train/SingMute.o
BIN_OBJECTS := $(shell find  ../../bin/ -mindepth 2 -name '*.o')

all: $(TARGET)

$(TARGET): $(OBJECTS) $(BIN_OBJECTS)
	$(CXX) $(CXXFLAGS) $^ -o $@ $(LDFLAGS)

../train/Trainer.o: ../train/Trainer.cpp
	$(CXX) $(CXXFLAGS) -c $< -o $@
#../train/SingleSelector.o: ../train/SingleSelector.cpp
#	$(CXX) $(CXXFLAGS) -c $< -o $@
../train/Predictor.o: ../train/Predictor.cpp
	$(CXX) $(CXXFLAGS) -c $< -o $@
../train/HandleSeq.o: ../train/HandleSeq.cpp
	$(CXX) $(CXXFLAGS) -c $< -o $@
../train/MultiMute.o: ../train/MultiMute.cpp
	$(CXX) $(CXXFLAGS) -c $< -o $@
../train/SingleMute.o: ../train/SingleMute.cpp
	$(CXX) $(CXXFLAGS) -c $< -o $@
bin/%.o: %.cpp
	mkdir -p $(@D)
	$(CXX) $(CXXFLAGS) -c $< -o $@

clean:
	$(RM) $(OBJECTS) $(TARGET)

install: $(TARGET)
	cp $(TARGET) ~/bin

.PHONY: all clean install
```
