## Supplementary Data 1 for "FASTCAR: Rapid alignment-free prediction of sequence alignment identity scores using self-supervised general linear models": bvec.cpp

```
/* -*- C++ -*-
 *
 * bvec.cpp
 *
 * Author: Benjamin T James
 *
 * Data structure - binned vector
 * Used in MeShClust for storing bulk sequences
 * which can be easily filtered by length
 */
#include "bvec.h"
#include <algorithm>
template<class T>
bvec<T>::bvec(vector<uint64_t>& lengths, uint64_t bin_size)
{
	uint64_t num_points = lengths.size();
	std::sort(std::begin(lengths), std::end(lengths));
	for (uint64_t i = 0; i < lengths.size(); i += bin_size) {
		begin_bounds.push_back(lengths[i]);
		uint64_t last_index = std::min((uint64_t)lengths.size() - 1,
					       i + bin_size - 1);
		end_bounds.push_back(lengths[last_index]);
		//std::cout << "[" << i << " " << last_index << "]" << std::endl;
	}
	data.reserve(begin_bounds.size());
	for (uint64_t i = 0; i < begin_bounds.size(); i++) {
		data.push_back({});
	}
}

template<class T>
Point<T>* bvec<T>::pop()
{
	for (auto& bin : data) {
		if (!bin.empty()) {
			Point<T>* p = bin[0].first;
			bin.erase(std::begin(bin));
			return p;
		}
	}
	return NULL;
}

template<class T>
Point<T>* bvec<T>::peek() const
{
	for (auto& bin : data) {
		if (!bin.empty()) {
			Point<T>* p = bin[0].first;
			return p;
		}
	}
	return NULL;
}

template<class T>
bool bvec<T>::inner_index_of(uint64_t length, size_t &idx, size_t *pfront, size_t *pback) const
{

	if (data.at(idx).empty() || idx == data.size()) {
		if (pfront) {
			for (size_t i = 0; i < data.size(); i++) {
				if (!data.at(i).empty()) {
					idx = i;
					*pfront = 0;
					break;
				}
			}
		}
		if (pback) {
			for (int i = data.size()-1; i >= 0; i--) {
				if (!data.at(i).empty()) {
					idx = i;
					*pback = 0;
					break;
				}
			}
		}
		return true;
	}
	size_t front = 0, back = 0;
	size_t low = 0, high = data.at(idx).size() - 1;
	bool found = false;
	if (length < data[idx][low].first->get_length() && pfront != NULL) {
		*pfront = low;
	}
	if (length > data[idx][high].first->get_length() && pback != NULL) {
		*pback = high;
	}
	for (;low <= high;) {
		size_t mid = (low + high) / 2;
		uint64_t d = data[idx][mid].first->get_length();
		if (d == length) {
			front = mid;
			back = mid;
			found = true;
			break;
		} else if (length < d) {
			high = mid;
		} else if (length > d) {
			low = mid + 1;
		}
		if (low == high) {
			found = true;
			front = low;
			back = high;
			break;
		}
	}
	if (pfront) {
		for (long i = front; i >= 0
			     && data[idx][i].first->get_length() == length; i--) {
			front = i;
		}
		*pfront = front;
	}
	if (pback) {
		for (long i = back; i < data[idx].size()
			     && data[idx][i].first->get_length() == length; i++) {
			back = i;
		}
		*pback = back;
	}
	return true;
}

template<class T>
bool bvec<T>::index_of(uint64_t point, size_t* pfront, size_t* pback) const
{
	size_t front = 0, back = 0;
        intmax_t low = 0, high = begin_bounds.size() - 1;
	bool found = false;
	if (point < begin_bounds[low] && pfront != NULL) {

		while (data[low].empty() && (low + 1) < data.size()) {
			low++;
		}
		*pfront = low;
		return true;
	}
	if (point > begin_bounds[high] && pback != NULL) {

		while (data[high].empty() && high > 0) {
			high--;
		}
		*pback = high;
		return true;
	}
	for (;low <= high;) {
		size_t mid = (low + high) / 2;
		if (begin_bounds.at(mid) <= point && end_bounds.at(mid) >= point) {
			front = mid;
			back = mid;
			found = true;
			break;
		} else if (point < begin_bounds[mid] && mid > 0) {
			high = mid - 1;
		} else if (point > end_bounds[mid] && mid < begin_bounds.size()-1) {
			low = mid + 1;
		} else {
			found = false;
			break; // not found
		}
	}
	if (!found) {
		std::cerr << "error: list not sorted" << std::endl;
		throw 100;
		return false;
	}
	if (pfront) {
		for (long i = front; i >= 0
			     && begin_bounds[i] <= point
			     && end_bounds[i] >= point; i--) {
			front = i;
		}
		// while (data[front].empty() && (front + 1) < data.size()) {
		// 	front++;
		// }
		*pfront = front;
	}
	if (pback) {
		for (long i = back; i < data.size()
			     && begin_bounds[i] <= point
			     && end_bounds[i] >= point; i++) {
			back = i;
		}
		// while (data[back].empty() && back > 0) {
		// 	back--;
		// }
		*pback = back;
	}
	return true;
}

template<class T>
void bvec<T>::insert(Point<T> *p)
{
	uint64_t len = p->get_length();
	size_t front = 0, back = 0;
	bool good = index_of(len, &front, &back);
	if (!good || front > back) {
		std::cerr << "error: list is not sorted" << std::endl;
	}
	std::vector<size_t> min_sizes;
	size_t minimum = std::numeric_limits<size_t>::max();
	for (size_t i = front; i <= back; i++) {
		size_t sz = data[i].size();
		if (sz < minimum) {
			minimum = sz;
			min_sizes.clear();
			min_sizes.push_back(i);
		} else if (sz == minimum) {
			min_sizes.push_back(i);
		}
	}
	if (min_sizes.empty()) {
		std::cerr << "error: no bins to insert into, item not inserted" << std::endl;
	}
	auto mid_min = min_sizes[min_sizes.size() / 2];
	data.at(mid_min).push_back(std::make_pair(p, false));
}

template<class T>
size_t bvec<T>::size() const
{
	size_t num_bins = data.size();
	size_t total_size = 0;
	for (size_t i = 0; i < num_bins; i++) {
		total_size += data[i].size();
	}
	return total_size;
}

template<class T>
size_t bvec<T>::report() const
{
	cout << "BVec: ";
	size_t num_bins = data.size();
	cout << "num_bins=" << num_bins << endl;
	size_t total_size = 0;
	for (size_t i = 0; i < num_bins; i++) {
		cout << "Bin " << i << ": [" << begin_bounds[i] << " " << end_bounds[i] << "] size=" << data[i].size() << endl;
		total_size += data[i].size();
	}
	cout << "total_size=" << total_size << endl;
	return total_size;
}
template<class T>
void bvec<T>::insert_finalize()
{
	auto sorter = [](const std::pair<Point<T>*,bool> a, const std::pair<Point<T>*,bool> b) {
		return a.first->get_length() < b.first->get_length();
	};
	for (size_t i = 0; i < data.size(); i++) {
		std::sort(std::begin(data[i]), std::end(data[i]), sorter);
		data[i].shrink_to_fit();
	}
}

template<class T>
bool bvec<T>::empty() const
{
	bool is_empty = true;
	for (auto bin : data) {
		if (!bin.empty()) {
			is_empty = false;
			break;
		}
	}
	return is_empty;
}


template<class T>
uint64_t bvec<T>::absolute_idx(bvec_idx_t idx) const
{
	uint64_t ptr = 0;
	for (int i = 0; i < idx.first; i++) {
		ptr += data[i].size();
	}
	ptr += idx.second;
	return ptr;
}

template<class T>
std::pair<bvec_idx_t, bvec_idx_t>
bvec<T>::get_range(uint64_t begin_len, uint64_t end_len) const
{
	/* perform binary search to find bin */
	bvec_idx_t front, back;
	front.first = 0;
	front.second = 0;
	back.first = data.size()-1;
	back.second = data[back.first].size() - 1;
	if (!index_of(begin_len, &front.first, NULL)) {
		throw 100;
	}
	if (!index_of(end_len, NULL, &back.first)) {
		throw 100;
	}
	if (!inner_index_of(begin_len, front.first, &front.second, NULL)) {
		throw 100;
	}
	if (!inner_index_of(end_len, back.first, NULL, &back.second)) {
		throw 100;
	}
	// if (back.first != data.size()) { // ++ to make it an end iterator
	// 	if (back.second != data[back.first].size()) {
	// 		back.second++;
	// 	} else {
	// 		back.first++;
	// 		back.second = 0;
	// 	}
	// } else {
	// 	throw 101;
	// }
	return std::make_pair(front, back);
}

template<class T>
void bvec<T>::erase(size_t r, size_t c)
{
	data.at(r).erase(data.at(r).begin() + c);
}

/*
 * TODO: change available to Center class so no intermediate copying is done
 */
template<class T>
void bvec<T>::remove_available(bvec_idx_t begin, bvec_idx_t end, std::vector<Point<T>*> &available)
{
	size_t a = begin.first;
	size_t b = end.first;
	int num = 0, new_num = 0;
	auto func = [](const bv_data_type<T> d) { return d.second; };
	auto inserter = [&](const std::pair<Point<T>*,bool> p) {
		if (p.second) {
#pragma omp critical
			available.push_back(p.first);
		}
	};
	#pragma omp parallel for
	for (size_t i = a; i <= b; i++) {
		/* move marked points to end of vector, then copy, then erase */
		//const auto last = std::remove_if(std::begin(data[i]), std::end(data[i]), func);
		for (int j = 0; j < data[i].size(); j++) {
			auto kv = data[i][j];
			if (kv.second) {
#pragma omp critical
				{
					available.push_back(kv.first);
				}
			}
		}
		data[i].erase(std::remove_if(std::begin(data[i]), std::end(data[i]), func), std::end(data[i]));
	}
}


template<class T>
bvec_iterator<T> bvec<T>::iter(bvec_idx_t idx)
{
	return bvec_iterator<T>(idx.first, idx.second, &data);
}


template class bvec<uint8_t>;
template class bvec<uint16_t>;
template class bvec<uint32_t>;
template class bvec<uint64_t>;
template class bvec<int>;
template class bvec<double>;
```
