## Supplementary Data 1 for "FASTCAR: Rapid alignment-free prediction of sequence alignment identity scores using self-supervised general linear models": bvec.h

```
/* -*- C++ -*-
 *
 * bvec.h
 *
 * Author: Benjamin T James
 *
 * Data structure - binned vector
 * Used in MeShClust for storing bulk sequences
 * which can be easily filtered by length
 */
#ifndef BVEC_H
#define BVEC_H

#include "Point.h"
#include "bvec_iterator.h"

typedef struct bvec_idx {
	size_t first, second;
} bvec_idx_t;

/*
 * operations needed:
 *
 * find bounds (range)
 * get available or min and remove
 *
 */
template<class T>
using bv_data_type = std::pair<Point<T>*, bool>;

template<class T>
using bv_row_type = vector<bv_data_type<T> >;

template<class T>
using bv_col_type = vector<bv_row_type<T> >;

template<class T>
class bvec {
public:
	bvec(vector<uint64_t>& lengths, uint64_t bin_size=1000);

	Point<T>* pop();
	Point<T>* peek() const;
	void insert(Point<T>* data);
	void insert_finalize(); /* sorts bins */


	bool index_of(uint64_t length, size_t* front, size_t* back) const;
	bool inner_index_of(uint64_t length, size_t& idx, size_t *front, size_t *back) const;
	bool empty() const;

	void check() const {
		for (int i = 0; i < data.size(); i++) {
			for (int j = 0; j < data[i].size(); j++) {
				if (data[i][j].first->get_header() == ">seq76 template_4") {
					cout << "seq76 is at index [" << i << "]["<< j << "]" << endl;
					return;
				}
			}

		}
		cout << "seq76 is NOT here" << endl;
	}
	std::pair<bvec_idx_t, bvec_idx_t>
	get_range(uint64_t begin_len, uint64_t end_len) const;

	void remove_available(bvec_idx_t begin, bvec_idx_t end, std::vector<Point<T>*> &);

	uint64_t absolute_idx(bvec_idx_t idx) const;

        bvec_iterator<T> iter(bvec_idx_t idx);
	typedef bvec_iterator<T> iterator;
	typedef bvec_iterator<T> const_iterator;

	size_t report() const;
	size_t size() const;

	void erase(size_t r, size_t c);
private:
        bv_col_type<T> data;
	vector<uint64_t> begin_bounds, end_bounds;
};


#endif
```
