## Supplementary Data 1 for "FASTCAR: Rapid alignment-free prediction of sequence alignment identity scores using self-supervised general linear models": bvec_iterator.cpp

```
/* -*- C++ -*-
 *
 * bvec_iterator.h
 *
 * Author: Benjamin T James
 *
 * Iterator for bvec
 */
#include "bvec_iterator.h"

template<class T>
bvec_iterator<T> bvec_iterator<T>::operator++()
{
	if (r != col->size()) {
		if (c + 1 < col->at(r).size()) {
			c++;
		} else {
			r++;
			c = 0;
			while (r < col->size() && col->at(r).empty()) {
				r++;
			}
		}
	} else {
		cerr << "tried incrementing null iterator" << endl;
		throw 10;
	}
	return *this;
}

template class bvec_iterator<uint8_t>;
template class bvec_iterator<uint16_t>;
template class bvec_iterator<uint32_t>;
template class bvec_iterator<uint64_t>;
template class bvec_iterator<int>;
template class bvec_iterator<double>;
```
