## Supplementary Data 1 for "FASTCAR: Rapid alignment-free prediction of sequence alignment identity scores using self-supervised general linear models": bvec_iterator.h

```
/* -*- C++ -*-
 *
 * bvec_iterator.h
 *
 * Author: Benjamin T James
 *
 * Iterator for bvec
 */
#include "bvec.h"
#ifndef BVEC_ITERATOR_H
#define BVEC_ITERATOR_H


template<class T>
class bvec_iterator {
public:
	// iterator: split ALL possible points into chunks by indices
	using dtype = std::pair<Point<T>*,bool>;
	using vtype = vector<vector<dtype> >;
	bvec_iterator(size_t _r,
		      size_t _c,
		      vtype* col_) : r(_r), c(_c), col(col_) {}

	bvec_iterator operator++();
	bvec_iterator operator++(int x) {
		return ++(*this);
	}
	dtype& operator*() {
		return col->at(r).at(c);
	}
	void operator+=(int64_t n) {
		if (n < 0) {
			throw "oops";
		}
		for (int i = 0; i < n; i++) {
			operator++();
		}
	}
	bool operator==(const bvec_iterator& rhs) const {
		return rhs.c == c && rhs.r == r;
	}
	bool operator<(const bvec_iterator& rhs) const {
		if (r < rhs.r) {
			return true;
		} else if (r == rhs.r) {
			return c < rhs.c;
		} else {
			return false;
		}
	}
	bool operator<=(const bvec_iterator& rhs) const {
		if (r < rhs.r) {
			return true;
		} else if (r == rhs.r) {
			return c <= rhs.c;
		} else {
			return false;
		}
	}
	bool operator!=(const bvec_iterator& rhs) const {
		return r != rhs.r || c != rhs.c;
	}
	int64_t operator-(const bvec_iterator& rhs) const {
		int64_t sum = 0;
		if (*this < rhs) {
			return -1 * (rhs - *this);
		}
		// subtract cols until last row is reached
		if (r == rhs.r) {
			return c - rhs.c;
		}
		sum += c;
		sum += col->at(rhs.r).size() - rhs.c;
		for (size_t i = rhs.r + 1; i < r; i++) {
			sum += col->at(i).size();
		}
		return sum;
	}
	// bvec_iterator operator[](uint64_t idx) {

	// }
//private:
	size_t r,c;
        vtype* col;
};
#endif
```
