## Supplementary Data 1 for "FASTCAR: Rapid alignment-free prediction of sequence alignment identity scores using self-supervised general linear models": Center.h

```
/* -*- C++ -*-
 *
 * Center.h
 *
 * Author: Benjamin T James
 *
 * Class representing a center in MeShClust,
 * which has a center and points which belong to the center
 */
#ifndef CENTER_H
#define CENTER_H

#include "Point.h"

template<class T>
struct Center {
	Center(Point<T>* c, const vector<Point<T>*> &pts) : center(c->clone()), points(pts), is_to_delete(false) {
	}
	Center(const Center<T> &cc) : center(cc.center->clone()), points(cc.points), is_to_delete(cc.is_to_delete) {}


	// Center(const Center<T>& c) {
	// 	center = c.get_clone();
	// 	points = c.getPoints_c();
	// 	is_to_delete = c.is_delete();
	// }
	~Center() { if (is_to_delete) { delete center; }}

	Point<T>* getCenter() { return center; }
	vector<Point<T>*> &getPoints() { return points; }

	const vector<Point<T>*> &getPoints_c() const { return points; };
	bool is_delete() const { return is_to_delete; }
	void lazy_remove() { is_to_delete = true; }
	size_t size() const { return points.size(); }
	bool empty() const { return points.empty(); }
	Point<T>* get_clone() const {
		return center->clone();
	}
	Point<T> *center;
	vector<Point<T>*> points;
	bool is_to_delete;
};

#endif
```
