## Supplementary Data 1 for "FASTCAR: Rapid alignment-free prediction of sequence alignment identity scores using self-supervised general linear models": ClusterFactory.cpp

```
/* -*- C++ -*-
 *
 * ClusterFactory.cpp
 *
 * Author: Benjamin T James
 * Contains code for building histograms, see get_divergence_point
 */

#ifndef HEADER_HACK
#include "ClusterFactory.h"
#endif

#include <assert.h>
#include <regex>
#include <tuple>
#include <queue>
#include <string>
#include <list>
#include "Histogram.h"
#include "../../nonltr/KmerHashTable.h"
#include "../../nonltr/ChromListMaker.h"
#include "DivergencePoint.h"
#include "Center.h"
#include "Progress.h"
//#include <omp.h>

template<class T>
T avg_distance(Point<T> &c, const std::vector<Point<T>*> &vec)
{
	T dist = 0;
	for (auto pt : vec) {
		dist += pt->distance(c);
	}
	return dist / vec.size();
}
template<class T>
Point<T>* find_center(const std::vector<Point<T>*> &vec)
{
	Point<T>* best = vec.front();
	T lowest = avg_distance(*best, vec);
	for (int i = 1; i < vec.size(); i++) {
		T dist = avg_distance(*vec[i], vec);
		if (dist < lowest) {
			best = vec[i];
			lowest = dist;
		}
	}
	return best;
}


template<class T>
void old_merge(vector<Point<T>*> &centers, map<Point<T>*,vector<Point<T>*>*> &clusters, T bandwidth)
{
	cout << "Merging points ... ";
	cout.flush();
	vector<Point<T>*> new_centers;
	vector<Point<T>*> to_delete;
	for (int i = 0; i < centers.size(); i++) {
		bool is_good = true;
		for (int j = i + 1; j < centers.size(); j++) {
			T dist = centers[i]->distance(*centers[j]);
			if (dist < bandwidth) {
				cout << "Merging centers " << centers[i]->get_header() << " and " << centers[j]->get_header() << endl;
				for (auto p : *clusters[centers[i]]) {
					clusters[centers[j]]->push_back(p);
				}
				delete clusters[centers[i]];
				clusters[centers[i]] = NULL;
				centers[i]->set_to_delete(true);
				to_delete.push_back(centers[i]);
				delete centers[i];
				is_good = false;
				break;
			}
		}
		if (is_good) {
			new_centers.push_back(centers[i]);
		}
	}
	for (auto it = clusters.begin(); it != clusters.end(); it++) {
		if ((*it).first->is_to_delete()) {
			clusters.erase(it);
		}
	}
	centers.clear();
	centers = new_centers;
	cout << "Done" << endl;
}

template<class T>
void sort_nn_func(std::vector<Point<T> *> &points, std::function<bool(const Point<T>&, const Point<T>&)> func, std::function<uint64_t(const Point<T>&, const Point<T>&)> distfunc) {
	if (points.empty()) {
		return;
	}
	cout << "Sorting points... ";
	cout.flush();
	list<int> good;
	int good_idx = 0;
	for (int i = points.size() - 1; i > 0; i--) { // start at idx 1 bc we don't want to redelete it
		good.push_front(i);
	}
	vector<Point<T>*> total_points;
	total_points.push_back(points[0]);
//	good.erase(good.begin());
	while (good.size() > 0) {
		auto last = total_points.back();
		auto smallest = good.begin();
		uint64_t small_dist = std::numeric_limits<uint64_t>::max();// / points[*smallest]->prob_under(*last);
		int count = 0;

		for (auto i = good.begin(); i != good.end(); i++) {
			if (func(*points[*i], *last)) {
//				cout << "Breaking loop " << points[*i]->get_length() << " " << last->get_length() << " -> " << count << endl;
				break;
			}
			uint64_t dist = distfunc(*points[*i],*last);// / points[*i]->prob_under(*last);
			if (dist < small_dist) {
				small_dist = dist;
				smallest = i;
			}
			count++;
		}
//		cout << "Number of gaps: " << num_gaps << endl;
		// if (func(*points[*smallest], *last)) {
		// 	cout << "Gap " << points[*smallest]->get_length() << " " << last->get_length() << endl;
		// }
//		cout << "Sorting: " << points[*smallest]->get_header() << endl;
		total_points.push_back(points[*smallest]);
		//	cout << points[*smallest]->get_header() << endl;
		// if (total_points.size() % 100 == 0) {
		// 	cout << "Size: " << total_points.size() << endl;
		// }
		good.erase(smallest);
	}
	assert(good.empty());
	assert(points.size() == total_points.size());
	points = total_points;
	cout << "Done" << endl;
}

template<class T>
void sort_nn_length(std::vector<Point<T>*> &points, double similarity) {
	similarity *= 0.90;
	if (points.empty()) {
		return;
	}
	cout << "Sorting points by length... ";
	cout.flush();
	list<int> good;
	int good_idx = 0;
	for (int i = 1; i < points.size(); i++) { // start at idx 1 bc we don't want to redelete it
		good.push_back(i);
	}
	vector<Point<T>*> total_points;
	total_points.push_back(points[0]);
//	good.erase(good.begin());
	bool working = true;
	while (working && good.size() > 0) {
		working = false;
		auto last = total_points.back();
		auto smallest = good.begin();
		T small_dist = points[*smallest]->distance(*last);// / points[*smallest]->prob_under(*last);
		for (auto i = good.begin(); i != good.end(); i++) {
			double ratio = 100.0 * (double)points[*i]->get_length() / last->get_length();
			if (ratio < similarity) {
				//	cout << "Length ratio " << ratio << " is less than " << similarity << "." << endl;
				break;
			}
			T dist = points[*i]->distance(*last);// / points[*i]->prob_under(*last);
			if (dist < small_dist) {
				small_dist = dist;
				smallest = i;
			}
		}
		total_points.push_back(points[*smallest]);
		//	cout << points[*smallest]->get_header() << endl;
		// if (total_points.size() % 100 == 0) {
		// 	cout << "Size: " << total_points.size() << endl;
		// }
		good.erase(smallest);
		working = true;
	}
	assert(good.empty());
	assert(points.size() == total_points.size());
	points = total_points;
	cout << "Done" << endl;
}


template<class T>
void calculate_gaps(const vector<Point<T>*> &vec, queue<int> &gaps, std::function<bool(const Point<T>&, const Point<T>&)> func)
{
	for (int i = 1; i < vec.size(); i++) {
		if (func(*vec[i], *vec[i-1])) {
			gaps.push(i);
		}
	}
}

// TODO: fix bounds
template<class T>
pair<int,int> find_bound(int idx, const vector<Point<T>*> &vec, double sim)
{
	size_t begin_len = vec[idx]->get_length();
	int begin_idx = idx;
	int end_idx = idx;
	#pragma omp parallel for
	for (int j = 0; j < 2; j++) {
		if (j == 1) {
			for (int i = idx - 1; i >= 0; i--) {
				if (vec[i]->get_id() == 0) {
					size_t len = vec[i]->get_length();
					if (begin_len < sim * len) {
						break;
					}
					begin_idx = i;
				}
			}
		} else {
			for (int i = idx + 1; i < vec.size(); i++) {
				if (vec[i]->get_id() == 0) {
					size_t len = vec[i]->get_length();
					if (len < sim * begin_len) {
						break;
					}
					end_idx = i;
				}
			}
		}
	}
	if (begin_idx < end_idx) {
		return make_pair(begin_idx, end_idx);
	} else {
		return make_pair(0, vec.size() - 1);
	}
}

template<class T>
vector<pair<int,double> > get_available_or_min(const vector<Point<T>*> &points, Point<T>* p, pair<int,int> bounds, const Trainer<T>& trn, bool& used_min)
{
	vector<pair<Point<T>*,int> > good;
	for (int i = bounds.first; i <= bounds.second; i++) {
		if (points[i]->get_id() == 0) {
			good.push_back(make_pair(points[i], i));
		}
	}
	bool f;
	vector<pair<int,double> > close;// = trn.get_close(p, good, f);
	used_min = f;
	return close;
	// if (used_min) {
	// 	used_min = true;
	// 	// find min
	// 	uintmax_t minimum = std::numeric_limits<uintmax_t>::max();
	// 	int min_index = -1;
	// 	vector<pair<int,int> > v;
	// 	for (int i = 0; i < good.size(); i++) {
	// 		uintmax_t u = p->distance(*good[i].first);
	// 		if (u < minimum) {
	// 			min_index = good[i].second;
	// 			minimum = u;
	// 		}
	// 	}
	// 	//std::cout << "none found, using minimum, dist = " << 1.0 - (double)minimum / 10000 << " " << points[min_index]->get_header() << endl;
	// 	//	v.push_back(make_pair(min_index, minimum));
	// 	v.push_back(make_pair(close[0], p->distance(*points[close[0]])));
	// 	return v;
	// } else
	// if (!close.empty()) {
	// 	vector<pair<int,int> > v(close.size());
	// 	#pragma omp parallel for
	// 	for (int i = 0; i < close.size(); i++) {
	// 		uintmax_t u = p->distance(*points[close[i]]);
	// 		v.at(i) = make_pair(close[i], u);
	// 	}
	// 	return v;
	// } else {
	// 	vector<pair<int,int> > v;
	// 	return v;
	// }
}

template<class T>
void mean_shift_update(vector<Center<T> > &part, int j, const Trainer<T>& trn, int delta)
{
	auto center = part[j].getCenter();

	int i_begin = std::max(0, j - delta);
	int i_end = std::min(j + delta, (int)part.size()-1);
	// if (i_begin == i_end) {
	// 	return;
	// }
	Point<double>* top = center->create_double();
	top->zero();
	Point<double>* temp = top->clone();
	uintmax_t bottom = 0;
	vector<pair<Point<T>*, bool> > good;
	for (int i = i_begin; i <= i_end; i++) {
//		const auto& vec = part.at(centers[i]);
		const auto& vec = part[i].getPoints();
		for (auto p : vec) {
			good.push_back(make_pair(p, false));
			// p->set_arg_to_this_d(*temp);
			// *top += *temp;
			// bottom++;
		}
	}
	trn.filter(center, good);
	if (!good.empty()) {
		for (auto p : good) {
			p.first->set_arg_to_this_d(*temp);
			*top += *temp;
			bottom++;
		}
		*top /= bottom;
		Point<T>* next = trn.closest(top, good);
		// Point<T> *next = NULL;
		// int next_dist = std::numeric_limits<int>::max();
		// for (int i = 0; i < N; i++) {
		// 	int dist = points[i]->distance_d(*top);
		// 	if (dist < next_dist) {
		// 		next_dist = dist;
		// 		next = points[i];
		// 	}
		// }
		if (next != NULL) {
			center->set(*next);
		} else {
			cerr << "mean shift: NULL" << endl;
		}
	}
	delete top;
	delete temp;
}

template<class T>
Point<T>* get_mean(vector<Point<T>*> &available, Point<T>& last, double bandwidth)
{
	Point<double>* top = last.create_double();
	top->zero();
	Point<double>* temp = top->clone();
	double bottom = 0;
	const int N = available.size();
	if (N == 0) {
		throw "N cannot be 0, bad";
	}
	bottom = available.size();
	// TODO: parallelize this loop
	for (int i = 0; i < N; i++) {
		available[i]->set_arg_to_this_d(*temp);
		*top += *temp;
	}

	if (bottom != 0) {
		*top /= bottom;
	} else {
		cerr << "No points in vector" << endl;
		throw 5;
	}
#pragma omp declare reduction(cmin:std::pair<Point<T>*,double>: \
			      omp_out = omp_in.second < omp_out.second ? omp_in : omp_out ) \
        initializer (omp_priv = std::make_pair((Point<T>*)NULL, (double)std::numeric_limits<double>::max()))				\

	std::pair<Point<T>*,double> result = std::make_pair((Point<T>*)NULL, (double)std::numeric_limits<double>::max());
	//todo: add pragma back in
#pragma omp parallel for reduction(cmin:result)
	for (int i = 0; i < available.size(); i++) {
		double dist = available[i]->distance_d(*top);
		if (dist < result.second) {
			result = std::make_pair(available[i], dist);
		}
	}
	delete top;
	delete temp;
	if (result.first == NULL && !available.empty()) {
		throw "not working";
	}
	return result.first;
}

template<class T>
bool merge(vector<Center<T> > &centers, const Trainer<T>& trn, int delta, int bandwidth)
{
	int num_merge = 0;
	for (int i = 0; i < centers.size(); i++) {
		long ret = trn.merge(centers, i, i + 1, std::min((int)centers.size()-1, i + delta));
		if (ret > i) {

			num_merge++;
			auto &to_add = centers[ret].getPoints();
			auto &to_del = centers[i].getPoints();
			to_add.insert(std::end(to_add), std::begin(to_del), std::end(to_del));
			centers[i].lazy_remove();
		}
		// vector<pair<Point<T>*,double> > to_merge;
		// for (int j = i + 1; j < std::min((int)centers.size(), i + 1 + delta); j++) {
		// 	to_merge.push_back(std::make_pair(centers[j].getCenter(), -1));
		// }
		// Point<T>* closest = trn.merge(centers[i].getCenter(), to_merge);
		// if (closest != NULL) {
		// 	#ifdef DEBUG
		// 	cout << "Merged center " << centers[i]->get_header() << " and " << closest->get_header() << endl;
		// 	#endif
		// 	num_merge++;
		// 	// auto& to_del = partition[centers[i]];
		// 	// auto& to_add = partition[closest];
		// 	// to_add.insert(std::end(to_add), std::begin(to_del), std::end(to_del));
		// 	// partition.erase(centers[i]);
		// 	// centers[i]->set_to_delete(true);
		// 	auto& to_del = partition[centers[i]];
		// 	auto& to_add = partition[closest];
		// 	to_add.insert(std::end(to_add), std::begin(to_del), std::end(to_del));
		// 	partition.erase(centers[i]);
		// 	centers[i]->set_to_delete(true);

		// }
	}
	//cout << "Merged " << num_merge << " centers" << endl;
	centers.erase(std::remove_if(centers.begin(), centers.end(), [](const Center<T>& p) {
			return p.is_delete();
		}), centers.end());
	return num_merge > 0;
}

template<class T>
void print_output(const string& output, vector<Center<T> > & partition)
{
	cout << "Printing output" << endl;
	std::ofstream ofs;
	ofs.open(output, std::ofstream::out);
	int counter = 0;
	for (auto& cen : partition) {
		if (cen.empty()) {
			continue;
		}
		ofs << ">Cluster " << counter << endl;
		int pt = 0;
		for (auto p : cen.getPoints()) {
			string s = p->get_header();
			ofs << pt << "\t" << p->get_length() << "nt, " << s << "... ";
			if (p->get_id() == cen.getCenter()->get_id()) {
				ofs << "*";
			}
			ofs << endl;
			pt++;
		}
		counter++;
	}
	ofs.close();
}

template<class T>
void sort(vector<Point<T>*> &points, vector<Point<T>*> &centers, int bandwidth, double sim, const Trainer<T>& trn, string output_file, int iter, int delta)
{
	int cur = 0;
	points[0]->set_id(points.size());
	cur++;
	int last = 0;
	vector<int> v;
	using partition = map<Point<T>*, vector<Point<T>*> >;
	partition part;
	centers.push_back(points.front()->clone());
	part[centers.front()].push_back(points.front());

	while (true) {
		pair<int,int> bounds = find_bound(last, points, sim);
		bool used_min;
		auto available = get_available_or_min(points, points[last], bounds, trn, used_min);
		// std::sort(available.begin(), available.end(),
		// 	  [](const pair<int,int> a, const pair<int,int> b) {
		// 		  return a.second < b.second;
		// 	  });
//		auto available = trn.get_close(points[last], points, bounds, used_min);
		if (available.empty()) {
			break;
		}
//		used_min = used_min && !v.empty();
		if (used_min) {
			if (!v.empty()) {
				auto c = points[last]->clone();
				centers.push_back(c);
				for (auto idx : v) {
					part[c].push_back(points[idx]);
				}
				v.clear();
			}
			last = available.back().first;
		}
		for (auto pr : available) {
			if (cur % 10000 == 0) {
				cout << "Placed " << cur << endl;
			}
			points[pr.first]->set_id(cur);
			v.push_back(pr.first);
			cur++;
		}
		if (!used_min) {
			last = get_mean(v, *points[last], points, bandwidth);
		}
	}
	auto c = points[last]->clone();
	centers.push_back(c);
	for (auto idx : v) {
		part[c].push_back(points[idx]);
	}
	points[0]->set_id(0);
	centers[0]->set_id(0);

	cout << "Found " << centers.size() << " initial centers" << endl;
	assert(centers.size() == part.size());
	std::sort(points.begin(), points.end(), [](const Point<T>* a, const Point<T>* b) {
			return a->get_id() < b->get_id();
		});
	std::sort(centers.begin(), centers.begin(), [](const Point<T>* a, const Point<T>* b) {
			return a->get_id() < b->get_id();
		});
	// for (int i = 0; i < points.size(); i++) {
	// 	cout << points[i]->get_header() << " ";
	// 	if (i == 0) {
	// 		cout << endl;
	// 		continue;
	// 	};
	// 	int last_dist = points[i]->distance(*points[i-1]);
	// 	cout << last_dist << endl;
	// }
	Point<T>* lastp = NULL;
	for (auto c : centers) {
		auto v = part[c];
		for (auto p : v) {
			cout << p->get_header() << " ";
			cout << c->get_header() << " ";
			cout << p->distance(*c) << " ";
			if (lastp == NULL) {
				cout << endl;
			} else {
				cout << p->distance(*lastp) << endl;
			}
			lastp = p;
		}
	}
	for (int i = 0; i < iter; i++) {
		print_output(output_file + to_string(i), part);
		cout << "Mean shift iteration " << i << endl;
		#pragma omp parallel for
		for (int j = 0; j < centers.size(); j++) {
			mean_shift_update(part, centers, j, trn, delta);
		}
		merge(centers, part, trn, delta, bandwidth);
		for (auto const& kv : part) {
			if (kv.second.empty()) {
				cerr << "Empty cluster " << kv.first->get_header() << endl;
				throw 0;
			}
		}
	}
	for (int j = 0; j < centers.size(); j++) {
		mean_shift_update(part, centers, j, trn, 0);
	}
	print_output(output_file, part);
}


/*
 * Accumulates points in a center until none are close,
 * then returns the next center (not cloned)
 */
template<class T>
size_t accumulate(Point<T>** last_ptr, bvec<T> &points, vector<Center<T> > &centers,
		const Trainer<T>& trn, double sim, double bandwidth, int total_iter)
{
	Point<T>* last = *last_ptr;
	vector<Point<T>*> current = {last};
	bool is_min = false;

	for (int num_iter=0; !is_min; num_iter++) {
		#ifdef DEBUG
		cout << num_iter << " last: " << last->get_header() << endl;
		#endif
		auto len = last->get_length();
		auto bounds = points.get_range(len * sim, len / sim);
		auto result = trn.get_close(last,
			      points.iter(bounds.first),
			      points.iter(bounds.second),
			      is_min);

		if (is_min) {
			Point<T>* new_pt = get<0>(result);
			//	cout << "minimum point: " << new_pt->get_header() << endl;
			size_t r = get<2>(result);
			size_t c = get<3>(result);
			#ifdef DEBUG
			cout << "center added" << endl;
			#endif
			// no close points left for center,
			// returned value is the next center (return this)
			//points.remove_available(bounds.first, bounds.second, newvec);
			if (new_pt == NULL) {
				// No points left in range, try 1st point
				*last_ptr = points.pop();
			} else {
				// New center
				*last_ptr = new_pt;
				points.erase(r, c);
			}
			vector<Point<T>*> newvec;
			points.remove_available(bounds.first, bounds.second, newvec); // DEBUGGING USE ONLY
			if (!newvec.empty()) {
				throw "this should never happen";
			}
		} else { // keep adding points, find new mean
			size_t prev_size = current.size();
			points.remove_available(bounds.first, bounds.second, current);

			last = get_mean(current, *last, bandwidth);
			size_t added_size = current.size() - prev_size;
			#ifdef DEBUG
			cout << "added new points (" << added_size << ")" << endl;
			#endif
			if (last == NULL) {
				cerr << "Last is null" << endl;
				throw 100;
			}
		}
	}
//	cout << "Pushed back center " << last->get_header() << endl;
	Center<T> cc(last, current);
	centers.push_back(Center<T>(cc));
//	Center<T> cen(last, current);
//	centers.emplace_back(last, current);
	// Point<T>* center = last->clone();
	// centers.push_back(center);
	// part[center] = current;
	#ifdef DEBUG
	for (auto p : current) {
		cout << total_iter << " Cluster " << last->get_header() << ": " << p->get_header() << endl;
	}
	#endif
        // if (points.empty()) {
	// 	return true;
	// } else {
	// 	return false;
	// }
	return current.size();
}


template<class T>
void ClusterFactory<T>::MS(bvec<T> &points, T bandwidth, double sim, const Trainer<T>& trn, string output, int iter, int delta)
{
	vector<Center<T> > part;
//	using partition = map<Point<T>*, vector<Point<T>*> >;
//	partition part;
	Progress pa(points.size(), "Accumulation");
	Point<T>* last = points.pop();

        for (int num = 0; last != NULL; num++) {
	        size_t n = accumulate(&last, points, part, trn, sim, bandwidth, num);
		pa += n;
	}
	pa.end();
//	points.check();
	size_t total = 0;
	for (auto cen  : part) {
		total += cen.getPoints().size();
	}
	cout << "total size: " << total << endl;
	Progress pu(iter, "Update");
	for (int i = 0; i < iter; i++) {
		// #ifdef DEBUG
		//print_output(output + to_string(i), part);
		// #endif
		//cout << "Mean shift iteration " << i << endl;
		#pragma omp parallel for
		for (int j = 0; j < part.size(); j++) {
			mean_shift_update(part, j, trn, delta);
		}
		merge(part, trn, delta, bandwidth);
		pu++;
	}

	#pragma omp parallel for
	for (int j = 0; j < m_centers.size(); j++) {
		mean_shift_update(part, j, trn, 0);
	}
	pu.end();
	print_output(output, part);
}

/*
 * This uses a callback to specify the specific type of point.
 *
 * To call this, use like:
 *
 * factory.build_points("input", &ClusterFactory<PtType>::get_histogram);
 */
template<class T>
std::vector<Point<T>*> ClusterFactory<T>::build_points(vector<string> fileList, std::function<Point<T>*(ChromosomeOneDigit *)> get_point)
{
	std::vector<Point<T>*> points;
	std::vector<Point<T>*> cpoints;
	unsigned fsize = fileList.size();
	std::vector<Point<T>*> initial_centers;
	std::stringstream buffer;
	buffer << "Counting " << k << "-mers";
	Progress p(fsize, buffer.str());
	for (unsigned i = 0; i < fsize; i++) {
		p++;
		ChromListMaker *maker = new ChromListMaker(fileList.at(i));
		const std::vector<Chromosome *> * chromList = maker->makeChromOneDigitDnaList();
		unsigned csize = chromList->size();
#pragma omp parallel for ordered
		for (unsigned h = 0; h < csize; h++) {
			ChromosomeOneDigit *chrom = dynamic_cast<ChromosomeOneDigit *>(chromList->at(h));
			if (chrom) {
				Point<T> *h = get_point(chrom);
				if (h != NULL) {
#pragma omp ordered
					{
						//	cout << "Header: " << h->get_header() << endl;
					points.push_back(h);
					}
				}
			} else {
				throw InvalidStateException(string("Dynamic cast failed"));
			}
		}
		delete maker;
	}
	return points;
//	std::random_shuffle(points.begin(), points.end());
//	queue<int> gaps;
//	calculate_gaps<T>(points, gaps, func);
	// for (int i = 1; i < points.size(); i++) {
	// 	int la = points[i]->get_length();
	// 	int lb = points[i-1]->get_length();
	// 	if (lb > la && 100.0 * la / lb < sim) {
	// 		gaps.push(i);
	// 	}
	// }


// 	vector<vector<Point<T>*>> p;
// 	vector<Point<T>*> tmp;
// 	tmp.push_back(points[0]);
// 	for (int j = 1; j < points.size(); j++) {

// 		int la = points[j]->get_length();
// 		int lb = points[j-1]->get_length();
// 		assert(lb >= la);
// 		if (lb > la && 100.0 * la / lb < sim) {
// 			p.push_back(tmp);
// 			cout << "Gap " << tmp.size() << endl;
// 			tmp.clear();
// 		}
// 		tmp.push_back(points[j]);
// 	}
// 	if (!tmp.empty()) {
// 		p.push_back(tmp);
// 	}

// //	calculate_distances(points);
// 	int idx = 0;
// 	for (auto &c : p) {
// 		sort_nn_func<T>(c, func);
// 		for (auto v : c) {
// 			v->set_id(idx++);
// 			cpoints.push_back(v);
// 		}
// 	}

	// sort_nn_func<T>(points,
	// 		[&](const Point<T>&a, const Point<T>&b) {
	// 			int la = a.get_length();
	// 			int lb = b.get_length();
	// 			return lb > la && 100.0 * la / lb < sim;
	// 		},
	// 		[](const Point<T>& a, const Point<T>& b) {
	// 		        return a.distance_k1(b);
	// 		});


	// // for(auto p : points){
	// //	cout << p->get_header() << endl;
	// // }


	// sort_nn_func<T>(points,
	// 		[&](const Point<T>& a, const Point<T>& b) {
	// 			int la = a.get_length();
	// 			int lb = b.get_length();
	// 			if (lb > la && 100.0 * la / lb < sim) {
	// 				double mono = a.distance_k1(b) * 100;
	// 				bool q = mono < sim;
	// 				/*
	// 				if (q) {
	// 					cout << "TRUE" << endl;
	// 				} else {
	// 					cout << "FALSE"<< endl;
	// 				}
	// 				*/
	// 				return q;
	// 			} else {
	// 				return false;
	// 			}
	// 		},
	// 		[](const Point<T>& a, const Point<T>& b) {
	// 			return a.distance(b);
	// 		});
	// uint64_t idx = 0;
	// for (auto v : points) {
	// 	v->set_id(idx++);

	// 	cpoints.push_back(v);
	// }
	// cout << "Points: " << cpoints.size() << endl;


	// for (int i = 0; i < points.size(); i++) {
	// 	cout << points[i]->get_header();
	// 	if (i > 0) {
	// 		cout << "  " << points[i]->distance(*points[i-1]);
	// 	}
	// 	cout << endl;
	// }


	// for (int i = 0; i < points.size(); i++) {
	// 	points[i]->set_id(i);
	// 	cpoints.push_back(points[i]);
	// 	assert(cpoints[i]->get_id() == i);
	// }
	return points;
}


// consider all from 'to', distances[].size must be >= to.size()
template<class T>
Point<T>* find_nearest(const std::vector<Point<T>*> &to,
		       vector<int> &good, const std::vector<Point<T>*> &from,
		       std::vector<int>* distances, int& last_idx)
{
	// Step 1. Fill the closest distance list
	int best_dist = 0;
	Point<T>* best_pt = NULL;
	std::vector<int>::iterator best_idx;
	last_idx %= to.size();
	for (auto idx = good.begin(); idx != good.end(); idx++) {
		int i = *idx;
		distances[last_idx][i] = to[last_idx]->distance(*from[i]);
		int dist = 0;
		for (int j = 0; j < to.size(); j++) {
			dist += distances[j][*idx];
		}
		if (best_pt == NULL || dist < best_dist) {
			best_pt = from[i];
			best_dist = dist;
			best_idx = idx;
		}
	}
	cout << "Dist: " << best_dist << endl;
	last_idx++;
	good.erase(best_idx);
	return best_pt;
}

template<class T>
void ClusterFactory<T>::sort_nn(std::vector<Point<T> *> &points, Point<T>* nearest_to, int arg) const
{

	if (points.empty()) {
		return;
	}
	cout << "Sorting points... ";
	cout.flush();
	vector<int> good;
	int good_idx = points.size() - 1;
	for (int i = 0; i < points.size(); i++) {
		if (nearest_to != NULL && nearest_to == points[i]) {
			good_idx = i;
		}
		good.push_back(i);
	}
	vector<Point<T>*> total_points;
	total_points.push_back(points[good_idx]);
	good.erase(good.begin() + good_idx);
	bool working = true;
	while (working && good.size() > 0) {
		working = false;
		auto last = total_points.back();
		auto smallest = good.begin();
		T small_dist = points[*smallest]->distance(*last);
		for (auto i = good.begin(); i != good.end(); i++) {
			T dist = points[*i]->distance(*last);// / points[*i]->prob_under(*last);
			if (dist < small_dist) {
				small_dist = dist;
				smallest = i;
			}
		}
		total_points.push_back(points[*smallest]);
		//	cout << points[*smallest]->get_header() << endl;
		if (total_points.size() % 100 == 0) {
			cout << "Size: " << total_points.size() << endl;
		}
		good.erase(smallest);
		working = true;
	}
	assert(good.empty());
	assert(points.size() == total_points.size());
	points = total_points;
	cout << "Done" << endl;
}

template<class T>
Point<T> *ClusterFactory<T>::get_divergence_point(ChromosomeOneDigit *chrom)
{
	if (chrom == NULL) {
		return NULL;
	}
	KmerHashTable<unsigned long, T> table(k, 1);
	KmerHashTable<unsigned long, uint64_t> table_k1(1, 0);
	std::vector<T> values;
	vector<uint64_t> values_k1;
	values.clear();
	fill_table<T>(table, chrom, values);
	fill_table<uint64_t>(table_k1, chrom, values_k1);
//	int tmplate = get_template(chrom->getHeader(), templates);
	Point<T> *p =  new DivergencePoint<T>(values, chrom->size());
//	cout << "mag: " << ((DivergencePoint<T>*)p)->getPseudoMagnitude() << std::endl;
	p->set_1mers(values_k1);
	p->set_header(chrom->getHeader());
	p->set_length(chrom->getBase()->length());
	p->set_data_str(*chrom->getBase());
	DivergencePoint<T>* q = dynamic_cast<DivergencePoint<T>*>(p);
	const auto N = q->points.size();
	double aq = (double)q->getPseudoMagnitude() / N;
	double sq = 0;
	for (auto i = 0; i < N; i++) {
		double qdiff = q->points[i] - aq;
		sq += qdiff * qdiff;
	}
	sq = sqrt(sq / N);
	q->set_stddev(sq);
	return p;
}


template<class T>
Point<T> *ClusterFactory<T>::get_histogram(ChromosomeOneDigit *chrom)
{
	if (chrom == NULL) {
		return NULL;
	}
        KmerHashTable<unsigned long, T> table(k, 0);
	std::vector<T> values;
	values.clear();
	fill_table(table, chrom, values);
//	int tmplate = get_template(chrom->getHeader(), templates);
//	Point<T> *p = new Histogram<T>(values);
	Point<T> *p = new DivergencePoint<T>(values, chrom->size());
	p->set_header(chrom->getHeader());
	p->set_length(chrom->getBase()->length());
	return p;
}

template<class T>
T ClusterFactory<T>::find_h(const std::vector<Point<T>*> &centers) const
{
	int size = centers.size();
	T div = 0;
	int num_divergence = 0;
	vector<T> divs;
	for (int i = 0; i < size; i++) {
		for (int j = 0; j < size; j++) {
			if (j == i) { continue; }
			divs.push_back(centers[i]->distance(*centers[j]));
//			num_divergence++;
		}
	}
	std::sort(divs.begin(), divs.end());
	int end = divs.size() / 50;
	for (int i = 0; i < end; i++) {
		div += divs[i];
	}
	return div / end / 100;
	if (divs.size() % 2 == 0) {
		return (divs[divs.size()/2 - 1] + divs[divs.size()/2]) / 2;
	} else {
		return divs[divs.size()/2];
	}
}
/*
template<class T>
std::vector<Point<T> *> ClusterFactory<T>::get_centers(const std::vector<Point<T> *> &points)
{
	std::vector<Point<T>*> centers;
	for (typename std::vector<Point<T>*>::const_iterator it = points.begin(); it != points.end(); ++it) {
		Point<T> *p = *it;
		if (choose_center(*p)) {
			centers.push_back(p->clone());
		}
	}

	return centers;
}
*/
#ifndef HEADER_HACK
template class ClusterFactory<double>;
template class ClusterFactory<int>;
template class ClusterFactory<uint64_t>;
template class ClusterFactory<uint32_t>;
template class ClusterFactory<uint16_t>;
template class ClusterFactory<uint8_t>;

#endif
```
