## Supplementary Data 1 for "FASTCAR: Rapid alignment-free prediction of sequence alignment identity scores using self-supervised general linear models": ClusterFactory.h

```
/* -*- C++ -*-
 *
 * ClusterFactory.h
 *
 * Author: Benjamin T James
 *
 * Contains code on building histograms from sequences (get_divergence_point)
 */

#ifndef CLUSTERFACTORY_H
#define CLUSTERFACTORY_H


#include <iostream>
#include <vector>
#include <functional>
#include <limits>
#include "../../nonltr/ChromosomeOneDigit.h"
#include "../../nonltr/KmerHashTable.h"
#include "Point.h"
#include "../../train/Trainer.h"
#include "bvec.h"

template<class T>
class ClusterFactory {
public:
	ClusterFactory(int k_len, int npp=std::numeric_limits<int>::max()) : k(k_len), num_per_partition(npp) {}
	std::vector<Point<T>*> build_points(vector<string> files, std::function<Point<T>*(ChromosomeOneDigit*)> get_point);
        Point<T>* get_histogram(ChromosomeOneDigit *chrom);
	Point<T>* get_divergence_point(ChromosomeOneDigit *chrom);
	T find_h(const std::vector<Point<T>*> &centers) const;
	void sort_nn(std::vector<Point<T>*> &points, Point<T>* nearest_to=NULL, int arg=3) const;
	void MS(bvec<T> &points, T bandwidth, double sim, const Trainer<T>& trn, string output, int iter, int delta);
private:
	vector<int> lookup_table;
	vector<Point<T>*> m_centers;
	const int num_per_partition;
	int k;
	//void fill_table(KmerHashTable<unsigned long, T> &table, ChromosomeOneDigit *chrom, std::vector<T>& values);
};

template<class V>
void fill_table(KmerHashTable<unsigned long, V> &table, ChromosomeOneDigit *chrom, std::vector<V>& values)
{

	const int k = table.getK();
	auto segment = chrom->getSegment();
	const char *seg_bases = chrom->getBase()->c_str();
	for (vector<int> *v : *segment) {
		int start = v->at(0);
		int end = v->at(1);
		table.wholesaleIncrement(seg_bases, start, end - k + 1);
	}
	unsigned long tableSize = table.getMaxTableSize();
	values.reserve(values.size() + tableSize);
	const V * valueArray = table.getValues();
	std::copy(&valueArray[0], &valueArray[tableSize], std::back_inserter(values));
}

#ifdef HEADER_HACK
#ifndef CLUSTERFACTORY_C
#define CLUSTERFACTORY_C
#include "ClusterFactory.cpp"
#endif
#endif

#endif
```
