## Supplementary Data 1 for "FASTCAR: Rapid alignment-free prediction of sequence alignment identity scores using self-supervised general linear models": DivergencePoint.cpp

```
/* -*- C++ -*-
 *
 * DivergencePoint.cpp
 *
 * Author: Benjamin T James
 *
 * Main histogram type, includes distance() which is intersection() in Feature.cpp
 */
#include "DivergencePoint.h"
#include <cmath>
#include <cstring>
#include <cfenv>
#include <iostream>


template<class T>
double DivergencePoint<T>::prob_under(Point<T> &p) const
{
	const DivergencePoint<T>& c = dynamic_cast<const DivergencePoint<T>&>(p);
	double sum = 0;
	const size_t s = points.size();
	double total = 0;
	std::feclearexcept(FE_OVERFLOW);
	std::feclearexcept(FE_UNDERFLOW);
	for (int i = 0; i < s; i++) {
		sum += c.points[i];
		if (i % 4 == 3) {
			for (int j = i - 3; j <= i; j++) {
				double prob = c.points[j] / sum;
				double log_prob = log(prob);
				total += (points[j] - 1) * log_prob;
				if ((bool)std::fetestexcept(FE_UNDERFLOW)) {
					cout << "Underflow!" << endl;
				}
				//	cond.push_back(log(prob)/log4);
			}
			sum = 0;
		}
	}
	// for (size_t q = 0; q < s; q += 4) {
	// 	double sum = 0;
	// 	for (int i = q; i < q + 4; i++) {
	// 		sum += c.points[i];
	// 	}
	// 	for (int i = q; i < q + 4; i++) {
	// 		double prob = c.points[i] / sum;
	// 		double log_prob = log(prob);
	// 		total += (points[i] - 1) * log_prob;
	// 	}
	// }
	return exp(total / s);
}

template<class T>
double DivergencePoint<T>::distance_d(Point<double>& p) const
{
	const DivergencePoint<double>& c = dynamic_cast<const DivergencePoint<double>&>(p);
	uint64_t dist = 0;
	uint64_t mag = 0;
	for (auto i = 0; i < points.size(); i++) {
		dist += 2 * min(points[i],(T)c.points[i]);
		mag += points[i] + c.points[i];
	}
	double frac = (double)dist / mag;
	return 10000.0 * (1.0 - frac * frac);
}


template<class T>
uint64_t DivergencePoint<T>::distance(const Point<T>& p) const
{
	const DivergencePoint<T>& c = dynamic_cast<const DivergencePoint<T>&>(p);
	uint64_t dist = 0;
	const uint64_t mag = getPseudoMagnitude() + c.getPseudoMagnitude();
	#pragma omp simd
	for (auto i = 0; i < points.size(); i++) {
		dist += min(points[i], c.points[i]);
	}
	dist *= 2;
	double frac = (double)dist / mag;
	return 10000.0 * (1.0 - frac * frac);
}

template<class T>
double DivergencePoint<T>::distance_k1(const Point<T> &p) const
{
	uint64_t dist = 0;

	auto a = Point<T>::get_1mers(), b = p.get_1mers();
	uint64_t mag = 0;
	for (auto i = 0; i < 4; i++) {
		dist += std::min(a[i], b[i]);
		mag += a[i];
	}
	return (double)dist / (double)mag;

}
template<class T>
DivergencePoint<T>::DivergencePoint(const std::vector<T>& pts, uint64_t len)
{
	mag = 0;
	for (unsigned int i = 0; i < pts.size(); i++) {
		points.push_back(pts.at(i));
		mag += pts.at(i);
	}
//	display();
	nucl_length = len;
	to_delete = false;
	id = 0;
}


template<class T>
DivergencePoint<T>::DivergencePoint(unsigned int size)
{
	for (unsigned int i = 0; i < size; i++) {
		points.push_back(0);
	}
	to_delete = false;
	nucl_length = 0;
	id = 0;
}

template<class T>
void DivergencePoint<T>::operator*=(double d)
{
	unsigned int size = points.size();
	for (auto& pt : points) {
		pt *= d;
	}
}

template<class T>
bool DivergencePoint<T>::operator<(Point<T>& p) const
{
	const DivergencePoint<T>& h = dynamic_cast<const DivergencePoint<T>&>(p);
	unsigned int size = std::min(points.size(),h.points.size());
	/*int boundary = 0;
	for (unsigned int i = 0; i < size; i++) {
		if (points.at(i) > h.points.at(i)) {
			boundary++;
		} else if (points.at(i) < h.points.at(i)) {
			boundary--;
		}
	}
	return boundary < 0;*/
	for (unsigned int i = 0; i < size; i++) {
		if (points.at(i) >= h.points.at(i)) {
			return false;
		}
	}
	return true;
}

template<class T>
void DivergencePoint<T>::operator/=(double d)
{
	unsigned int size = points.size();
	for (unsigned int i = 0; i < size; i++) {
		points[i] /= d;
	}
//	cout << endl;
}

template<class T>
void DivergencePoint<T>::operator+=(Point<T>& p)
{
	const DivergencePoint<T>& h = dynamic_cast<const DivergencePoint<T>&>(p);
	unsigned int size = std::min(points.size(),h.points.size());
	for (unsigned int i = 0; i < size; i++) {
		points.at(i) += h.points.at(i);
	}
}

template<class T>
uint64_t DivergencePoint<T>::operator-(const Point<T>& p) const
{
	return distance(p);
}

template<class T>
void DivergencePoint<T>::set(Point<T>& p)
{
	const DivergencePoint<T>& h = dynamic_cast<const DivergencePoint<T>&>(p);
	points = std::vector<T>(h.points);
	set_length(h.get_length());
	to_delete = h.to_delete;
	Point<T>::set_header(h.get_header());
	set_id(h.get_id());
}

template<class T>
void DivergencePoint<T>::display() const
{
	unsigned size = points.size();
	for (unsigned i = 0; i < size; i++) {
		std::cout << points.at(i) << " ";
	}
	std::cout << std::endl;
}

template<class T>
void DivergencePoint<T>::zero()
{
	for (auto &i : points) {
		i = 0;
	}
}

template<class T>
void DivergencePoint<T>::addOne()
{
	for (auto& a : points) {
		a++;
	}
}

template<class T>
void DivergencePoint<T>::subOne()
{
	for (auto& a : points) {
		a--;
	}
}

/*
 * p(y|x) = cond_p
 * q(y|x) = cond_p
 */
template<class T>
double DivergencePoint<T>::divergence(Point<T>& p) const
{
	const DivergencePoint<T>& d = dynamic_cast<const DivergencePoint<T>&>(p);
	T sum4_p = 0,      sum4_q = 0;                 // Sum for every 4 nucleotides
        double total_sum_p = 0, total_sum_q = 0;       // Total running sum of all nucleotides
	double outer_sum_p = 0, outer_sum_q = 0;       // Prior K-mer sum
	for (int i = 0; i < points.size(); i++) { // Compute divergence for P and Q simultaneously
		sum4_p += points[i];
		sum4_q += d.points[i];
		if (i % 4 == 3) { //finished counting word, now compute probabilities
			double inner_sum_p = 0;        // Sum of p(X|Y) * log(p(X|Y) / q(X|Y))
			double inner_sum_q = 0;        // Sum of q(X|Y) * log(q(X|Y) / p(X|Y))
			for (int j = i - 3; j <= i; j++) {
				double conditional_p =   points[j] / sum4_p;
				double conditional_q = d.points[j] / sum4_q;
				double lg = log(conditional_p) - log(conditional_q);
				inner_sum_p +=      conditional_p * lg;
				inner_sum_q += -1 * conditional_q * lg;
			}
			outer_sum_p += sum4_p * inner_sum_p;
			outer_sum_q += sum4_q * inner_sum_q;

			total_sum_p += sum4_p;
			total_sum_q += sum4_q;
			sum4_p = 0;
			sum4_q = 0;
		}
	}
	double left = outer_sum_p / total_sum_p;
	double right = outer_sum_q / total_sum_q;
	return (left + right) / 2.0;
}

template<class T>
uint64_t DivergencePoint<T>::getPseudoMagnitude() const
{
	return mag;
}


template<class T>
uint64_t DivergencePoint<T>::getRealMagnitude() const
{
	return mag - points.size();
}

#ifndef HEADER_HACK
template class DivergencePoint<int>;
template class DivergencePoint<double>;
template class DivergencePoint<uint64_t>;
template class DivergencePoint<uint32_t>;
template class DivergencePoint<uint16_t>;
template class DivergencePoint<uint8_t>;
#endif
```
