## Supplementary Data 1 for "FASTCAR: Rapid alignment-free prediction of sequence alignment identity scores using self-supervised general linear models": DivergencePoint.h

```
/* -*- C++ -*-
 *
 * DivergencePoint.h
 *
 * Author: Benjamin T James
 *
 * Header for most often used k-mer histogram type
 */
#ifndef DIVERGENCE_POINT_H
#define DIVERGENCE_POINT_H
#include "Point.h"
#include <vector>
template<class T>
class DivergencePoint : public Point<T> {
public:
	DivergencePoint(const std::vector<T>& pts, uint64_t len);
	DivergencePoint(unsigned int size);
	~DivergencePoint() { points.clear(); }
	void operator*=(double d);
	void operator/=(double d);
	uint64_t operator-(const Point<T>& p) const;
	bool operator<(Point<T>& p) const;
	void operator+=(Point<T>& p);
	void set(Point<T>& p);
	void display() const;
	void zero();
	void addOne();
	void subOne();
	double prob_under(Point<T>& p) const;
	uint64_t getRealMagnitude() const;
	uint64_t getPseudoMagnitude() const;
//	T magnitude() const { return getRealMagnitude(); };
	double distance_k1(const Point<T>& p) const;
	double get_stddev() const { return s_dev; };
	DivergencePoint* clone() const {
		auto d = new DivergencePoint(points, to_delete);
		d->set_header(Point<T>::get_header());
		d->set_id(get_id());
		d->set_length(get_length());
		d->set_stddev(get_stddev());
		d->set_data_str(Point<T>::get_data_str());
		return d;
	}
	DivergencePoint* create() const {
		return new DivergencePoint(points.size());
	}
	Point<double>* create_double() const {
		vector<double> v;
		for (auto val : points) {
			v.push_back(val);
		}
		return new DivergencePoint<double>(v, nucl_length);
	}
	void set_arg_to_this_d(Point<double>& p) const {
		DivergencePoint<double>& c = dynamic_cast< DivergencePoint<double>&>(p);
		for (int i = 0; i < points.size(); i++) {
			c.points[i] = points[i];
		}
		c.set_id(id);
	};


	bool is_to_delete() const {
		return to_delete;
	}
	void set_to_delete(bool b) {
		to_delete = b;
	}
	double divergence(Point<T>& p) const;
	double distance_d(Point<double>& p) const;
	uint64_t distance(const Point<T>& p) const;
	const vector<T>& get_data() const { return points; }
	void set_id(uintmax_t c_id) { id = c_id; };
	const uintmax_t get_id() const { return id; };

	void set_length(unsigned long len) { nucl_length = len; };
	void set_stddev(double s_dev_) { s_dev = s_dev_; };
	unsigned long get_length() const { return nucl_length; };
	unsigned long size() const { return points.size(); };
	std::vector<T> points;

private:
	uintmax_t mag;
	bool to_delete;
	uint64_t id;
	uint64_t nucl_length;
	double s_dev;
};

#endif
```
