## Supplementary Data 1 for "FASTCAR: Rapid alignment-free prediction of sequence alignment identity scores using self-supervised general linear models": Feature.cpp

```
/* -*- C++ -*-
 *
 * Feature.cpp
 *
 * Author: Benjamin T James
 *
 * Raw feature methods are here.
 * Duplicates exist of many of the functions
 * exist because I was lazy and couldn't get
 * anonymous functions to work with the hashing
 */

#include <map>
#include <iterator>
#include <algorithm>
#include "Feature.h"

using namespace std;

template<class T>
Feature<T>::Feature(const Feature<T>& feat_) : k(feat_.get_k())
{
	flags = feat_.get_flags();
	mins = feat_.get_mins();
	maxs = feat_.get_maxs();
	is_sims = feat_.get_sims();
	combos = feat_.get_combos();
	lookup = feat_.get_lookup();
	is_finalized = feat_.get_finalized();
	do_save = false;
	auto freverse = [](int idx, int k) {
		int sum = 0;
		for (int i = 0; i < k; i++) {
			int rem = idx % 4;
			idx /= 4;
			sum = 4 * sum + rem;

		}
		return sum;
	};
	auto freverse_complement = [](int idx, int k) {
		std::vector<int> v;
		for (int i = 0; i < k; i++) {
			v.push_back(3 - idx % 4);
			idx /= 4;
		}
		int sum = 0;
		for (auto val : v) {
			sum = 4 * sum + val;
		}
		return sum;
	};
	for (auto f : lookup) {
		raw_funcs.push_back(get_func(f));
	}
}


// void * __gxx_personality_v0=0;
// void * _Unwind_Resume =0;
template<class T>
Feature<T> Feature<T>::operator=(const Feature<T>& feat_)
{
	k = feat_.get_k();
	flags = feat_.get_flags();
	mins = feat_.get_mins();
	maxs = feat_.get_maxs();
	is_sims = feat_.get_sims();
	combos = feat_.get_combos();
	lookup = feat_.get_lookup();
	is_finalized = feat_.get_finalized();
	do_save = false;
	auto freverse = [](int idx, int k) {
		int sum = 0;
		for (int i = 0; i < k; i++) {
			int rem = idx % 4;
			idx /= 4;
			sum = 4 * sum + rem;

		}
		return sum;
	};
	auto freverse_complement = [](int idx, int k) {
		std::vector<int> v;
		for (int i = 0; i < k; i++) {
			v.push_back(3 - idx % 4);
			idx /= 4;
		}
		int sum = 0;
		for (auto val : v) {
			sum = 4 * sum + val;
		}
		return sum;
	};
	for (auto f : lookup) {
		raw_funcs.push_back(get_func(f));
	}
	return *this;
}

template<class T>
void Feature<T>::add_feature(uint64_t f_flags, Combo combo)
{
	//	cout << "Adding combo " << f_flags << endl;
	if (combo != Combo::xy && combo != Combo::x2y && combo != Combo::xy2 && combo != Combo::x2y2) {
		throw "invalid combo";
	}
	vector<int> indices;
	for (uint64_t f = 1; f <= f_flags; f = (f << 1)) {
		// it is in the new parameter but not currently in store
		if ((f_flags & f) != 0) {
			if ((flags & f) == 0) {
				lookup.push_back(f);
				raw_funcs.push_back(get_func(f));
				//cout << "new single feature " << f << endl;
				mins.push_back(std::numeric_limits<double>::max());
				maxs.push_back(std::numeric_limits<double>::min());
				is_sims.push_back(feat_is_sim(f));
				is_finalized.push_back(false);
				flags |= f;
			}
			indices.push_back(index_of(f));
		}
	}
	combos.push_back(std::make_pair(combo, indices));
}

template<class T>
void Feature<T>::finalize()
{
	for (size_t i = 0; i < is_finalized.size(); i++) {
		is_finalized[i] = true;
	}
}
template<class T>
void Feature<T>::normalize_cache(vector<double> &cache) const
{
	for (size_t i = 0; i < lookup.size(); i++) {
		double val = (cache[i] - mins[i]) / (maxs[i] - mins[i]);

		// Hani Z. Girgis added this test
		if(isnan(val)){
			cerr << "Got NAN from max " << maxs[i] << " min " << mins[i] << endl;
			throw std::exception();
		}

		if (is_sims[i]) {
			cache[i] = val;
		} else {
			cache[i] = 1 - val;
		}
	}
}
template<class T>
vector<double> Feature<T>::compute_all_raw(Point<T> &p, Point<T> &q)
{
	vector<double> cache(lookup.size());
	uint64_t done = 0;
#ifdef FEATURE_OMP
#pragma omp parallel for
#endif
	for (size_t i = 0; i < lookup.size(); i++) {
		if ((lookup[i] & done) == 0) {
//			auto rres = get_func(lookup[i])(p, q);
			auto rres = raw_funcs[i](p, q);
			cache[i] = rres;
		}
	}
	return cache;
}

template<class T>
void Feature<T>::set_normal(uint64_t single_flag, double min_, double max_)
{
	int idx = index_of(single_flag);
	mins.at(idx) = min_;
	maxs.at(idx) = max_;
	is_finalized.at(idx) = true;
}

/*
template<class T>
vector<double> Feature<T>::get_raw(const vector<pair<Point<T>*,Point<T>*> > &vec, int index) const
{
	std::vector<double> results(vec.size(), 0);
	auto func = raw_funcs[index];

	#pragma omp parallel for
	for (size_t i = 0; i < vec.size(); i++) {
		results[i] = func(*vec[i].first, *vec[i].second);
	}

	double vmin, vmax;
        auto mm = std::minmax_element(results.begin(), results.end());
	vmin = *(mm.first);
	vmax = *(mm.second);
	for (auto &v : results) {
		v = (v - vmin) / (vmax - vmin);
		if (! is_sims[index]) {
			v = 1 - v;
		}
	}
	return results;
}
*/

template<class T>
void Feature<T>::normalize(const vector<pra<T> > &pairs)
{

	for (size_t i = 0; i < lookup.size(); i++) {
		double small = mins[i], big = maxs[i];
		if (lookup[i] == FEAT_ALIGN) {
			mins[i] = 0;
			maxs[i] = 1;
			continue;
		}
		if (is_finalized[i]) {
			continue;
		}
// #ifdef FEATURE_OMP
// #pragma omp parallel for reduction(min:small), reduction(max:big)
// #endif
		auto func = raw_funcs[i];//
		// get_func(lookup[i]);
		for (size_t j = 0; j < pairs.size(); j++) {
			double val = func(*pairs[j].first, *pairs[j].second);
			if (val < small) {
				small = val;
			}
			if (val > big) {
				big = val;
			}
		}

		mins[i] = small;
		maxs[i] = big;

		// Hani Z. Girgis added this tests
		if(abs(maxs[i] - mins[i]) <= 0.000000001){
			cerr << "Error of feature: " << feat_names().at(i) << ". ";
			cerr << "The maximum distance cannot be zero.";
			cerr << endl;
			throw std::exception();
		}

		if(isinf(maxs[i])){
			cerr << "Error of feature: " << feat_names().at(i) << ". ";
			cerr << "Maximum is " << maxs[i] << endl;
			throw std::exception();
		}

		if(isinf(mins[i])){
			cerr << "Error of feature: " << feat_names().at(i) << ". ";
			cerr << "Minimum is " << mins[i] << endl;
			throw std::exception();
		}

	}
};

template<class T>
vector<std::string> Feature<T>::feat_names()
{
	std::vector<std::string> vec;
	for (int i = 0; i < combos.size(); i++) {
		auto indices = combos[i].second;
		std::vector<std::string> names;
		for (auto s : indices) {
			names.push_back(feat_name(lookup[s]));
		}
		std::string str = "";
		auto combo = combos[i].first;
		if (combo == Combo::xy) {
			str = names[0];
			for (int j = 1; j < indices.size(); j++) {
				str += " * " + names[j];
			}
		} else if (combo == Combo::xy2 && indices.size() == 2) {
			str = names[0] + " * " + names[1] + "^2";
		} else if (combo == Combo::x2y && indices.size() == 2) {
			str = names[0] + "^2 * " + names[1];
		} else if (combo == Combo::x2y2) {
			str = names[0] + "^2";
			for (int j = 1; j < indices.size(); j++) {
				str += " * " + names[j] + "^2";
			}
		}
		vec.push_back(str);
	}
	return vec;
}

template<class T>
std::string Feature<T>::feat_name(uint64_t single_flag)
{
	if (single_flag == FEAT_ALIGN) {
		return "align";
	} else if (single_flag == FEAT_HELLINGER) {
		return "hellinger";
	} else if (single_flag == FEAT_MANHATTAN) {
		return "manhattan";
	} else if (single_flag == FEAT_EUCLIDEAN) {
		return "euclidean";
	} else if (single_flag == FEAT_CHI_SQUARED) {
		return "chi_squared";
	} else if (single_flag == FEAT_NORMALIZED_VECTORS) {
		return "normalized_vectors";
	} else if (single_flag == FEAT_HARMONIC_MEAN) {
		return "harmonic_mean";
	} else if (single_flag == FEAT_JEFFEREY_DIV) {
		return "jefferey_divergence";
	} else if (single_flag == FEAT_K_DIV) {
		return "k_divergence";
	} else if (single_flag == FEAT_PEARSON_COEFF) {
		return "pearson";
	} else if (single_flag == FEAT_SQCHORD) {
		return "squared_chord";
	} else if (single_flag == FEAT_KL_COND) {
		return "kl_conditional";
	} else if (single_flag == FEAT_MARKOV) {
		return "markov";
	} else if (single_flag == FEAT_INTERSECTION) {
		return "intersection";
	} else if (single_flag == FEAT_RRE_K_R) {
		return "rre_k_r";
	} else if (single_flag == FEAT_D2z) {
		return "d2z";
	} else if (single_flag == FEAT_SIM_MM) {
		return "sim_mm";
	} else if (single_flag == FEAT_EUCLIDEAN_Z) {
		return "euclidean_z";
	} else if (single_flag == FEAT_EMD) {
		return "emd";
	} else if (single_flag == FEAT_SPEARMAN) {
		return "spearman";
	} else if (single_flag == FEAT_JACCARD) {
		return "jaccard";
	} else if (single_flag == FEAT_LENGTHD) {
		return "length_difference";
	} else if (single_flag == FEAT_D2s) {
		return "d2s";
	} else if (single_flag == FEAT_AFD) {
		return "afd";
	} else if (single_flag == FEAT_MISMATCH) {
		return "mismatch";
	} else if (single_flag == FEAT_CANBERRA) {
		return "canberra";
	} else if (single_flag == FEAT_KULCZYNSKI1) {
		return "kulczynski1";
	} else if (single_flag == FEAT_KULCZYNSKI2) {
		return "kulczynski2";
	} else if (single_flag == FEAT_SIMRATIO) {
		return "simratio";
	} else if (single_flag == FEAT_JENSEN_SHANNON) {
		return "jensen_shannon";
	} else if (single_flag == FEAT_D2_star) {
		return "d2_star";
	} else if (single_flag == FEAT_N2R) {
		return "n2r";
	} else if (single_flag == FEAT_N2RC) {
		return "n2rc";
	} else if (single_flag == FEAT_N2RRC) {
		return "n2rrc";
	} else {
		return "unknown";
	}
}

template<class T>
std::function<double(Point<T>&,Point<T>&)> Feature<T>::get_func_(uint64_t single_flag)
{
	std::function<double(Point<T>&,Point<T>&)> func = [&](Point<T>&,Point<T>&)->double {
		cerr << "Unknown single flag " << single_flag << endl;
		throw "Function not set";
	};
	if (single_flag == FEAT_ALIGN) {
		func = [&](Point<T>& a,Point<T>& b) {
			return align(a, b, atable);
		};
	} else if (single_flag == FEAT_HELLINGER) {
		func = hellinger;
	} else if (single_flag == FEAT_MANHATTAN) {
		func = manhattan;
	} else if (single_flag == FEAT_EUCLIDEAN) {
		func = euclidean;
	} else if (single_flag == FEAT_CHI_SQUARED) {
		func = chi_squared;
	} else if (single_flag == FEAT_NORMALIZED_VECTORS) {
		func = normalized_vectors;
	} else if (single_flag == FEAT_HARMONIC_MEAN) {
		func = harmonic_mean;
	} else if (single_flag == FEAT_JEFFEREY_DIV) {
		func = jefferey_divergence;
	} else if (single_flag == FEAT_K_DIV) {
		func = k_divergence;
	} else if (single_flag == FEAT_PEARSON_COEFF) {
		func = pearson;
	} else if (single_flag == FEAT_SQCHORD) {
		func = squaredchord;
	} else if (single_flag == FEAT_KL_COND) {
		func = kl_conditional;
	} else if (single_flag == FEAT_MARKOV) {
		func = markov;
	} else if (single_flag == FEAT_INTERSECTION) {
		func = intersection;
	} else if (single_flag == FEAT_RRE_K_R) {
		func = rre_k_r;
	} else if (single_flag == FEAT_D2z) {
		func = d2z;
	} else if (single_flag == FEAT_SIM_MM) {
		func = sim_mm;
	} else if (single_flag == FEAT_EUCLIDEAN_Z) {
		func = euclidean_z;
	} else if (single_flag == FEAT_EMD) {
		func = emd;
	} else if (single_flag == FEAT_SPEARMAN) {
		func = spearman;
	} else if (single_flag == FEAT_JACCARD) {
		func = jaccard;
	} else if (single_flag == FEAT_LENGTHD) {
		func = length_difference;
	} else if (single_flag == FEAT_D2s) {
		func = d2s;
	} else if (single_flag == FEAT_AFD) {
		func = afd;
	} else if (single_flag == FEAT_MISMATCH) {
		func = mismatch;
	} else if (single_flag == FEAT_CANBERRA) {
		func = canberra;
	} else if (single_flag == FEAT_KULCZYNSKI1) {
		func = kulczynski1;
	} else if (single_flag == FEAT_KULCZYNSKI2) {
		func = kulczynski2;
	} else if (single_flag == FEAT_SIMRATIO) {
		func = simratio;
	} else if (single_flag == FEAT_JENSEN_SHANNON) {
		func = [&](Point<T>&a, Point<T>&b) { return jensen_shannon(a, b); };
	} else if (single_flag == FEAT_D2_star) {
		func = d2_star;
	} else if (single_flag == FEAT_N2R) {
		func = [&](Point<T>&a, Point<T>&b) { return n2r(a, b); };
	} else if (single_flag == FEAT_N2RC) {
		func = [&](Point<T>&a, Point<T>&b) { return n2rc(a, b); };
	} else if (single_flag == FEAT_N2RRC) {
		func = [&](Point<T>&a, Point<T>&b) { return n2rrc(a, b); };
	}
	return func;
}

template<class T>
std::function<double(Point<T>&,Point<T>&)> Feature<T>::get_func(uint64_t single_flag)
{
//	cout << "SINGLE FLAG: " << single_flag << ": " << Feature<T>::log2(single_flag) << endl;
	if (!do_save) {
		return get_func_(single_flag);
	} else if (single_flag == FEAT_HELLINGER) {
		return [&](Point<T>& a, Point<T>& b) { return c_hellinger(a,b); };
	} else if (single_flag == FEAT_MANHATTAN) {
		return [&](Point<T>& a, Point<T>& b) { return c_manhattan(a,b); };
	} else if (single_flag == FEAT_EUCLIDEAN) {
		return [&](Point<T>& a, Point<T>& b) { return c_euclidean(a,b); };
	} else if (single_flag == FEAT_CHI_SQUARED) {
		return [&](Point<T>& a, Point<T>& b) { return c_chi_squared(a,b); };
	} else if (single_flag == FEAT_NORMALIZED_VECTORS) {
		return [&](Point<T>& a, Point<T>& b) { return c_normalized_vectors(a,b); };
	} else if (single_flag == FEAT_HARMONIC_MEAN) {
		return [&](Point<T>& a, Point<T>& b) { return c_harmonic_mean(a,b); };
	} else if (single_flag == FEAT_JEFFEREY_DIV) {
		return [&](Point<T>& a, Point<T>& b) { return c_jefferey_divergence(a,b); };
	} else if (single_flag == FEAT_K_DIV) {
		return [&](Point<T>& a, Point<T>& b) { return c_k_divergence(a,b); };
	} else if (single_flag == FEAT_PEARSON_COEFF) {
		return [&](Point<T>& a, Point<T>& b) { return c_pearson(a,b); };
	} else if (single_flag == FEAT_SQCHORD) {
		return [&](Point<T>& a, Point<T>& b) { return c_squaredchord(a,b); };
	} else if (single_flag == FEAT_KL_COND) {
		return [&](Point<T>& a, Point<T>& b) { return c_kl_conditional(a,b); };
	} else if (single_flag == FEAT_MARKOV) {
		return [&](Point<T>& a, Point<T>& b) { return c_markov(a,b); };
	} else if (single_flag == FEAT_INTERSECTION) {
		return [&](Point<T>& a, Point<T>& b) { return c_intersection(a,b); };
	} else if (single_flag == FEAT_RRE_K_R) {
		return [&](Point<T>& a, Point<T>& b) { return c_rre_k_r(a,b); };
	} else if (single_flag == FEAT_D2z) {
		return [&](Point<T>& a, Point<T>& b) { return c_d2z(a,b); };
	} else if (single_flag == FEAT_SIM_MM) {
		return [&](Point<T>& a, Point<T>& b) { return c_sim_mm(a,b); };
	} else if (single_flag == FEAT_EUCLIDEAN_Z) {
		return [&](Point<T>& a, Point<T>& b) { return c_euclidean_z(a,b); };
	} else if (single_flag == FEAT_EMD) {
		return [&](Point<T>& a, Point<T>& b) { return c_emd(a,b); };
	} else if (single_flag == FEAT_SPEARMAN) {
		return [&](Point<T>& a, Point<T>& b) { return c_spearman(a,b); };
	} else if (single_flag == FEAT_JACCARD) {
		return [&](Point<T>& a, Point<T>& b) { return c_jaccard(a,b); };
	} else if (single_flag == FEAT_LENGTHD) {
		return [&](Point<T>& a, Point<T>& b) { return length_difference(a,b); };
	} else if (single_flag == FEAT_D2s) {
		return [&](Point<T>& a, Point<T>& b) { return c_d2s(a,b); };
	} else if (single_flag == FEAT_AFD) {
		return [&](Point<T>& a, Point<T>& b) { return c_afd(a,b); };
	} else if (single_flag == FEAT_MISMATCH) {
		return [&](Point<T>& a, Point<T>& b) { return c_mismatch(a,b); };
	} else if (single_flag == FEAT_CANBERRA) {
		return [&](Point<T>& a, Point<T>& b) { return c_canberra(a,b); };
	} else if (single_flag == FEAT_KULCZYNSKI1) {
		return [&](Point<T>& a, Point<T>& b) { return c_kulczynski1(a,b); };
	} else if (single_flag == FEAT_KULCZYNSKI2) {
		return [&](Point<T>& a, Point<T>& b) { return c_kulczynski2(a,b); };
	} else if (single_flag == FEAT_SIMRATIO) {
		return [&](Point<T>& a, Point<T>& b) { return c_simratio(a,b); };
	} else if (single_flag == FEAT_JENSEN_SHANNON) {
		return [&](Point<T>& a, Point<T>& b) { return c_jensen_shannon(a,b); };
	} else if (single_flag == FEAT_D2_star) {
		return [&](Point<T>& a, Point<T>& b) { return c_d2_star(a,b); };
	} else if (single_flag == FEAT_N2R) {
		return [&](Point<T>& a, Point<T>& b) { return c_n2r(a,b); };
	} else if (single_flag == FEAT_N2RC) {
		return [&](Point<T>& a, Point<T>& b) { return c_n2rc(a,b); };
	} else if (single_flag == FEAT_N2RRC) {
		return [&](Point<T>& a, Point<T>& b) { return c_n2rrc(a,b); };
	} else {
		throw "err";
	}
	return get_func_(single_flag);
}
template<class T>
vector<int> Feature<T>::multi_to_log(uint64_t multi)
{
	vector<int> ret;
	for (uint64_t i = 1; i <= 33; i++) {
		if (((1UL << i) & multi) != 0) {
			ret.push_back(i);
		}
	}
	return ret;
}

template<class T>
bool Feature<T>::feat_is_sim(uint64_t single_flag) const
{
	bool unknown = true;
	bool is_sim = true;
	switch (single_flag) {
	case FEAT_ALIGN:
		is_sim = true;
		break;
	case FEAT_HELLINGER:
		is_sim = false;
		break;
	case FEAT_MANHATTAN:
		is_sim = false;
		break;
	case FEAT_EUCLIDEAN:
		is_sim = false;
		break;
	case FEAT_CHI_SQUARED:
		is_sim = false;
		break;
	case FEAT_NORMALIZED_VECTORS:
		is_sim = true;
		break;
	case FEAT_HARMONIC_MEAN:
		is_sim = true;
		break;
	case FEAT_JEFFEREY_DIV:
		is_sim = false;
		break;
	case FEAT_K_DIV:
		is_sim = false;
		break;
	case FEAT_PEARSON_COEFF:
		is_sim = true;
		break;
	case FEAT_SQCHORD:
		is_sim = false;
		break;
	case FEAT_KL_COND:
		is_sim = false;
		break;
	case FEAT_MARKOV:
		is_sim = true;
		break;
	case FEAT_INTERSECTION:
		is_sim = true;
		break;
	case FEAT_RRE_K_R:
		is_sim = false;
		break;
	case FEAT_D2z:
		is_sim = true;
		break;
	case FEAT_SIM_MM:
		is_sim = true;//probably yes
		break;
	case FEAT_EUCLIDEAN_Z:
		is_sim = false;
		break;
	case FEAT_EMD:
		is_sim = false;
		break;
	case FEAT_SPEARMAN:
		is_sim = false; // Hani Z. Girgis modified the boolean
		//is_sim = true;
		break;
	case FEAT_JACCARD:
		is_sim = true;
		break;
	case FEAT_LENGTHD:
		is_sim = false;
		break;
	case FEAT_D2s:
		is_sim = true;
		break;
	case FEAT_AFD:
		is_sim = false;
		break;
	case FEAT_MISMATCH:
		is_sim = false;
		break;
	case FEAT_CANBERRA:
		is_sim = false;
		break;
	case FEAT_KULCZYNSKI1:
		is_sim = false;
		break;
	case FEAT_KULCZYNSKI2:
		is_sim = true;
		break;
	case FEAT_SIMRATIO:
		is_sim = true;
		break;
	case FEAT_JENSEN_SHANNON:
		is_sim = false;
		break;
	case FEAT_D2_star:
		is_sim = true;
		break;
	case FEAT_N2R:
		is_sim = true;
		break;
	case FEAT_N2RC:
		is_sim = true;
		break;
	case FEAT_N2RRC:
		is_sim = true;
		break;
	default:
		cerr << "bad feature flag " << single_flag << " aka 2^" << log(single_flag) << endl;

		throw single_flag;
	}
	return is_sim;
}


template<class T>
double Feature<T>::c_kulczynski2(Point<T>& a, Point<T>& b) {

	auto aid = a.get_id();
	auto bid = b.get_id();
	auto tup = std::tuple<uintmax_t, uintmax_t, uint8_t>(aid, bid, Feature<T>::log2(FEAT_KULCZYNSKI2));
	if (ltable.find(tup) == ltable.end()) {
		double val = kulczynski2(a, b);
		ltable.insert({tup, val});
		return val;
	} else {
		return ltable.at(tup);
	}
}

template<class T>
double Feature<T>::kulczynski2(Point<T> &a, Point<T> &b)
{
	const DivergencePoint<T>& p = dynamic_cast<const DivergencePoint<T>&>(a);
	const DivergencePoint<T>& q = dynamic_cast<const DivergencePoint<T>&>(b);
	const auto N = p.points.size();
	uint64_t min_sum = 0;
	double ap = (double)p.getPseudoMagnitude() / N;
	double aq = (double)q.getPseudoMagnitude() / N;
	for (auto i = 0; i < N; i++) {
		min_sum += std::min(p.points[i], q.points[i]);
	}
	double coeff = N * (ap + aq) / (2 * ap * aq);
	return coeff * min_sum;
}
template<class T>
double Feature<T>::align(Point<T> &a, Point<T> &b, std::map<std::pair<uintmax_t, uintmax_t>, double> &atbl)
{
	auto ai = a.get_id();
	auto bi = b.get_id();
	std::pair<uintmax_t, uintmax_t> pr = ai < bi ? std::make_pair(ai, bi) : std::make_pair(bi, ai);
	auto res = atbl.find(pr);
	if (res == atbl.end()) {
		auto sa = a.get_data_str();
		auto sb = b.get_data_str();
		int la = sa.length();
		int lb = sb.length();
		GlobAlignE galign(sa.c_str(), 0, la-1,
				  sb.c_str(), 0, lb-1,
				  1, -1, 2, 1);
		double val = galign.getIdentity();
#pragma omp critical
		atbl[pr] = val;
		return val;
	} else {
		return res->second;
	}
}

template<class T>
double Feature<T>::c_squaredchord(Point<T>& a, Point<T>& b) {

	auto aid = a.get_id();
	auto bid = b.get_id();
	auto tup = std::tuple<uintmax_t, uintmax_t, uint8_t>(aid, bid, Feature<T>::log2(FEAT_SQCHORD));
	if (ltable.find(tup) == ltable.end()) {
		double val = squaredchord(a, b);
		ltable.insert({tup, val});
		return val;
	} else {
		return ltable.at(tup);
	}
}

template<class T>
double Feature<T>::squaredchord(Point<T> &a, Point<T> &b)
{
	const DivergencePoint<T>& p = dynamic_cast<const DivergencePoint<T>&>(a);
	const DivergencePoint<T>& q = dynamic_cast<const DivergencePoint<T>&>(b);
	const auto N = p.points.size();
	double sum = 0;
	for (auto i = 0; i < N; i++) {
		sum += p.points[i] + q.points[i] - 2 * sqrt(p.points[i] * q.points[i]);
	}
	return sum;
}

template<class T>
double Feature<T>::c_intersection(Point<T>& a, Point<T>& b) {

	auto aid = a.get_id();
	auto bid = b.get_id();
	auto tup = std::tuple<uintmax_t, uintmax_t, uint8_t>(aid, bid, Feature<T>::log2(FEAT_INTERSECTION));
	if (ltable.find(tup) == ltable.end()) {
		double val = intersection(a, b);
		ltable.insert({tup, val});
		return val;
	} else {
		return ltable.at(tup);
	}
}

template<class T>
double Feature<T>::intersection(Point<T> &a, Point<T> &b)
{
	const DivergencePoint<T>& p = dynamic_cast<const DivergencePoint<T>&>(a);
	const DivergencePoint<T>& q = dynamic_cast<const DivergencePoint<T>&>(b);
	const auto N = p.points.size();
	uintmax_t dist = 0;
	uintmax_t mag = p.getPseudoMagnitude() + q.getPseudoMagnitude();
	#pragma omp simd
	for (auto i = 0; i < N; i++) {
		dist += 2 * std::min(p.points[i], q.points[i]);
	}

	return (double)dist / (double)mag;
}

template<class T>
double Feature<T>::c_pearson(Point<T>& a, Point<T>& b) {

	auto aid = a.get_id();
	auto bid = b.get_id();
	auto tup = std::tuple<uintmax_t, uintmax_t, uint8_t>(aid, bid, Feature<T>::log2(FEAT_PEARSON_COEFF));
	if (ltable.find(tup) == ltable.end()) {
		double val = pearson(a, b);
		ltable.insert({tup, val});
		return val;
	} else {
		return ltable.at(tup);
	}
}

template<class T>
double Feature<T>::pearson(Point<T> &a, Point<T> &b)
{
	const DivergencePoint<T>& p = dynamic_cast<const DivergencePoint<T>&>(a);
	const DivergencePoint<T>& q = dynamic_cast<const DivergencePoint<T>&>(b);
	const auto N = p.points.size();
	double dap = (double)p.getPseudoMagnitude() / N;
	double daq = (double)q.getPseudoMagnitude() / N;
	double dot = 0, np = 0, nq = 0;
	for (auto i = 0; i < N; i++) {
	        double dp = p.points[i] - dap;
	        double dq = q.points[i] - daq;
		np += dp * dp;
		nq += dq * dq;
		dot += dp * dq;
	}
	return dot / sqrt(np * nq);
}

template<class T>
double Feature<T>::c_simratio(Point<T>& a, Point<T>& b) {

	auto aid = a.get_id();
	auto bid = b.get_id();
	auto tup = std::tuple<uintmax_t, uintmax_t, uint8_t>(aid, bid, Feature<T>::log2(FEAT_SIMRATIO));
	if (ltable.find(tup) == ltable.end()) {
		double val = simratio(a, b);
		ltable.insert({tup, val});
		return val;
	} else {
		return ltable.at(tup);
	}
}

template<class T>
double Feature<T>::simratio(Point<T> &a, Point<T> &b)
{
	const DivergencePoint<T>& p = dynamic_cast<const DivergencePoint<T>&>(a);
	const DivergencePoint<T>& q = dynamic_cast<const DivergencePoint<T>&>(b);
	const auto N = p.points.size();
	uintmax_t dot = 0, norm2 = 0;
	for (auto i = 0; i < N; i++) {
		intmax_t diff = p.points[i] - q.points[i];
		dot += p.points[i] * q.points[i];
		norm2 += diff * diff;
	}
	return dot / (dot + sqrt(norm2));
}

template<class T>
double Feature<T>::c_manhattan(Point<T>& a, Point<T>& b) {

	auto aid = a.get_id();
	auto bid = b.get_id();
	auto tup = std::tuple<uintmax_t, uintmax_t, uint8_t>(aid, bid, Feature<T>::log2(FEAT_MANHATTAN));
	if (ltable.find(tup) == ltable.end()) {
		double val = manhattan(a, b);
		ltable.insert({tup, val});
		return val;
	} else {
		return ltable.at(tup);
	}
}

template<class T>
double Feature<T>::manhattan(Point<T>& a, Point<T>& b)
{
	const DivergencePoint<T>& p = dynamic_cast<const DivergencePoint<T>&>(a);
	const DivergencePoint<T>& q = dynamic_cast<const DivergencePoint<T>&>(b);
	auto N = p.points.size();
	int sum = 0;
	#pragma omp simd
	for (auto i = 0; i < N; i++) {
		sum += p.points[i] > q.points[i] ? p.points[i] - q.points[i] : q.points[i] - p.points[i];
	}
//	std::cout << "manhattan: " << sum << std::endl;
	return sum;
}

template<class T>
double Feature<T>::length_difference(Point<T>& a, Point<T>& b)
{
	const DivergencePoint<T>& p = dynamic_cast<const DivergencePoint<T>&>(a);
	const DivergencePoint<T>& q = dynamic_cast<const DivergencePoint<T>&>(b);
	auto lp = p.get_length();
	auto lq = q.get_length();
	if (lp == 0 || lq == 0) {
		cerr << "lp: " << lp << " lq: " << lq << endl;
		throw 123;
	}
	auto ret = (lp > lq) ? (lp - lq) : (lq - lp);
//	std::cout << "length difference: " << ret << std::endl;
	return ret;
}


double neighbor(double *cp, double *cq, double ap, double aq, const size_t N)
{
	double sp = 0, sq = 0;
	#pragma omp simd
	for (auto i = 0; i < N; i++) {
		double dp = cp[i] - ap;
		double dq = cq[i] - aq;
		sp += dp * dp;
		sq += dq * dq;
	}
	sp = sqrt(sp / N);
	sq = sqrt(sq / N);
	double psum = 0, qsum = 0;
	#pragma omp simd
	for (auto i = 0; i < N; i++) {
		cp[i] = (cp[i] - ap) / sp;
		cq[i] = (cq[i] - aq) / sq;
		psum += cp[i] * cp[i];
		qsum += cq[i] * cq[i];
	}
	double total = 0;
	psum = sqrt(psum);
	qsum = sqrt(qsum);
	#pragma omp simd
	for (auto i = 0; i < N; i++) {
		cp[i] /= psum;
		cq[i] /= qsum;
		total += cp[i] * cq[i];
	}
	return total;
}

template<class T>
double Feature<T>::c_n2rrc(Point<T>& a, Point<T>& b) {

	auto aid = a.get_id();
	auto bid = b.get_id();
	auto tup = std::tuple<uintmax_t, uintmax_t, uint8_t>(aid, bid, Feature<T>::log2(FEAT_N2RRC));
	if (ltable.find(tup) == ltable.end()) {
		double val = n2rrc(a, b);
		ltable.insert({tup, val});
		return val;
	} else {
		return ltable.at(tup);
	}
}

template<class T>
double Feature<T>::n2rrc(Point<T>& a, Point<T>& b) const
{
	if(!Util::isDna){
		cerr << "n2rrc cannot be calculated on protein sequences." << endl;
		throw std::exception();
	}

	const DivergencePoint<T>& p = dynamic_cast<const DivergencePoint<T>&>(a);
	const DivergencePoint<T>& q = dynamic_cast<const DivergencePoint<T>&>(b);
	const auto N = p.points.size();
	double *cp = new double[N];
	double *cq = new double[N];
	double ap = 0, aq = 0;
	for (auto i = 0; i < N; i++) {
		int j = reverse.at(i);
		int h = reverse_complement.at(i);
		cp[i] = p.points[h] + p.points[i] + p.points[j];
		cq[i] = q.points[h] + q.points[i] + q.points[j];
		ap += cp[i];
		aq += cq[i];
	}
	ap /= N;
	aq /= N;
	double total = neighbor(cp, cq, ap, aq, N);
	delete[] cp;
	delete[] cq;
//	std::cout << "n2rrc: " << total << std::endl;
	return total;
}

template<class T>
double Feature<T>::c_jensen_shannon(Point<T>& a, Point<T>& b) {

	auto aid = a.get_id();
	auto bid = b.get_id();
	auto tup = std::tuple<uintmax_t, uintmax_t, uint8_t>(aid, bid, Feature<T>::log2(FEAT_JENSEN_SHANNON));
	if (ltable.find(tup) == ltable.end()) {
		double val = jensen_shannon(a, b);
		ltable.insert({tup, val});
		return val;
	} else {
		return ltable.at(tup);
	}
}

template<class T>
double Feature<T>::jensen_shannon(Point<T> &a, Point<T> &b) const
{
	const DivergencePoint<T>& p = dynamic_cast<const DivergencePoint<T>&>(a);
	const DivergencePoint<T>& q = dynamic_cast<const DivergencePoint<T>&>(b);
	uint64_t mp = p.getPseudoMagnitude();
	uint64_t mq = q.getPseudoMagnitude();
	double sum = 0;
	const auto N = p.points.size();
    #pragma omp simd reduction(+:sum)
	for (auto i = 0; i < N; i++) {
		double pp = (double)p.points[i] / mp;
		double pq = (double)q.points[i] / mq;
		double avg = 0.5 * (pp + pq);
		#ifndef USETBL
		double lp = // tbl[(int)(coeff * pp / avg)];
			log(pp / avg);
		double lq = // tbl[(int)(coeff * pq / avg)];
			log(pq / avg);
		#else
		double lp = tbl[(int)(coeff * pp / avg)];
		double lq = tbl[(int)(coeff * pq / avg)];
		#endif
	        sum += pp * lp + pq * lq;
	}
	return sum / 2;
}

template<class T>
double Feature<T>::c_rre_k_r(Point<T>& a, Point<T>& b) {

	auto aid = a.get_id();
	auto bid = b.get_id();
	auto tup = std::tuple<uintmax_t, uintmax_t, uint8_t>(aid, bid, Feature<T>::log2(FEAT_RRE_K_R));
	if (ltable.find(tup) == ltable.end()) {
		double val = rre_k_r(a, b);
		ltable.insert({tup, val});
		return val;
	} else {
		return ltable.at(tup);
	}
}

// This statistics uses conditional probability
// Modified by Hani Z. Girgis on Oct 7 2018 to enable processing protein sequences
template<class T>
double Feature<T>::rre_k_r(Point<T>& a, Point<T>& b)
{
	const DivergencePoint<T>& p = dynamic_cast<const DivergencePoint<T>&>(a);
	const DivergencePoint<T>& q = dynamic_cast<const DivergencePoint<T>&>(b);
	const auto N = p.points.size();
	const auto A = Util::getAlphabetSize();

	double op = 0, oq = 0;
	const double l4 = log(A);
	uint64_t sum4_p = 0, sum4_q = 0;

	for (auto i = 0; i < N; i++) {
		sum4_p += p.points[i];
		sum4_q += q.points[i];
		if (i % A == (A-1)) {
			double inner_sum_p = 0;
			double inner_sum_q = 0;
			for (auto j = i - (A-1); j <= i; j++) {
				double conditional_p = (double)p.points[j] / sum4_p;
				double conditional_q = (double)q.points[j] / sum4_q;
				double avg = 0.5 * (conditional_p + conditional_q);
				inner_sum_p += (double)(p.points[j])
					* log(conditional_p / avg) / sum4_p;
				inner_sum_q += (double)(q.points[j])
					* log(conditional_q / avg) / sum4_q;
			}
			op += inner_sum_p;
			oq += inner_sum_q;
			sum4_p = 0;
			sum4_q = 0;
		}
	}

    double val = 0.5 * (op + oq);
	return val;
}


template<class T>
double Feature<T>::c_hellinger(Point<T>& a, Point<T>& b) {
	auto aid = a.get_id();
	auto bid = b.get_id();
	auto tup = std::tuple<uintmax_t, uintmax_t, uint8_t>(aid, bid, Feature<T>::log2(FEAT_HELLINGER));
	if (ltable.find(tup) == ltable.end()) {
		double val = hellinger(a, b);
		ltable.insert({tup, val});
		return val;
	} else {
		return ltable.at(tup);
	}
}

template<class T>
double Feature<T>::hellinger(Point<T>& a, Point<T>& b)
{
	const DivergencePoint<T>& p = dynamic_cast<const DivergencePoint<T>&>(a);
	const DivergencePoint<T>& q = dynamic_cast<const DivergencePoint<T>&>(b);
	const auto N = p.points.size();
	double ap = (double)p.getPseudoMagnitude() / N;
	double aq = (double)q.getPseudoMagnitude() / N;
	double sum = 0;
	for (auto i = 0; i < N; i++) {
		double diff = sqrt(p.points[i] / ap) - sqrt(q.points[i] / aq);
		sum += diff * diff;
	}
	return sqrt(2 * sum);
}

template<class T>
double Feature<T>::c_euclidean(Point<T>& a, Point<T>& b) {

	auto aid = a.get_id();
	auto bid = b.get_id();
	auto tup = std::tuple<uintmax_t, uintmax_t, uint8_t>(aid, bid, Feature<T>::log2(FEAT_EUCLIDEAN));
	if (ltable.find(tup) == ltable.end()) {
		double val = euclidean(a, b);
		ltable.insert({tup, val});
		return val;
	} else {
		return ltable.at(tup);
	}
}

template<class T>
double Feature<T>::euclidean(Point<T>& a, Point<T>& b)
{
	const DivergencePoint<T>& p = dynamic_cast<const DivergencePoint<T>&>(a);
	const DivergencePoint<T>& q = dynamic_cast<const DivergencePoint<T>&>(b);
	const auto N = p.points.size();
	uintmax_t sum = 0;
	for (auto i = 0; i < N; i++) {
		auto diff = p.points[i] - q.points[i];
		sum += diff * diff;
	}
	return sqrt(sum);
}

template<class T>
double Feature<T>::c_chi_squared(Point<T>& a, Point<T>& b) {

	auto aid = a.get_id();
	auto bid = b.get_id();
	auto tup = std::tuple<uintmax_t, uintmax_t, uint8_t>(aid, bid, Feature<T>::log2(FEAT_CHI_SQUARED));
	if (ltable.find(tup) == ltable.end()) {
		double val = chi_squared(a, b);
		ltable.insert({tup, val});
		return val;
	} else {
		return ltable.at(tup);
	}
}

template<class T>
double Feature<T>::chi_squared(Point<T>& a, Point<T>& b)
{
	const DivergencePoint<T>& p = dynamic_cast<const DivergencePoint<T>&>(a);
	const DivergencePoint<T>& q = dynamic_cast<const DivergencePoint<T>&>(b);
	const auto N = p.points.size();
	double sum = 0;
	for (auto i = 0; i < N; i++) {
		auto diff = p.points[i] - q.points[i];
		sum += (double)(diff * diff) / (p.points[i] + q.points[i]);
	}
	return sum;
}

template<class T>
double Feature<T>::c_normalized_vectors(Point<T>& a, Point<T>& b) {

	auto aid = a.get_id();
	auto bid = b.get_id();
	auto tup = std::tuple<uintmax_t, uintmax_t, uint8_t>(aid, bid, Feature<T>::log2(FEAT_NORMALIZED_VECTORS));
	if (ltable.find(tup) == ltable.end()) {
		double val = normalized_vectors(a, b);
		ltable.insert({tup, val});
		return val;
	} else {
		return ltable.at(tup);
	}
}

template<class T>
double Feature<T>::normalized_vectors(Point<T>& a, Point<T>& b)
{
	const DivergencePoint<T>& p = dynamic_cast<const DivergencePoint<T>&>(a);
	const DivergencePoint<T>& q = dynamic_cast<const DivergencePoint<T>&>(b);
	const auto N = p.points.size();
	uintmax_t sum = 0;
	uintmax_t d1 = 0, d2 = 0;
	for (auto i = 0; i < N; i++) {
		sum += p.points[i] * q.points[i];
		d1 += p.points[i] * p.points[i];
		d2 += q.points[i] * q.points[i];
	}
	return (double)sum / sqrt(d1 * d2);
}

template<class T>
double Feature<T>::c_harmonic_mean(Point<T>& a, Point<T>& b) {

	auto aid = a.get_id();
	auto bid = b.get_id();
	auto tup = std::tuple<uintmax_t, uintmax_t, uint8_t>(aid, bid, Feature<T>::log2(FEAT_HARMONIC_MEAN));
	if (ltable.find(tup) == ltable.end()) {
		double val = harmonic_mean(a, b);
		ltable.insert({tup, val});
		return val;
	} else {
		return ltable.at(tup);
	}
}

template<class T>
double Feature<T>::harmonic_mean(Point<T>& a, Point<T>& b)
{
	const DivergencePoint<T>& p = dynamic_cast<const DivergencePoint<T>&>(a);
	const DivergencePoint<T>& q = dynamic_cast<const DivergencePoint<T>&>(b);
	const auto N = p.points.size();
	double sum = 0;
	for (auto i = 0; i < N; i++) {
		double numer = p.points[i] * q.points[i];
		sum += numer / (p.points[i] + q.points[i]);
	}
	return 2 * sum;
}

template<class T>
double Feature<T>::c_jefferey_divergence(Point<T>& a, Point<T>& b) {

	auto aid = a.get_id();
	auto bid = b.get_id();
	auto tup = std::tuple<uintmax_t, uintmax_t, uint8_t>(aid, bid, Feature<T>::log2(FEAT_JEFFEREY_DIV));
	if (ltable.find(tup) == ltable.end()) {
		double val = jefferey_divergence(a, b);
		ltable.insert({tup, val});
		return val;
	} else {
		return ltable.at(tup);
	}
}

template<class T>
double Feature<T>::jefferey_divergence(Point<T>& a, Point<T>& b)
{
	const DivergencePoint<T>& p = dynamic_cast<const DivergencePoint<T>&>(a);
	const DivergencePoint<T>& q = dynamic_cast<const DivergencePoint<T>&>(b);
	uint64_t mp = p.getPseudoMagnitude();
	uint64_t mq = q.getPseudoMagnitude();
	double sum = 0;
	const auto N = p.points.size();
	for (auto i = 0; i < N; i++) {
		double pp = (double)p.points[i] / mp;
		double pq = (double)q.points[i] / mq;
		// if (q.points[i] == 0) {
		// 	cout << "Error for sequence " << q.get_header() << endl;
		// 	for (int j = 0; j < q.points.size(); j++) {
		// 		cout << q.points.at(j) << " ";
		// 	}
		// 	cout << endl;
		// 	exit(1);
		// }
		// if (p.points[i] == 0) {
		// 	cout << "Error for sequence " << p.get_header() << endl;
		// 	for (int j = 0; j < p.points.size(); j++) {
		// 		cout << (int)p.points.at(j) << " ";
		// 	}
		// 	cout << endl;
		// 	exit(1);
		// }
		double diff = pp - pq;
		//	cout << "pp: " << pp << " pq: " << pq << " pp/pq: " << pp / pq << endl;
	        sum += diff * log(pp / pq);
	}
	return sum;
}

template<class T>
double Feature<T>::c_k_divergence(Point<T>& a, Point<T>& b) {

	auto aid = a.get_id();
	auto bid = b.get_id();
	auto tup = std::tuple<uintmax_t, uintmax_t, uint8_t>(aid, bid, Feature<T>::log2(FEAT_K_DIV));
	if (ltable.find(tup) == ltable.end()) {
		double val = k_divergence(a, b);
		ltable.insert({tup, val});
		return val;
	} else {
		return ltable.at(tup);
	}
}

template<class T>
double Feature<T>::k_divergence(Point<T>& a, Point<T>& b)
{
	const DivergencePoint<T>& p = dynamic_cast<const DivergencePoint<T>&>(a);
	const DivergencePoint<T>& q = dynamic_cast<const DivergencePoint<T>&>(b);
	uint64_t mp = p.getPseudoMagnitude();
	uint64_t mq = q.getPseudoMagnitude();
	double sum = 0;
	const auto N = p.points.size();
	for (auto i = 0; i < N; i++) {
		double pp = (double)p.points[i] / mp;
		double pq = (double)q.points[i] / mq;
		double avg = 0.5 * (pp + pq);
	        sum += pp * log(pp / avg);
	}
	return sum;
}

template<class T>
double Feature<T>::c_kl_conditional(Point<T>& a, Point<T>& b) {

	auto aid = a.get_id();
	auto bid = b.get_id();
	auto tup = std::tuple<uintmax_t, uintmax_t, uint8_t>(aid, bid, Feature<T>::log2(FEAT_KL_COND));
	if (ltable.find(tup) == ltable.end()) {
		double val = kl_conditional(a, b);
		ltable.insert({tup, val});
		return val;
	} else {
		return ltable.at(tup);
	}
}

// Modified by Hani Z Girgis on Oct 7 2018.
template<class T>
double Feature<T>::kl_conditional(Point<T>& a, Point<T>& b)
{
	const DivergencePoint<T>& p = dynamic_cast<const DivergencePoint<T>&>(a);
	const DivergencePoint<T>& q = dynamic_cast<const DivergencePoint<T>&>(b);
	uint64_t sum4_p = 0,    sum4_q = 0;            // Sum for every 4 nucleotides or 22 a.a.
	double outer_sum_p = 0, outer_sum_q = 0;       // Prior K-mer sum

	const auto N = p.points.size();
	const auto A = Util::getAlphabetSize();

	for (auto i = 0; i < N; i++) {
		sum4_p += p.points[i];
		sum4_q += q.points[i];

		if (i % A == A-1) { //finished counting word, now compute probabilities
			double inner_sum_p = 0;        // Sum of p(X|Y) * log(p(X|Y) / q(X|Y))
			double inner_sum_q = 0;        // Sum of q(X|Y) * log(q(X|Y) / p(X|Y))
			for (auto j = i - (A-1); j <= i; j++) {
				double conditional_p = (double)p.points[j] / sum4_p;
				double conditional_q = (double)q.points[j] / sum4_q;
				double lg = log(conditional_p / conditional_q);
				inner_sum_p +=      conditional_p * lg;
				inner_sum_q += -1 * conditional_q * lg;
			}
			outer_sum_p += sum4_p * inner_sum_p;
			outer_sum_q += sum4_q * inner_sum_q;

			sum4_p = 0;
			sum4_q = 0;
		}
	}
	double left = outer_sum_p / p.getPseudoMagnitude();
	double right = outer_sum_q / q.getPseudoMagnitude();
	return (left + right) / 2.0;
}

template<class T>
double Feature<T>::c_markov(Point<T>& a, Point<T>& b) {

	auto aid = a.get_id();
	auto bid = b.get_id();
	auto tup = std::tuple<uintmax_t, uintmax_t, uint8_t>(aid, bid, Feature<T>::log2(FEAT_MARKOV));
	if (ltable.find(tup) == ltable.end()) {
		double val = markov(a, b);
		ltable.insert({tup, val});
		return val;
	} else {
		return ltable.at(tup);
	}
}

template<class T>
double Feature<T>::markov(Point<T>& a, Point<T>& b)
{
	const DivergencePoint<T>& q = dynamic_cast<const DivergencePoint<T>&>(a);
	const DivergencePoint<T>& p = dynamic_cast<const DivergencePoint<T>&>(b);
	double total = 0;       // Prior K-mer sum

	// Hani Z. Girgis modified this code on Oct 2 2018
	// to adapt this feature to proteins
	const auto N = p.points.size();
	const auto A = Util::getAlphabetSize();

	for (auto i = 0; i < N; i += A) {
		uint64_t psum = 0, qsum = 0;
		for (auto j = 0; j < A; j++) {
			psum += p.points[i+j];
			qsum += q.points[i+j];
		}
		double lpsum = log(psum);
		double lqsum = log(qsum);
		for (auto j = 0; j < A; j++) {
			total += (q.points[i+j]-1) * (log(p.points[i+j]) - lpsum);
			total += (p.points[i+j]-1) * (log(q.points[i+j]) - lqsum);
		}
    }

	return total / 2;
}

template<class T>
double Feature<T>::c_d2z(Point<T>& a, Point<T>& b) {

	auto aid = a.get_id();
	auto bid = b.get_id();
	auto tup = std::tuple<uintmax_t, uintmax_t, uint8_t>(aid, bid, Feature<T>::log2(FEAT_D2z));
	if (ltable.find(tup) == ltable.end()) {
		double val = d2z(a, b);
		ltable.insert({tup, val});
		return val;
	} else {
		return ltable.at(tup);
	}
}

template<class T>
double Feature<T>::d2z(Point<T>& a, Point<T>& b)
{
	const DivergencePoint<T>& p = dynamic_cast<const DivergencePoint<T>&>(a);
	const DivergencePoint<T>& q = dynamic_cast<const DivergencePoint<T>&>(b);
	double sum = 0;
	const auto N = p.points.size();
	double ap = (double)p.getPseudoMagnitude() / N;
	double aq = (double)q.getPseudoMagnitude() / N;
	double sp = p.get_stddev(), sq = q.get_stddev();
	for (auto i = 0; i < N; i++) {
		double pz = (p.points[i] - ap) / sp;
		double qz = (q.points[i] - aq) / sq;
		sum += pz * qz;
    }
	return sum;
}

template<class T>
double d_markov(Point<T>& a, Point<T>& b)
{
	const DivergencePoint<T>& q = dynamic_cast<const DivergencePoint<T>&>(b);
	return log(Feature<T>::markov(b, a) / Feature<T>::markov(b, b)) / q.getRealMagnitude();
}

template<class T>
double Feature<T>::c_sim_mm(Point<T>& a, Point<T>& b) {

	auto aid = a.get_id();
	auto bid = b.get_id();
	auto tup = std::tuple<uintmax_t, uintmax_t, uint8_t>(aid, bid, Feature<T>::log2(FEAT_SIM_MM));
	if (ltable.find(tup) == ltable.end()) {
		double val = sim_mm(a, b);
		ltable.insert({tup, val});
		return val;
	} else {
		return ltable.at(tup);
	}
}

template<class T>
double Feature<T>::sim_mm(Point<T>& a, Point<T>& b)
{
	return 1 - exp(0.5 * (d_markov(a, b) + d_markov(b, a)));
}

template<class T>
double Feature<T>::c_euclidean_z(Point<T>& a, Point<T>& b) {

	auto aid = a.get_id();
	auto bid = b.get_id();
	auto tup = std::tuple<uintmax_t, uintmax_t, uint8_t>(aid, bid, Feature<T>::log2(FEAT_EUCLIDEAN_Z));
	if (ltable.find(tup) == ltable.end()) {
		double val = euclidean_z(a, b);
		ltable.insert({tup, val});
		return val;
	} else {
		return ltable.at(tup);
	}
}

template<class T>
double Feature<T>::euclidean_z(Point<T>& a, Point<T>& b)
{
	const DivergencePoint<T>& p = dynamic_cast<const DivergencePoint<T>&>(a);
	const DivergencePoint<T>& q = dynamic_cast<const DivergencePoint<T>&>(b);
	double sum = 0;
	const auto N = p.points.size();
	double ap = (double)p.getPseudoMagnitude() / N;
	double aq = (double)q.getPseudoMagnitude() / N;
	double sp = p.get_stddev(), sq = q.get_stddev();
	for (auto i = 0; i < N; i++) {
		double pz = (p.points[i] - ap) / sp;
		double qz = (q.points[i] - aq) / sq;
		sum += (pz - qz) * (pz - qz);
        }
	return sqrt(sum);
}

template<class T>
double Feature<T>::c_emd(Point<T>& a, Point<T>& b) {

	auto aid = a.get_id();
	auto bid = b.get_id();
	auto tup = std::tuple<uintmax_t, uintmax_t, uint8_t>(aid, bid, Feature<T>::log2(FEAT_EMD));
	if (ltable.find(tup) == ltable.end()) {
		double val = emd(a, b);
		ltable.insert({tup, val});
		return val;
	} else {
		return ltable.at(tup);
	}
}

template<class T>
double Feature<T>::emd(Point<T>& a, Point<T>& b)
{
	const DivergencePoint<T>& p = dynamic_cast<const DivergencePoint<T>&>(a);
	const DivergencePoint<T>& q = dynamic_cast<const DivergencePoint<T>&>(b);
	const auto N = p.points.size();
	uintmax_t cp = 0, cq = 0;
	uintmax_t dist = 0;
	for (auto i = 0; i < N; i++) {
		cp += p.points[i];
		cq += q.points[i];
		dist += cp > cq ? cp - cq : cq - cp;
	}
	return (double)dist;
}

// Commented by Hani Z. Girgis
// template<class T>
// std::vector<size_t> tiedrank(const Point<T>& a)
// {
// 	const DivergencePoint<T>& p = dynamic_cast<const DivergencePoint<T>&>(a);
// 	const auto N = p.points.size();
// 	vector<size_t> ip(N, 0);
// 	std::iota(std::begin(ip), std::end(ip), 0);
// 	std::sort(std::begin(ip), std::end(ip), [&](size_t x, size_t y) {
// 			return p.points[x] < p.points[y];
// 	});

// 	for(auto elm : ip){
// 		cerr << elm << endl;
// 	}
// 	exit(9);
// 	return ip;
// }

// Added by Hani Z. Girgis
template<class T>
std::vector<double> tiedrank(const Point<T>& a){
	// Initialize multimap
	const DivergencePoint<T>& p = dynamic_cast<const DivergencePoint<T>&>(a);
	unsigned int n = p.points.size();

	std::multimap<T, double > mmap;
	for(unsigned i = 0; i < n; i++){
		mmap.insert(pair<T,double>( p.points[i] , i));
	}

	// Set ranks without ties
	int lastRank = 0;
	// std::multimap<T,double>::iterator
	for (auto it=mmap.begin(); it!=mmap.end(); ++it){
	 	(*it).second = ++lastRank;
	}

	for (auto it=mmap.begin(); it!=mmap.end(); it=mmap.upper_bound((*it).first)){
	 	auto ret = mmap.equal_range((*it).first);

    	// Calculate the average rank
    	double rankTotal = 0;
    	double count = 0;
    	for (auto it1=ret.first; it1 != ret.second; ++it1){
    		count++;
    		rankTotal += (*it1).second;
    	}

    	// Assign the average rank
    	double meanRank = rankTotal / count;
    	for (auto it1=ret.first; it1 != ret.second; ++it1){
    		(*it1).second = meanRank;
    		// cout << (*it).first << " => " << (*it1).second << endl;
    	}
	}

	std::vector<double> r(n, 0);
	for(unsigned int i = 0; i < n; i++){
		r[i] = mmap.find(p.points[i])->second;
	}

	// For testing
	// for(unsigned int i = 0; i < n; i++){
	// 	cout << r[i] << endl;
	// }

	return r;
}

template<class T>
double Feature<T>::c_spearman(Point<T>& a, Point<T>& b) {

	auto aid = a.get_id();
	auto bid = b.get_id();
	auto tup = std::tuple<uintmax_t, uintmax_t, uint8_t>(aid, bid, Feature<T>::log2(FEAT_SPEARMAN));
	if (ltable.find(tup) == ltable.end()) {
		double val = spearman(a, b);
		ltable.insert({tup, val});
		return val;
	} else {
		return ltable.at(tup);
	}
}

/*
template<class T>
double Feature<T>::spearman(Point<T>& a, Point<T>& b)
{
	const DivergencePoint<T>& p = dynamic_cast<const DivergencePoint<T>&>(a);
	const DivergencePoint<T>& q = dynamic_cast<const DivergencePoint<T>&>(b);
	const auto N = p.points.size();
	vector<size_t> ip(N, 0);
	vector<size_t> iq(N, 0);
	std::iota(std::begin(ip), std::end(ip), 0);
	std::iota(std::begin(iq), std::end(iq), 0);
	std::sort(std::begin(ip), std::end(ip), [&](size_t x, size_t y) {
			return p.points[x] < p.points[y];
		});

	double expected = (N+1) / 2.0;
	double cov = 0;
	double sp = 0;
	double sq = 0;
	for (auto i = 0; i < N; i++) {
		cov += (ip[i] - expected) * (iq[i] - expected);
		sp += (ip[i] - expected) * (ip[i] - expected);
		sq += (iq[i] - expected) * (iq[i] - expected);
    }

    cout << "N: "   << N   << endl;
    cout << "Cov: " << cov << endl;
    cout << "Sp: "  << sp  << endl;
    cout << "Sq: "  << sq  << endl;

    double results = (N * cov) / (sp * sq);

	return log(results);
}
*/


template<class T>
double Feature<T>::spearman(Point<T>& a, Point<T>& b)
{
	vector<double> ip = tiedrank(a);
	vector<double> iq = tiedrank(b);
	const auto N = iq.size();

	double expected = (N+1) / 2.0;
	double cov = 0;
	double sp = 0;
	double sq = 0;
	for (auto i = 0; i < N; i++) {
		cov += (ip[i] - expected) * (iq[i] - expected);
		sp += (ip[i] - expected) * (ip[i] - expected);
		sq += (iq[i] - expected) * (iq[i] - expected);
    }

    double result = 1 - ( cov / ( sqrt(sp) * sqrt(sq) ));
    // cerr << result << endl;
	return result;
}

template<class T>
double Feature<T>::c_jaccard(Point<T>& a, Point<T>& b) {

	auto aid = a.get_id();
	auto bid = b.get_id();
	auto tup = std::tuple<uintmax_t, uintmax_t, uint8_t>(aid, bid, Feature<T>::log2(FEAT_JACCARD));
	if (ltable.find(tup) == ltable.end()) {
		double val = jaccard(a, b);
		ltable.insert({tup, val});
		return val;
	} else {
		return ltable.at(tup);
	}
}

template<class T>
double Feature<T>::jaccard(Point<T>& a, Point<T>& b)
{
	const DivergencePoint<T>& p = dynamic_cast<const DivergencePoint<T>&>(a);
	const DivergencePoint<T>& q = dynamic_cast<const DivergencePoint<T>&>(b);
	const auto N = p.points.size();
	uint64_t sum = 0;
	for (auto i = 0; i < N; i++) {
		if (p.points[i] == q.points[i] && p.points[i] > 1) {
			sum++;
		}
	}
	return (double)sum / N;
}

template<class T>
double Feature<T>::c_d2s(Point<T>& a, Point<T>& b) {

	auto aid = a.get_id();
	auto bid = b.get_id();
	auto tup = std::tuple<uintmax_t, uintmax_t, uint8_t>(aid, bid, Feature<T>::log2(FEAT_D2s));
	if (ltable.find(tup) == ltable.end()) {
		double val = d2s(a, b);
		ltable.insert({tup, val});
		return val;
	} else {
		return ltable.at(tup);
	}
}

// Modified by Hani Z. Girgis on Oct 07 2018 to enable comparing protein sequences
// Note: This feature cannot be used if k is 1.
template<class T>
double Feature<T>::d2s(Point<T>& a, Point<T>& b)
{
	const DivergencePoint<T>& p = dynamic_cast<const DivergencePoint<T>&>(a);
	const DivergencePoint<T>& q = dynamic_cast<const DivergencePoint<T>&>(b);
	const auto N = p.points.size();
	const auto A = Util::getAlphabetSize();

	// Commented out by Hani Z Girgis and replaced by the line next to it.
	// const int k = (int)(log(N) / log(4));
	int k = a.getK();
	if(k==1){
		cerr << "D2s is skipped because it cannot be applied when k is 1.";
		cerr << endl;
		throw std::exception();
	}

	const auto p1 = p.get_1mers();
	const auto q1 = q.get_1mers();
	const double pmag = p.getPseudoMagnitude();
	const double qmag = q.getPseudoMagnitude();
	double sum = 0;
	for (size_t i = 0; i < N; i++) {
		double p1i = 1;
	    double q1i = 1;
	    size_t idx = i;
		for (int j = 0; j < k; j++) {
			int i1 = idx % A;
			idx /= A;
			p1i *= (double)p1[i1] / pmag;
			q1i *= (double)q1[i1] / qmag;
		}

		// Post conditions the probabilities
		if(p1i > 1 || p1i < 0){
			cerr << "p1i is too big or too small." << endl;
			throw std::exception();
		}
		if(q1i > 1 || q1i < 0){
			cerr << "pq1i is too big or too small." << endl;
			throw std::exception();
		}

		//double hp = p.points[i] - pmag * p1i;
		//double hq = q.points[i] - qmag * q1i;
		double hp = p.points[i] - (p.getRealMagnitude() * p1i + 1);
		double hq = q.points[i] - (q.getRealMagnitude() * q1i + 1);
		double denom = hypot(hp, hq);
		if (denom != 0 ) {
			sum += (hp * hq) / denom;
		}
	}
	return sum;
}

template<class T>
double Feature<T>::c_d2_star(Point<T>& a, Point<T>& b) {

	auto aid = a.get_id();
	auto bid = b.get_id();
	auto tup = std::tuple<uintmax_t, uintmax_t, uint8_t>(aid, bid, Feature<T>::log2(FEAT_D2_star));
	if (ltable.find(tup) == ltable.end()) {
		double val = d2_star(a, b);
		ltable.insert({tup, val});
		return val;
	} else {
		return ltable.at(tup);
	}
}

// Modified by Hani Z. Girgis on Oct 07 2018 to enable comparing protein sequences
// This method is rewriten based on the d2s code.
// Note: This feature cannot be used if k is 1.
template<class T>
double Feature<T>::d2_star(Point<T>& a, Point<T>& b)
{
	const DivergencePoint<T>& p = dynamic_cast<const DivergencePoint<T>&>(a);
	const DivergencePoint<T>& q = dynamic_cast<const DivergencePoint<T>&>(b);
	const auto N = p.points.size();
	const auto A = Util::getAlphabetSize();

	// Commented out by Hani Z Girgis and replaced by the line next to it.
	// const int k = (int)(log(N) / log(4));
	int k = a.getK();
	if(k==1){
		cerr << "D2_star cannot be applied when k is 1.";
		cerr << endl;
		throw std::exception();
	}

	const auto p1 = p.get_1mers();
	const auto q1 = q.get_1mers();
	const double pmag = p.getPseudoMagnitude();
	const double qmag = q.getPseudoMagnitude();
	const double pq_len = sqrt(p.getRealMagnitude() * q.getRealMagnitude());

	double sum = 0;
	for (size_t i = 0; i < N; i++) {
		double p1i  = 1;
	    double q1i  = 1;
	    double pq1i = 1;
	    size_t idx  = i;
		for (int j = 0; j < k; j++) {
			int i1 = idx % A;
			idx  /= A;
			p1i  *= (double) p1.at(i1) / pmag;
			q1i  *= (double) q1.at(i1) / qmag;
			pq1i *= ((double) p1.at(i1) + q1.at(i1)) / (pmag + qmag);
		}

		// Post conditions the probabilities
		if(p1i > 1 || p1i < 0){
			cerr << "p1i is too big or too small." << endl;
			throw std::exception();
		}
		if(q1i > 1 || q1i < 0){
			cerr << "pq1i is too big or too small." << endl;
			throw std::exception();
		}
		if(pq1i > 1 || pq1i < 0){
			cerr << "pq1i is too big or too small." << endl;
			throw std::exception();
		}

		double hp = p.points[i] - (p.getRealMagnitude() * p1i + 1);
		double hq = q.points[i] - (q.getRealMagnitude() * q1i + 1);
		double e = (p.getRealMagnitude() + q.getRealMagnitude()) * pq1i + 1;

		// Post conditions on the expected value
		if(e > p.getRealMagnitude() + q.getRealMagnitude()){
			cerr << "E is too big." << endl;
			throw std::exception();
		}
		if(e < 0){
			cerr << "E is too small." << endl;
			throw std::exception();
		}

		double denom =  e * pq_len;
		if (denom > 0) {
			sum += hp * hq / denom;
		}
	}

	return sum;
}

template<class T>
double Feature<T>::c_afd(Point<T>& a, Point<T>& b) {

	auto aid = a.get_id();
	auto bid = b.get_id();
	auto tup = std::tuple<uintmax_t, uintmax_t, uint8_t>(aid, bid, Feature<T>::log2(FEAT_AFD));
	if (ltable.find(tup) == ltable.end()) {
		double val = afd(a, b);
		ltable.insert({tup, val});
		return val;
	} else {
		return ltable.at(tup);
	}
}

// Modified by Hani Z. Girgis to enable processing protein sequences on Oct 9 2018.
// Must be used when k = 2; otherwise, an exception is thrown.
template<class T>
double Feature<T>::afd(Point<T>& a, Point<T>& b)
{
	const DivergencePoint<T>& p = dynamic_cast<const DivergencePoint<T>&>(a);
	const DivergencePoint<T>& q = dynamic_cast<const DivergencePoint<T>&>(b);
	const auto N = p.points.size();
	const auto A = Util::getAlphabetSize();

	const int k = a.getK();
	if(k != 2){
		cerr << "AFD cannot be calculated for k other than 2: Received: "  << k << endl;
		throw std::exception();
	}

	const auto p1 = p.get_1mers();
	const auto q1 = q.get_1mers();
	const auto pmag = p.getPseudoMagnitude();
	const auto qmag = q.getPseudoMagnitude();

	double sum = 0;
	const auto nMinusOne = N / A;
	const auto nMinusTwo = nMinusOne / A;
	int first_i = 0;
	for (auto i = 0; i < N; i += nMinusTwo) {
		// 16 iterations total, iterating through all 2-mers
		uint64_t psum = 0, qsum = 0;
		for (auto j = i; j < i + nMinusTwo; j++) {
			psum += p.points.at(j);
			qsum += q.points.at(j);
		}
		double x = (double)psum / p1.at(first_i / A);
		double y = (double)qsum / q1.at(first_i / A);
		first_i++;
		double diff = abs(x - y);
	    double unsquared = (diff * pow(1+diff, -14));
	    // Hani Z. Girgis modified this line
		// double unsquared = (diff * pow(1+diff, -2));

		sum += unsquared * unsquared;

		if(isinf(sum)){
			cerr << x << " " << y << " " << diff << " " << unsquared << endl;
			throw std::exception();
		}
	}

	return sum;
}

template<class T>
double Feature<T>::c_mismatch(Point<T>& a, Point<T>& b) {

	auto aid = a.get_id();
	auto bid = b.get_id();
	auto tup = std::tuple<uintmax_t, uintmax_t, uint8_t>(aid, bid, Feature<T>::log2(FEAT_MISMATCH));
	if (ltable.find(tup) == ltable.end()) {
		double val = mismatch(a, b);
		ltable.insert({tup, val});
		return val;
	} else {
		return ltable.at(tup);
	}
}

template<class T>
double Feature<T>::mismatch(Point<T>& a, Point<T>& b)
{
	const DivergencePoint<T>& p = dynamic_cast<const DivergencePoint<T>&>(a);
	const DivergencePoint<T>& q = dynamic_cast<const DivergencePoint<T>&>(b);
	const auto N = p.points.size();
	uint64_t sum = 0;
	#pragma omp simd
	for (auto i = 0; i < N; i++) {
		sum += (p.points[i] != q.points[i]);
	}
	return sum;
}

template<class T>
double Feature<T>::c_canberra(Point<T>& a, Point<T>& b) {

	auto aid = a.get_id();
	auto bid = b.get_id();
	auto tup = std::tuple<uintmax_t, uintmax_t, uint8_t>(aid, bid, Feature<T>::log2(FEAT_CANBERRA));
	if (ltable.find(tup) == ltable.end()) {
		double val = canberra(a, b);
		ltable.insert({tup, val});
		return val;
	} else {
		return ltable.at(tup);
	}
}

template<class T>
double Feature<T>::canberra(Point<T>& a, Point<T>& b)
{
	const DivergencePoint<T>& p = dynamic_cast<const DivergencePoint<T>&>(a);
	const DivergencePoint<T>& q = dynamic_cast<const DivergencePoint<T>&>(b);
	const auto N = p.points.size();
	double sum = 0;
	#pragma omp simd
	for (auto i = 0; i < N; i++) {
		auto numer = p.points[i] > q.points[i] ? p.points[i] - q.points[i] : q.points[i] - p.points[i];
		auto denom = p.points[i] + q.points[i];
		sum += (double)numer / denom;
	}
	return sum;
}

template<class T>
double Feature<T>::c_kulczynski1(Point<T>& a, Point<T>& b) {

	auto aid = a.get_id();
	auto bid = b.get_id();
	auto tup = std::tuple<uintmax_t, uintmax_t, uint8_t>(aid, bid, Feature<T>::log2(FEAT_KULCZYNSKI1));
	if (ltable.find(tup) == ltable.end()) {
		double val = kulczynski1(a, b);
		ltable.insert({tup, val});
		return val;
	} else {
		return ltable.at(tup);
	}
}

template<class T>
double Feature<T>::kulczynski1(Point<T> &a, Point<T> &b)
{
	const DivergencePoint<T>& p = dynamic_cast<const DivergencePoint<T>&>(a);
	const DivergencePoint<T>& q = dynamic_cast<const DivergencePoint<T>&>(b);
	const auto N = p.points.size();
	double sum = 0;
	for (auto i = 0; i < N; i++) {
		auto numer = p.points[i] > q.points[i] ? p.points[i] - q.points[i] : q.points[i] - p.points[i];
		auto denom = std::min(p.points[i], q.points[i]);
		sum += (double)numer / denom;
	}
	return sum;
}


// // Modified by Hani Z. Girgis on Oct 7 2018 to enable processing protine sequence.
// // Failed——needs understanding of the implementation.
// template<class T>
// double Feature<T>::d2_star(Point<T>& a, Point<T>& b)
// {
// 	const DivergencePoint<T>& p = dynamic_cast<const DivergencePoint<T>&>(a);
// 	const DivergencePoint<T>& q = dynamic_cast<const DivergencePoint<T>&>(b);
// 	const auto N = p.points.size();

// 	// const int k = (int)(log(N) / log(4));
// 	int k = a.getK();
// 	if(k==1){
// 		cerr << "D2s is skipped because it cannot be applied when k is 1.";
// 		cerr << endl;
// 		throw std::exception();
// 	}
// 	const int Alpha = Util::getAlphabetSize();

// 	const auto p1 = p.get_1mers();
// 	const auto q1 = q.get_1mers();

// 	const auto pmag = p.getPseudoMagnitude();
// 	const auto qmag = q.getPseudoMagnitude();
// 	double sum = 0;

// 	vector<double> tilde(Alpha, 0);
// 	for (int i = 0; i < Alpha; i++) {
// 		tilde[i] = (double)(p1[i] + q1[i]) / (pmag + qmag);
// 		cerr << "tilde[i]: " << tilde[i] << endl;
// 	}
// 	const double L = sqrt(pmag * qmag);
// 	for (auto i = 0; i < N; i++) {
// 		double p1i = 1;
// 	    double q1i = 1;
// 		double tilde_i = 1;
// 	    auto idx = i;
// 		for (int j = 0; j < k; j++) {
// 			auto i1 = idx % Alpha;
// 			idx /= Alpha;
// 			p1i *= (double)p1[i1] / pmag;
// 			q1i *= (double)q1[i1] / qmag;
// 			tilde_i *= tilde[i1];
// 		}
// 		double hp = p.points[i] - pmag * p1i;
// 		double hq = q.points[i] - qmag * q1i;
// 		sum += hp * hq / (L * tilde_i);
// 	}

// cerr << "L: " << L << endl;

// 	return sum;
// }


template<class T>
double Feature<T>::c_n2r(Point<T>& a, Point<T>& b) {

	auto aid = a.get_id();
	auto bid = b.get_id();
	auto tup = std::tuple<uintmax_t, uintmax_t, uint8_t>(aid, bid, Feature<T>::log2(FEAT_N2R));
	if (ltable.find(tup) == ltable.end()) {
		double val = n2r(a, b);
		ltable.insert({tup, val});
		return val;
	} else {
		return ltable.at(tup);
	}
}

template<class T>
double Feature<T>::n2r(Point<T>& a, Point<T>& b) const
{
	const DivergencePoint<T>& p = dynamic_cast<const DivergencePoint<T>&>(a);
	const DivergencePoint<T>& q = dynamic_cast<const DivergencePoint<T>&>(b);
	const auto N = p.points.size();
	double *cp = new double[N];
	double *cq = new double[N];
	double ap = 0, aq = 0;
	for (auto i = 0; i < N; i++) {
		int j = reverse.at(i);
		cp[i] = p.points[i] + p.points[j];
		cq[i] = q.points[i] + q.points[j];
		ap += cp[i];
		aq += cq[i];
	}
	ap /= N;
	aq /= N;
	double total = neighbor(cp, cq, ap, aq, N);
	delete[] cp;
	delete[] cq;
	return total;
}

template<class T>
double Feature<T>::c_n2rc(Point<T>& a, Point<T>& b) {

	auto aid = a.get_id();
	auto bid = b.get_id();
	auto tup = std::tuple<uintmax_t, uintmax_t, uint8_t>(aid, bid, Feature<T>::log2(FEAT_N2RC));
	if (ltable.find(tup) == ltable.end()) {
		double val = n2rc(a, b);
		ltable.insert({tup, val});
		return val;
	} else {
		return ltable.at(tup);
	}
}

template<class T>
double Feature<T>::n2rc(Point<T>& a, Point<T>& b) const
{
	if(!Util::isDna){
		cerr << "n2rc cannot be calculated on protein sequences." << endl;
		throw std::exception();
	}

	const DivergencePoint<T>& p = dynamic_cast<const DivergencePoint<T>&>(a);
	const DivergencePoint<T>& q = dynamic_cast<const DivergencePoint<T>&>(b);
	const auto N = p.points.size();
	double *cp = new double[N];
	double *cq = new double[N];
	double ap = 0, aq = 0;
	for (auto i = 0; i < N; i++) {
		int h = reverse_complement.at(i);
		cp[i] = p.points[h] + p.points[i];
		cq[i] = q.points[h] + q.points[i];
		ap += cp[i];
		aq += cq[i];
	}
	ap /= N;
	aq /= N;
	double total = neighbor(cp, cq, ap, aq, N);
	delete[] cp;
	delete[] cq;
	return total;
}

// template<class T>
// void Feature<T>::safe_insert(std::tuple<uintmax_t, uintmax_t, uint8_t> k, double v){
// 	# pragma omp critical
// 	{
// 		ltable.insert({k, v});
// 	}
// }

template class Feature<uint8_t>;
template class Feature<uint16_t>;
template class Feature<uint32_t>;
template class Feature<uint64_t>;
template class Feature<int>;
template class Feature<double>;
```
