## Supplementary Data 1 for "FASTCAR: Rapid alignment-free prediction of sequence alignment identity scores using self-supervised general linear models": Feature.h

```
/* -*- C++ -*-
 *
 * Feature.h
 *
 * Author: Benjamin T James
 *
 * Class containing all features and a glue to bind them together,
 * shared indivual features can be shared through hashing if sequence
 * id's are set.
 */
#ifndef FEATURE_H
#define FEATURE_H

// #include "SingleFeature.h"

#include <cmath>
#include <algorithm>
#include <map>
#include <functional>
#include <numeric>
#include <limits>
#include <vector>
#include <iostream>
#include <iterator>

#include "DivergencePoint.h"
#include "../../utility/GlobAlignE.h"

using namespace std;

#define	FEAT_ALIGN               (1UL << 0)
#define	FEAT_HELLINGER           (1UL << 1)
#define	FEAT_MANHATTAN           (1UL << 2)
#define	FEAT_EUCLIDEAN           (1UL << 3)
#define	FEAT_CHI_SQUARED         (1UL << 4)
#define	FEAT_NORMALIZED_VECTORS  (1UL << 5)
#define	FEAT_HARMONIC_MEAN       (1UL << 6)
#define	FEAT_JEFFEREY_DIV        (1UL << 7)
#define	FEAT_K_DIV               (1UL << 8)
#define	FEAT_PEARSON_COEFF       (1UL << 9)
#define	FEAT_SQCHORD             (1UL << 10)
#define	FEAT_KL_COND             (1UL << 11)
#define	FEAT_MARKOV              (1UL << 12)
#define	FEAT_INTERSECTION        (1UL << 13)
#define	FEAT_RRE_K_R             (1UL << 14)
#define	FEAT_D2z                 (1UL << 15)
#define	FEAT_SIM_MM              (1UL << 16)
#define	FEAT_EUCLIDEAN_Z         (1UL << 17)
#define	FEAT_EMD                 (1UL << 18)
#define	FEAT_SPEARMAN            (1UL << 19)
#define	FEAT_JACCARD             (1UL << 20)
#define	FEAT_LENGTHD             (1UL << 21)
#define	FEAT_D2s                 (1UL << 22)
#define	FEAT_AFD                 (1UL << 23)
#define	FEAT_MISMATCH            (1UL << 24)
#define	FEAT_CANBERRA            (1UL << 25)
#define	FEAT_KULCZYNSKI1         (1UL << 26)
#define	FEAT_KULCZYNSKI2         (1UL << 27)
#define	FEAT_SIMRATIO            (1UL << 28)
#define	FEAT_JENSEN_SHANNON      (1UL << 29)
#define	FEAT_D2_star             (1UL << 30)
#define	FEAT_N2R                 (1UL << 31)
#define	FEAT_N2RC                (1UL << 32)
#define	FEAT_N2RRC               (1UL << 33)

enum class Combo {
	xy,
	x2y2,
	xy2,
	x2y
};

template<class T>
struct pra {
	Point<T>* first;
	Point<T>* second;
	double val;
	pra() {}
	pra(const pra<T>&f) : first(f.first), second(f.second), val(f.val) {}
	pra(Point<T>* a, Point<T>* b, double c) : first(a), second(b), val(c) {}
	pra<T> deep_clone() const {
		return pra(first->clone(), second->clone(), val);
	}
};

/*
 * Usage:
 *   add_feature(FEAT_LD | FEAT_INTERSECTION, COMBO_SELF);
 *   add_feature(FEAT_LD | FEAT_JENSONSHANNON, COMBO_SELF);
 *
 *   normalize(some_pairs_to_normalize)
 *   normalize(more_pairs_to_normalize)
 *   finalize()
 *
 *   add_feature(....);
 *
 *   normalize(some_pairs_to_normalize)
 *   normalize(more_pairs_to_normalize)
 *   finalize()
 *
 *   compute(p,q)
 *   for (size_t i = 0; i < feature.size(); i++) {
 *       cout << feature[i] << endl;
 *   }
 */
template<class T>
class Feature {
public:
	Feature(const Feature<T>& feat_);
	Feature<T> operator=(const Feature<T>& feat_);
	Feature(const int k_) : k(k_) {
		flags = 0;

		// Modified by Hani Z. Girgis on Oct 9 2018 to enable processing protein
		auto freverse = [](int idx, int k) {
			int sum = 0;
			const auto A = Util::getAlphabetSize();
			for (int i = 0; i < k; i++) {
				int rem = idx % A;
				idx /= A;
				sum = A * sum + rem;
			}
			return sum;
		};

		auto freverse_complement = [](int idx, int k) {
			std::vector<int> v;
			for (int i = 0; i < k; i++) {
				v.push_back(3 - idx % 4);
				idx /= 4;
			}
			int sum = 0;
			for (auto val : v) {
				sum = 4 * sum + val;
			}
			return sum;
		};

		uint64_t k4_22 = 1;
		for (int i = 0; i < k; i++) {
			k4_22 *= Util::getAlphabetSize();
		}

		for (int i = 0; i < k4_22; i++) {
			reverse.push_back(freverse(i, k));
		}

		if(Util::isDna){
			for (int i = 0; i < k4_22; i++) {
				reverse_complement.push_back(freverse_complement(i, k));
			}
		}
	}
	void add_feature(uint64_t f_flags, Combo combo=Combo::xy);
	static vector<int> multi_to_log(uint64_t multi);
	vector<std::string> feat_names();
	static std::string feat_name(uint64_t single);
	void finalize();
	// std::vector<double> get_raw(const vector<pair<Point<T>*,Point<T>*> >&, int index) const;
	void remove_feature() { // Tear down features SPECIFIC to last pairing
		// auto indices_to_rm = combos.back().second;
		// combos.pop_back();
		// uint64_t feat_flags;


		// TO_DEL = TO_RM & (INDICES - REST)

		vector<int> vec = combos.back().second;
		combos.pop_back();
		for (auto combo : combos) {
			for (auto idx : combo.second) {
				vec.erase(std::remove(vec.begin(), vec.end(), idx), vec.end());
			}
		}
		std::sort(vec.begin(), vec.end(), std::greater<int>());
		for (int idx : vec) {
			flags ^= lookup[idx];
			lookup.erase(lookup.begin() + idx);
			raw_funcs.erase(raw_funcs.begin() + idx);
			mins.erase(mins.begin() + idx);
			maxs.erase(maxs.begin() + idx);
			is_sims.erase(is_sims.begin() + idx);
			is_finalized.erase(is_finalized.begin() + idx);
		}
		// flags ^= lookup[idx]
		// lookup[IDX]
		// raw_funcs[IDX]
		// mins[idx]
		// maxs[idx]
		// is_sims[idx]
		// is_finalized[idx]
	}
	void normalize(const vector<pra<T> > &pairs);
	void set_normal(uint64_t single_flag, double min, double max);
	vector<double> compute(Point<T>& p, Point<T>& q) {
		vector<double> cache = compute_all_raw(p, q);
		normalize_cache(cache);
		return cache;
	};

	// This should be called on the singles, which can be calculated
	// using the compute method
	double operator()(int col, const vector<double>& cache) const {
		auto pr = combos.at(col);
		Combo combo = pr.first;
		auto indices = pr.second;
		if (combo == Combo::xy) {
			double prod = 1;
			for (auto idx : indices) {
				prod *= cache[idx];
			}
			return prod;
		} else if (combo == Combo::x2y2) {
			double prod = 1;
			for (auto idx : indices) {
				prod *= cache[idx] * cache[idx];
			}
			return prod;
		} else if (combo == Combo::xy2) {
			if (indices.size() != 2) {
				cerr << "index size: " << indices.size() << endl;
				throw "invalid";
			}
			auto i0 = indices[0];
			auto i1 = indices[1];
			return cache[i0] * cache[i1] * cache[i1];
		} else if (combo == Combo::x2y) {
			if (indices.size() != 2) {
				throw "invalid";
			}
			auto i0 = indices[0];
			auto i1 = indices[1];
			return cache[i0] * cache[i0] * cache[i1];
		} else {
			throw "invalid combo";
		}
	}
	size_t size() const { return combos.size(); }
	void print_bounds() const {
		for (size_t i = 0; i < lookup.size(); i++) {
			cout << "bounds[" << i << "]: " << mins[i] << " to " << maxs[i] << endl;
		}
	}
	static int log2(uint64_t feature_) {
		for (size_t i = 0; i < 33; i++) {
			if (feature_ & (1UL << i)) {
				return i;
			}
		}
		return 0;
	}
	static double hellinger(Point<T>& p, Point<T>& q);
	double c_hellinger(Point<T>& p, Point<T>& q);
	static double manhattan(Point<T>& p, Point<T>& q);
	double c_manhattan(Point<T>& p, Point<T>& q);
	static double euclidean(Point<T>& p, Point<T>& q);
	double c_euclidean(Point<T>& p, Point<T>& q);
	static double chi_squared(Point<T>& p, Point<T>& q);
	double c_chi_squared(Point<T>& p, Point<T>& q);
	static double normalized_vectors(Point<T>& p, Point<T>& q);
	double c_normalized_vectors(Point<T>& p, Point<T>& q);
	static double harmonic_mean(Point<T>& p, Point<T>& q);
	double c_harmonic_mean(Point<T>& p, Point<T>& q);
	static double jefferey_divergence(Point<T>& p, Point<T>& q);
	double c_jefferey_divergence(Point<T>& p, Point<T>& q);
	static double k_divergence(Point<T>& p, Point<T>& q);
	double c_k_divergence(Point<T>& p, Point<T>& q);
	static double pearson(Point<T>& p, Point<T>& q);
	double c_pearson(Point<T>& p, Point<T>& q);
	static double squaredchord(Point<T>& a, Point<T>& b);
	double c_squaredchord(Point<T>& a, Point<T>& b);
	static double kl_conditional(Point<T>& a, Point<T>& b);
	double c_kl_conditional(Point<T>& a, Point<T>& b);
	static double markov(Point<T>& a, Point<T>& b);
	double c_markov(Point<T>& a, Point<T>& b);
	static double intersection(Point<T>& p, Point<T>& q);
	double c_intersection(Point<T>& p, Point<T>& q);
	static double rre_k_r(Point<T>& p, Point<T>& q);
	double c_rre_k_r(Point<T>& p, Point<T>& q);
	static double d2z(Point<T>& p, Point<T>& q);
	double c_d2z(Point<T>& p, Point<T>& q);
	static double sim_mm(Point<T>& p, Point<T>& q);
	double c_sim_mm(Point<T>& p, Point<T>& q);
	static double euclidean_z(Point<T>& p, Point<T>& q);
	double c_euclidean_z(Point<T>& p, Point<T>& q);
	static double emd(Point<T>& p, Point<T>& q);
	double c_emd(Point<T>& p, Point<T>& q);
	static double spearman(Point<T>& p, Point<T>& q);
	double c_spearman(Point<T>& p, Point<T>& q);
	static double jaccard(Point<T>& p, Point<T>& q);
	double c_jaccard(Point<T>& p, Point<T>& q);
	static double length_difference(Point<T>& p, Point<T>& q);
	static double d2s(Point<T>& p, Point<T>& q);
	double c_d2s(Point<T>& p, Point<T>& q);
	static double afd(Point<T>& p, Point<T>& q);
	double c_afd(Point<T>& p, Point<T>& q);
	static double mismatch(Point<T>& p, Point<T>& q);
	double c_mismatch(Point<T>& p, Point<T>& q);
	static double canberra(Point<T>& p, Point<T>& q);
	double c_canberra(Point<T>& p, Point<T>& q);
	static double kulczynski1(Point<T>& a, Point<T>& b);
	double c_kulczynski1(Point<T>& a, Point<T>& b);
	static double kulczynski2(Point<T>& a, Point<T>& b);
	double c_kulczynski2(Point<T>& a, Point<T>& b);
	static double simratio(Point<T>& a, Point<T>& b);
	double c_simratio(Point<T>& a, Point<T>& b);
	double jensen_shannon(Point<T>& p, Point<T>& q) const;
	double c_jensen_shannon(Point<T>& p, Point<T>& q);
	static double d2_star(Point<T>& p, Point<T>& q);
	double c_d2_star(Point<T>& p, Point<T>& q);
	double n2r(Point<T>& p, Point<T>& q) const;
	double c_n2r(Point<T>& p, Point<T>& q);
	double n2rc(Point<T>& p, Point<T>& q) const;
	double c_n2rc(Point<T>& p, Point<T>& q);
	double n2rrc(Point<T>& p, Point<T>& q) const;
	double c_n2rrc(Point<T>& p, Point<T>& q);

	static double align(Point<T>& a, Point<T>& b, std::map<std::pair<uintmax_t, uintmax_t>, double> &atable);
	std::function<double(Point<T>&,Point<T>&)> get_func(uint64_t single_feat);
	std::function<double(Point<T>&,Point<T>&)> get_func_(uint64_t single_feat);
	bool feat_is_sim(uint64_t single_flag) const;
	bool get_save() const { return do_save; }
	void set_save(bool save_) {
		do_save = save_;
		if (!save_) {
			ltable.clear();
		}
	}
	std::vector<std::pair<Combo, std::vector<int> > > get_combos() const { return combos; }
	std::vector<double> get_mins() const { return mins; };
	std::vector<double> get_maxs() const { return maxs; };
	std::vector<uint64_t> get_lookup() const { return lookup; };
private:

	vector<double> compute_all_raw(Point<T>& p, Point<T>& q);
	void normalize_cache(vector<double>& cache) const;


	// double raw(uint64_t single_flag, Point<T>& a, Point<T>& b);
	int index_of(uint64_t single_flag) const {
		for (size_t i = 0; i < lookup.size(); i++) {
			if (lookup[i] == single_flag) {
				return i;
			}
		}
		return -1;
	}
	void reset_funcs() {
		raw_funcs.clear();
		for (auto f : lookup) {
			raw_funcs.push_back(get_func(f));
		}
	}
	uint64_t get_flags() const { return flags; };


	std::vector<bool> get_sims() const { return is_sims; };
	std::vector<bool> get_finalized() const { return is_finalized; };

	int k;
	int get_k() const { return k; };
	uint64_t flags;
	bool do_save;
	std::vector<std::pair<Combo,
			      std::vector<int>
			      > > combos;

	std::vector<double> mins, maxs;
	std::vector<bool> is_sims, is_finalized;
	std::vector<uint64_t> lookup;
	std::vector<int> reverse, reverse_complement;
	std::vector<std::function<double(Point<T>&,Point<T>&)> > raw_funcs;

	std::map<std::pair<uintmax_t,uintmax_t>, double> atable;
	std::map<std::tuple<uintmax_t, uintmax_t, uint8_t>, double> ltable;

	// Added by Hani Z. Girgis
	// std::vector<double> tiedrank(const Point<T>& a);

//	std::map<std::tuple<uintmax_t, uintmax_t, uint8_t>, double> * get_table() const { return ltable; }
};

// template<class T>
// class Feature {
// public:
// 	Feature(std::function<double(vector<double>)> combination, std::vector<SingleFeature<T> > sf)
// 		: features(sf), combo(combination) {}
// 	double operator()(Point<T>*, Point<T>*) const;


// 	static double manhattan(Point<T>& p, Point<T>& q);
// 	static double length_difference(Point<T>& p, Point<T>& q);
// 	static double n2rrc(Point<T>& p, Point<T>& q, const vector<int>&, const vector<int> &);
// 	static double rre_k_r(Point<T>& p, Point<T>& q);
// 	static double intersection(Point<T>& p, Point<T>& q);
// 	static double jenson_shannon(Point<T>& p, Point<T>& q);
// 	static double pearson(Point<T>& p, Point<T>& q);
// 	static double simratio(Point<T>& a, Point<T>& b);
// 	static double squaredchord(Point<T>& a, Point<T>& b);
// private:
// 	vector<SingleFeature<T> > features;
// 	std::function<double(vector<double>)> combo;
// };

//#include "Feature.cpp"

#endif
```
