## Supplementary Data 1 for "FASTCAR: Rapid alignment-free prediction of sequence alignment identity scores using self-supervised general linear models": GLM.cpp

```
/*
 * glm.cpp
 *
 * Created on: May 29, 2017
 * Author: Robert Geraghty, The Bioinformatics Toolsmith Laboratory, The University of Tulsa
 *
 * Modified by Benjamin T James
 */

#include "GLM.h"
#include "Matrix.h"

#include <math.h>
#include <iostream>
using namespace std;
// using namespace matrix;

namespace matrix{

void GLM::train(Matrix& features, Matrix& labels){
	weights = features.transpose() * features;
	weights = weights.pseudoInverse() * features.transpose() * labels;
}


double GLM::logistic(double x)
{
	return 1.0 / (1 + exp(-x));
}
Matrix GLM::predict(Matrix& features) const {
	Matrix labels;
	labels	= features * weights;
	double log;
	for(int i = 0; i < labels.getNumRow(); i++){
		//log = round(1/(1 + exp(-(labels.get(i,0)))) + 0.1);
		labels.set(i,0, round(logistic(labels.get(i, 0))));
	}
	return labels;
}

std::tuple<double,double,double> GLM::accuracy(Matrix& oLabels, Matrix& pLabels) const {
	int sum = 0;
	int negSum = 0;
	int negSame = 0;
	int posSum = 0;
	int posSame = 0;
	for(int i = 0; i < oLabels.getNumRow(); i++){
		if(oLabels.get(i,0) == -1){
			negSum++;
			if(oLabels.get(i,0) == pLabels.get(i, 0)){
				sum++;
				negSame++;
			}
		}else{
			posSum++;
			if(oLabels.get(i,0) == pLabels.get(i, 0)){
				sum++;
				posSame++;
			}
		}
	}
	double acc = (((double)sum*100)/(oLabels.getNumRow()));
	double sens =  (((double)posSame*100)/(posSum));
	double spec = (((double)negSame*100)/(negSum));
	// cout << "Accuracy: " << acc << "% ";
	// cout << "Sensitivity: " << sens << "% ";
	// cout << "Specificity: " << spec << "% " << endl;
	return make_tuple(acc, sens, spec);
}

}
```
