## Supplementary Data 1 for "FASTCAR: Rapid alignment-free prediction of sequence alignment identity scores using self-supervised general linear models": GLM.h

```
/*
 * glm.h
 *
 * Created on: May 29, 2017
 * Author: Robert Geraghty, The Bioinformatics Toolsmith Laboratory, The University of Tulsa
 *
 * Modified by Benjamin T James
 */

#ifndef SRC_MATRIX_GLM_H_
#define SRC_MATRIX_GLM_H_

#include "Matrix.h"
#include <tuple>
namespace matrix {

class GLM {
private:
	Matrix weights;

public:
	void load(Matrix weights_) { weights = weights_; }
	void train(matrix::Matrix& features, matrix::Matrix& labels);
	Matrix predict(matrix::Matrix& features) const;
	static double logistic(double x);
	static double linear(double x);
	std::tuple<double,double,double> accuracy(matrix::Matrix& oLabels, matrix::Matrix& pLabels) const;
	const Matrix& get_weights() const { return weights; };
};

}

#endif /* SRC_MATRIX_GLM_H_ */
```
