## Supplementary Data 1 for "FASTCAR: Rapid alignment-free prediction of sequence alignment identity scores using self-supervised general linear models": Histogram.cpp

```
/* -*- C++ -*-
 *
 * Histogram.cpp
 *
 * Author: Benjamin T James
 *
 * Artifact from early development of MeShClust
 */
#ifndef HEADER_HACK
#include "Histogram.h"
#endif

#include <vector>
#include <iostream>

template<class T>
double Histogram<T>::distance_k1(const Point<T> &p) const
{
	throw "Not implemented";
	const Histogram<T>& h = dynamic_cast<const Histogram<T>&>(p);
	uint64_t dist = 0;
        auto size = std::min(points.size(),h.points.size());
/*
	for (unsigned int i = 0; i < size; i++) {
		T l = points.at(i);
		T r = h.points.at(i);
		dist += (l > r) ? (l - r) : (r - l);
	}
*/
	uint64_t avg_mag = (magnitude() + h.magnitude()) / 2.0;
	for (auto i = 0; i < size; i++) {
		T l = points[i];
		T r = h.points[i];
		dist += min(l, r);
	}
	return 1.0 - dist / avg_mag;
}
template<class T>
Histogram<T>::Histogram(std::vector<T> pts, char mark)
{
	for (T t : pts) {
		points.push_back(t);
	}
	to_delete = false;
}
template<class T>
Histogram<T>::Histogram(std::vector<T> pts)
{
	for (T t : pts) {
		points.push_back(t);
	}
	to_delete = false;
}

template<class T>
Histogram<T>::Histogram(std::vector<T> pts, bool toDelete)
{
	for (T t : pts) {
		points.push_back(t);
	}
	to_delete = toDelete;
}

template<class T>
Histogram<T>::Histogram(unsigned int size)
{
	for (unsigned int i = 0; i < size; i++) {
		points.push_back(0);
	}
	to_delete = false;
}

template<class T>
void Histogram<T>::operator*=(double d)
{
	for (T &t : points) {
		t *= d;
	}
}

template<class T>
bool Histogram<T>::operator<(Point<T>& p) const
{
	const Histogram<T>& h = dynamic_cast<const Histogram<T>&>(p);
	unsigned int size = std::min(points.size(),h.points.size());
	for (unsigned int i = 0; i < size; i++) {
		if (points.at(i) >= h.points.at(i)) {
			return false;
		}
	}
	return true;
}

template<class T>
void Histogram<T>::operator/=(double d)
{
	unsigned int size = points.size();
	for (unsigned int i = 0; i < size; i++) {
		points.at(i) = points.at(i) / d;
	}
}

template<class T>
void Histogram<T>::operator+=(Point<T>& p)
{
	const Histogram<T>& h = dynamic_cast<const Histogram<T>&>(p);
	unsigned int size = std::min(points.size(),h.points.size());
	for (unsigned int i = 0; i < size; i++) {
		points.at(i) += h.points.at(i);
	}
}

template<class T>
uint64_t Histogram<T>::operator-(const Point<T>& p) const
{
	return distance(p);
}

template<class T>
void Histogram<T>::set(Point<T>& p)
{
	const Histogram<T>& h = dynamic_cast<const Histogram<T>&>(p);
	points = h.points;
}

template<class T>
void Histogram<T>::display() const
{
	unsigned size = points.size();
	for (unsigned i = 0; i < size; i++) {
		std::cout << points.at(i) << " ";
	}
	std::cout << std::endl;
}

template<class T>
void Histogram<T>::addOne()
{
	for (auto &a : points) {
		a++;
	}
}
template<class T>
void Histogram<T>::subOne()
{
	for (auto &a : points) {
		a--;
	}
}

template<class T>
void Histogram<T>::zero()
{
	for (typename std::vector<T>::iterator it = points.begin(); it != points.end(); ++it) {
		*it = 0;
	}
}

template<class T>
uint64_t Histogram<T>::distance(const Point<T>& p) const
{
/*
	// Vectors should be the same width
	const Histogram<T>& h = dynamic_cast<const Histogram<T>&>(p);
	T dist = 0;
	unsigned int size = std::min(points.size(),h.points.size());
	for (unsigned int i = 0; i < size; i++) {
		T l = points.at(i);
		T r = h.points.at(i);
		dist += (l > r) ? (l - r) : (r - l);
	}
	return dist;
*/
	throw "Not implemented";
	return 0;
}

template<class T>
uint64_t Histogram<T>::magnitude() const
{
	uint64_t dist = 0;
	for (auto const& p : points) {
		dist += p;
	}
	return dist;
}

#ifndef HEADER_HACK
template class Histogram<int>;
template class Histogram<double>;
template class Histogram<uint64_t>;
template class Histogram<uint32_t>;
template class Histogram<uint16_t>;
template class Histogram<uint8_t>;
#endif
```
