## Supplementary Data 1 for "FASTCAR: Rapid alignment-free prediction of sequence alignment identity scores using self-supervised general linear models": Histogram.h

```
/* -*- C++ -*-
 *
 * Histogram.h
 *
 * Author: Benjamin T James
 *
 * Artifact from early development of MeShClust
 */
#ifndef HISTOGRAM_H
#define HISTOGRAM_H
#include <vector>
#include "Point.h"

template<class T>
class Histogram : public Point<T> {
public:
	Histogram(std::vector<T> pts);
	Histogram(std::vector<T> pts, char marker);
	Histogram(std::vector<T> pts, bool to_delete);
	Histogram(unsigned int size);
	~Histogram() {}
	void operator*=(double d);
	void operator/=(double d);
	uint64_t operator-(const Point<T>& p) const;
	bool operator<(Point<T>& p) const;
	void operator+=(Point<T>& p);
	void set(Point<T>& p);
	void display() const;
	void zero();
	void addOne();
	void subOne();
	double distance_k1(const Point<T>& p) const;
	double prob_under(Point<T>& p) const { return distance(p); };
	uint64_t distance(const Point<T>& p) const;
	uint64_t magnitude() const;
	uint64_t getRealMagnitude() const { return 0; };
	double distance_d(Point<double>& p) const {
		throw "not implemented";
		return 0;
	}
	void set_arg_to_this_d(Point<double>& p) const {
		throw "not implemented";
	}
	Point<double>* create_double() const {
		throw "not implemented";
		return NULL;
	}
	Histogram* clone() const {
		return new Histogram(points, to_delete);
	}
	Histogram* create() const {
		return new Histogram(points.size());
	}
	bool is_to_delete() const {
		return to_delete;
	}
	void set_to_delete(bool b) {
		to_delete = b;
	}
	const vector<T>& get_data() const { return points; }
	void set_id(uintmax_t c_id) { id = c_id; };
	const uintmax_t get_id() const { return id; };
	void set_length(unsigned long len) { nucl_length = len; };
	unsigned long get_length() const { return nucl_length; };
        unsigned long size() const { return points.size(); };
private:
	std::vector<T> points;
	bool to_delete;
	uintmax_t id;
	unsigned long nucl_length;
};

#ifdef HEADER_HACK
#ifndef HISTOGRAM_C
#define HISTORGRAM_C
#include "Histogram.cpp"
#endif
#endif

#endif
```
