## Supplementary Data 1 for "FASTCAR: Rapid alignment-free prediction of sequence alignment identity scores using self-supervised general linear models": Loader.cpp

```
/* -*- C++ -*-
 *
 * Loader.cpp
 *
 * Author: Benjamin T James
 *
 * Class which can 'preload' chunks of sequences from a file list,
 * and then count the k-mers separately, which can be done in
 * multiple threads
 */
#include "Loader.h"
#include "Runner.h"

static uint64_t num_overflow = 0;
std::string next_histogram(std::string cur_type)
{
	if (cur_type == "uint8_t") {
		return "uint16_t";
	} else if (cur_type == "uint16_t") {
		return "uint32_t";
	} else {
		return "uint64_t";
	}
}

template<class T>
std::string Loader<T>::get_warning()
{
	if (num_overflow == 0) {
		return "";
	} else {
		std::ostringstream oss;
		oss << "For " << num_overflow << " sequences, the histogram type " << Runner::get_datatype() << " was too small for holding sequences." << endl;
		oss << "Performance may be slightly hindered, but can be improved by increasing the integral type (--datatype " << next_histogram(Runner::get_datatype())  << ")" << endl;
		return oss.str();
	}
}


template<class V>
void Loader<V>::fill_table(KmerHashTable<unsigned long, V> &table, ChromosomeOneDigit *chrom, std::vector<V>& values)
{
	const int k = table.getK();
	auto segment = chrom->getSegment();
	const char *seg_bases = chrom->getBase()->c_str();
	for (vector<int> *v : *segment) {
		int start = v->at(0);
		int end = v->at(1);

		// Hani Z Girgis added the following line
		// It is possible
		if(end - start + 1 >= k){
			int r = table.wholesaleIncrementNoOverflow(seg_bases, start, end - k + 1);
			if (r == -1) {
				num_overflow++;
				// #pragma omp critical
				// {
				// 	std::ostringstream oss;
				// 	oss << "In header \"" << chrom->getHeader() << "\"" << endl;
				// 	oss << "Histogram type " << Runner::get_datatype() << " is too small for holding sequences." << endl;
				// 	oss << "Performance may be slightly hindered, but can be improved by increasing the integral type (--datatype " << next_histogram(Runner::get_datatype())  << ")" << endl;
				// 	_loader_warning = oss.str();
				// 	cerr << get_warning() << endl;
				// }
			}
		}
	}

	std::string header = chrom->getHeader();
	header = header.substr(1, header.find(' ')-1);
	// Hani Z. Girgis added the following lines on 10/3/2018
	// This should result in significant speed up.
	unsigned long tableSize = table.getMaxTableSize();
	values.reserve(values.size() + tableSize);
	const V * valueArray = table.getValues();

	copy(&valueArray[0], &valueArray[tableSize], back_inserter(values));

    // Commented out by Hani Z. Girgis on 10/3/2018 and replaced by the code above
	// std::vector<std::string> *keys = table.getKeys();
	// for (std::string str : *keys) {
	// 	values.push_back(table.valueOf(str.c_str()));
	// }
	// keys->clear();
	// delete keys;
}

template<class T>
bool Loader<T>::done() const
{
	return file_idx == files.size();
}

template<class T>
void Loader<T>::preload(int tid)
{
	if (file_idx == files.size()) {
		return;
	}
	for (uint64_t j = 0; j < chunk_size; j++) {
		auto chrom = next();
		if (chrom.first == "") {
			return;
		}
		cache_list.at(tid).emplace_back(chrom.first, chrom.second);
	}
}


// Modified by Hani Z. Girgis on Oct 2, 2018
template<class T>
Point<T>* Loader<T>::get_point(std::string header, const std::string &base, uintmax_t& id, int k)
{
	std::string obase = "";
	for (int i = 0; i < base.length(); i++) {
		if (base[i] == 'A' || base[i] == 'C' ||
		    base[i] == 'G' || base[i] == 'T') {
			obase += base[i];
		}
	}
	ChromosomeOneDigit * chrom;
	if(Util::isDna){
		chrom = new ChromosomeOneDigitDna();
	}else{
		chrom = new ChromosomeOneDigitProtein();
	}

	chrom->setHeader(header);
	chrom->appendToSequence(obase);
	chrom->finalize();
	Point<T> *p = Loader<T>::get_point(chrom, id, k);
	delete chrom;
	return p;
}

// Modified by Hani Z. Girgis on Oct 2, 2018
template<class T>
Point<T>* Loader<T>::get_point(ChromosomeOneDigit* chrom, uintmax_t& id, int k)
{

	KmerHashTable<unsigned long, T> table(k, 1);
	// Hani Z. Girgis changed the following line
	// The table_k1 was initialized from 0 now it is 1
	KmerHashTable<unsigned long, uint64_t> table_k1(1, 1);
	std::vector<T> values;
	vector<uint64_t> values_k1;
	// values.clear();

	Loader<T>::fill_table(table, chrom, values);
	Loader<uint64_t>::fill_table(table_k1, chrom, values_k1);
//	int tmplate = get_template(chrom->getHeader(), templates);
	Point<T> *p = new DivergencePoint<T>(values, chrom->size());
//	cout << "mag: " << ((DivergencePoint<T>*)p)->getPseudoMagnitude() << std::endl;
	p->set_1mers(values_k1);
	p->set_header(chrom->getHeader());
	p->set_length(chrom->getBase()->length());
	p->set_data_str(*chrom->getBase());
	// Added by Hani Z. Girgis on Oct 7 2018
	p->setK(k);
	DivergencePoint<T>* q = dynamic_cast<DivergencePoint<T>*>(p);
	const auto N = q->points.size();
	double aq = (double) q->getPseudoMagnitude() / N;
	double sq = 0;
	for (auto i = 0; i < N; i++) {
		double qdiff = q->points[i] - aq;
		sq += qdiff * qdiff;
	}
	sq = sqrt(sq / N);
	q->set_stddev(sq);
	p->set_id(id);
	#pragma omp atomic
	id++;

	// Clean

	return p;
}


template<class T>
std::vector<Point<T>*> Loader<T>::load_next(int tid)
{
	std::vector<Point<T>*> points;
	for (size_t i = 0; i < cache_list.at(tid).size(); i++) {
	        auto pr = cache_list.at(tid).at(i);
		Point<T>* p = get_point(pr.first, *pr.second, id_list.at(tid), k);
		points.push_back(p);
		delete pr.second;
	}
	cache_list.at(tid).clear();
	return points;
}

template<class T>
std::pair<std::string,std::string*> Loader<T>::next()
{
	auto n = maker->next();
	if (n.first != "") {
		return n;
	}
	delete maker;
	maker = NULL;
	file_idx++;
	if (file_idx >= files.size()) {
		return n;
	}
	maker = new SingleFileLoader(files.at(file_idx));
	return maker->next();
}

template class Loader<double>;
template class Loader<int>;
template class Loader<uint64_t>;
template class Loader<uint32_t>;
template class Loader<uint16_t>;
template class Loader<uint8_t>;
```
