## Supplementary Data 1 for "FASTCAR: Rapid alignment-free prediction of sequence alignment identity scores using self-supervised general linear models": Loader.h

```
/* -*- C++ -*-
 *
 * Loader.h
 *
 * Author: Benjamin T James
 *
 * Class which can 'preload' chunks of sequences from a file list,
 * and then count the k-mers separately, which can be done in
 * multiple threads
 */
#ifndef LOADER_H
#define LOADER_H

#include <omp.h>

#include "SingleFileLoader.h"
#include "Point.h"
#include "DivergencePoint.h"
#include "../../nonltr/KmerHashTable.h"
// Add by Hani Z. Girgis, PhD on Oct 2, 2018
#include "../../nonltr/ChromosomeOneDigit.h"
#include "../../nonltr/ChromosomeOneDigitDna.h"
#include "../../nonltr/ChromosomeOneDigitProtein.h"


template<class T>
class Loader {
public:
	Loader(std::vector<std::string> files_,
	       uint64_t total_num_points_,
	       uint64_t chunk_size_,
	       int num_threads_,
	       int k_,
	       uint64_t start_id=0)
		:
		chunk_size(chunk_size_),
		num_threads(num_threads_),
		k(k_),
		files(files_) {

		maker = new SingleFileLoader(files.at(0));
		uint64_t total_id = start_id;
		for (int i = 0; i < num_threads_; i++) {
			id_list.push_back(total_id);
			total_id += total_num_points_;
			cache_list.push_back(std::vector<std::pair<std::string,std::string*> >());
		}
//		preload();
	};

	~Loader() {
		if (get_warning() != "") {
			cerr << get_warning() << endl;
		}
		cache_list.clear();
		id_list.clear();
		if (maker != NULL) {
			delete maker;
		}
	}

	// single threaded
	void preload(int tnum);

	bool done() const;
	// multi-thread accessible
	std::vector<Point<T>*> load_next(int tid);

	static Point<T>* get_point(std::string header, const std::string &base, uintmax_t& id, int k);
	static Point<T>* get_point(ChromosomeOneDigit* dna, uintmax_t& id, int k);

	static void fill_table(KmerHashTable<unsigned long, T> &table, ChromosomeOneDigit *chrom, std::vector<T>& values);
	static std::string get_warning();
private:

	std::pair<std::string,std::string*> next();

	uint64_t chunk_size;
	int num_threads, k;

	std::vector<std::vector<std::pair<std::string,std::string*> > > cache_list;
	std::vector<uint64_t> id_list;

	std::vector<std::string> files;
	size_t file_idx = 0;
	SingleFileLoader *maker = NULL;

};

#endif
```
