## Supplementary Data 1 for "FASTCAR: Rapid alignment-free prediction of sequence alignment identity scores using self-supervised general linear models": Matrix.cpp

```
/*
 * matrix.cpp
 *
 * Created on: May 10, 2017
 * Author: Robert Geraghty, The Bioinformatics Toolsmith Laboratory, The University of Tulsa
 */

#include "Matrix.h"

#include <vector>
#include <iostream>
#include <fstream>
#include <sstream>
#include <string>
#include <iomanip>
#include <stdlib.h>
#include <math.h>

using namespace std;

namespace matrix {

Matrix::Matrix(vector<double> vec) : numRow(1), numCol(vec.size()) {
       	m.at(0) = vector<double>(vec.size());
       	for (int i = 0; i < vec.size(); i++) {
       		set(0, i, vec[i]);
       	}
}

Matrix::Matrix(int r, int c) :
		numRow(r), numCol(c) {
	m.resize(r);
	for (int i = 0; i < r; i++) {
		m.at(i) = vector<double>(c);
	}
}
Matrix::Matrix() :
		numRow(0), numCol(0) {

}

Matrix::~Matrix() {

}

Matrix Matrix::operator+(Matrix n) {
	if (numCol == n.numCol && numRow == n.numRow) {
		Matrix mat = Matrix(numRow, numCol);
		for (int i = 0; i < mat.numRow; i++) {
			for (int j = 0; j < mat.numCol; j++) {
				mat.set(i, j, (get(i, j) + n.get(i, j)));
			}
		}
		return mat;
	} else {
		cerr << "Invalid input: array dimension mismatch." << endl;
		throw exception();
	}
}

Matrix Matrix::operator-(Matrix n) {
	if (numCol == n.numCol && numRow == n.numRow) {
		Matrix mat = Matrix(numRow, numCol);
		for (int i = 0; i < mat.numRow; i++) {
			for (int j = 0; j < mat.numCol; j++) {
				mat.set(i, j, (get(i, j) - n.get(i, j)));
			}
		}
		return mat;
	} else {
		cerr << "Invalid input: array dimension mismatch." << "\n";
		throw exception();
	}
}

Matrix Matrix::operator*(Matrix n) {

	if (numCol == n.numRow) {
		double curSum = 0;
		Matrix mat = Matrix(numRow, n.numCol);
////#pragma omp parallel for collapse(2)
		for (int i = 0; i < mat.numRow; i++) {
			for (int j = 0; j < mat.numCol; j++) {
				curSum = 0;
				for (int k = 0; k < numCol; k++) {
					curSum = curSum + get(i, k) * n.get(k, j);
				}
				mat.set(i, j, curSum);
			}
		}
		return mat;
	} else {
		cerr << "Invalid input: array dimension mismatch." << endl;
		throw exception();
	}
}

Matrix Matrix::transpose() {
	Matrix temp = Matrix(numCol, numRow);
	for (int i = 0; i < numRow; i++) {
		for (int j = 0; j < numCol; j++) {
			temp.set(j, i, get(i, j));
		}
	}
	return temp;

}

Matrix Matrix::gaussJordanInverse() {
	if (numRow == numCol) {			//Checks if matrix is square
		Matrix invert = Matrix(numRow, numCol);
		Matrix temp = Matrix(numRow, numCol);
		double pivotVal;

		temp.m = m;

		for (int i = 0; i < numRow; i++) {//Creates identity Matrix, which will become inverse matrix
			invert.set(i, i, 1);
		}

		for (int i = 0; i < numRow; i++) {
			if (get(i, i) != 1) {				//Checks if the pivot point is 1
				if (get(i, i) != 0) {//Check if the pivot point is 0, if not it performs a type 2 row operation to set the pivot point to 1
					pivotVal = get(i, i);
					for (int j = 0; j < numCol; j++) {
						set(i, j, (get(i, j) / pivotVal));
						invert.set(i, j, (invert.get(i, j) / pivotVal));
					}
				} else {//If the pivot point is zero, it performs a type 1 row operation
					bool properSwap = false;
					int row = i + 1;
					double valSwap;
					double valSwap2;
					while (!properSwap && row < numRow) {
						if (get(row, i) != 0) {
							properSwap = true;
						} else {
							row++;
						}
					}
					if (properSwap) {
						for (int j = 0; j < numCol; j++) {
							valSwap = get(i, j);
							valSwap2 = invert.get(i, j);
							set(i, j, get(row, j));
							invert.set(i, j, (invert.get(row, j)));
							set(row, j, valSwap);
							invert.set(row, j, valSwap2);
						}
					} else {//If it cannot perform a type 1 row swap with a non zero pivot value, the Inverse does not exist.
						cout << "Inverse does not exist\n";
						m = temp.m;
						return temp;
					}
					pivotVal = get(i, i);
					for (int j = 0; j < numCol; j++) {//Now perform a type 2 row operation to set the new pivot point to 1
						set(i, j, (get(i, j) / pivotVal));
						invert.set(i, j, (invert.get(i, j) / pivotVal));
					}
				}
			}
			for (int below = i + 1; below < numRow; below++) { //Iterate through the elements below the pivot, performing type 3 row operations to set each to 0
				if (get(below, i) != 0) {
					pivotVal = get(below, i);
					for (int j = 0; j < numCol; j++) {
						set(below, j, (get(below, j) - (pivotVal * get(i, j))));
						invert.set(below, j,
								(invert.get(below, j)
										- (pivotVal * invert.get(i, j))));
					}
				}
			}
		}
		//		cout << "\n\n";
		for (int i = numRow - 1; i >= 0; i--) {	//Now perform the same step as the last except on the elements above the pivot.
			for (int above = 0; above < i; above++) {
				if (get(above, i) != 0) {
					pivotVal = get(above, i);
					for (int j = 0; j < numCol; j++) {
						set(above, j, (get(above, j) - (pivotVal * get(i, j))));
						invert.set(above, j,
								(invert.get(above, j)
										- (pivotVal * invert.get(i, j))));
					}
				}
			}
		}
		for (int i = 0; i < numRow; i++) {//Now check to make sure the original matrix is an identity matrix.
			for (int j = 0; j < numCol; j++) {
				if (i == j && get(i, j) != 1) {
					cout << "Inverse does not exist\n";
					m = temp.m;
					return temp;
				}
				if (i != j && get(i, j) != 0) {
					cout << "Inverse does not exist\n";
					m = temp.m;
					return temp;
				}
			}
		}
		m = temp.m;				//Reset the original matrix
		return invert;
	}
	cerr << "Invalid dimensions" << endl;
	throw exception();
}

Matrix Matrix::pseudoInverse() {
	if (numRow >= numCol) {
		Matrix temp = transpose();
		Matrix transByOrig = temp * *this;
		Matrix psuedoInv = (transByOrig.gaussJordanInverse()) * temp;
		return psuedoInv;
	} else {
		Matrix temp = transpose();
		Matrix origByTrans = *this * temp;
		Matrix psuedoInv = temp * (origByTrans.gaussJordanInverse());
		return psuedoInv;
	}
}

double Matrix::get(int r, int c) const {
	return m.at(r).at(c);
}

void Matrix::set(int r, int c, double val) {
	m.at(r).at(c) = val;
	//m[r][c] = val;
}

void Matrix::print() {
	for (int i = 0; i < numRow; i++) {
		for (int j = 0; j < numCol; j++) {
			cout << right << fixed;
			cout << "[" << setprecision(4) << setw(7) << get(i, j) << "] ";
		}
		cout << endl;
	}
	cout << endl;
}

void Matrix::printToFile(string fileName) {
	ofstream outSequence(fileName.c_str());

	for (int i = 0; i < numRow; i++) {
		for (int j = 0; j < numCol; j++) {
			outSequence << right << fixed;
			outSequence << "[" << setprecision(4) << setw(7) << get(i, j)
					<< "] ";
		}
		outSequence << endl;
	}
	outSequence << endl;

	outSequence.close();
}

void Matrix::randFill(double low, double high) {
	double x;
	for (int i = 0; i < numRow; i++) {
		for (int j = 0; j < numCol; j++) {
			x = ((double) rand() * (high - low)) / (double) RAND_MAX + low;
			set(i, j, x);
		}
	}
}

void Matrix::userFill() {
	double val;
	for (int i = 0; i < numRow; i++) {
		for (int j = 0; j < numCol; j++) {
			cout << "input value for cell (" << i << ", " << j << ")?\n";
			cin >> val;
			cout << endl;
			set(i, j, val);
		}
	}
}

void Matrix::fileFill(string filename) {
	ifstream infile(filename.c_str());
	if (!infile) {
		cerr << "file read fail" << endl;
		throw exception();
	}
	string line;
	int i = -1;
	while (getline(infile, line)) {
		i++;
		if (i >= numRow) {
			addRow(0);
		}
		double num;
		istringstream iss(line);
		int j = -1;
		while (iss >> num) {
			j++;
			if (j >= numCol) {
				addCol(0);
			}
			//cout << num << endl;
			set(i, j, num);
		}
		j = 0;
	}
	i = 0;
}

void Matrix::addRow(double val) {
	numRow++;
	vector<double> temp = vector<double>(numCol, val);
	m.push_back(temp);
}

void Matrix::addCol(double val) {
	numCol++;
	for (int i = 0; i < numRow; i++) {
		m.at(i).push_back(val);
	}
}

void Matrix::normalize(double a, double b) {
	for (int j = 0; j < numCol; j++) {
		int min = get(0, j);
		int max = min;
		for (int i = 1; i < numRow; i++) {
			if (get(i, j) < min) {
				min = get(i, j);
			} else if (get(i, j) > max) {
				max = get(i, j);
			}
		}
		for (int i = 0; i < numRow; i++) {
			set(i, j, (b - a) * ((get(i, j) - min) / (max - min)) + a);
		}
	}
}

void Matrix::rowToVector(int row, vector<double>& v) {
	if (row >= numRow || row < 0) {
		cerr << "Invalid Row (rowToVector)" << endl;
		throw exception();
	} else {
		v = m.at(row);
	}
}

void Matrix::colToVector(int col, vector<double>& v) {
	if (col >= numCol || col < 0) {
		cerr << "Invalid Column (colToVector)" << endl;
		throw exception();
	} else {
		for (int j = 0; j < numRow; j++) {
			v.push_back(m.at(j).at(col));
		}
	}
}

int Matrix::getNumRow() const {
	return numRow;
}

}
```
