## Supplementary Data 1 for "FASTCAR: Rapid alignment-free prediction of sequence alignment identity scores using self-supervised general linear models": Matrix.h

```
/*
 * matrix.h
 *
 * Created on: May 10, 2017
 * Author: Robert Geraghty, The Bioinformatics Toolsmith Laboratory, The University of Tulsa
 * Modified by Benjamin T James
 */


#ifndef MATRIX_H_
#define MATRIX_H_

#include <vector>
#include <string>

namespace matrix {

class Matrix
{
private:
	std::vector<std::vector<double> > m;
	int numRow;
	int numCol;


public:
	Matrix(std::vector<double> m);
	Matrix(int r, int c);
	Matrix();
	~Matrix();
	Matrix operator+(Matrix n);
	Matrix operator-(Matrix n);
	Matrix operator*(Matrix n);
	Matrix transpose();
	Matrix gaussJordanInverse();
	Matrix pseudoInverse();
	void userFill();
	double determinant();
	double get(int r, int c) const;
	void set(int r, int c, double val);
	void addRow(double);
	void addCol(double);
	void print();
	void printToFile(std::string);
	void randFill(double low, double high);
	void fileFill(std::string filename);
	void normalize(double a, double b);
	void rowToVector(int, std::vector<double>&);
	void colToVector(int, std::vector<double>&);
	int getNumRow() const;
	int getNumCol() const { return numCol; };
};
}
#endif /* MATRIX_H_ */
```
