## Supplementary Data 1 for "FASTCAR: Rapid alignment-free prediction of sequence alignment identity scores using self-supervised general linear models": Point.h

```
/* -*- C++ -*-
 *
 * Point.h
 *
 * Author: Benjamin T James
 *
 * For some reason this class was made pure virtual
 * in early development of MeShClust, making Histogram
 * and DivergencePoint both derivatives that essentially
 * did the same thing
 */
#ifndef POINT_H
#define POINT_H

#include <string>
#include "../../nonltr/ChromosomeOneDigit.h"

/*
 * Pure virtual class that defines behavior for
 * points. Has clone() and create() that allow for
 * polymorphic behavior
 */
template<class T>
class Point {
public:
	virtual ~Point() { data.clear(); };
	virtual void operator*=(double d) = 0;
	virtual void operator/=(double d) = 0;
	virtual bool operator<(Point<T>& p) const = 0;
	virtual uint64_t operator-(const Point<T>& p) const = 0;
	virtual void operator+=(Point<T>& p) = 0;
	virtual void set(Point<T>& p) = 0;
	virtual void display() const = 0;
	virtual uint64_t distance(const Point<T>& p) const = 0;
	virtual double distance_d(Point<double>& p) const = 0;
	virtual Point* clone() const = 0;
	virtual Point* create() const = 0;

	virtual void zero() = 0;
	virtual void addOne() = 0;
	virtual double distance_k1(const Point<T>& p) const = 0;
	virtual double prob_under(Point<T>& center) const = 0;
	virtual void subOne() = 0;
	virtual uint64_t getRealMagnitude() const = 0;
//	virtual T magnitude() const = 0;
	virtual bool is_to_delete() const = 0;
	virtual void set_to_delete(bool b) = 0;

	virtual Point<double>* create_double() const = 0;
	virtual void set_arg_to_this_d(Point<double>& p) const = 0;

	virtual const vector<T>& get_data() const = 0;

	void set_header(const std::string c) { header = string(c); };
	const std::string get_header() const { return header; };

	void set_data_str(const std::string& c) { data = c; };
	const std::string & get_data_str() const { return data; };

	void set_1mers(const vector<uint64_t> &vec) {
		// for (auto i = 0; i < Util::getAlphabetSize(); i++) {
		// 	one_mers[i] = vec[i];
		// }
		one_mers = vector<uint64_t>(vec);
	}

	vector<uint64_t> get_1mers() const {
		// vector<uint64_t> vec;
		// for (auto i = 0; i < Util::getAlphabetSize(); i++) {
		// 	vec.push_back(one_mers[i]);
		// }
		// return vec;
		return one_mers;
	}
	virtual unsigned long size() const = 0;
	virtual void set_id(uintmax_t c_id) = 0;//{ id = c_id; };
	virtual const uintmax_t get_id() const = 0;//{ return id; };
	virtual void set_length(unsigned long len) = 0;
	virtual unsigned long get_length() const = 0;

	// Added by Hani Z. Girgis on Oct 7 2018
	int getK(){
		return k;
	}
	void setK(int k){
		this->k = k;
	}

private:
	vector<uint64_t> one_mers;
    std::string header;
	std::string data;
	// Added by Hani Z. Girgis on Oct 7 2018
	// The k in k-mer used to build the table
	int k;
};

#endif
```
