## Supplementary Data 1 for "FASTCAR: Rapid alignment-free prediction of sequence alignment identity scores using self-supervised general linear models": Progress.cpp

```
/* -*- C++ -*-
 *
 * Progress.cpp
 *
 * Author: Benjamin T James
 *
 * Progress bar that uses carriage return '\r'
 * to seek to the beginning of a line to redraw
 */
#include "Progress.h"
#include <iostream>
Progress::Progress(long num, std::string prefix_)
{
	pmax = num;
	ended = 0;
	pcur = 0;
	old_prog = -1;
	prefix = prefix_;
	barWidth = 70 - (prefix.size()+1);
	print();
}

void Progress::print()
{
	#ifndef NOPROG
	double prog = (double)pcur / pmax;
	if (old_prog != int(prog * 100)) {
		std::cout << prefix << " [";
		int pos = barWidth * prog;
		for (int i = 0; i < barWidth; i++) {
			if (i < pos) {
				std::cout << "=";
			} else if (i == pos) {
				std::cout << ">";
			} else {
				std::cout << " ";
			}
		}
		std::cout << "] " << int(prog * 100.0) << " %\r";
		std::cout.flush();
	}
	old_prog = int(prog * 100);
	#endif
}

void Progress::end()
{
	if (!ended) {
		pcur = pmax;
		print();
		std::cout << std::endl;
	}
	ended = true;
}

void Progress::operator++()
{
	pcur++;
	print();
}
void Progress::operator++(int)
{
	print();
	pcur++;
}


void Progress::operator+=(size_t num)
{
	pcur += num;
	print();
}
```
