## Supplementary Data 1 for "FASTCAR: Rapid alignment-free prediction of sequence alignment identity scores using self-supervised general linear models": Progress.h

```
/* -*- C++ -*-
 *
 * Progress.h
 *
 * Author: Benjamin T James
 *
 * Progress bar that uses carriage return '\r'
 * to seek to the beginning of a line to redraw
 *
 */
#include <iostream>
#ifndef PROGRESS_H
#define PROGRESS_H

class Progress {
public:
	Progress(long num, std::string prefix_);
	~Progress() { end(); }
	void end();
	void operator++();
	void operator++(int);
	void operator+=(size_t);
private:
	void print();
	long pmax;
	long pcur;
	long old_prog;
	bool ended;
	std::string prefix;
	int barWidth;
};
#endif
```
