## Supplementary Data 1 for "FASTCAR: Rapid alignment-free prediction of sequence alignment identity scores using self-supervised general linear models": Runner.cpp

```
/* -*- C++ -*-
 *
 * Runner.cpp
 *
 * Author: Benjamin T James
 *
 * Runner class that parses options and controls
 * the process of the program.
 */
#include <vector>
#include <algorithm>
#include <cmath>
#include <cstring>
#include <sys/stat.h>
#include <sys/types.h>
#include <sys/sysinfo.h>
#include <cstdlib>
#include "../../nonltr/ChromListMaker.h"
#include "DivergencePoint.h"
#include "../../utility/AffineId.h"
#include "Runner.h"
#include "../../train/Predictor.h"
#include "Loader.h"
#include "bvec.h"
#include "Progress.h"
#include <omp.h>


Runner::Runner(int argc, char **argv)
{
	get_opts(argc, argv);
	srand(10);
}

int parseLine(char* line) {
	int i = strlen(line);
	const char* p = line;
	while (*p < '0' || *p > '9') p++;
	line[i-3] = '\0';
	i = atoi(p);
	return i;
}

void mem_used(std::string prefix)
{
	struct sysinfo memInfo;
	sysinfo(&memInfo);
	FILE* file = fopen("/proc/self/status", "r");
	int result = -1;
	char line[128];
	while (fgets(line, 128, file)) {
		if (strncmp(line, "VmSize:", 7) == 0) {
			result = parseLine(line);
			break;
		}
	}
	fclose(file);
	cout << prefix << ": used memory: " << result << " KB" << endl;
}

int Runner::run()
{
	if (pred64) {
		k = pred64->get_k();
	} else if (k == -1) {
		uintmax_t total_length = 0;
		uintmax_t total_num_seq = 0;
		largest_count = 0;
		Progress progress(files.size(), "Reading in sequences");
		uintmax_t num_seq = 10000;
		for (auto i = 0; i < files.size(); i++) {
			auto f = files.at(i);
			SingleFileLoader maker(f);

			progress++;
			uint64_t local_largest_count = 0;
			std::pair<std::string,std::string*> pr;
			while ((pr = maker.next()).first != "" && total_num_seq++ < num_seq) {
				total_length += pr.second->length();
			}
		}
		progress.end();
		double avg_length = (double)total_length / total_num_seq;
		k = std::max((int)(ceil(log(avg_length) / log(4)) - 1), 2);
	}
	cout << "K: " << k << endl;
// #pragma omp parallel for reduction(max:largest_count)
// 	for (size_t i = 0; i < sequences.size(); i++) {
// 		std::vector<uint64_t> values;
// 		KmerHashTable<unsigned long, uint64_t> table(k, 1);
// 		ChromosomeOneDigitDna chrom;
// 		chrom.setSequence(*sequences[i].second);
// 		chrom.setHeader(sequences[i].first);
// 		chrom.finalize();
// 		fill_table<uint64_t>(table, &chrom, values);
// 		uint64_t l_count = 0;
// 		for (auto elt : values) {
// 			if (elt > l_count) {
// 				l_count = elt;
// 			}
// 		}
// 		if (l_count > largest_count) {
// 			largest_count = l_count;
// 		}
// 		values.clear();
// 	}
// 	largest_count *= 2;
	uint64_t cap = 10000;
	std::vector<ChromosomeOneDigit* > sequences(cap);
	if (pred64 == NULL || Runner::get_datatype() == "") {
		uint64_t idx = 0;
		Progress progress(cap, "Reading in sequences");
		uint64_t largest_count = 0;


		for (auto i = 0; i < files.size(); i++) {
			auto f = files.at(i);
			SingleFileLoader maker(f);
			ChromosomeOneDigitDna* chrom = NULL;
			while ((chrom = maker.nextChrom()) != NULL && idx < cap) {

				sequences[idx] = chrom;
				idx++;
				progress++;
			}
		}
		sequences.resize(idx);

#pragma omp parallel for reduction(max:largest_count)
		for (int i = 0; i < sequences.size(); i++) {
			auto chrom = sequences[i];
			std::vector<uint64_t> values;
			KmerHashTable<unsigned long, uint64_t> table(k, 1);
			Loader<uint64_t>::fill_table(table, chrom, values);
			uint64_t l_count = *std::max_element(std::begin(values), std::end(values));
			if (l_count > largest_count) {
				largest_count = l_count;
			}
		}
		progress.end();
	} else if (pred64 != NULL) {
		sequences.clear();
		Runner::set_datatype(pred64->get_datatype());
		similarity = pred64->get_id();
	}
	if (Runner::get_datatype() != "") {
		std::string type = Runner::get_datatype();
		if (type == "uint8_t") {
			largest_count = std::numeric_limits<uint8_t>::max();
		} else if (type == "uint16_t") {
			largest_count = std::numeric_limits<uint16_t>::max();
		} else if (type == "uint32_t") {
			largest_count = std::numeric_limits<uint32_t>::max();
		} else if (type == "uint64_t") {
			largest_count = std::numeric_limits<uint64_t>::max();
		}
	}
	if (largest_count <= std::numeric_limits<uint8_t>::max()) {
		Runner::set_datatype("uint8_t");
		cout << "Using 8 bit histograms" << endl;
		return do_run<uint8_t>(sequences);
	} else if (largest_count <= std::numeric_limits<uint16_t>::max()) {
		Runner::set_datatype("uint16_t");
		cout << "Using 16 bit histograms" << endl;
		return do_run<uint16_t>(sequences);
	} else if (largest_count <= std::numeric_limits<uint32_t>::max()){
		Runner::set_datatype("uint32_t");
	       	cout << "Using 32 bit histograms" << endl;
		return do_run<uint32_t>(sequences);
	} else if (largest_count <= std::numeric_limits<uint64_t>::max()) {
		Runner::set_datatype("uint64_t");
	       	cout << "Using 64 bit histograms" << endl;
		return do_run<uint64_t>(sequences);
	} else {
		throw "Too big sequence";
	}
}


void Runner::usage(std::string progname) const
{
	int num_threads = omp_get_max_threads();
	std::cout << "Usage: " << progname << " *.fasta --query queryFile.fasta --id 0.90 [optional_arguments]" << std::endl << std::endl;
	std::cout << "Options: " << std::endl;
	std::cout << "\t" << "--id        "<<"\t" <<"identityValue" << "\t\t" << "Use this alignment identity (0.0 to 1.0) for classification" << std::endl;
	std::cout << "\t" << "-q|--query  "<<"\t" <<"queryFile.fasta" << "\t\t" << "Run the database against this query file" << std::endl;
	std::cout << "\t" << "-k|--kmer   "<<"\t" << "N"<<"\t\t\t" << "Usually calculated by going through the data and finding the ceil(log_4(Length_avg))-1,"<< std::endl;
	std::cout << "\t\t\t\t\t\t    " << "so if provided, it can save computational time. Increasing the k-mer increases memory usage four-fold."<< std::endl;
        std::cout << "\t" << "--datatype  "<<"\t" <<"uintX_t" << "\t\t\t" << "If provided, instead of running through the data another time," << std::endl;
	std::cout << "\t\t\t\t\t\t    " << "provide the maximum data type to not overflow, one of {uint8_t, uint16_t, uint32_t, uint64_t}" << std::endl;
        std::cout << "\t" << "-c|--chunk  "<<"\t" << chunk_size << "\t\t\t" << "Process N (a positive integer number) sequences at once in the multithreading model." << std::endl;
	std::cout << "\t" << "--dump      "<<"\t" <<"weights.txt" << "\t\t" << "Instead of running, only train the model(s) and dump the weights" <<  std::endl;
	std::cout << "\t" << "--no-format "<<"\t\t\t\t" << "Print the full header instead of the abbreviated header when printing output" <<  std::endl;


	std::cout << "\t" << "-o|--output "<<"\t" <<"output.search" << "\t\t" << "Output file, to which numbers 0 through [num_threads] are appended. Each file contains data computed by each thread." << std::endl;
        std::cout << "\t" << "-r|--recover"<<"\t" <<"weights.txt" << "\t\t" << "Instead of training, use a pre-computed weights file to avoid re-training" << std::endl;
	std::cout << "\t" << "-f|--feat   "<<"\t" <<"fast" << "\t\t\t"<<"Use a small,fast set of possible features (fast) or a larger, slower-to-train set of possible features (slow)"<<std::endl;
	std::cout << "\t" << "-m|--mode   "<<"\t" <<"rc"   << "\t\t\t"<<"Use the provided mode, either \"c\" for classification (print all pairs above threshold, but no provided alignment value)," << std::endl;
	std::cout << "\t\t\t\t\t\t    " << "\"r\" for regression only, meaning all pairs are printed, or" << std::endl;
	std::cout << "\t\t\t\t\t\t    " << "\"rc\" for both (default), printing all pairs above the threshold with alignment identity predictions." <<  std::endl;
	std::cout << "\t" << "-s|--sample "<<"\t" << sample_size << "\t\t\t" << "Use this many template sequences, from which 5 positive (above the --id threshold)" << std::endl;
	std::cout << "\t\t\t\t\t\t    " << "and 10 negative alignments will be generated, yielding ([sample_size] x 15) total training points." << std::endl;
        std::cout << "\t" << "--mut-type  "<<"\t" << "single"    << "\t\t\t" << "Use this mutation type to generate synthetic alignments." << std::endl;
	std::cout << "\t\t\t\t\t\t    " << "Options for mutation type are \"single\", \"nonsingle-typical\", \"both\" (for single and nonsingle-typical)," << std::endl;
	std::cout << "\t\t\t\t\t\t    " << "\"nonsingle-all\", and \"all\" (single, nonsingle, and atypical nonsingle)." <<  std::endl;
	std::cout << "\t" << "-t|--threads"<<"\t" << num_threads << "\t\t\t" << "Set the number of threads used from this number to a lower number." << std::endl << std::endl;
#ifndef VERSION
        #define VERSION "(undefined)"
        #endif
        std::cout << "Version " << VERSION << " compiled on " << __DATE__ << " " << __TIME__;
        #ifdef _OPENMP
        std::cout << " with OpenMP " << _OPENMP;
        #else
        std::cout << " without OpenMP";
        #endif
	std::cout << std::endl;
}

void Runner::get_opts(int argc, char **argv)
{
	for (int i = 1; i < argc; i++) {
		string arg = argv[i];
		if (arg == "--id" && i + 1 < argc) {
			try {
				std::string opt = argv[i+1];
				similarity = std::stod(opt);
				if (similarity <= 0 || similarity >= 1) {
					throw std::invalid_argument("");
				}
			} catch(std::exception e) {
				cerr << "Similarity must be between 0 and 1" << endl;
				exit(EXIT_FAILURE);
			}
			i++;
		} else if ((arg == "-c" || arg == "--chunk") && i + 1 < argc) {
			chunk_size = strtol(argv[i+1], NULL, 10);
			if (errno) {
				perror(argv[i+1]);
				exit(EXIT_FAILURE);
			} else if (chunk_size <= 0) {
				fprintf(stderr, "Chunk size must be greater than 0.\n");
				exit(EXIT_FAILURE);
			}
			i++;
		} else if ((arg == "--dump") && i + 1 < argc) {
			dump_str = argv[++i];
			dump = true;
		} else if (arg == "--noformat" || arg == "--no-format") {
			format = false;
		} else if ((arg == "--datatype") && i + 1 < argc) {
			std::string val = argv[++i];
			if (val == "uint8_t" || val == "8" || val == "uint8") {
				Runner::set_datatype("uint8_t");
			} else if (val == "uint16_t" || val == "16" || val == "uint16") {
				Runner::set_datatype("uint16_t");
			} else if (val == "uint32_t" || val == "32" || val == "uint32") {
				Runner::set_datatype("uint32_t");
			} else if (val == "uint64_t" || val == "64" || val == "uint64") {
				Runner::set_datatype("uint64_t");
			} else {
				cerr << "Histogram data type must have a valid data type or size: one of 8, 16, 32, 64" << endl;
				exit(EXIT_FAILURE);
			}
		} else if ((arg == "-k" || arg == "--kmer") && i + 1 < argc) {
			k = strtol(argv[i+1], NULL, 10);
			if (errno) {
				perror(argv[i+1]);
				exit(EXIT_FAILURE);
			} else if (k <= 0) {
				fprintf(stderr, "K must be greater than 0.\n");
				exit(EXIT_FAILURE);
			}
			align = false;
			i++;
		} else if ((arg == "-o" || arg == "--output") && i + 1 < argc) {
			output = string(argv[i+1]);
			i++;
		} else if ((arg == "-q" || arg == "--query") && i + 1 < argc) {
			char* qfile = argv[++i];
			struct stat st;
			stat(qfile, &st);
			if (S_ISREG(st.st_mode)) {
				qfiles.emplace_back(qfile);
			} else {
				usage(*argv);
				exit(EXIT_FAILURE);
			}
		} else if ((arg == "-r" || arg == "--recover") && i + 1 < argc) {
			recover = true;
			dump_str = argv[++i];
			pred64 = new Predictor<uint64_t>(dump_str);
			similarity = pred64->get_id();
			k = pred64->get_k();
		} else if ((arg == "-f" || arg == "--feat") && i + 1 < argc) {
			std::string val = argv[++i];
			if (val == "fast") {
				feats = PRED_FEAT_FAST;
			} else if (val == "slow") {
				feats = PRED_FEAT_FAST | PRED_FEAT_DIV;
			} else {
				cerr << "Features must be either \"fast\" or \"slow\"" << endl;
			}
		} else if ((arg == "-m" || arg == "--mode") && i + 1 < argc) {
			std::string val = argv[++i];
			if (val == "c") {
				mode |= PRED_MODE_CLASS;
			} else if (val == "r") {
				mode |= PRED_MODE_REGR;
			} else if (val == "cr" || val == "rc") {
				mode |= PRED_MODE_CLASS | PRED_MODE_REGR;
			} else {
				cerr << "Mode must be either c, r, or a combination" << endl;
				exit(EXIT_FAILURE);
			}
		} else if ((arg == "-s" || arg == "--sample") && i + 1 < argc) {
			sample_size = strtol(argv[i+1], NULL, 10);
			if (errno) {
				perror(argv[i+1]);
				exit(EXIT_FAILURE);
			} else if (sample_size <= 0) {
				fprintf(stderr, "Sample size must be greater than 0.\n");
				exit(EXIT_FAILURE);
			}
			i++;
		} else if ((arg == "--mut-type") && i + 1 < argc) {
			std::string opt = argv[i+1];
			if (opt == "all") {
				mut_type = HandleSeq::BOTH | HandleSeq::ATYPICAL;
			} else if (opt == "both") {
				mut_type = HandleSeq::BOTH;
			} else if (opt == "snp" || opt == "single") {
				mut_type = HandleSeq::SINGLE;
			} else if (opt == "nonsingle-typical") {
				mut_type = HandleSeq::NON_SINGLE;
			} else if (opt == "nonsingle-all") {
				mut_type = HandleSeq::NON_SINGLE | HandleSeq::ATYPICAL;
			} else if (opt == "all-but-reversion") {
				mut_type = HandleSeq::BOTH | HandleSeq::TRANSLOCATION;
			} else if (opt == "all-but-translocation") {
				mut_type = HandleSeq::BOTH | HandleSeq::REVERSION;
			} else {
				cerr << "Options for mutation type are \"single\", \"nonsingle-typical\", \"both\" (for single and nonsingle-typical), \"nonsingle-all\", and \"all\" (single, nonsingle, and atypical nonsingle)." << endl;
				exit(1);
			}
			i++;
		} else if ((arg == "-t" || arg == "--threads") && i + 1 < argc) {
			try {
				std::string opt = argv[i+1];
				int threads = std::stoi(opt);
				if (threads <= 0) {
					throw std::invalid_argument("");
				}
				#ifdef _OPENMP
				omp_set_num_threads(threads);
				#endif
			} catch (std::exception e) {
				cerr << "Number of threads must be greater than 0." << endl;
				exit(1);
			}

			i++;

		} else if ((arg == "-h") || (arg == "--help")) {
			usage(*argv);
			exit(EXIT_FAILURE);
		} else {
			struct stat st;
			if (stat(argv[i], &st) == 0 && S_ISREG(st.st_mode)) {
				files.push_back(argv[i]);
			} else {
				usage(*argv);
				exit(EXIT_FAILURE);
			}
		}
	}
	if (files.empty()) {
		usage(*argv);
		exit(EXIT_FAILURE);
	}
}


double global_mat[4][4] = {{1, -1, -1, -1},
			   {-1, 1, -1, -1},
			   {-1, -1, 1, -1},
			   {-1, -1, -1, 1}};
double global_sigma = -2;
double global_epsilon = -1;

template<class T>
long bin_search(const std::vector<Point<T>*> &points, size_t begin, size_t last, size_t length)
{
	if (last < begin) {
		return 0;
	}
	size_t idx = begin + (last - begin) / 2;
	if (points.at(idx)->get_length() == length) {
		while (idx > 0 && points[idx-1]->get_length() == length) {
			idx--;
		}
		return idx;
	} else if (points.at(idx)->get_length() > length) {
		if (begin == idx) { return idx; }
		return bin_search(points, begin, idx-1, length);
	} else {
		return bin_search(points, idx+1, last, length);
	}
}

std::string format_header(std::string hdr)
{
	long len = hdr.length();
	long b_idx = 0;
	if (hdr[0] == '>') {
		b_idx++;
	}
	for (long i = b_idx; i < len; i++) {
		if (hdr[i] == ' ' || hdr[i] == '\t') {
			len = i + 1;
			break;
		}
	}
	return hdr.substr(b_idx, len - b_idx);
}

template<class T>
void work(const std::vector<Point<T>*> &queries, const std::vector<Point<T>*> &pts, double similarity, Predictor<T>* pred, std::string delim, std::ofstream &out, uintmax_t &num_pred_pos, bool format)
{
	if (pts.empty()) {
		return;
	}
	uint8_t mode = pred->get_mode();
	for (auto query : queries) {
		size_t q_len = query->get_length();
		size_t begin_length = q_len * similarity;
		size_t end_length = q_len / similarity;
		size_t start = bin_search(pts, 0, pts.size()-1,
					  begin_length);

		for (size_t i = start;
		     i < pts.size() && pts[i]->get_length() <= end_length;
		     i++) {
			double sim = 0.0;
			bool cls = true;

			if (mode & PRED_MODE_CLASS) {
				cls = pred->close(pts[i], query);

			}
			if (!cls) {
				continue;
			}
			num_pred_pos++;
			if (mode & PRED_MODE_REGR) {
				sim = pred->similarity(pts[i], query);
			} else {
				sim = 1;
			}
			if (mode & PRED_MODE_CLASS) {
//				sim = (sim > similarity) ? sim : 0;
			}
			if (sim > 0) {
				if (format) {
					out << format_header(query->get_header()) << delim << format_header(pts[i]->get_header()) << delim << 100 * sim << endl;
				} else {
					out << query->get_header() << delim << pts[i]->get_header() << delim << 100 * sim << endl;
				}
			}
		}
	}
}

template<class T>
int Runner::do_run(std::vector<ChromosomeOneDigit* > &seqs)
{
	using DNA=ChromosomeOneDigit;
	using pvec = vector<Point<T> *>;
	using pmap = map<Point<T>*, pvec*>;
	srand(0xFF);
	mem_used("before do_run");
	size_t num_points = 0;
	uintmax_t _id = 0;


		// Sorting all sequences based on length
	std::sort(seqs.begin(), seqs.end(), [](DNA* a, DNA* b) {
			return a->getBase()->length() < b->getBase()->length();
		});
		cout << "sample_size: " << sample_size << endl;
		double increment = std::max(1.0, (double)seqs.size() / sample_size);
		for (double i = 0; round(i) < seqs.size(); i += increment) {
			indices.push_back(round(i));
			//	cout << "index: " << round(i) << " length: " << seqs.at(round(i)).second->length() << endl;
		}
		std::vector<Point<T>*> trpoints(indices.size());
		#pragma omp parallel for
		for (size_t i = 0; i < indices.size(); i++) {
			auto chrom = seqs.at(indices.at(i));
			Point<T>* p = Loader<T>::get_point(chrom, _id, k);
			trpoints[i] = p;
		}
		for (auto p : seqs) {
			delete p;
		}
		seqs.clear();

	indices.clear();
	mem_used("after selection");
	cout << "TRpoints.size(): " << trpoints.size() << endl;

	// std::sort(trpoints.begin(), trpoints.end(), [](const Point<T>* a, const Point<T>* b) {
	// 		return a->get_length() < b->get_length(); });

	int n_threads = omp_get_max_threads();
	Predictor<T> *pred = NULL;
	if (recover) {
		pred = new Predictor<T>(dump_str);

	} else {
		if (mode == 0) {
			cout << "No mode specified, using regression and classification by default" << endl;
			mode = PRED_MODE_REGR | PRED_MODE_CLASS;
		}
		if (feats == 0) {
			cout << "No feature set specified, using fast features by default" << endl;
			feats = PRED_FEAT_FAST;
		}
		if ((mode & PRED_MODE_CLASS) == PRED_MODE_CLASS && similarity < 0) {
			cout << "Classification specified, but no identity score given. Please supply a cutoff with \"--id\"" << endl;
			exit(EXIT_FAILURE);
		} else if (similarity < 0) {
			similarity = 0.9;
		}

		pred = new Predictor<T>(k, similarity, mode, feats, mut_type, 4);
		auto before = clock();
		mem_used("before predictor training");
		pred->train(trpoints, _id, sample_size);

		double elapsed = (clock() - before);
		elapsed /= CLOCKS_PER_SEC;
		cout << "Training time: " << elapsed << endl;
		for (auto p : trpoints) {
			delete p;
		}
		trpoints.clear();
		if (dump) {
			pred->save(dump_str, Runner::get_datatype());
			exit(0);
		}
	}
	mem_used("after predictor training");

	std::vector<std::ofstream> output_list;
	for (int i = 0; i < n_threads; i++) {
		std::ostringstream oss;
		oss << output << i;
		output_list.emplace_back(oss.str());
	}


	string delim = "\t";
	if (!format) {
		delim = "!";
	}
	uint64_t query_id_start = num_points;
	int num_query = num_points;
	Loader<T> qloader(qfiles, n_threads * num_points, chunk_size, 1, k, query_id_start);
	mem_used("before loop");
	uintmax_t num_pred_pos = 0;
	while (!qloader.done()) {
		qloader.preload(0);
		auto queries = qloader.load_next(0);
		Loader<T> loader(files, 0, chunk_size, n_threads, k);


		while (!loader.done()) {
			int n_iter = n_threads;
			mem_used("during inner loop");
			for (int h = 0; h < n_iter; h++) {
				loader.preload(h);
			}
			#pragma omp parallel for
			for (int h = 0; h < n_iter; h++) {
				int tid = omp_get_thread_num();
				auto pts = loader.load_next(tid);
				std::sort(std::begin(pts), std::end(pts), [](Point<T>*a, Point<T>*b) {
						return a->get_length() < b->get_length();
					});
				work(queries, pts, similarity, pred, delim, output_list[tid], num_pred_pos, format);
				for (auto p : pts) {
					delete p;
				}
			}
		}

		for (auto q : queries) {
			delete q;
		}
		mem_used("mid loop");
	}
	mem_used("after loop");
	cout << "# of predicted positive: " << num_pred_pos << endl;
	std::string warn = Loader<T>::get_warning();
	if (warn != "") {
		cout << warn << endl;
	}
	return 0;
}


template<class T>
void Runner::print_output(const map<Point<T>*, vector<Point<T>*>*> &partition) const
{
	cout << "Printing output" << endl;
	std::ofstream ofs;
	ofs.open(output, std::ofstream::out);
	int counter = 0;
	for (auto const& kv : partition) {
		if (kv.second->size() == 0) {
			continue;
		}
		ofs << ">Cluster " << counter << endl;
		int pt = 0;
		for (auto p : *kv.second) {
			string s = p->get_header();
			ofs << pt << "\t"  << p->get_length() << "nt, " << s << "... " << endl;
			pt++;
		}
		counter++;
	}
	ofs.close();
}


std::string _runner_datatype = "";

std::string Runner::get_datatype()
{
	return _runner_datatype;
}

void Runner::set_datatype(std::string s)
{
	_runner_datatype = s;
}
```
