## Supplementary Data 1 for "FASTCAR: Rapid alignment-free prediction of sequence alignment identity scores using self-supervised general linear models": Runner.h

```
/* -*- C++ -*-
 *
 * Runner.h
 *
 * Author: Benjamin T James
 *
 * Runner class, sets default params
 * and runs program
 */
#ifndef RUNNER_H
#define RUNNER_H

#include <iostream>
#include <map>
#include <set>
#include "Point.h"
#include "../../train/Predictor.h"
#include "../../train/HandleSeq.h"
#include "../../nonltr/ChromosomeOneDigitDna.h"
using namespace std;

class Runner {
public:
	Runner(int argc, char** argv);
	~Runner() { indices.clear(); files.clear(); qfiles.clear(); if (pred64) {delete pred64;}};
	int run();
	static std::string get_datatype();
private:
	void usage(std::string progname) const;
	template<class T> int do_run(std::vector<ChromosomeOneDigit*> &sequences);
	template<class T> void print_output(const map<Point<T>*, vector<Point<T>*>*> &m) const;
	static void set_datatype(std::string);
	int k = -1;
        int bandwidth;
	double similarity = -1;
	long largest_count = 0;
	bool align = false;
	bool recover = false;
	int sample_size = 300;
	int mut_type = HandleSeq::SINGLE;
	uint8_t mode = 0;
	uint64_t feats = 0;
	uint64_t chunk_size = 10000;
	std::vector<std::string> files, qfiles;
	std::vector<size_t> indices;
	bool dump = false;
	bool format = true;
	string output = "output.search";
	string dump_str = "weights.txt";
	void get_opts(int argc, char** argv);
	Predictor<uint64_t> *pred64 = NULL;


};
#endif
```
