## Supplementary Data 1 for "FASTCAR: Rapid alignment-free prediction of sequence alignment identity scores using self-supervised general linear models": SingleFeature.cpp

```
/* -*- C++ -*-
 *
 * SingleFeature.cpp
 *
 * Author: Benjamin T James
 *
 * Old method of normalizing features from MeShClust
 */
#include "SingleFeature.h"

template<class T>
void SingleFeature<T>::normalize(const vector<pair<Point<T>*,Point<T>*> > &pairs)
{
	for (auto p : pairs) {
		double d;
		if (rc.empty()) {
			d = raw(p.first, p.second);
		} else {
			d = rraw(p.first, p.second, rc, rv);
		}
		if (!min_set || d < min) {
			min = d;
			min_set = true;
		}
		if (!max_set || d > max) {
			max = d;
			max_set = true;
		}
	}
}

template<class T>
double SingleFeature<T>::operator()(Point<T> *a, Point<T> *b) const
{
	double d;
	if (rc.empty()) {
		d = raw(a, b);
	} else {
		d = rraw(a, b, rc, rv);
	}
//	std::cout << "Raw: " << d << std::endl;
	double f = (d - min) / (max - min);
//	std::cout << "Normalized: " << f << std::endl;
	f = std::min(1.0, std::max(0.0, f));
	if (is_sim) {
		return f;
	} else {
		return 1.0 - f;
	}
}


template class SingleFeature<uint8_t>;
template class SingleFeature<uint16_t>;
template class SingleFeature<uint32_t>;
template class SingleFeature<uint64_t>;
template class SingleFeature<int>;
template class SingleFeature<double>;
```
