## Supplementary Data 1 for "FASTCAR: Rapid alignment-free prediction of sequence alignment identity scores using self-supervised general linear models": SingleFeature.h

```
/* -*- C++ -*-
 *
 * SingleFeature.h
 *
 * Author: Benjamin T James
 *
 * Old method of normalizing features from MeShClust
 */
#ifndef SINGLEFEATURE_H
#define SINGLEFEATURE_H

#include "Point.h"
#include <functional>

template<class T>
class SingleFeature {
public:
	SingleFeature(std::function<double(Point<T>*, Point<T>*)> f, bool is_sim_=true)
		: raw(f), is_sim(is_sim_), min_set(false), max_set(false) {}
	SingleFeature(std::function<double(Point<T>*, Point<T>*, const vector<int>&, const vector<int>&)> f, vector<int> rrv, vector<int> rrc, bool is_sim_=true)
		: rraw(f), is_sim(is_sim_), min_set(false), max_set(false), rv(rrv), rc(rrc) {}
	void normalize(const vector<pair<Point<T>*,Point<T>*> > &pairs);
	double operator()(Point<T>*, Point<T>*) const;
	double min, max;
private:
	std::function<double(Point<T>*, Point<T>*)> raw;
	std::function<double(Point<T>*, Point<T>*, const vector<int>&, const vector<int>&)> rraw;
	vector<int> rv, rc;
	const bool is_sim;
	bool max_set, min_set;

};

#endif
```
