## Supplementary Data 1 for "FASTCAR: Rapid alignment-free prediction of sequence alignment identity scores using self-supervised general linear models": SingleFileLoader.cpp

```
/* -*- C++ -*-
 *
 * SingleFileLoader.cpp
 *
 * Author: Benjamin T James
 *
 * Reads sequences one by one from a file
 */
#include "SingleFileLoader.h"
#include <sstream>
#include <iostream>

std::istream& safe_getline(std::istream& is, std::string& t)
{
	t.clear();
	std::istream::sentry se(is, true);
	std::streambuf* sb = is.rdbuf();
	for(;;) {
		int c = sb->sbumpc();
		switch (c) {
		case '\n':
			return is;
		case '\r':
			if (sb->sgetc() == '\n') {
				sb->sbumpc();
			}
			return is;
		case std::streambuf::traits_type::eof():
			if (t.empty()) {
				is.setstate(std::ios::eofbit);
			}
			return is;
		default:
			t += (char)c;
		}
	}
}


SingleFileLoader::SingleFileLoader(std::string filename)
{
	in = new std::ifstream(filename);
	is_first = true;
}
std::pair<std::string, std::string*> SingleFileLoader::next()
{
	std::pair<std::string,std::string*> ret = std::make_pair("", (std::string*)NULL);
	if (!in->good()) {
		return ret;
	}
	clock_t begin = clock();
	ret.second = new std::string("");
	if (is_first) {
		safe_getline(*in, buffer);
		is_first = false;
	}
	do {
		if (buffer[0] == '>') {
			if (ret.first != "")  {
				return ret;
			}
			ret.first = buffer;
		} else if (buffer[0] == ' ' || buffer[0] == '\t') {
			bool all_spaces = true;
			for (auto c : buffer) {
				if (c != ' ' && c != '\t') {
					all_spaces = false;
				}
			}
			if (!all_spaces) {
				std::ostringstream oss;
				oss << ret.first << buffer;
				std::string new_header = oss.str();
				ret.first = new_header;
			}
		} else {
			ret.second->append(buffer);
		}
		safe_getline(*in, buffer);
	} while (in->good());
	double diff = clock() - begin;
//	std::cout << "next(): " << diff / CLOCKS_PER_SEC << std::endl;
	return ret;
}
ChromosomeOneDigitDna* SingleFileLoader::nextChrom()
{
	ChromosomeOneDigitDna* ret = NULL;
	if (!in->good()) {
		return ret;
	}
	if (is_first) {
		safe_getline(*in, buffer);
		is_first = false;
	}
	do {
		if (buffer[0] == '>') {
			if (ret != NULL)  {
				ret->finalize();
				return ret;
			}
			ret = new ChromosomeOneDigitDna();
			ret->setHeader(buffer);
		} else if (buffer[0] == ' ' || buffer[0] == '\t') {
			bool all_spaces = true;
			for (auto c : buffer) {
				if (c != ' ' && c != '\t') {
					all_spaces = false;
				}
			}
			if (!all_spaces) {
				std::ostringstream oss;
				oss << ret->getHeader() << buffer;
				std::string new_header = oss.str();
				ret->setHeader(new_header);
			}
		} else {
			ret->appendToSequence(buffer);
		}
		safe_getline(*in, buffer);
	} while (in->good());
	ret->finalize();
	return ret;
}
```
