## Supplementary Data 1 for "FASTCAR: Rapid alignment-free prediction of sequence alignment identity scores using self-supervised general linear models": SingleFileLoader.h

```
/* -*- C++ -*-
 *
 * SingleFileLoader.h
 *
 * Author: Benjamin T James
 *
 * A way of reading in 1 sequence at a time
 * from FASTA, sequence is heap allocated
 */
#ifndef SINGLEFILELOADER_H
#define SINGLEFILELOADER_H

#include <fstream>
#include "../../nonltr/ChromosomeOneDigitDna.h"
class SingleFileLoader {
public:
	SingleFileLoader(std::string file);
	~SingleFileLoader() {
		if (in != NULL) {
			delete in;
		}
	}
	std::pair<std::string,std::string*> next();
	ChromosomeOneDigitDna* nextChrom();
private:
	std::ifstream *in;
	std::string buffer;
	bool is_first;
};
#endif
```
