## Supplementary Data 1 for "FASTCAR: Rapid alignment-free prediction of sequence alignment identity scores using self-supervised general linear models": FileDoesNotExistException.cpp

```
/*
 * FileDoesNotExistException.cpp
 *
 *  Created on: Apr 30, 2012
 *      Author: Hani Zakaria Girgis, PhD
 */

#include "FileDoesNotExistException.h"

#include <iostream>
#include <string>

using namespace std;

namespace exception{

FileDoesNotExistException::FileDoesNotExistException(string massage) {
	cerr << "File Does Not Exist Exception" << endl;
	cerr << massage << endl;
}

FileDoesNotExistException::~FileDoesNotExistException() {
	// TODO Auto-generated destructor stub
}
}
```
