## Supplementary Data 1 for "FASTCAR: Rapid alignment-free prediction of sequence alignment identity scores using self-supervised general linear models": FileDoesNotExistException.h

```
/*
 * FileDoesNotExistException.h
 *
 *  Created on: Apr 30, 2012
 *      Author: Hani Zakaria Girgis, PhD
 */

#ifndef FILEDOESNOTEXISTEXCEPTION_H_
#define FILEDOESNOTEXISTEXCEPTION_H_

#include <string>

using namespace std;

namespace exception {
	class FileDoesNotExistException {
	public:
		FileDoesNotExistException(string);
		~FileDoesNotExistException();
	};
}

#endif /* FILEDOESNOTEXISTEXCEPTION_H_ */
```
