## Supplementary Data 1 for "FASTCAR: Rapid alignment-free prediction of sequence alignment identity scores using self-supervised general linear models": InvalidInputException.cpp

```
/*
 * InvalidInputException.cpp
 *
 *  Created on: May 1, 2012
 *      Author: Hani Zakaria Girgis, PhD
 */

#include "InvalidInputException.h"

#include <string>
#include <iostream>

using namespace std;
namespace exception{

InvalidInputException::InvalidInputException(string msg) {
	cerr << "Invalid Input Exception" << endl;
	cerr << msg << endl;
}

InvalidInputException::~InvalidInputException() {
	// TODO Auto-generated destructor stub
}
}
```
