## Supplementary Data 1 for "FASTCAR: Rapid alignment-free prediction of sequence alignment identity scores using self-supervised general linear models": InvalidInputException.h

```
/*
 * InvalidInputException.h
 *
 *  Created on: May 1, 2012
 *      Author: Hani Zakaria Girgis, PhD
 */

#ifndef INVALIDINPUTEXCEPTION_H_
#define INVALIDINPUTEXCEPTION_H_

#include<string>

using namespace std;

namespace exception {
	class InvalidInputException {
	public:
		InvalidInputException(string);
		~InvalidInputException();
	};
}

#endif /* INVALIDINPUTEXCEPTION_H_ */
```
