## Supplementary Data 1 for "FASTCAR: Rapid alignment-free prediction of sequence alignment identity scores using self-supervised general linear models": InvalidOperationException.cpp

```
/*
 * InvalidOperationException.cpp
 *
 *  Created on: Dec 20, 2012
 *      Author: Hani Zakaria Girgis, PhD
 */

#include <iostream>
#include "InvalidOperationException.h"


namespace exception {

InvalidOperationException::InvalidOperationException(string msg) : std::runtime_error(msg) {
	cerr << "Invalid Operation Exception." << endl;
	cerr << what() << endl;
}

}
```
