## Supplementary Data 1 for "FASTCAR: Rapid alignment-free prediction of sequence alignment identity scores using self-supervised general linear models": InvalidOperationException.h

```
/*
 * InvalidOperationException.h
 *
 *  Created on: Dec 20, 2012
 *      Author: Hani Zakaria Girgis, PhD
 */

#ifndef INVALIDOPERATIONEXCEPTION_H_
#define INVALIDOPERATIONEXCEPTION_H_

#include <string>
#include <stdexcept>

using namespace std;

namespace exception {

class InvalidOperationException : public std::runtime_error{
public:
	InvalidOperationException(string msg);
	//virtual ~InvalidOperationException();
};

}

#endif /* INVALIDOPERATIONEXCEPTION_H_ */
```
