## Supplementary Data 1 for "FASTCAR: Rapid alignment-free prediction of sequence alignment identity scores using self-supervised general linear models": InvalidOrderOfOperationsException.cpp

```
/*
 * InvalidOrderOfOperationsException.cpp
 *
 *  Created on: Apr 26, 2012
 *      Author: Hani Zakaria Girgis, PhD
 */

#include "InvalidOrderOfOperationsException.h"

#include <string>
#include <iostream>

using namespace std;
namespace exception{

InvalidOrderOfOperationsException::InvalidOrderOfOperationsException(string massage) {
	cerr << "Invalid Order Of Operations Exception" << endl;
	cerr << massage << endl;
}

InvalidOrderOfOperationsException::~InvalidOrderOfOperationsException() {
	// TODO Auto-generated destructor stub
}
}
```
