## Supplementary Data 1 for "FASTCAR: Rapid alignment-free prediction of sequence alignment identity scores using self-supervised general linear models": InvalidOrderOfOperationsException.h

```
/*
 * InvalidOrderOfOperationsException.h
 *
 *  Created on: Apr 26, 2012
 *      Author: Hani Zakaria Girgis, PhD
 */

#ifndef INVALIDORDEROFOPERATIONSEXCEPTION_H_
#define INVALIDORDEROFOPERATIONSEXCEPTION_H_

#include <string>

using namespace std;

namespace exception{
	class InvalidOrderOfOperationsException {
	public:
		InvalidOrderOfOperationsException(string);
		~InvalidOrderOfOperationsException();
	};
}

#endif /* INVALIDORDEROFOPERATIONSEXCEPTION_H_ */
```
