## Supplementary Data 1 for "FASTCAR: Rapid alignment-free prediction of sequence alignment identity scores using self-supervised general linear models": InvalidScoreException.cpp

```
/*
 * InvalidScoreException.cpp
 *
 *  Created on: Apr 27, 2012
 *      Author: Hani Zakaria Girgis, PhD
 */

#include "InvalidScoreException.h"

#include <string>
#include <iostream>

using namespace std;
namespace exception{

InvalidScoreException::InvalidScoreException(string massage) {
	cerr << "Invalid Score Exception." << endl;
	cerr << massage << endl;
}

InvalidScoreException::~InvalidScoreException() {
	// TODO Auto-generated destructor stub
}
}
```
