## Supplementary Data 1 for "FASTCAR: Rapid alignment-free prediction of sequence alignment identity scores using self-supervised general linear models": InvalidScoreException.h

```
/*
 * InvalidScoreException.h
 *
 *  Created on: Apr 27, 2012
 *      Author: Hani Zakaria Girgis, PhD
 */

#ifndef INVALIDSCOREEXCEPTION_H_
#define INVALIDSCOREEXCEPTION_H_

#include <string>

using namespace std;

namespace exception{
	class InvalidScoreException {
	public:
		InvalidScoreException(string);
		virtual ~InvalidScoreException();
	};
}

#endif /* INVALIDSCOREEXCEPTION_H_ */
```
