## Supplementary Data 1 for "FASTCAR: Rapid alignment-free prediction of sequence alignment identity scores using self-supervised general linear models": InvalidStateException.cpp

```
/*
 * InvalidStateException.cpp
 *
 *  Created on: Aug 9, 2012
 *      Author: Hani Zakaria Girgis, PhD
 */

#include <iostream>
#include <string>
#include "InvalidStateException.h"

using namespace std;


namespace exception {
InvalidStateException::InvalidStateException(string msg) :
		std::runtime_error(msg) {
	cerr << "Invalid State Exception." << endl;
	cerr << what() << endl;
}
}

//InvalidStateException::~InvalidStateException() {
// TODO Auto-generated destructor stub
//}
```
