## Supplementary Data 1 for "FASTCAR: Rapid alignment-free prediction of sequence alignment identity scores using self-supervised general linear models": InvalidStateException.h

```
/*
 * InvalidStateException.h
 *
 *  Created on: Aug 9, 2012
 *      Author: Hani Zakaria Girgis, PhD
 */

#ifndef INVALIDSTATEEXCEPTION_H_
#define INVALIDSTATEEXCEPTION_H_

#include <string>
#include <stdexcept>

using namespace std;

namespace exception {
	class InvalidStateException : public std::runtime_error{
	public:
		InvalidStateException(string);
	};
}

#endif /* INVALIDSTATEEXCEPTION_H_ */
```
