## Supplementary Data 1 for "FASTCAR: Rapid alignment-free prediction of sequence alignment identity scores using self-supervised general linear models": Makefile

```
# CXX = /usr/bin/c++
CXX ?= g++

CXXFLAGS =	-g -Wall -O3 -march=native

#
# Objects
#

ORed  = ../bin/Red.o

# Exception
OInvalidInputException  = ../bin/exception/InvalidInputException.o
OInvalidStateException  = ../bin/exception/InvalidStateException.o
OFileDoesNotExistException  = ../bin/exception/FileDoesNotExistException.o
OInvalidOrderOfOperationsException  = ../bin/exception/InvalidOrderOfOperationsException.o
OInvalidScoreException  = ../bin/exception/InvalidScoreException.o
OInvalidOperationException = ../bin/exception/InvalidOperationException.o

# Utility
OUtil = ../bin/utility/Util.o
OLocation = ../bin/utility/Location.o
OEmptyLocation = ../bin/utility/EmptyLocation.o
OLCSLen = ../bin/utility/LCSLen.o
OAffineId = ../bin/utility/AffineId.o
OGlobAlignE = ../bin/utility/GlobAlignE.o

# Non TR
OChromosome  = ../bin/nonltr/Chromosome.o
OChromosomeOneDigit = ../bin/nonltr/ChromosomeOneDigit.o
OChromosomeOneDigitDna = ../bin/nonltr/ChromosomeOneDigitDna.o
OChromosomeOneDigitProtein = ../bin/nonltr/ChromosomeOneDigitProtein.o
OChromosomeRandom = ../bin/nonltr/ChromosomeRandom.o
OChromListMaker = ../bin/nonltr/ChromListMaker.o
OTableBuilder = ../bin/nonltr/TableBuilder.o
OScorer = ../bin/nonltr/Scorer.o
ODetectorMaxima = ../bin/nonltr/DetectorMaxima.o
OChromDetectorMaxima =  ../bin/nonltr/ChromDetectorMaxima.o
OHMM = ../bin/nonltr/HMM.o
OScanner = ../bin/nonltr/Scanner.o
OTrainer = ../bin/nonltr/Trainer.o
OLocationList = ../bin/nonltr/LocationList.o
OLocationListCollection = ../bin/nonltr/LocationListCollection.o

OBJS = $(ORed) $(OInvalidInputException) $(OInvalidStateException) $(OFileDoesNotExistException) $(OInvalidOrderOfOperationsException) $(OInvalidOperationException) $(OInvalidScoreException) $(OUtil) $(OLocation) $(OEmptyLocation) $(OChromosome) $(OChromosomeOneDigit) $(OChromosomeRandom) $(OChromListMaker) $(OTableBuilder) $(OScorer) $(ODetectorMaxima) $(OChromDetector) $(OChromDetectorMaxima) $(OHMM) $(OScanner) $(OTrainer) $(OLocationList) $(OLocationListCollection) $(OLCSLen) $(OAffineId) $(OGlobAlignE) $(OChromosomeOneDigitProtein) $(OChromosomeOneDigitDna)

#
# Target
#

TRed = ../bin/Red

#
# Make RepeatsDetector
#

$(TRed): $(OBJS)
	$(CXX) -o $(TRed) $(OBJS)

#
# RepeatsDetector
#

$(ORed): RepeatsDetector.cpp nonltr/KmerHashTable.h nonltr/KmerHashTable.cpp nonltr/TableBuilder.h nonltr/HMM.h nonltr/Scanner.h nonltr/Trainer.h utility/Util.h
	$(CXX) $(CXXFLAGS) -c RepeatsDetector.cpp -o $(ORed)

#
# Exception
#
$(OInvalidInputException): exception/InvalidInputException.cpp exception/InvalidInputException.h
	$(CXX) $(CXXFLAGS) -c exception/InvalidInputException.cpp -o $(OInvalidInputException)

$(OInvalidStateException): exception/InvalidStateException.cpp exception/InvalidStateException.h
	$(CXX) $(CXXFLAGS) -c exception/InvalidStateException.cpp -o $(OInvalidStateException)

$(OFileDoesNotExistException): exception/FileDoesNotExistException.cpp exception/FileDoesNotExistException.h
	$(CXX) $(CXXFLAGS) -c exception/FileDoesNotExistException.cpp -o $(OFileDoesNotExistException)

$(OInvalidOrderOfOperationsException): exception/InvalidOrderOfOperationsException.cpp exception/InvalidOrderOfOperationsException.h
	$(CXX) $(CXXFLAGS) -c exception/InvalidOrderOfOperationsException.cpp -o $(OInvalidOrderOfOperationsException)

$(OInvalidScoreException): exception/InvalidScoreException.cpp exception/InvalidScoreException.h
	$(CXX) $(CXXFLAGS) -c exception/InvalidScoreException.cpp -o $(OInvalidScoreException)

$(OInvalidOperationException): exception/InvalidOperationException.cpp exception/InvalidOperationException.h
	$(CXX) $(CXXFLAGS) -c exception/InvalidOperationException.cpp -o $(OInvalidOperationException)

#
# Utility
#

$(OUtil): utility/Util.cpp utility/Util.h utility/Location.h exception/FileDoesNotExistException.h
	$(CXX) $(CXXFLAGS) -c utility/Util.cpp -o $(OUtil)

$(OLocation): utility/Location.cpp utility/Location.h utility/ILocation.h exception/InvalidInputException.h utility/Util.h
	$(CXX) $(CXXFLAGS) -c utility/Location.cpp -o $(OLocation)

$(OEmptyLocation): utility/EmptyLocation.cpp utility/EmptyLocation.h utility/ILocation.h exception/InvalidOperationException.h
	$(CXX) $(CXXFLAGS) -c utility/EmptyLocation.cpp -o $(OEmptyLocation)

$(OLCSLen): utility/LCSLen.cpp utility/LCSLen.h
	$(CXX) $(CXXFLAGS) -c utility/LCSLen.cpp -o $(OLCSLen)

$(OAffineId): utility/AffineId.cpp utility/AffineId.h
	$(CXX) $(CXXFLAGS) -c utility/AffineId.cpp -o $(OAffineId)

$(OGlobAlignE): utility/GlobAlignE.cpp utility/GlobAlignE.h
	$(CXX) $(CXXFLAGS) -c utility/GlobAlignE.cpp -o $(OGlobAlignE)
#
# Non LTR
#

$(OChromosome): nonltr/Chromosome.cpp nonltr/Chromosome.h nonltr/IChromosome.h utility/Util.h exception/InvalidInputException.h exception/InvalidOperationException.h
	$(CXX) $(CXXFLAGS) -c nonltr/Chromosome.cpp -o $(OChromosome)

$(OChromosomeOneDigit): nonltr/ChromosomeOneDigit.cpp nonltr/ChromosomeOneDigit.h nonltr/Chromosome.h exception/InvalidInputException.h
	$(CXX) $(CXXFLAGS) -c nonltr/ChromosomeOneDigit.cpp	-o $(OChromosomeOneDigit)

$(OChromosomeOneDigitDna): nonltr/ChromosomeOneDigitDna.cpp nonltr/ChromosomeOneDigitDna.h nonltr/Chromosome.h exception/InvalidInputException.h
	$(CXX) $(CXXFLAGS) -c nonltr/ChromosomeOneDigitDna.cpp	-o $(OChromosomeOneDigitDna)

$(OChromosomeOneDigitProtein): nonltr/ChromosomeOneDigitProtein.cpp nonltr/ChromosomeOneDigitProtein.h nonltr/Chromosome.h exception/InvalidInputException.h
	$(CXX) $(CXXFLAGS) -c nonltr/ChromosomeOneDigitProtein.cpp	-o $(OChromosomeOneDigitProtein)

$(OChromosomeRandom): nonltr/ChromosomeRandom.cpp nonltr/ChromosomeRandom.h nonltr/IChromosome.h exception/InvalidInputException.h exception/InvalidStateException.h utility/Util.h
	$(CXX) $(CXXFLAGS) -c nonltr/ChromosomeRandom.cpp -o $(OChromosomeRandom)

$(OTableBuilder): nonltr/TableBuilder.cpp nonltr/TableBuilder.h utility/Util.h nonltr/ChromosomeOneDigit.h nonltr/ITableView.h nonltr/KmerHashTable.h nonltr/KmerHashTable.cpp nonltr/EnrichmentMarkovView.h nonltr/EnrichmentMarkovView.cpp exception/InvalidStateException.h nonltr/ChromListMaker.h nonltr/IChromosome.h
	$(CXX) $(CXXFLAGS) -c nonltr/TableBuilder.cpp -o $(OTableBuilder)

$(OScorer): nonltr/Scorer.cpp nonltr/Scorer.h nonltr/ChromosomeOneDigit.h utility/Util.h exception/InvalidStateException.h
	$(CXX) $(CXXFLAGS) -c nonltr/Scorer.cpp -o $(OScorer)

$(ODetectorMaxima): nonltr/DetectorMaxima.cpp nonltr/DetectorMaxima.h utility/ILocation.h exception/InvalidStateException.h
	$(CXX) $(CXXFLAGS) -c nonltr/DetectorMaxima.cpp -o $(ODetectorMaxima)

$(OChromDetectorMaxima): nonltr/ChromDetectorMaxima.cpp nonltr/ChromDetectorMaxima.h nonltr/DetectorMaxima.h nonltr/ChromosomeOneDigit.h utility/Util.h utility/ILocation.h utility/Location.h
	$(CXX) $(CXXFLAGS) -c nonltr/ChromDetectorMaxima.cpp -o $(OChromDetectorMaxima)

$(OHMM): nonltr/HMM.cpp nonltr/HMM.h utility/ILocation.h exception/InvalidStateException.h exception/InvalidInputException.h exception/FileDoesNotExistException.h exception/InvalidOperationException.h
	$(CXX) $(CXXFLAGS) -c nonltr/HMM.cpp -o $(OHMM)

$(OScanner): nonltr/Scanner.cpp nonltr/Scanner.h nonltr/Chromosome.h nonltr/ChromosomeOneDigit.h nonltr/HMM.h nonltr/ITableView.h nonltr/Scorer.h utility/Util.h utility/ILocation.h exception/InvalidInputException.h exception/InvalidStateException.h exception/FileDoesNotExistException.h exception/InvalidOperationException.h
	$(CXX) $(CXXFLAGS) -c nonltr/Scanner.cpp -o $(OScanner)

$(OTrainer): nonltr/Trainer.cpp nonltr/Trainer.h nonltr/TableBuilder.h nonltr/KmerHashTable.h nonltr/KmerHashTable.cpp nonltr/HMM.h nonltr/ChromDetectorMaxima.h nonltr/Scorer.h nonltr/ChromListMaker.h utility/Util.h nonltr/LocationListCollection.h
	$(CXX) $(CXXFLAGS) -c nonltr/Trainer.cpp -o $(OTrainer)

$(OChromListMaker): nonltr/ChromListMaker.cpp nonltr/ChromListMaker.h nonltr/Chromosome.h nonltr/ChromosomeOneDigit.h utility/Util.h
	$(CXX) $(CXXFLAGS) -c nonltr/ChromListMaker.cpp -o $(OChromListMaker)

$(OCluster): nonltr/Cluster.cpp nonltr/Cluster.h utility/Util.h exception/InvalidStateException.h exception/InvalidInputException.h
	$(CXX) $(CXXFLAGS) -c nonltr/Cluster.cpp -o $(OCluster)

$(OLocationList): nonltr/LocationList.cpp nonltr/LocationList.h utility/ILocation.h utility/Location.h exception/InvalidStateException.h
	$(CXX) $(CXXFLAGS) -c nonltr/LocationList.cpp -o $(OLocationList)

$(OLocationListCollection): nonltr/LocationListCollection.cpp nonltr/LocationListCollection.h utility/Location.h exception/InvalidStateException.h
	$(CXX) $(CXXFLAGS) -c nonltr/LocationListCollection.cpp -o $(OLocationListCollection)


#
# Make binary directories
#

red: $(TRed)

#
# Make Red
#

bin:
	mkdir ../bin
	mkdir ../bin/exception
	mkdir ../bin/utility
	mkdir ../bin/nonltr

#
# Make clean
#

clean:
	rm -f ../bin/*.o ../bin/exception/*.o ../bin/ms/*.o ../bin/nonltr/*.o ../bin/test/*.o ../bin/utility/*.o ../bin/tr/*.o *.o $(TRed)
```
