## Supplementary Data 1 for "FASTCAR: Rapid alignment-free prediction of sequence alignment identity scores using self-supervised general linear models": ChromDetector.cpp

```
/*
 * ChromDetector.cpp
 *
 *  Created on: Nov 8, 2012
 *      Author: Hani Zakaria Girgis, PhD
 */

#include <vector>

#include "ChromDetector.h"
#include "Detector.h"
#include "../utility/Util.h"

using namespace std;
using namespace nonltr;
using namespace utility;

ChromDetector::ChromDetector(double s, double w, double pDelta, double b,
		double mDelta, vector<int> * scores,
		const vector<vector<int> *> * segmentList) {

	regions = new vector<vector<int> *>();

	for (int i = 0; i < segmentList->size(); i++) {
		Detector * detector = new Detector(segmentList->at(i)->at(0),
				segmentList->at(i)->at(1), s, w, pDelta, b, mDelta, scores);
		vector<vector<int> *> * segRegions = detector->getRegions();
		regions->insert(regions->end(), segRegions->begin(), segRegions->end());
		delete detector;
	}
}

ChromDetector::~ChromDetector() {
	Util::deleteInVector(regions);
	regions->clear();
	delete regions;
}

vector<vector<int> *> * ChromDetector::getRegions() {
	return regions;
}
```
