## Supplementary Data 1 for "FASTCAR: Rapid alignment-free prediction of sequence alignment identity scores using self-supervised general linear models": ChromDetector.h

```
/*
 * ChromDetector.h
 *
 *  Created on: Nov 8, 2012
 *      Author: Hani Zakaria Girgis, PhD
 */

#ifndef CHROMDETECTOR_H_
#define CHROMDETECTOR_H_

#include <vector>

using namespace std;

namespace nonltr{
class ChromDetector {

private:
	vector<vector<int> *> * regions;

public:
	ChromDetector(double, double, double, double, double, vector<int> *,
			const vector<vector<int> *> *);
	virtual ~ChromDetector();
	vector<vector<int> *> * getRegions();
};
}

#endif /* CHROMDETECTOR_H_ */
```
