## Supplementary Data 1 for "FASTCAR: Rapid alignment-free prediction of sequence alignment identity scores using self-supervised general linear models": ChromDetectorMaxima.cpp

```
/*
 * ChromDetectorMaxima.cpp
 *
 *  Created on: Jun 6, 2013
 *      Author: Hani Zakaria Girgis, PhD
 */

#include "ChromDetectorMaxima.h"

namespace nonltr {

ChromDetectorMaxima::ChromDetectorMaxima(double s, double w, double m,
		double t, double p, int e, vector<int> * oScores,
		ChromosomeOneDigit * chrom) {
	header = chrom->getHeader();
	start(s, w, m, t, p, e, oScores, chrom->getSegment());

}

ChromDetectorMaxima::ChromDetectorMaxima(double s, double w, double m,
		double t, double p, int e, vector<int> * oScores, const vector<vector<
				int> *> * segmentList) {
	header = string("chrUnknown");
	start(s, w, m, t, p, e, oScores, segmentList);
}

void ChromDetectorMaxima::start(double s, double w, double m, double t,
		double p, int e, vector<int> * oScores,
		const vector<vector<int> *> * segmentList) {

	regionList = new vector<ILocation *> ();

	int segmentCount = segmentList->size();
	for (int i = 0; i < segmentCount; i++) {
		int segStart = segmentList->at(i)->at(0);
		int segEnd = segmentList->at(i)->at(1);

		// The effective length is shorter than the actual length by 2w
		int effLen = 2 * w + 10;
		int segLen = segEnd - segStart + 1;

		if (segLen > effLen) {
			DetectorMaxima * detector = new DetectorMaxima(segStart, segEnd, s,
					w, m, t, p, e, oScores);

			const vector<ILocation *> * segRegions = detector->getRegionList();
			int segRegionCount = segRegions->size();
			for (int h = 0; h < segRegionCount; h++) {
				regionList->push_back(new Location(*(segRegions->at(h))));
			}

			delete detector;
		} else {
			cout << "\tSkipping a short segment: ";
			cout << segStart << "-" << segEnd << endl;
		}
	}
}

ChromDetectorMaxima::~ChromDetectorMaxima() {
	Util::deleteInVector(regionList);
	regionList->clear();
	delete regionList;
}

void ChromDetectorMaxima::printIndex(string outputFile) {
	printIndex(outputFile, false);
}

void ChromDetectorMaxima::printIndex(string outputFile, bool canAppend) {
	ofstream outIndex;

	if (canAppend) {
		outIndex.open(outputFile.c_str(), ios::out | ios::app);
	} else {
		outIndex.open(outputFile.c_str(), ios::out);
	}

	// Write the index of the repeat segment [x,y[
	for (int j = 0; j < regionList->size(); j++) {
		outIndex << header << ":";
		outIndex << ((int) (regionList->at(j)->getStart())) << "-";
		outIndex << ((int) (regionList->at(j)->getEnd() + 1)) << " ";
		outIndex << endl;
	}

	outIndex.close();
}

const vector<ILocation*>* ChromDetectorMaxima::getRegionList() const {
	return regionList;
}

} /* namespace nonltr */
```
