## Supplementary Data 1 for "FASTCAR: Rapid alignment-free prediction of sequence alignment identity scores using self-supervised general linear models": ChromDetectorMaxima.h

```
/*
 * ChromDetectorMaxima.h
 *
 *  Created on: Jun 6, 2013
 *      Author: Hani Zakaria Girgis, PhD
 */

#ifndef CHROMDETECTORMAXIMA_H_
#define CHROMDETECTORMAXIMA_H_

#include <fstream>
#include <vector>

#include "ChromosomeOneDigit.h"
#include "DetectorMaxima.h"

#include "../utility/Util.h"
#include "../utility/ILocation.h"
#include "../utility/Location.h"

using namespace std;
using namespace utility;

namespace nonltr {

class ChromDetectorMaxima {
private:
	vector<ILocation *> * regionList;
	string header;

	void start(double, double, double, double, double, int, vector<int> *,
			const vector<vector<int> *> *);

public:
	ChromDetectorMaxima(double, double, double, double, double, int,
			vector<int> *, ChromosomeOneDigit *);
	ChromDetectorMaxima(double, double, double, double, double, int,
			vector<int> *, const vector<vector<int> *> *);
	virtual ~ChromDetectorMaxima();
	const vector<ILocation*>* getRegionList() const;
	void printIndex(string);
	void printIndex(string, bool);

};

} /* namespace nonltr */
#endif /* CHROMDETECTORMAXIMA_H_ */
```
