## Supplementary Data 1 for "FASTCAR: Rapid alignment-free prediction of sequence alignment identity scores using self-supervised general linear models": ChromListMaker.cpp

```
/*
 * ChromListMaker.cpp
 *
 *  Created on: Mar 13, 2014
 *      Author: Hani Zakaira Girgis
 */

#include "ChromListMaker.h"

namespace nonltr {

ChromListMaker::ChromListMaker(string seqFileIn) {
	seqFile = seqFileIn;
	chromList = new vector<Chromosome *>();
}

ChromListMaker::~ChromListMaker() {
	Util::deleteInVector(chromList);
	delete chromList;
}


std::istream& safe_getline(std::istream& is, std::string& t)
{
	t.clear();
	std::istream::sentry se(is, true);
	std::streambuf* sb = is.rdbuf();
	for(;;) {
		int c = sb->sbumpc();
		switch (c) {
		case '\n':
			return is;
		case '\r':
			if (sb->sgetc() == '\n') {
				sb->sbumpc();
			}
			return is;
		case std::streambuf::traits_type::eof():
			if (t.empty()) {
				is.setstate(std::ios::eofbit);
			}
			return is;
		default:
			t += (char)c;
		}
	}
}

const vector<Chromosome *> * ChromListMaker::makeChromList() {
	ifstream in(seqFile.c_str());
	bool isFirst = true;
	Chromosome * chrom;

	while (in.good()) {
		string line;
		safe_getline(in, line);
		if (line[0] == '>') {
			if (!isFirst) {
				chrom->finalize();
				chromList->push_back(chrom);
			} else {
				isFirst = false;
			}

			chrom = new Chromosome();
			chrom->setHeader(line);
		} else if (line[0] == ' ' || line[0] == '\t') {
			bool all_spaces = true;
			for (auto c : line) {
				if (c != ' ' && c != '\t') {
					all_spaces = false;
				}
			}
			if (all_spaces) {
				continue;
			}
			std::ostringstream oss;
			oss << chrom->getHeader() << line;
			std::string new_header = oss.str();
			chrom->setHeader(new_header);
		} else {
			chrom->appendToSequence(line);
		}
	}
	chrom->finalize();
	chromList->push_back(chrom);
	in.close();

	return chromList;
}

const vector<Chromosome *> * ChromListMaker::makeChromOneDigitDnaList() {
	ifstream in(seqFile.c_str());
	bool isFirst = true;
	ChromosomeOneDigitDna * chrom;

	while (in.good()) {
		string line;
		safe_getline(in, line);
		if (line[0] == '>') {
			if (!isFirst) {
				chrom->finalize();
				chromList->push_back(chrom);
			} else {
				isFirst = false;
			}

			chrom = new ChromosomeOneDigitDna();
			chrom->setHeader(line);
		} else {
			chrom->appendToSequence(line);
		}
	}

	chrom->finalize();
	chromList->push_back(chrom);
	in.close();

	return chromList;
}

const vector<Chromosome *> * ChromListMaker::makeChromOneDigitProteinList() {
	ifstream in(seqFile.c_str());
	bool isFirst = true;
	ChromosomeOneDigitProtein * chrom;

	while (in.good()) {
		string line;
		safe_getline(in, line);
		if (line[0] == '>') {
			if (!isFirst) {
				chrom->finalize();
				chromList->push_back(chrom);
			} else {
				isFirst = false;
			}

			chrom = new ChromosomeOneDigitProtein();
			chrom->setHeader(line);
		} else {
			chrom->appendToSequence(line);
		}
	}

	chrom->finalize();
	chromList->push_back(chrom);
	in.close();

	return chromList;
}

}
/* namespace nonltr */
```
