## Supplementary Data 1 for "FASTCAR: Rapid alignment-free prediction of sequence alignment identity scores using self-supervised general linear models": ChromListMaker.h

```
/*
 * ChromListMaker.h
 *
 *   Created on: Mar 13, 2014
 *  Modified on: Oct 2, 2018
 *       Author: Hani Zakaria Girgis, PhD
 */

#ifndef CHROMLISTMAKER_H_
#define CHROMLISTMAKER_H_

#include <string>
#include <vector>

#include "Chromosome.h"
#include "ChromosomeOneDigitDna.h"
#include "ChromosomeOneDigitProtein.h"

#include "../utility/Util.h"

using namespace std;
using namespace utility;

namespace nonltr {

class ChromListMaker {
private:
	vector<Chromosome *> * chromList;
	string seqFile;

public:
	ChromListMaker(string);
	virtual ~ChromListMaker();
	const vector<Chromosome *> * makeChromList();
	const vector<Chromosome *> * makeChromOneDigitDnaList();
	const vector<Chromosome *> * makeChromOneDigitProteinList();

};

} /* namespace nonltr */
#endif /* CHROMLISTMAKER_H_ */
```
