## Supplementary Data 1 for "FASTCAR: Rapid alignment-free prediction of sequence alignment identity scores using self-supervised general linear models": Chromosome.cpp

```
/*
 * Chromosome.cpp
 *
 *  Created on: Mar 26, 2012
 *      Author: Hani Zakaria Girgis, PhD - NCBI/NLM/NIH
 */
#include "Chromosome.h"

Chromosome::Chromosome() {
	header = string("");
	base = string("");
	isHeaderReady = false;
	isBaseReady = false;
	isFinalized = false;
}

Chromosome::Chromosome(string fileName) {
	chromFile = fileName;
	readFasta();
	help(1000000, true);
}

Chromosome::Chromosome(string fileName, bool canMerge) {
	chromFile = fileName;
	readFasta();
	help(1000000, canMerge);
}

Chromosome::Chromosome(string fileName, int len) {
	chromFile = fileName;
	readFasta();
	help(len, true);
}

Chromosome::Chromosome(string fileName, int len, int maxLength) {
	chromFile = fileName;
	readFasta(maxLength);
	help(len, true);
}

Chromosome::Chromosome(string &seq, string &info) {
	header = info;
	base = seq;
	help(1000000, true);
}

Chromosome::Chromosome(string &seq, string &info, int len) {
	header = info;
	base = seq;
	help(len, true);
}

void Chromosome::setHeader(string& info) {
	if (isFinalized) {
		string msg("This chromosome has been finalized. ");
		msg.append("The header cannot be modified.");
		throw InvalidOperationException(msg);
	} else {
		header = info;
		isHeaderReady = true;
	}
}

/**
 * This method can waste memory if the sequence is large.
 * Consider using the method appendToSequence instead
 */
void Chromosome::setSequence(string& seq) {
	if (isFinalized) {
		string msg("This chromosome has been finalized. ");
		msg.append("The sequence cannot be modified.");
		throw InvalidOperationException(msg);
	} else {
		base = seq;
		isBaseReady = true;
	}
}

void Chromosome::appendToSequence(const string& line) {
	if (isFinalized) {
		string msg("This chromosome has been finalized. ");
		msg.append("The sequence cannot be modified.");
		throw InvalidOperationException(msg);
	} else {
		base.append(line);
		isBaseReady = true;
	}
}

void Chromosome::finalize() {
	if (isFinalized) {
		string msg("This chromosome has been already finalized. ");
		msg.append("Finalize can be only called once.");
		throw InvalidOperationException(msg);
	} else if (!(isHeaderReady && isBaseReady)) {
		string msg(
				"The header and the sequence must be set before calling finalize");
		throw InvalidOperationException(msg);
	} else {
		help(1000000, true);
		isFinalized = true;
	}
}

void Chromosome::help(int len, bool canMerge) {
	canClean = true;

	effectiveSize = 0;
	segLength = len;
	segment = new vector<vector<int> *>();
	//segment->reserve(100);

	toUpperCase();

	if(Util::isDna){
		baseCount = new vector<int>(4, 0);
		makeBaseCount();
	}

	removeAmbiguous();

	if (Util::isDna && (canMerge && base.size() > 20)) {
		mergeSegments();
	}

	makeSegmentList();
	calculateEffectiveSize();

}

Chromosome::~Chromosome() {
	base.clear();

	//cerr << "~Chromosome() 1" << endl;

	if (canClean) {
		while (!segment->empty()) {
			segment->back()->clear();
			delete segment->back();
			segment->pop_back();
		}
		segment->clear();

		// Util::deleteInVector(segment);
		delete segment;
		if(Util::isDna){
			baseCount->clear();
			delete baseCount;
		}
	}
	//cerr << "~Chromosome() 2" << endl;
}

void Chromosome::readFasta() {
	bool isFirst = true;
	header = string("");
	base = string("");

	ifstream in(chromFile.c_str());
	if (in.fail()) {
		string msg("Cannot open ");
		msg.append(chromFile);
		msg.append(". System code is: ");
		msg.append(Util::int2string(errno));
		throw InvalidInputException(msg);
	}

	while (in.good()) {
		string line;
		getline(in, line);
		if (line[0] == '>') {
			if (!isFirst) {
				string msg = "Chromosome file: ";
				msg = msg + chromFile;
				msg =
						msg
								+ " must have one sequence only. But it has more than one.";
				throw InvalidInputException(msg);
			} else {
				header = line;
				isFirst = false;
			}
		} else {
			base.append(line);
		}
	}
	in.close();
}

void Chromosome::readFasta(int maxLength) {
	bool isFirst = true;
	header = string("");
	base = string("");

	ifstream in(chromFile.c_str());
	if (in.fail()) {
		string msg("Cannot open ");
		msg.append(chromFile);
		msg.append(". System code is: ");
		msg.append(Util::int2string(errno));
		throw InvalidInputException(msg);
	}

	while (in.good() && base.size() < maxLength) {
		string line;
		getline(in, line);
		if (line[0] == '>') {
			if (!isFirst) {
				string msg = "Chromosome file: ";
				msg = msg + chromFile;
				msg =
						msg
								+ " must have one sequence only. But it has more than one.";
				throw InvalidInputException(msg);
			} else {
				header = line;
				isFirst = false;
			}
		} else {
			base.append(line);
		}
	}
	in.close();
}

/**
 * Convert alphabet to upper case if it has not been done before
 **/
void Chromosome::toUpperCase() {
	for (int i = 0; i < base.length(); i++) {
		base[i] = toupper(base[i]);
	}
}

/**
 * Segment coordinates are inclusive [s,e]
 **/
void Chromosome::removeAmbiguous() {
	// Store non-N index
	int start = -1;
	char uncertainChar = Util::isDna? 'N' : 'X';
	for (int i = 0; i < base.size(); i++) {
		if (base[i] != uncertainChar && start == -1) {
			start = i;
		} else if (base[i] == uncertainChar && start != -1) {
			vector<int> * v = new vector<int>();
			v->push_back(start);
			v->push_back(i - 1);
			segment->push_back(v);

			start = -1;
		} else if (i == base.size() - 1 && base[i] != uncertainChar && start != -1) {
			vector<int> * v = new vector<int>();
			v->push_back(start);
			v->push_back(i);

			segment->push_back(v);
			start = -1;
		}
	}

	// Test code
	// for(auto seg : *segment){
	// 	cerr << seg->at(0) << "-" << seg->at(1) << endl;
	// }
}

/**
 * Applied to DNA only--not proteins.
 * If the gap between two consecutive segments is less than 10 bp.
 * Segments that are shorter than 20 bp are not added.
 */
void Chromosome::mergeSegments() {

	// cout << "Segment size is " << segment->size() << endl;
	// cout << base << endl;

	if (segment->size() > 0) {
		vector<vector<int> *> * mSegment = new vector<vector<int> *>();
		int s = segment->at(0)->at(0);
		int e = segment->at(0)->at(1);

		for (int i = 1; i < segment->size(); i++) {
			int s1 = segment->at(i)->at(0);
			int e1 = segment->at(i)->at(1);

			/*
			 if(e1 - s1 + 1 <= 2000){
			 cout << "s1:" << s1 << " e1: " << e1 << endl;
			 }
			 */

			if (s1 - e < 10) {
				e = e1;
			} else {
				if (e - s + 1 >= 20) {
					vector<int> * seg = new vector<int>();
					seg->push_back(s);
					seg->push_back(e);
					mSegment->push_back(seg);
				}

				// Test start
				/*
				 if (e - s + 1 <= 100) {
				 cout << "Removing: " << base.substr(s, e - s + 1) << endl;
				 }
				 */
				// Test end
				s = s1;
				e = e1;
			}
		}

		// Handle the last index
		if (e - s + 1 >= 20) {
			vector<int> * seg = new vector<int>();
			seg->push_back(s);
			seg->push_back(e);
			mSegment->push_back(seg);
		}

		Util::deleteInVector(segment);
		segment->clear();
		delete segment;
		segment = mSegment;
	}
}

void Chromosome::makeSegmentList() {
	vector<vector<int> *> * segmentList = new vector<vector<int> *>();
	int segmentCount = segment->size();
	for (int oo = 0; oo < segmentCount; oo++) {
		int s = segment->at(oo)->at(0);
		int e = segment->at(oo)->at(1);

		if (e - s + 1 > segLength) {
			int fragNum = (int) (e - s + 1) / segLength;

			for (int h = 0; h < fragNum; h++) {
				int fragStart = s + (h * segLength);
				int fragEnd =
						(h == fragNum - 1) ? e : fragStart + segLength - 1;
				vector<int> * v = new vector<int>();
				v->push_back(fragStart);
				v->push_back(fragEnd);
				segmentList->push_back(v);
			}
		} else {
			vector<int> * v = new vector<int>();
			v->push_back(segment->at(oo)->at(0));
			v->push_back(segment->at(oo)->at(1));
			segmentList->push_back(v);
		}
	}

	Util::deleteInVector(segment);
	delete segment;
	segment = segmentList;
}

const string* Chromosome::getBase() {
	return &base;
}

string& Chromosome::getBaseRef() {
	return base;
}

string& Chromosome::getHeaderRef() {
	return header;
}

const vector<vector<int> *> * Chromosome::getSegment() {
	return segment;
}

void Chromosome::printSegmentList() {
	int l = segment->size();
	cout << "Segment list size = " << l << endl;
	for (int i = 0; i < l; i++) {
		cout << segment->at(i)->at(0) << "\t";
		cout << segment->at(i)->at(1) << endl;
	}
}

string Chromosome::getHeader() {
	return header;
}

int Chromosome::size() {
	return base.size();
}

void Chromosome::calculateEffectiveSize() {
	int segmentCount = segment->size();
	for (int oo = 0; oo < segmentCount; oo++) {
		int s = segment->at(oo)->at(0);
		int e = segment->at(oo)->at(1);
		effectiveSize += (e - s + 1);
	}
}

int Chromosome::getEffectiveSize() {
	return effectiveSize;
}

int Chromosome::getGcContent() {
	if(!Util::isDna){
		cerr << "Calculating GC content on a protein sequence is not allowed." << endl;
		throw std::exception();
	}

	int gc = 0;
	int size = base.size();
	for (int i = 0; i < size; i++) {
		char n = base.at(i);
		if (n == 'C' || n == 'G') {
			gc++;
		}
	}
	return gc;
}

void Chromosome::makeBaseCount() {
	if(!Util::isDna){
		cerr << "Counting nucleotides in a protein sequence is not allowed." << endl;
		throw std::exception();
	}

	int size = base.size();
	for (int i = 0; i < size; i++) {
		switch (base.at(i)) {
		case 'A':
			baseCount->at(0)++;
			break;
;		case 'C':
			baseCount->at(1)++;
			break;
		case 'G':
			baseCount->at(2)++;
			break;
		case 'T':
			baseCount->at(3)++;
			break;
		}
	}
}

vector<int> * Chromosome::getBaseCount() {
	if(!Util::isDna){
		cerr << "Counting nucleotides in a protein sequence is not allowed." << endl;
		throw std::exception();
	}

	return baseCount;
}
```
