## Supplementary Data 1 for "FASTCAR: Rapid alignment-free prediction of sequence alignment identity scores using self-supervised general linear models": Chromosome.h

```
/*
 * Chromosome.h
 *
 *  Created on: Mar 26, 2012
 *      Author: Hani Zakaria Girgis, PhD - NCBI/NLM/NIH
 */
#ifndef CHROMOSOME_H_
#define CHROMOSOME_H_

#include <string>
#include <fstream>
#include <vector>
#include <iostream>
#include <map>

#include "IChromosome.h"
#include "../exception/InvalidOperationException.h"
#include "../exception/InvalidInputException.h"
#include "../utility/Util.h"

using namespace std;
using namespace nonltr;
using namespace utility;
using namespace exception;

namespace nonltr {
class Chromosome: public IChromosome {
public:
	Chromosome();
	Chromosome(string);
	Chromosome(string, bool);
	Chromosome(string, int);
	Chromosome(string, int, int);
	Chromosome(string &, string&);
	Chromosome(string &, string&, int);

	int getGcContent();

	virtual ~Chromosome();

	virtual string& getBaseRef();
	virtual string& getHeaderRef();

	virtual const string* getBase();
	virtual const vector<vector<int> *> * getSegment();
	virtual void printSegmentList();
	virtual string getHeader();
	virtual int size();
	virtual int getEffectiveSize();
	virtual void setHeader(string&);
	virtual void setSequence(string&);
	virtual void appendToSequence(const string&);
	virtual void finalize();
	virtual vector<int> * getBaseCount();

protected:
	string chromFile;
	string header;
	string base;
	int effectiveSize;
	int segLength;

	vector<vector<int> *> * segment;
	void readFasta();
	void readFasta(int);
	void toUpperCase();
	void removeAmbiguous();
	void mergeSegments();
	virtual void help(int, bool);
	void makeSegmentList();
	void calculateEffectiveSize();

private:
	bool isHeaderReady;
	bool isBaseReady;
	bool isFinalized;
	bool canClean = false;

	void reverseSegments();
	void makeBaseCount();
	vector<int> * baseCount;
};
}

#endif /* CHROMOSOME_H_ */
```
