## Supplementary Data 1 for "FASTCAR: Rapid alignment-free prediction of sequence alignment identity scores using self-supervised general linear models": ChromosomeOneDigit.cpp

```
/*
 * ChromosomeOneDigit.cpp
 *
 *  Created on: Jul 31, 2012
 *      Author: Hani Zakaria Girgis, PhD at the NCB1/NLM/NIH
 */
#include <iostream>
#include <map>

#include "Chromosome.h"
#include "ChromosomeOneDigit.h"
#include "../exception/InvalidInputException.h"

using namespace exception;

namespace nonltr {

ChromosomeOneDigit::ChromosomeOneDigit() :
		Chromosome() {
	//cout << "The no args constructor is called" << endl;
}

ChromosomeOneDigit::ChromosomeOneDigit(string fileName) :
		Chromosome(fileName) {
	help();
}

ChromosomeOneDigit::ChromosomeOneDigit(string fileName, int segmentLength,
		int maxLength) :
		Chromosome(fileName, segmentLength, maxLength) {
	help();
}

ChromosomeOneDigit::ChromosomeOneDigit(string& seq, string& info) :
		Chromosome(seq, info) {
	//cout << "Two string constructor is called" << endl;
	help();
}

ChromosomeOneDigit::ChromosomeOneDigit(string& seq, string& info, int length) :
		Chromosome(seq, info, length) {
	//cout << "Two string constructor is called" << endl;
	help();
}

void ChromosomeOneDigit::finalize() {
	Chromosome::finalize();
	help();
}

void ChromosomeOneDigit::help() {
	// Can delete the codes
	canClean = true;

	// Make map
	codes = new map<char, char>();

	// Build codes
	buildCodes();
	// Modify the sequence in the super class
	encode();
}

ChromosomeOneDigit::~ChromosomeOneDigit() {
	if (canClean) {
		codes->clear();
		delete codes;
	} 
}

/**
 * This method converts nucleotides in the segments to single digit codes
 */
void ChromosomeOneDigit::encode() {

	for (int s = 0; s < segment->size(); s++) {
		int segStart = segment->at(s)->at(0);
		int segEnd = segment->at(s)->at(1);
		for (int i = segStart; i <= segEnd; i++) {

			if (codes->count(base[i]) > 0) {
				base[i] = codes->at(base[i]);
			} else {
				string msg = "Invalid nucleotide: ";
				msg.append(1, base[i]);
				throw InvalidInputException(msg);
			}
		}
	}

	// Digitize skipped segments
	char uncertainChar = Util::isDna? 'N' : 'X';
	int segNum = segment->size();
	if (segNum > 0) {
		// The first interval - before the first segment
		int segStart = 0;
		int segEnd = segment->at(0)->at(0) - 1;

		for (int s = 0; s <= segNum; s++) {
			for (int i = segStart; i <= segEnd; i++) {
				char c = base[i];

				if (c != uncertainChar) {
					if (codes->count(c) > 0) {
						base[i] = codes->at(c);
					} else {
						string msg = "ChromosomeOneDigit::encode() found invalid letter: ";
						msg.append(1, c);
						throw InvalidInputException(msg);
					}
				}
			}

			// The regular intervals between two segments
			if (s < segNum - 1) {
				segStart = segment->at(s)->at(1) + 1;
				segEnd = segment->at(s + 1)->at(0) - 1;
			}
			// The last interval - after the last segment
			else if (s == segNum - 1) {
				segStart = segment->at(s)->at(1) + 1;
				segEnd = base.size() - 1;
			}
		}
	}
}

}
```
