## Supplementary Data 1 for "FASTCAR: Rapid alignment-free prediction of sequence alignment identity scores using self-supervised general linear models": ChromosomeOneDigit.h

```
/*
 * ChromosomeOneDigit.h
 *
 *  Created on: Jul 31, 2012
 *      Author: Hani Zakaria Girgis, PhD - NCBI/NLM/NIH
 */

#ifndef CHROMOSOMEONEDIGIT_H_
#define CHROMOSOMEONEDIGIT_H_

#include <map>
#include "Chromosome.h"

namespace nonltr {
class ChromosomeOneDigit: public Chromosome {

private:
	void encode();
	void help();


protected:
	bool canClean = false;
	map<char, char> * codes;
	virtual void buildCodes() = 0;


public:
	/* Methods */
	ChromosomeOneDigit();
	ChromosomeOneDigit(string);
	ChromosomeOneDigit(string, int, int);
	ChromosomeOneDigit(string&, string&);
	ChromosomeOneDigit(string&, string&, int);
	virtual ~ChromosomeOneDigit();
	virtual void finalize();


};
}

#endif /* CHROMOSOMEONEDIGIT_H_ */
```
