## Supplementary Data 1 for "FASTCAR: Rapid alignment-free prediction of sequence alignment identity scores using self-supervised general linear models": ChromosomeOneDigitDna.cpp

```
#include "ChromosomeOneDigitDna.h"

namespace nonltr{

ChromosomeOneDigitDna::ChromosomeOneDigitDna() : ChromosomeOneDigit() {}

ChromosomeOneDigitDna::ChromosomeOneDigitDna(string fileName) : 
	ChromosomeOneDigit(fileName){

}

ChromosomeOneDigitDna::ChromosomeOneDigitDna(string fileName, int segmentLength, int maxLength) : 
	ChromosomeOneDigit(fileName, segmentLength, maxLength) {

}

ChromosomeOneDigitDna::ChromosomeOneDigitDna(string& seq, string& info) : 
	ChromosomeOneDigit(seq, info){

}

ChromosomeOneDigitDna::ChromosomeOneDigitDna(string& seq, string& info, int length) : 
	ChromosomeOneDigit(seq, info, length) {
}

ChromosomeOneDigitDna::~ChromosomeOneDigitDna(){

}

/**
 * A	A
 * T	T
 * G	G
 * C	C
 * R	G or A
 * Y	T or C
 * M	A or C
 * K	G or T
 * S	G or C
 * W	A or T
 * H	A or C or T
 * B	G or T or C
 * V	G or C or A
 * D	G or T or A
 * N	G or T or A or C
 */
void ChromosomeOneDigitDna::buildCodes() {
	// Certain nucleotides
	codes->insert(map<char, char>::value_type('A', (char) 0));
	codes->insert(map<char, char>::value_type('C', (char) 1));
	codes->insert(map<char, char>::value_type('G', (char) 2));
	codes->insert(map<char, char>::value_type('T', (char) 3));

	// Uncertain nucleotides
	codes->insert(map<char, char>::value_type('R', codes->at('G')));
	codes->insert(map<char, char>::value_type('Y', codes->at('C')));
	codes->insert(map<char, char>::value_type('M', codes->at('A')));
	codes->insert(map<char, char>::value_type('K', codes->at('T')));
	codes->insert(map<char, char>::value_type('S', codes->at('G')));
	codes->insert(map<char, char>::value_type('W', codes->at('T')));
	codes->insert(map<char, char>::value_type('H', codes->at('C')));
	codes->insert(map<char, char>::value_type('B', codes->at('T')));
	codes->insert(map<char, char>::value_type('V', codes->at('A')));
	codes->insert(map<char, char>::value_type('D', codes->at('T')));
	codes->insert(map<char, char>::value_type('N', codes->at('C')));
	codes->insert(map<char, char>::value_type('X', codes->at('G')));
}

/**
 * Cannot be called on already finalized object.
 */
void ChromosomeOneDigitDna::makeR() {
	//cout << "Making reverse ..." << endl;
	makeReverse();
	reverseSegments();
}

/**
 * Cannot be called on already finalized object.
 */
void ChromosomeOneDigitDna::makeRC() {
	//cout << "Making reverse complement ..." << endl;
	makeComplement();
	makeReverse();
	reverseSegments();
}

void ChromosomeOneDigitDna::makeComplement() {
	map<char, char> complement;

	// Certain nucleotides
	complement.insert(map<char, char>::value_type((char) 0, (char) 3));
	complement.insert(map<char, char>::value_type((char) 1, (char) 2));
	complement.insert(map<char, char>::value_type((char) 2, (char) 1));
	complement.insert(map<char, char>::value_type((char) 3, (char) 0));

	// Unknown nucleotide
	complement.insert(map<char, char>::value_type('N', 'N'));
	// complement.insert(map<char, char>::value_type((char) 4, (char) 4));

	// Convert a sequence to its complement
	int seqLen = base.size();
	for (int i = 0; i < seqLen; i++) {
		if (complement.count(base[i]) > 0) {
			base[i] = complement.at(base[i]);
		} else {
			cerr << "Error: The digit " << (char) base[i];
			cerr << " does not represent a base." << endl;
			exit(2);
		}
	}
}

void ChromosomeOneDigitDna::makeReverse() {
	int last = base.size() - 1;

	// Last index to be switched
	int middle = base.size() / 2;

	for (int i = 0; i < middle; i++) {
		char temp = base[last - i];
		base[last - i] = base[i];
		base[i] = temp;
	}
}

void ChromosomeOneDigitDna::reverseSegments() {
	int segNum = segment->size();
	int lastBase = size() - 1;

	// Calculate the coordinate on the main strand
	for (int i = 0; i < segNum; i++) {
		vector<int> * seg = segment->at(i);

		int s = lastBase - seg->at(1);
		int e = lastBase - seg->at(0);
		seg->clear();
		seg->push_back(s);
		seg->push_back(e);
	}

	// Reverse the regions within the list
	int lastRegion = segNum - 1;
	int middle = segNum / 2;
	for (int i = 0; i < middle; i++) {
		vector<int> * temp = segment->at(lastRegion - i);
		(*segment)[lastRegion - i] = segment->at(i);
		(*segment)[i] = temp;
	}
}


}
```
