## Supplementary Data 1 for "FASTCAR: Rapid alignment-free prediction of sequence alignment identity scores using self-supervised general linear models": ChromosomeOneDigitDna.h

```
/*
 * ChromosomeOneDigitDna.h
 * Created on: September 28, 2018
 *     Author: Hani Z. Girgis, PhD
 */

 #ifndef HROMOSOMEONEDIGITDNA_H_
 #define HROMOSOMEONEDIGITDNA_H_

#include "ChromosomeOneDigit.h"

namespace nonltr{
	class ChromosomeOneDigitDna: public ChromosomeOneDigit{
		private:
			void makeReverse();
			void makeComplement();
			void reverseSegments();

		protected:
			virtual void buildCodes();

		public:
			ChromosomeOneDigitDna();
			ChromosomeOneDigitDna(string);
			ChromosomeOneDigitDna(string, int, int);
			ChromosomeOneDigitDna(string&, string&);
			ChromosomeOneDigitDna(string&, string&, int);
			virtual ~ChromosomeOneDigitDna();

			void makeR();
			void makeRC();
	};
}

#endif
```
