## Supplementary Data 1 for "FASTCAR: Rapid alignment-free prediction of sequence alignment identity scores using self-supervised general linear models": ChromosomeOneDigitProtein.cpp

```
#include "ChromosomeOneDigitProtein.h"

namespace nonltr{

ChromosomeOneDigitProtein::ChromosomeOneDigitProtein() : 
	ChromosomeOneDigit() {

}

ChromosomeOneDigitProtein::ChromosomeOneDigitProtein(string fileName) : 
	ChromosomeOneDigit(fileName){

}

ChromosomeOneDigitProtein::ChromosomeOneDigitProtein(string fileName, int segmentLength, int maxLength) : 
	ChromosomeOneDigit(fileName, segmentLength, maxLength) {

}

ChromosomeOneDigitProtein::ChromosomeOneDigitProtein(string& seq, string& info) : 
	ChromosomeOneDigit(seq, info){

}

ChromosomeOneDigitProtein::ChromosomeOneDigitProtein(string& seq, string& info, int length) : 
	ChromosomeOneDigit(seq, info, length) {
}

ChromosomeOneDigitProtein::~ChromosomeOneDigitProtein(){

}

void ChromosomeOneDigitProtein::buildCodes() {
	// https://en.wikipedia.org/wiki/Proteinogenic_amino_acid
	codes->insert(map<char, char>::value_type('A', (char) 0));
	codes->insert(map<char, char>::value_type('C', (char) 1));
	codes->insert(map<char, char>::value_type('D', (char) 2));
	codes->insert(map<char, char>::value_type('E', (char) 3));
	codes->insert(map<char, char>::value_type('F', (char) 4));
	codes->insert(map<char, char>::value_type('G', (char) 5));
	codes->insert(map<char, char>::value_type('H', (char) 6));
	codes->insert(map<char, char>::value_type('I', (char) 7));
	codes->insert(map<char, char>::value_type('K', (char) 8));
	codes->insert(map<char, char>::value_type('L', (char) 9));
	codes->insert(map<char, char>::value_type('M', (char) 10));
	codes->insert(map<char, char>::value_type('N', (char) 11));
	codes->insert(map<char, char>::value_type('O', (char) 12));
	codes->insert(map<char, char>::value_type('P', (char) 13));
	codes->insert(map<char, char>::value_type('Q', (char) 14));
	codes->insert(map<char, char>::value_type('R', (char) 15));
	codes->insert(map<char, char>::value_type('S', (char) 16));
	codes->insert(map<char, char>::value_type('T', (char) 17));
	codes->insert(map<char, char>::value_type('U', (char) 18));
	codes->insert(map<char, char>::value_type('V', (char) 19));
	codes->insert(map<char, char>::value_type('W', (char) 20));
	codes->insert(map<char, char>::value_type('Y', (char) 21));

	// Uncertain uncleotides
	codes->insert(map<char, char>::value_type('B', codes->at('D')));
	codes->insert(map<char, char>::value_type('Z', codes->at('E')));
	codes->insert(map<char, char>::value_type('J', codes->at('L')));
}

}// End namespace
```
