## Supplementary Data 1 for "FASTCAR: Rapid alignment-free prediction of sequence alignment identity scores using self-supervised general linear models": ChromosomeOneDigitProtein.h

```
/*
 * ChromosomeOneDigitProtein.h
 * Created on: October 2, 2018
 *     Author: Hani Z. Girgis, PhD
 */

 #ifndef HROMOSOMEONEDIGITPROTEIN_H_
 #define HROMOSOMEONEDIGITPROTEIN_H_

#include "ChromosomeOneDigit.h"

namespace nonltr{
	class ChromosomeOneDigitProtein: public ChromosomeOneDigit{

		protected:
			virtual void buildCodes();

		public:
			ChromosomeOneDigitProtein();
			ChromosomeOneDigitProtein(string);
			ChromosomeOneDigitProtein(string, int, int);
			ChromosomeOneDigitProtein(string&, string&);
			ChromosomeOneDigitProtein(string&, string&, int);
			virtual ~ChromosomeOneDigitProtein();
	};
}

#endif
```
