## Supplementary Data 1 for "FASTCAR: Rapid alignment-free prediction of sequence alignment identity scores using self-supervised general linear models": ChromosomeRandom.cpp

```
/*
 * ChromosomeRandom.cpp
 *
 *  Created on: Feb 4, 2013
 *      Author: Hani Zakaria Girgis, PhD
 *
 */

#include <string>
#include <iostream>
#include <fstream>
#include <vector>
#include <cstdlib>
#include <ctime>

#include "ChromosomeRandom.h"
#include "../exception/InvalidInputException.h"
#include "../exception/InvalidStateException.h"
#include "../utility/Util.h"

using namespace std;
using namespace exception;
using namespace utility;

namespace nonltr {

ChromosomeRandom::ChromosomeRandom(int nIn, IChromosome* oChromIn,
		char unreadIn, vector<char>* alphaIn) {
	// Check the order
	if (nIn < 0) {
		string msg("The Markov order must be non-negative. ");
		msg.append("The order received is: ");
		msg.append(Util::int2string(nIn));
		msg.append(".");
		throw InvalidInputException(msg);
	}

	// n here is the length of the word, i.e. the order + 1
	n = nIn + 1;
	oChrom = oChromIn;
	unread = unreadIn;
	alpha = alphaIn;

	// Initialize the random sequence
	int size = oChrom->getBase()->size();
	rBase = new string(size, unread);

	// Initialize key list
	keyList = new vector<string>();

	// Initialize the table
	table = new map<string, double>();

	// Handle unusual characters in the first word of a segment
	// Make map
	codes = new map<char, char>();
	codes->insert(map<char, char>::value_type('A', 'A'));
	codes->insert(map<char, char>::value_type('C', 'C'));
	codes->insert(map<char, char>::value_type('G', 'G'));
	codes->insert(map<char, char>::value_type('T', 'T'));
	codes->insert(map<char, char>::value_type('R', 'G'));
	codes->insert(map<char, char>::value_type('Y', 'C'));
	codes->insert(map<char, char>::value_type('M', 'A'));
	codes->insert(map<char, char>::value_type('K', 'T'));
	codes->insert(map<char, char>::value_type('S', 'G'));
	codes->insert(map<char, char>::value_type('W', 'T'));
	codes->insert(map<char, char>::value_type('H', 'C'));
	codes->insert(map<char, char>::value_type('B', 'T'));
	codes->insert(map<char, char>::value_type('V', 'A'));
	codes->insert(map<char, char>::value_type('D', 'T'));
	codes->insert(map<char, char>::value_type('N', 'C'));
	codes->insert(map<char, char>::value_type('X', 'G'));

	// Start operations
	cout << "\tFilling key list ..." << endl;
	fillKeyList();

	cout << "\tInitializing table ..." << endl;
	initializeTable();

	cout << "\tCounting words ..." << endl;
	countWords();

	cout << "\tCalculating probabilities ..." << endl;
	convertToProbabilities();

	//cout << "\tPrinting the table ..." << endl;
	//printTable();

	cout << "\tGenerating the random sequence ..." << endl;
	generateRandomSequence();
}

ChromosomeRandom::~ChromosomeRandom() {
	codes->clear();
	delete codes;

	keyList->clear();
	delete keyList;

	table->clear();
	delete table;

	delete rBase;
}

void ChromosomeRandom::fillKeyList() {
	// Collect keys
	int alphaCount = alpha->size();

	// Order 0

	for (int h = 0; h < alphaCount; h++) {
		string s("");
		s.append(1, alpha->at(h));
		keyList->push_back(s);
	}

	// Order 1 and higher
	for (int g = 1; g < n; g++) {
		vector<string> o;
		int keyListSize = keyList->size();
		for (int i = 0; i < keyListSize; i++) {
			for (int j = 0; j < alphaCount; j++) {
				string s(keyList->at(i));
				s.append(1, alpha->at(j));
				o.push_back(s);
			}
		}
		keyList->clear();
		(*keyList) = o;
	}
}

void ChromosomeRandom::initializeTable() {
	int keyListSize = keyList->size();
	for (int i = 0; i < keyListSize; i++) {
		table->insert(valType(keyList->at(i), 1));
	}
}

void ChromosomeRandom::countWords() {
	// Get the original sequence
	const string* oBase = oChrom->getBase();

	// Count words
	const vector<vector<int> *> * segmentList = oChrom->getSegment();
	int segmentCount = segmentList->size();
	for (int i = 0; i < segmentCount; i++) {
		int s = segmentList->at(i)->at(0);
		int e = segmentList->at(i)->at(1);
		if (e - s + 1 >= n) {

			int limit = e - n + 1;

			for (int h = s; h <= limit; h++) {
				// Check if the current base is a standard one.
				// Words including non-standard bases are not counted.

				char c = oBase->at(h);

				int alphaCount = alpha->size();
				bool isStandard = false;
				for (int a = 0; a < alphaCount; a++) {
					if (alpha->at(a) == c) {
						isStandard = true;
						break;
					}
				}

				// Increment the count
				if (isStandard) {
					string word = oBase->substr(h, n);
					if (table->count(word) > 0) {
						(*table)[word] = table->at(word) + 1;
					} else {
						cout << "\t\tIgnoring " << word << endl;
					}
				}
			}
		}
	}
}

void ChromosomeRandom::convertToProbabilities() {
	int alphaCount = alpha->size();
	int keyListSize = keyList->size();
	for (int i = 0; i < keyListSize; i += alphaCount) {
		double sum = 0;
		for (int j = 0; j < alphaCount; j++) {
			string key = keyList->at(i + j);
			sum += table->at(key);
		}
		for (int j = 0; j < alphaCount; j++) {
			string key = keyList->at(i + j);
			(*table)[key] = ((double) table->at(key)) / sum;
		}
	}
}

void ChromosomeRandom::generateRandomSequence() {
	// Get the original sequence
	const string* oBase = oChrom->getBase();

	// Alphabet count
	int alphaCount = alpha->size();

	// Get the original segments
	const vector<vector<int> *> * segmentList = oChrom->getSegment();
	int segmentCount = segmentList->size();

	// Generate random segments
	for (int i = 0; i < segmentCount; i++) {
		int s = segmentList->at(i)->at(0);
		int e = segmentList->at(i)->at(1);

		if (e - s + 1 > n) {
			//string order = oBase->substr(s, n - 1);
			string order("");
			// The first order is based on the original sequence.
			for (int w = s; w < s + n - 1; w++) {
				(*rBase)[w] = codes->at(oBase->at(w));
				order.append(1, codes->at(oBase->at(w)));
			}

			for (int h = s + n - 1; h <= e; h++) {
				// Subsequent orders are based on the random sequence.
				order = rBase->substr(h - n + 1, n - 1);
				vector<vector<int> > lottery;
				int chanceSoFar = 0;
				for (int k = 0; k < alphaCount; k++) {
					string temp = order;
					temp.append(1, alpha->at(k));
					if (table->count(temp) > 0) {
						int periodStart = chanceSoFar;
						int periodEnd = periodStart + (100 * table->at(temp));
						chanceSoFar = periodEnd + 1;
						vector<int> entry;
						entry.push_back(alpha->at(k));
						entry.push_back(periodStart);
						entry.push_back(periodEnd);
						lottery.push_back(entry);
					} else {
						string msg("This word must exist in the table: ");
						msg.append(temp);
						msg.append(".");
						throw InvalidStateException(msg);
					}
				}

				if (lottery.size() > 0) {
					int randInt = rand() % chanceSoFar;

					for (int tt = 0; tt < alphaCount; tt++) {
						vector<int> entry = lottery.at(tt);
						if (randInt >= entry.at(1) && randInt <= entry.at(2)) {
							(*rBase)[h] = entry.at(0);
							break;
						}
					}
					lottery.clear();
				} else {
					string msg("The lottery vector cannot be empty.");
					throw InvalidStateException(msg);
				}
			}
		}
	}

	// Make sure that the generated sequence has the same length as the original sequence
	if (oBase->size() != rBase->size()) {
		cerr << "The original sequence and the random sequence ";
		cerr << "do not have the same size." << endl;
		cerr << "Original sequence size is: " << oBase->size() << endl;
		cerr << "Generated sequence size is: " << rBase->size() << endl;
	}
}

void ChromosomeRandom::printTable() {
	map<string, double>::iterator iterStart = table->begin();
	map<string, double>::iterator iterEnd = table->end();
	while (iterStart != iterEnd) {
		cout << (*iterStart).first << " -> " << (*iterStart).second << endl;
		iterStart++;
	}
}

/**
 * Returns the segments of the original chromosome
 */
const vector<vector<int> *> * ChromosomeRandom::getSegment() {
	return oChrom->getSegment();
}

/**
 * Returns the random sequence
 */
const string* ChromosomeRandom::getBase() {
	return rBase;
}

/**
 * Returns the header indicating the order of the Markov chain
 */
string ChromosomeRandom::getHeader() {
	string header = oChrom->getHeader();
//header.append(" - Random based on ");
//header.append(Util::int2string(n - 1));
//header.append("-order Markov chain.");
	return header;
}

void ChromosomeRandom::printEffectiveSequence(string outputFile) {
	int totalSize = rBase->size();
	string * effectiveRBase = new string("");
	for (int i = 0; i < totalSize; i++) {
		char b = rBase->at(i);
		if (b != unread) {
			effectiveRBase->append(1, b);
		}
	}

	// Make sure that the effective sequence is shorter than the original
	// length
	if (effectiveRBase->size() > totalSize) {
		cerr << "The effective length must be <= the original length." << endl;
		cerr << "Generated sequence size is: " << totalSize << endl;
		cerr << "The effective size is: " << effectiveRBase->size() << endl;

	}

	printSequence(outputFile, effectiveRBase);

	delete effectiveRBase;
}

void ChromosomeRandom::printSequence(string outputFile) {
	printSequence(outputFile, rBase);
}

void ChromosomeRandom::printSequence(string outputFile, string * baseToPrint) {
	cout << "Printing chromosome to file ..." << endl;
	ofstream outSequence;
	outSequence.open(outputFile.c_str(), ios::out);

	int step = 50;

	outSequence << getHeader() << endl;
	int len = baseToPrint->size();

	for (int i = 0; i < len; i = i + step) {
		int e = (i + step - 1 > len - 1) ? len - 1 : i + step - 1;
		for (int k = i; k <= e; k++) {
			outSequence << baseToPrint->at(k);
		}
		outSequence << endl;
	}
	outSequence << endl;

	outSequence.close();
}

} /* namespace nonltr */
```
