## Supplementary Data 1 for "FASTCAR: Rapid alignment-free prediction of sequence alignment identity scores using self-supervised general linear models": ChromosomeRandom.h

```
/*
 * ChromosomeRandom.h
 *
 *  Created on: Feb 4, 2013
 *      Author: Hani Zakaria Girgis, PhD
 */

#ifndef CHROMOSOMERANDOM_H_
#define CHROMOSOMERANDOM_H_

#include <map>

#include "IChromosome.h"

namespace nonltr {

class ChromosomeRandom: public nonltr::IChromosome {
	// Key-value pair type.
	typedef map<string, double>::value_type valType;

private:
	int n;
	char unread;
	IChromosome * oChrom;
	vector<char> * alpha;
	map<string, double> * table;
	string * rBase;
	vector<string> * keyList;
	map<char, char> * codes;

	void fillKeyList();
	void initializeTable();
	void countWords();
	void convertToProbabilities();
	void printTable();
	void generateRandomSequence();

public:
	ChromosomeRandom(int, IChromosome*, char, vector<char>*);
	virtual ~ChromosomeRandom();

	virtual const string* getBase();
	virtual const vector<vector<int> *> * getSegment();
	virtual string getHeader();
	virtual void printSequence(string);
	void printSequence(string, string *);
	void printEffectiveSequence(string);
};

} /* namespace nonltr */
#endif /* CHROMOSOMERANDOM_H_ */
```
