## Supplementary Data 1 for "FASTCAR: Rapid alignment-free prediction of sequence alignment identity scores using self-supervised general linear models": DetectorMaxima.cpp

```
/*
 * DetectorMaxima.cpp
 *
 *  Created on: May 31, 2013
 *      Author: Hani Zakaria Girgis, PhD
 */

#include "DetectorMaxima.h"
#include "../utility/Util.h"
#include "../utility/Location.h"
#include "../exception/InvalidStateException.h"

#include <cmath>
// Delete start
#include <iostream>
using namespace std;
// Delete end

using namespace exception;

namespace nonltr {

DetectorMaxima::DetectorMaxima(int segStartIn, int segEndIn, double sIn,
		double wIn, double mIn, double tIn, double pIn, int eIn,
		vector<int> * oScoresIn) {

	// ToDo: make sure that segStart and segEnd are within the input scores.
	segStart = segStartIn;
	segEnd = segEndIn;
	s = sIn;
	w = wIn;
	m = mIn;
	t = tIn;
	p = pIn;
	e = eIn;
	oScores = oScoresIn;

	halfS = s;
	//s / 2;

	mask = new vector<double>();
	// Complete
	scores = new vector<double>();

	// Trimmed on both sides
	first = new vector<double>();

	// Trimmed on both sides
	second = new vector<double>();

	// Coordinates according to the complete sequence
	maxima = new vector<int>();

	// Coordinates according to the complete sequence
	// allMaxima = new vector<vector<double> *>();

	// Coordinates according to the complete sequence
	separatorList = new vector<ILocation *>();

	// Coordinates according to the complete sequence
	regionList = new vector<ILocation *>();

	makeMask();

	smooth();

	deriveFirst();

	deriveSecond();

	// Free memory start
	mask->clear();
	delete mask;
	scores->clear();
	delete scores;
	// Free memory end

	findMaxima();

	// Free memory start
	first->clear();
	delete first;
	second->clear();
	delete second;
	// Free memory end

	findSeparators();

	findRegions();

	// Free memory start
	maxima->clear();
	delete maxima;
	Util::deleteInVector(separatorList);
	separatorList->clear();
	delete separatorList;
	// Free memory end

	extendRegions();
}

/*
 const vector<vector<double> *>* DetectorMaxima::getAllMaxima() const {
 return allMaxima;
 }
 */

const vector<double>* DetectorMaxima::getFirst() const {
	return first;
}

const vector<double>* DetectorMaxima::getSecond() const {
	return second;
}

const vector<ILocation*> * DetectorMaxima::getRegionList() const {
	return regionList;
}

DetectorMaxima::~DetectorMaxima() {
	/*
	 Util::deleteInVector (allMaxima);
	 allMaxima->clear();
	 delete allMaxima;
	 */

	Util::deleteInVector(regionList);
	regionList->clear();
	delete regionList;
}

void DetectorMaxima::makeMask() {
	const double PI = 3.14159265358979323846;
	double sigma = (double) s / 3.5;
	const double PART_1 = 1 / sqrt(2 * PI * pow(sigma, 2));

	int l = 2 * s + 1;
	for (int i = 0; i < l; i++) {
		double g = PART_1 * exp(-1 * pow(i - s, 2) / (2 * pow(sigma, 2)));
		mask->push_back(g);
	}

	// For testing only
	/*
	for (int i = 0; i < l; i++) {
		cout << i << "\t" << mask->at(i) << endl;
	}
	cout << endl;
	cout << endl;
	*/
	// End testing
}

void DetectorMaxima::smooth() {
	for (int i = segStart; i <= segEnd; i++) {
		int winS = i - s;
		int maskS = 0;
		if (winS < segStart) {
			maskS = -1 * (winS - segStart);
			winS = segStart;
		}

		int winE = (i + s > segEnd) ? segEnd : i + s;
		// int winL = winE - winS + 1;

		double sum = 0.0;
		double maskSum = 0.0;

		int j = winS;
		int h = maskS;

		while (j <= winE) {
			double weight = mask->at(h);
			sum += oScores->at(j) * weight;
			maskSum += weight;

			j++;
			h++;
		}

		if (maskSum <= 0.0) {
			string msg("The sum of the weights in the mask must be > 0");
			throw InvalidStateException(msg);
		}

		scores->push_back(sum / maskSum);
		// scores->push_back(sum / winL);
	}

	// Testing - start
	/*
	cout << "The smoothed scores ... " << endl;
	for (int k = 0; k < scores->size(); k++) {
		if (k % 25 == 0) {
			cout << endl;
		}
		cout << scores->at(k) << " ";
	}
	cout << endl;
	cout << endl;
	*/
	// Testing - end
}

void DetectorMaxima::deriveFirst() {
	double l = 0.0;
	double r = 0.0;

	for (int i = 0; i < w; i++) {
		l += scores->at(i);
	}

	for (int i = w + 1; i <= 2 * w; i++) {
		r += scores->at(i);
	}

	first->push_back(round(-1 * l + r));

	for (int i = w + 1; i < scores->size() - w; i++) {
		l -= scores->at(i - w - 1);
		l += scores->at(i - 1);
		r -= scores->at(i);
		r += scores->at(i + w);
		first->push_back(round(-1 * l + r));
	}

	// For testing only
	/*
	 for (int i = 0; i < first->size(); i++) {
	 cout << first->at(i) << " ";
	 }
	 cout << endl;
	 */
}

void DetectorMaxima::deriveSecond() {
	double l = 0.0;
	double r = 0.0;
	double d = 2 * w;

	for (int i = 0; i < w; i++) {
		l += scores->at(i);
	}

	for (int i = w + 1; i <= 2 * w; i++) {
		r += scores->at(i);
	}

	second->push_back(round(l + r - d * scores->at(w)));

	for (int i = w + 1; i < scores->size() - w; i++) {
		l -= scores->at(i - w - 1);
		l += scores->at(i - 1);
		r -= scores->at(i);
		r += scores->at(i + w);
		second->push_back(round(l + r - d * scores->at(i)));
	}

	// For testing only
	/*
	 for (int i = 0; i < second->size(); i++) {
	 cout << second->at(i) << " ";
	 }
	 cout << endl;
	 */
}

void DetectorMaxima::findMaxima() {
	int firstSize = first->size();

	for (int i = 1; i < firstSize; i++) {
		double magnitude = abs(first->at(i - 1) - first->at(i));

		if (first->at(i) == 0 || (first->at(i - 1) < 0 & first->at(i) > 0)
				|| (first->at(i - 1) > 0 && first->at(i) < 0)) {
			if (second->at(i) < 0) {
				// Adjust index
				int peakIndex = i + w + segStart;

				// Record the index of the peak and its magnitude
				/*
				 vector<double> * pair = new vector<double>();
				 pair->push_back(peakIndex);
				 pair->push_back(magnitude);
				 allMaxima->push_back(pair);
				 */

				// Make sure that the peak is in a high-scoring region of width s centered on the peak
				if (magnitude > m) {
					// Make sure that the peak is in a high-scoring region of width s centered on the peak
					int peakStart = peakIndex - halfS;
					if (peakStart < segStart) {
						peakStart = segStart;
					}
					int peakEnd = peakIndex + halfS;
					if (peakEnd > segEnd) {
						peakEnd = segEnd;
					}

					double count = countLessThan(oScores, peakStart, peakEnd,
							t);
					double v = (100.00 * count)
							/ ((double) peakEnd - peakStart + 1);
					if (v < p) {
						maxima->push_back(peakIndex);
					}
				}
			}
		}
	}

	// Testing - start
	/*
	cout << "Maxima: " << endl;
	for (int i = 0; i < maxima->size(); i++) {
		cout << maxima->at(i) << " ";
	}
	cout << endl << endl;
	*/
	// Testing - end
}

int DetectorMaxima::countLessThan(vector<int> * list, int s, int e, double t) {
	int count = 0;
	for (int u = s; u <= e; u++) {
		if (list->at(u) < t) {
			count++;
		}
	}
	return count;
}

void DetectorMaxima::findSeparators() {
	int n = maxima->size();

	if (n > 0) {
		for (int i = 0; i < n - 1; i++) {
			int j = i + 1;
			int s = maxima->at(i);
			int e = maxima->at(j);

			double count = countLessThan(oScores, s, e, t);
			double v = (100.00 * count) / ((double) e - s + 1);
			if (v >= p) {
				separatorList->push_back(new Location(s, e));
			}
		}
	}

	// For testing only
	/*
	 cout << "Separators: " << endl;
	 for (int h = 0; h < separatorList->size(); h++) {
	 cout << separatorList->at(h)->toString() << endl;
	 }
	 cout << endl;
	 */
}

void DetectorMaxima::findRegions() {
	// Determine regions
	int maximaCount = maxima->size();
	if (maximaCount > 0) {
		int segStart = maxima->at(0);
		int separatorCount = separatorList->size();
		for (int k = 0; k < separatorCount; k++) {
			int segEnd = separatorList->at(k)->getStart();
			regionList->push_back(new Location(segStart, segEnd));
			segStart = separatorList->at(k)->getEnd();
		}
		regionList->push_back(
				new Location(segStart, maxima->at(maximaCount - 1)));
	}

	// For testing only
	/*
	 cout << "Regions: " << endl;
	 for (int r = 0; r < regionList->size(); r++) {
	 cout << regionList->at(r)->toString() << endl;
	 }
	 cout << endl;
	 */
	// End testing
}

/*
 *
 */
void DetectorMaxima::extendRegions() {
	int regionCount = regionList->size();
	int gg = 0;
	while (gg < regionCount) {
		ILocation * region = regionList->at(gg);

		int regionStart = region->getStart();
		int regionEnd = region->getEnd();

		// Handle the case where the region is made of one nucleotide
		if (regionStart == regionEnd) {
			regionStart = regionStart - halfS;
			if (regionStart < segStart) {
				regionStart = segStart;
			}
			region->setStart(regionStart);

			regionEnd = regionEnd + halfS;
			if (regionEnd > segEnd) {
				regionEnd = segEnd;
			}
			region->setEnd(regionEnd);
		}

		// Left end: Extend step by step
		int lEnd = (gg == 0) ? segStart : regionList->at(gg - 1)->getEnd();
		for (int u = regionStart; u >= lEnd; u = u - e) {
			int d = u - e + 1;
			if (d < lEnd) {
				d = lEnd;
			}
			double v = (100.0 * countLessThan(oScores, d, u, t)) / ((double) e);
			if (v >= p) {
				break;
			} else {
				regionStart = d;
			}
		}

		// Left end: Extend or erode base by base
		if (oScores->at(regionStart) < t) {
			for (int a = regionStart; a < regionEnd; a++) {
				if (oScores->at(a) >= t) {
					regionStart = a;
					break;
				}
			}
		} else {
			for (int a = regionStart; a >= lEnd; a--) {
				if (oScores->at(a) >= t) {
					regionStart = a;
				} else {
					break;
				}
			}
		}

		// Set new start to check for validity
		region->setStart(regionStart);

		// Right end: extend to the right step by step
		int rEnd =
				(gg == regionCount - 1) ?
						segEnd : regionList->at(gg + 1)->getStart();
		for (int u = regionEnd; u <= rEnd; u = u + e) {
			int d = u + e - 1;
			if (d > rEnd) {
				d = rEnd;
			}
			double v = (100.0 * countLessThan(oScores, u, d, t)) / ((double) e);
			if (v >= p) {
				break;
			} else {
				regionEnd = d;
			}
		}

		// Right end: extend or erod base by base
		if (oScores->at(regionEnd) < t) {
			for (int a = regionEnd; a > regionStart; a--) {
				if (oScores->at(a) >= t) {
					regionEnd = a;
					break;
				}
			}
		} else {
			for (int a = regionEnd; a <= rEnd; a++) {
				if (oScores->at(a) >= t) {
					regionEnd = a;
				} else {
					break;
				}
			}
		}

		// Set new end to check for validity
		region->setEnd(regionEnd);

		// Merge overlapping regions
		if (gg > 0) {
			ILocation * pRegion = regionList->at(gg - 1);
			int pStart = pRegion->getStart();
			int pEnd = pRegion->getEnd();

			if (Util::isOverlapping(pStart, pEnd, regionStart, regionEnd)) {
				pRegion->setEnd(regionEnd);
				regionList->erase(regionList->begin() + gg);
				regionCount = regionList->size();
			} else {
				gg++;
			}
		}

		if (gg == 0) {
			gg++;
		}
	}

	// Testing - Start
	/*
	 cout << "Extended regions: " << endl;
	 for (int r = 0; r < regionList->size(); r++) {
	 cout << regionList->at(r)->toString() << endl;
	 }
	 cout << endl;
	 */
	// Testing - End
}

} /* namespace nonltr */
```
