## Supplementary Data 1 for "FASTCAR: Rapid alignment-free prediction of sequence alignment identity scores using self-supervised general linear models": DetectorMaxima.h

```
/*
 * DetectorMaxima.h
 *
 *  Created on: May 31, 2013
 *      Author: Hani Zakaria Girgis, PhD
 */

#ifndef DETECTORMAXIMA_H_
#define DETECTORMAXIMA_H_

#include <vector>
#include <math.h>

#include "../utility/ILocation.h"

using namespace std;
using namespace utility;

namespace nonltr {

class DetectorMaxima {
private:

	int segStart;
	int segEnd;
	double s;
	double w;
	double m;
	double t;
	double p;
	int e;
	int halfS;

	vector<int> * oScores;
	vector<double> * scores;
	vector<double> * mask;
	vector<double> * first;
	vector<double> * second;
	vector<int> * maxima;
	// vector<vector<double> *> * allMaxima;

	vector<ILocation *> * separatorList;
	vector<ILocation *> * regionList;

	void makeMask();
	void smooth();
	void deriveFirst();
	void deriveSecond();
	void findMaxima();

	void findSeparators();
	void findRegions();

	void extendRegions();

	int countLessThan(vector<int> *, int, int, double);

	/**
	 * Credit: http://stackoverflow.com/questions/554204/where-is-round-in-c
	 */
	inline double round(double number) {
		return number < 0.0 ? ceil(number - 0.5) : floor(number + 0.5);
	}

public:
	DetectorMaxima(int, int, double, double, double, double, double, int,
			vector<int> *);
	virtual ~DetectorMaxima();
	const vector<ILocation*>* getRegionList() const;
	const vector<double>* getFirst() const;
	const vector<double>* getSecond() const;

	// const vector<vector<double> *>* getAllMaxima() const;
};

} /* namespace nonltr */
#endif /* DETECTORMAXIMA_H_ */
```
