## Supplementary Data 1 for "FASTCAR: Rapid alignment-free prediction of sequence alignment identity scores using self-supervised general linear models": EnrichmentMarkovView.cpp

```
/*
 * EnrichmentMarkovView.cpp
 *
 *  Created on: Apr 17, 2013
 *      Author: Hani Zakaria Girgis, PhD
 */

namespace nonltr {

/**
 * The Markov order. It start at 0.
 */
template<class I, class V>
EnrichmentMarkovView<I, V>::EnrichmentMarkovView(int k, int order, int m) :
		minObs(m), factor(10000.00), KmerHashTable<I, V>(k) {
	initialize(order);
}

template<class I, class V>
EnrichmentMarkovView<I, V>::EnrichmentMarkovView(int k, V initValue, int order,
		int m) :
		minObs(m), factor(10000.00), KmerHashTable<I, V>(k, initValue) {
	initialize(order);
}

template<class I, class V>
void EnrichmentMarkovView<I, V>::initialize(int order) {
	// Test start
	// cout << "Testing: " << minObs << endl;
	// Test end

	o = order;
	if (o < 0) {
		string msg("The Markov order must be non-negative integer. ");
		msg.append("The invalid input is: ");
		msg.append(Util::int2string(o));
		msg.append(".");
		throw InvalidInputException(msg);
	}

	if (o >= KmerHashTable<I, V>::k) {
		string msg("The Markov order cannot be >= k (k-mer).");
		throw InvalidInputException(msg);
	}

	l = 0;
	modelList = new vector<KmerHashTable<int, int> *>();

	for (int i = 1; i <= o + 1; i++) {
		modelList->push_back(new KmerHashTable<int, int>(i));
	}
}

template<class I, class V>
EnrichmentMarkovView<I, V>::~EnrichmentMarkovView() {
	Util::deleteInVector(modelList);
	delete modelList;
}

/**
 * This method count words of size 1 to order+1 in the input sequence.
 * In other words, it updates the background tables. In addition, it
 * updates the length of the genome.
 *
 * sequence: is the input sequence.
 * start: the start index - inclosing.
 * end: the end index - inclosing.
 */
template<class I, class V>
void EnrichmentMarkovView<I, V>::count(const char * sequence, int start,
		int end) {

	// Multiple by 2 if scanning the forward strand and its reverse complement
	// l = l + (2 * (end - start + 1));
	l = l + (end - start + 1);

	int modelNumber = modelList->size();
	for (int i = 0; i < modelNumber; i++) {
		KmerHashTable<int, int> * t = modelList->at(i);
		t->wholesaleIncrement(sequence, start, end - i);
	}
}

/**
 * Normalize the count of words in each model.
 * Values stored in these models are multiplied by "factor."
 */
template<class I, class V>
void EnrichmentMarkovView<I, V>::generateProbapilities() {
	int modelNumber = modelList->size();

	for (int m = 0; m < modelNumber; m++) {
		KmerHashTable<int, int> * t = modelList->at(m);
		int tSize = t->getMaxTableSize();

		for (int i = 0; i < tSize; i += 4) {
			double sum = 0.0;

			for (int j = i; j < i + 4; j++) {
				sum += t->valueOf(j);
			}

			for (int j = i; j < i + 4; j++) {
				t->insert(j, round(factor * ((double) t->valueOf(j) / sum)));
			}
		}
	}
}

template<class I, class V>
void EnrichmentMarkovView<I, V>::processTable() {
	char base = 4;
	int modelNumber = modelList->size();

	// Make a zero in quaternary form as a string of length k.
	string q("");
	for (int x = 0; x < KmerHashTable<I, V>::k; x++) {
		q.append(1, 0);
	}

	double lowerP;
	double upperP;
	for (I y = 0; y < KmerHashTable<I, V>::maxTableSize; y++) {
		if (y % 10000000 == 0) {
			cout << "Processing " << y << " keys out of "
					<< KmerHashTable<I, V>::maxTableSize;
			cout << endl;
		}

		const char * qc = q.c_str();

		// Calculate the expected number of occurrences.

		// a. Calculate probability from lower order models.
		// Lower probabilities are the same for four consecutive words of length of k-1
		if (y % 4 == 0) {
			lowerP = 1.0;
			for (int m = 0; m < modelNumber - 1; m++) {
				KmerHashTable<int, int> * oTable = modelList->at(m);
				lowerP *= (((double) oTable->valueOf(qc, 0)) / factor);
			}
		}

		// b. Calculate probability based on the specified order.
		KmerHashTable<int, int> * oTable = modelList->at(modelNumber - 1);
		int resultsSize = KmerHashTable<I, V>::k - o - 1;

		// Upper probabilities are the same for four consecutive words of length of k-1
		// The scanning of words or length corresponding to the highest order + 1
		// This step is not needed if k = o + 1, i.e. resultsSize = 0.
		if (y % 4 == 0) {
			if (resultsSize > 0) {
				//Initialize the elements of the vector invalid index
				vector<int> results = vector<int>(resultsSize, -987);
				oTable->wholesaleValueOf(qc, 0, resultsSize - 1, &results, 0);

				upperP = 1.0;
				for (int i = 0; i < resultsSize; i++) {
					upperP *= (((double) results.at(i)) / factor);
				}
				results.clear();

			} else {
				upperP = 1.0;
			}
		}

		// The expected number of occurances
		double exp = l * lowerP * upperP
				* (((double) oTable->valueOf(qc, resultsSize)) / factor);

		// Calculate the enrichment value.
		// Log value
		// values[y] = round((log((double) values[y] + 1.0) - log(exp + 1.0)));

		// Raw value
		// Requirement: if observed is >= 5 && observed > expected then the value is the difference
		// otherwise the value is zero

		V observed = KmerHashTable<I, V>::values[y];

		if (observed >= minObs && observed > exp) {

			KmerHashTable<I, V>::values[y] = round(observed - exp);
		} else {
			KmerHashTable<I, V>::values[y] = 0;
		}

		/*
		 KmerHashTable<I, V>::values[y] =
		 round(
		 (((double) KmerHashTable<I, V>::values[y] + 1.0)
		 / (exp + 1.0)));
		 */

		// Increment the quaternary number:
		// 1 - guard against overflow.
		if (q[0] == base - 1) {
			string z("");
			z.append(1, 0);
			q = z + q;
		}

		// 2 - increment the quaternary number by 1.
		int qLen = q.size();
		for (int i = qLen - 1; i >= 0; i--) {
			if (q[i] + 1 < base) {
				q[i] = q[i] + 1;
				break;
			} else {
				q[i] = 0;
			}
		}
	}
}

} /* namespace nonltr */
```
