## Supplementary Data 1 for "FASTCAR: Rapid alignment-free prediction of sequence alignment identity scores using self-supervised general linear models": EnrichmentMarkovView.h

```
/*
 * EnrichmentMarkovView.h
 *
 *  Created on: Apr 17, 2013
 *      Author: Hani Zakaria Girgis, PhD
 */

#ifndef ENRICHMENTMARKOVVIEW_H_
#define ENRICHMENTMARKOVVIEW_H_

#include <cmath>
#include <vector>
#include <iostream>

#include "KmerHashTable.h"
#include "../utility/Util.h"
#include "../exception/InvalidInputException.h"

using namespace std;
using namespace utility;
using namespace exception;

namespace nonltr {

template<class I, class V>
class EnrichmentMarkovView: public KmerHashTable<I,V>{

private:
	// The minimum number of the observed k-mers
	const int minObs;

	// This template specification should work up to order of 14,
	// i.e. word length = 15
	vector<KmerHashTable<int,int> *> * modelList;

	// Markov order
	int o;

	// Total length
	long l;

	// Multiplied the probability of word by this factor
	// Equivalent to four decimal points.
	const double factor;	// = 10000.00;

	// Initialize data members
	void initialize(int);

	/**
	 * Credit: http://stackoverflow.com/questions/554204/where-is-round-in-c
	 */
	inline double round(double number) {
		return number < 0.0 ? ceil(number - 0.5) : floor(number + 0.5);
	}

public:
	EnrichmentMarkovView(int, int, int);
	EnrichmentMarkovView(int, V, int, int);
	virtual ~EnrichmentMarkovView();

	void count(const char *, int, int);
	void generateProbapilities();
	void processTable();
};
} /* namespace nonltr */

#include "EnrichmentMarkovView.cpp"

#endif /* ENRICHMENTMARKOVVIEW_H_ */
```
