## Supplementary Data 1 for "FASTCAR: Rapid alignment-free prediction of sequence alignment identity scores using self-supervised general linear models": HMM.cpp

```
/*
 * HMM.cpp
 *
 *  Created on: Jun 21, 2013
 *      Author: Hani Zakaria Girgis, PhD
 */

#include "HMM.h"

#include <iostream>
#include <fstream>

#include "../utility/Util.h"
#include "../exception/InvalidStateException.h"
#include "../exception/InvalidInputException.h"
#include "../exception/FileDoesNotExistException.h"
#include "../exception/InvalidOperationException.h"

using namespace std;
using namespace utility;
using namespace exception;

namespace nonltr {

HMM::HMM(string hmmFile) :
		PRECISION(numeric_limits<double>::digits10 + 1) {
	// ToDo: Fix this operation
	string msg("Reading HMM from file is temporarily disabled.");
	throw InvalidOperationException(msg);

	cerr << "Building HMM from: " << hmmFile << endl;

	ifstream in(hmmFile.c_str());
	in.precision(PRECISION);

	if (in) {
		string token;
		bool isLogBase = false;
		bool isStates = false;
		bool isPriors = false;
		bool isTransition = false;

		while (in >> token) {
			if (isLogBase) {
				base = atof(token.c_str());

				checkBase(base);

				logBase = log(base);
				isLogBase = false;
			} else if (isStates) {
				stateNumber = atoi(token.c_str());
				positiveStateNumber = stateNumber / 2;
				initializeHelper();

				isStates = false;
			} else if (isPriors) {
				//Skip state names
				for (int i = 1; i < stateNumber; i++) {
					in >> token;
				}
				for (int i = 0; i < stateNumber; i++) {
					in >> token;
					(*pList)[i] = atof(token.c_str());
				}

				isPriors = false;
			} else if (isTransition) {
				//Skip state names
				for (int i = 1; i < stateNumber; i++) {
					in >> token;
				}

				for (int i = 0; i < stateNumber; i++) {
					//Skip the name of the state at the beginning of the line
					for (int j = -1; j < stateNumber; j++) {
						in >> token;
						if (j > -1) {
							(*(tList->at(i)))[j] = atof(token.c_str());
						}
					}
				}

				isTransition = false;
			}

			if (token.compare("Base") == 0) {
				isLogBase = true;
			} else if (token.compare("States") == 0) {
				isStates = true;
			} else if (token.compare("Priors") == 0) {
				isPriors = true;
			} else if (token.compare("Transition") == 0) {
				isTransition = true;
			}
		}

		in.close();
	} else {
		string msg(hmmFile);
		msg.append(" does not exist.");
		throw FileDoesNotExistException(msg);
	}
	in.close();

	//print("/Users/zakarota/Data/HgTest/Rep/Test/genome/hmmTest.txt");
}

/**
 * Use this constructor to train on the entire genome.
 * The client has to call train on each chromosome.
 * base is the threshold.

 */
HMM::HMM(double base, int stateNumber) :
		PRECISION(numeric_limits<double>::digits10 + 1) {
	initialize(base, stateNumber);
}

void HMM::initialize(double baseIn, int stateNumberIn) {
	base = baseIn;
	checkBase(base);

	logBase = log(baseIn);

	stateNumber = stateNumberIn;
	// Make sure that the number of states is even and > 0
	if (stateNumber % 2 != 0 || stateNumber == 0) {
		string msg("The number of states must be even and > zero.");
		throw InvalidInputException(msg);
	}

	positiveStateNumber = stateNumber / 2;
	cout << "The number of states is: " << stateNumber << endl;

	initializeHelper();
}

/**
 * This method makes sure that the base is not zero.
 */
void HMM::checkBase(double base) {
	if (fabs(base - 0.0) < std::numeric_limits<double>::epsilon()) {
		string msg("The base cannot be zero because log(base) is not defined.");
		throw InvalidInputException(msg);
	}
}

void HMM::initializeHelper() {
	// Ensure that the number of the states is positive
	if (stateNumber < 1) {
		string msg("The number of states must be positive.");
		throw InvalidStateException(msg);
	}

	pList = new vector<double>(stateNumber, 1);
	tList = new vector<vector<double> *>;
	for (int i = 0; i < stateNumber; i++) {
		tList->push_back(new vector<double>(stateNumber, 1));
	}
	oList = new vector<double>(stateNumber, 1);

	// Check if infinity can be handled
	if (!std::numeric_limits<double>::has_infinity) {
		string msg("This compiler does not handle infinite values. ");
		msg.append(string("The decoding algorithm will not function."));
		throw InvalidStateException(msg);
	} else {
		minusInf = -1.0 * std::numeric_limits<double>::infinity();
	}
}

HMM::~HMM() {
	pList->clear();
	delete pList;

	Util::deleteInVector(tList);
	delete tList;

	oList->clear();
	delete oList;
}

void HMM::train(vector<int> * scoreListIn,
		const vector<vector<int> *> * segmentListIn,
		const vector<ILocation*> * candidateListIn) {

	scoreList = scoreListIn;
	segmentList = segmentListIn;
	candidateList = candidateListIn;

	int candidateCount = candidateList->size();
	if (candidateCount > 0) {
		int firstCandIndex = 0;
		int lastCandIndex = 0;
		int segmentNumber = segmentList->size();
		for (int i = 0; i < segmentNumber; i++) {
			vector<int> * s = segmentList->at(i);
			ILocation * c = candidateList->at(firstCandIndex);
			// A segment may have no detections
			if (Util::isOverlapping(s->at(0), s->at(1), c->getStart(),
					c->getEnd())) {
				lastCandIndex = trainHelper1(s->at(0), s->at(1),
						firstCandIndex);
				trainHelper2(s->at(0), s->at(1), firstCandIndex, lastCandIndex);
				firstCandIndex = lastCandIndex + 1;
				if (firstCandIndex >= candidateCount) {
					break;
				}
			}
		}
	}
}

int HMM::trainHelper1(int segStart, int segEnd, int firstCandIndex) {
	ILocation * cand = candidateList->at(firstCandIndex);
	if (!Util::isOverlapping(segStart, segEnd, cand->getStart(),
			cand->getEnd())) {
		string msg("The first candidate is not overlapping with the segment. ");
		msg.append("Candidate location is: ");
		msg.append(cand->toString());
		msg.append(" Segment location is: ");
		msg.append(Util::int2string(segStart));
		msg.append("-");
		msg.append(Util::int2string(segEnd));
		throw InvalidInputException(msg);
	}

	int lastCandIndex = -1;
	int candidateNumber = candidateList->size();
	for (int c = firstCandIndex; c < candidateNumber; c++) {
		ILocation * cand = candidateList->at(c);
		if (Util::isOverlapping(segStart, segEnd, cand->getStart(),
				cand->getEnd())) {
			lastCandIndex = c;
		} else {
			break;
		}
	}

	if (lastCandIndex < 0) {
		string msg("The index of the last candidate cannot be negative.");
		throw InvalidStateException(msg);
	}

	return lastCandIndex;
}

void HMM::trainHelper2(int segStart, int segEnd, int firstCandIndex,
		int lastCandIndex) {
	ILocation * f = candidateList->at(firstCandIndex);

	// First negative region if present
	int fStart = f->getStart();
	if (fStart > segStart) {
		trainNegative(segStart, fStart - 1);
		move(getNgtvState(fStart - 1), getPstvState(fStart));
	}

	// Alternating positive and negative regions
	for (int i = firstCandIndex; i < lastCandIndex; i++) {
		ILocation * c = candidateList->at(i);
		int cStart = c->getStart();
		int cEnd = c->getEnd();
		trainPositive(cStart, cEnd);
		move(getPstvState(cEnd), getNgtvState(cEnd + 1));

		int nextStart = candidateList->at(i + 1)->getStart();
		trainNegative(cEnd + 1, nextStart - 1);
		move(getNgtvState(nextStart - 1), getPstvState(nextStart));
	}

	// Last positive region
	ILocation * l = candidateList->at(lastCandIndex);
	int lEnd = l->getEnd();
	trainPositive(l->getStart(), lEnd);

	// Last negative region if present
	if (segEnd > lEnd) {
		move(getPstvState(lEnd), getNgtvState(lEnd + 1));
		trainNegative(lEnd + 1, segEnd);
	}
}

void HMM::trainPositive(int s, int e) {
	int pIndex = getPstvState(s);
	(*pList)[pIndex] = pList->at(pIndex) + 1;

	for (int i = s; i <= e; i++) {
		int index = getPstvState(i);
		(*oList)[index] = oList->at(index) + 1;
	}

	for (int i = s; i < e; i++) {
		move(getPstvState(i), getPstvState(i + 1));
	}
}

void HMM::trainNegative(int s, int e) {
	int pIndex = getNgtvState(s);
	(*pList)[pIndex] = pList->at(pIndex) + 1;

	for (int i = s; i <= e; i++) {
		int index = getNgtvState(i);
		(*oList)[index] = oList->at(index) + 1;
	}

	for (int i = s; i < e; i++) {
		move(getNgtvState(i), getNgtvState(i + 1));
	}
}

void HMM::move(int state1, int state2) {
	vector<double> * state1Row = tList->at(state1);
	(*state1Row)[state2] = state1Row->at(state2) + 1;
}

void HMM::normalize() {
// Priors
	double sum = 0.0;
	for (int i = 0; i < stateNumber; i++) {
		sum += pList->at(i);
	}
	for (int i = 0; i < stateNumber; i++) {
		(*pList)[i] = log(pList->at(i) / sum);
	}

// Output
	for (int i = 0; i < stateNumber; i++) {
		(*oList)[i] = log(1.0);
	}

// Transition
	for (int i = 0; i < stateNumber; i++) {
		vector<double> * row = tList->at(i);
		double sum = 0.0;
		for (int j = 0; j < stateNumber; j++) {
			sum += row->at(j);
		}

		for (int j = 0; j < stateNumber; j++) {
			(*row)[j] = log(row->at(j) / sum);
		}
	}
}

void HMM::print() {
	cout.precision(PRECISION);

	// State names
	vector<string> v;
	for (int j = 0; j < positiveStateNumber; j++) {
		v.push_back(Util::int2string(j));
	}
	string m("-");
	for (int j = 0; j < positiveStateNumber; j++) {
		v.push_back(m + Util::int2string(j));
	}

	cout << "Priors:" << endl;
	for (int g = 0; g < 2; g++) {
		for (int i = 0; i < positiveStateNumber; i++) {
			cout << v.at(i + (g * positiveStateNumber)) << "\t";
		}

		for (int i = 0; i < positiveStateNumber; i++) {
			cout << pList->at(i + (g * positiveStateNumber)) << "\t";
		}
		cout << endl;
	}
	cout << endl;

	/*
	 cout << "Output:" << endl;
	 for (int i = 0; i < v.size(); i++) {
	 cout << v.at(i) << "\t";
	 }
	 cout << endl;
	 for (int i = 0; i < stateNumber; i++) {
	 cout << oCountList->at(i) << "\t";
	 }
	 cout << endl << endl;
	 */

	cout << "Transition:" << endl << "\t";
	for (int i = 0; i < v.size(); i++) {
		cout << v.at(i) << "\t";
	}
	cout << endl;

	for (int i = 0; i < stateNumber; i++) {
		vector<double> * row = tList->at(i);
		cout << v.at(i) << "\t";
		for (int j = 0; j < stateNumber; j++) {
			cout << row->at(j) << "\t";
		}
		cout << endl;
	}
	cout << endl << endl;
}

void HMM::print(string hmo) {
	ofstream out(hmo.c_str());
	out.precision(PRECISION);

	out << "Base" << endl << base << endl;

	out << "States" << endl << stateNumber << endl;

	vector<string> v;
	for (int j = 0; j < positiveStateNumber; j++) {
		v.push_back(Util::int2string(j));
	}
	string m("-");
	for (int j = 0; j < positiveStateNumber; j++) {
		v.push_back(m + Util::int2string(j));
	}

	out << "Priors" << endl;
	for (int i = 0; i < v.size(); i++) {
		out << v.at(i) << "    ";
	}
	out << endl;

	for (int i = 0; i < v.size(); i++) {
		out << pList->at(i) << "    ";
	}
	out << endl;

	out << "Transition" << endl << "\t";
	for (int i = 0; i < v.size(); i++) {
		out << v.at(i) << "\t";
	}
	out << endl;

	for (int i = 0; i < stateNumber; i++) {
		vector<double> * row = tList->at(i);
		out << v.at(i) << "\t";
		for (int j = 0; j < stateNumber; j++) {
			out << row->at(j) << "\t";
		}
		out << endl;
	}
	out << endl << endl;

	out.close();
}

/**
 * This method will append the state sequence to the end of the input state list
 * This method returns the log likelihood
 */
double HMM::decode(int rStart, int rEnd, vector<int> * scoreListIn,
		vector<int>& stateList) {
	scoreList = scoreListIn;

	// Make sure that the coordinates represent valid location
	Location check(rStart, rEnd);
	// End check

	vector<vector<double> > v(stateNumber);
	int size = rEnd - rStart + 1;
	for (int i = 0; i < stateNumber; i++) {
		v[i] = vector<double>(size, minusInf);
	}

	vector<vector<int> > p(stateNumber);
	for (int i = 0; i < stateNumber; i++) {
		p[i] = vector<int>(size, -1);
	}

	// Initialize
	int firstPstvState = getPstvState(rStart);
	int firstNgtvState = positiveStateNumber + firstPstvState;
	v[firstPstvState][0] = pList->at(firstPstvState);
	v[firstNgtvState][0] = pList->at(firstNgtvState);

	// Recurs
	for (int i = rStart + 1; i <= rEnd; i++) {
		int vIndex = i - rStart;

		// Obtain states from scores
		int pPstvState = getPstvState(i - 1);
		int pNgtvState = positiveStateNumber + pPstvState;
		int cPstvState = getPstvState(i);
		int cNgtvState = positiveStateNumber + cPstvState;

		// Set positive state
		double p1 = v[pPstvState][vIndex - 1]
				+ (*(*tList)[pPstvState])[cPstvState];
		double p2 = v[pNgtvState][vIndex - 1]
				+ (*(*tList)[pNgtvState])[cPstvState];
		if (p1 > p2) {
			v[cPstvState][vIndex] = p1;
			p[cPstvState][vIndex] = pPstvState;
		} else {
			v[cPstvState][vIndex] = p2;
			p[cPstvState][vIndex] = pNgtvState;
		}

		// Set negative state
		double p3 = v[pPstvState][vIndex - 1]
				+ (*(*tList)[pPstvState])[cNgtvState];
		double p4 = v[pNgtvState][vIndex - 1]
				+ (*(*tList)[pNgtvState])[cNgtvState];
		if (p3 > p4) {
			v[cNgtvState][vIndex] = p3;
			p[cNgtvState][vIndex] = pPstvState;
		} else {
			v[cNgtvState][vIndex] = p4;
			p[cNgtvState][vIndex] = pNgtvState;
		}
	}

	// Decode
	int lastBestState = 0;
	double lastBestValue = v[0][size - 1];
	for (int i = 1; i < stateNumber; i++) {
		double currentValue = v[i][size - 1];
		if (currentValue > lastBestValue) {
			lastBestState = i;
			lastBestValue = currentValue;
		}
	}

	int stateListOriginalSize = stateList.size();
	for (int i = stateListOriginalSize; i < stateListOriginalSize + size; i++) {
		stateList.push_back(-1);
	}

	stateList[stateListOriginalSize + size - 1] = lastBestState;
	for (int i = size - 1; i > 0; i--) {
		lastBestState = p[lastBestState][i];
		stateList[stateListOriginalSize + i - 1] = lastBestState;
	}

	// Make sure that no state in the results has the value of -1
	for (int i = stateListOriginalSize; i < stateListOriginalSize + size; i++) {
		if (stateList[i] == -1) {
			string msg("At least one state was not determined properly.");
			throw InvalidStateException(msg);
		}
	}

	// Test - start
	/*
	 bool canPrint = false;
	 for (int i = stateListOriginalSize; i < stateListOriginalSize + size; i++) {
	 if (stateList.at(i) >= positiveStateNumber) {
	 canPrint = true;
	 }
	 }
	 if (canPrint) {
	 for (int i = rStart; i <= rEnd; i++) {
	 cout << scoreList->at(i) << " ";
	 }
	 cout << endl;

	 for (int i = stateListOriginalSize; i < stateListOriginalSize + size;
	 i++) {
	 if (stateList.at(i) < positiveStateNumber) {
	 cout << "+";
	 } else {
	 cout << "-";
	 //cout << stateList.at(i) << " ";
	 }
	 }
	 cout << endl;
	 }
	 */

	// Test - end
	return lastBestValue;
}

/**
 * Append positive regions at the end of regionList
 */
double HMM::decode(int rStart, int rEnd, vector<int> * scoreListIn,
		vector<ILocation *>& regionList) {

	vector<int> stateList;
	double logLikelihood = decode(rStart, rEnd, scoreListIn, stateList);

	int size = stateList.size();
	bool inRpt = false;
	bool canFill = false;
	int s = -1;
	int e = -1;

	for (int i = 0; i < size; i++) {
		// Start a new repeat
		if (stateList.at(i) < positiveStateNumber && !inRpt) {
			inRpt = true;
			s = i;
		}
		// End a the current repeat
		else if (stateList.at(i) >= positiveStateNumber && inRpt) {
			e = i - 1;
			inRpt = false;
			canFill = true;
		}
		// If the current repeat at the end of the segment
		else if (i == size - 1 && inRpt) {
			e = i;
			inRpt = false;
			canFill = true;
		}
		// Extract features of the just recognized repeat
		if (canFill) {
			regionList.push_back(new Location(s + rStart, e + rStart));
			s = -1;
			e = -1;
			canFill = false;
		}
	}

	return logLikelihood;
}

int HMM::getPositiveStateNumber() {
	return positiveStateNumber;
}

double HMM::getBase() {
	return base;
}

}
/* namespace nonltr */
```
