## Supplementary Data 1 for "FASTCAR: Rapid alignment-free prediction of sequence alignment identity scores using self-supervised general linear models": HMM.h

```
/*
 * HMM.h
 *
 *  Created on: Jun 21, 2013
 *      Author: Hani Zakaria Girgis, PhD
 */

#ifndef HMM_H_
#define HMM_H_

#include <vector>
#include <math.h>
#include <limits>
#include <stdlib.h>

#include "../utility/ILocation.h"

using namespace std;
using namespace utility;

namespace nonltr {

class HMM {
private:
	const int PRECISION;
	double minusInf;
	vector<double> * pList;
	vector<vector<double> *> * tList;
	vector<double> * oList;

	void initializeHelper();
	// Returns the index of the last candidate in the segment
	int trainHelper1(int, int, int);
	void trainHelper2(int, int, int, int);
	void trainPositive(int, int);
	void trainNegative(int, int);
	void move(int, int);
	void checkBase(double);

	/*
	 inline int getPstvState(int score) {
	 int state = ceil(log(score) / logBase);
	 if (state < 0) {
	 state = 0;
	 }
	 return state;
	 }

	 inline int getNgtvState(int score) {
	 int state = ceil(log(score) / logBase);
	 if (state < 0) {
	 state = 0;
	 }
	 return state + positiveStateNumber;
	 }
	 */

	inline int getPstvState(int index) {
		int state = scoreList->at(index);
		return state;
	}

	inline int getNgtvState(int index) {
		int state = scoreList->at(index);
		return state + positiveStateNumber;
	}

protected:
	double base;
	double logBase;
	int stateNumber;
	int positiveStateNumber;

	vector<int> * scoreList;
	const vector<vector<int> *> * segmentList;
	const vector<ILocation*> * candidateList;

	void initialize(double, int);
	/**
	 * Credit: http://stackoverflow.com/questions/554204/where-is-round-in-c
	 */
	inline double round(double number) {
		return number < 0.0 ? ceil(number - 0.5) : floor(number + 0.5);
	}

public:
	HMM(string); // Build a model from file
	HMM(double, int);
	// HMM(vector<int> *, const vector<vector<int> *> *,
	//		const vector<ILocation*> *, double);
	virtual ~HMM();
	void train(vector<int> *, const vector<vector<int> *> *, const vector<ILocation*> *);
	void normalize();
	double decode(int, int, vector<int> *, vector<int>&);
	double decode(int, int, vector<int> *, vector<ILocation *>&);
	int getPositiveStateNumber();
	void print();
	void print(string);
	double getBase();
};

} /* namespace nonltr */
#endif /* HMM_H_ */
```
