## Supplementary Data 1 for "FASTCAR: Rapid alignment-free prediction of sequence alignment identity scores using self-supervised general linear models": IChromosome.h

```
/*
 * IChromosome.h
 *
 *  Created on: Feb 4, 2013
 *      Author: Hani Zakaria Girgis, PhD
 */

#ifndef ICHROMOSOME_H_
#define ICHROMOSOME_H_

#include <string>
#include <vector>

using namespace std;

namespace nonltr {

class IChromosome {
public:
	//IChromosome();
	//virtual ~IChromosome();
	virtual const string* getBase() = 0;
	virtual const vector<vector<int> *> * getSegment() = 0;
	virtual string getHeader() = 0;
};

} /* namespace tr */
#endif /* ICHROMOSOME_H_ */
```
