## Supplementary Data 1 for "FASTCAR: Rapid alignment-free prediction of sequence alignment identity scores using self-supervised general linear models": ITableView.h

```
/*
 * ITableView.h
 *
 *  Created on: Aug 9, 2012
 *      Author: Hani Zakaria Girgis, PhD
 */

#ifndef ITABLEVIEW_H_
#define ITABLEVIEW_H_

#include <vector>

using namespace std;

namespace nonltr {

template<class I, class V>
class ITableView {
public:
	virtual V valueOf(const char*) = 0 ;
	virtual V valueOf(const char*, int) = 0;
	virtual V valueOf(I) = 0;

	virtual int getK() = 0;
	virtual I getMaxTableSize() = 0;
	virtual const V * getValues() const = 0;

	virtual void wholesaleValueOf(const char *, int, int, vector<V> *) = 0;
	virtual void wholesaleValueOf(const char *, int, int, vector<V> *, int) = 0;
};

}

#endif /* ITABLEVIEW_H_ */
```
