## Supplementary Data 1 for "FASTCAR: Rapid alignment-free prediction of sequence alignment identity scores using self-supervised general linear models": KmerHashTable.cpp

```
/*
 * KmerHashTable.cpp
 *
 *  Created on: Jul 25, 2012
 *      Author: Hani Zakaria Girgis, PhD
 */
#include <stdio.h>
#include <math.h>
#include <iostream>
#include <fstream>
#include <limits>

#include "../utility/Util.h"
#include "../exception/InvalidInputException.h"
#include "../exception/InvalidStateException.h"

using namespace std;
using namespace exception;
using namespace nonltr;
using namespace utility;

template<class I, class V>
KmerHashTable<I, V>::KmerHashTable(int keyLength) {
	initialize(keyLength, 0);
}

template<class I, class V>
KmerHashTable<I, V>::KmerHashTable(int keyLength, V initValue) {
	initialize(keyLength, initValue);
}

template<class I, class V>
void KmerHashTable<I, V>::initialize(int keyLength, V initialValueIn) {
	/*
	 if (keyLength > maxKeyLength) {
	 string msg = "The maximum size (k) of the k-mer is ";
	 char temp[3];
	 sprintf(temp, "%d", maxKeyLength);
	 msg += temp;
	 throw InvalidInputException(msg);
	 }
	 */

	k = keyLength;
	initialValue = initialValueIn;

	// Initialize bases
	bases = new I[k];
	for (int i = k - 1; i >= 0; i--) {
		bases[k - 1 - i] = (I) pow(4.0, i);
	}

	// Initialize mMinusOne
	mMinusOne = new I[4];
	for (int i = 0; i < 4; i++) {
		mMinusOne[i] = i * bases[0];
	}

	// Get maximum size of table
	char * temp = new char[k];
	for (int i = 0; i < k; i++) {
		temp[i] = 3;
	}

	maxTableSize = hash(temp) + 1;
	delete[] temp;

	// Initialize values
	values = new V[maxTableSize];
	for (I i = 0; i < maxTableSize; i++) {
		values[i] = initialValue;
	}

	// Test
	/*
	 char key[] = { 3, 3, 3, 3, 0, 0, 0, 0 };
	 long value = 100;
	 insert(key, 4, value);
	 long index = hash(key, 4);
	 cout << "Index: " << index << " " << values[index] << endl;
	 cout << "Index: " << index << " " << valueOf(key, 4) << endl;
	 cout << "Number of filled entries: " << countNonZeroEntries() << endl;
	 */
}

template<class I, class V>
KmerHashTable<I, V>::~KmerHashTable() {
	delete[] bases;
	delete[] mMinusOne;
	delete[] values;
}

/**
 * word: an array of characters.
 * The maximum integer value is 3 and the minimum is 0
 */
template<class I, class V>
I KmerHashTable<I, V>::hash(const char * key) {
	return hash(key, 0);
}

/**
 * seq: an array of characters e.g. [0,0,1,1,1,3,2].
 * start: the start index of the key.
 * This method is designed to process a long sequence.
 */
template<class I, class V>
I KmerHashTable<I, V>::hash(const char * sequence, int keyStart) {
	I index = 0;
	for (int i = 0; i < k; i++) {
		char nucleotide = sequence[keyStart + i];
		if (nucleotide >= 0 && nucleotide <= 3) {
			index += bases[i] * sequence[keyStart + i];
		} else {
			string msg("The value of the char representing the nucleotide ");
			msg.append("must be between 0 and 3.");
			msg.append("The int value is ");
			msg.append(Util::int2string((int) nucleotide));
			msg.append(" of nucleotide at index ");
			msg.append(Util::int2string(keyStart + i));

			for (int h = 0 + keyStart; h < k + keyStart; h++) {
				cerr << (int) sequence[h];
			}
			cerr << endl;

			throw InvalidInputException(msg);
		}
	}
	return index;
}

template<class I, class V>
void KmerHashTable<I, V>::hash(const char * sequence, int start, int end,
		vector<I> * hashList) {

	for (int i = start; i <= end; i++) {
		char nucleotide = sequence[i];
		if (!(nucleotide >= 0 && nucleotide <= 3)) {
			string msg("The value of the char representing the nucleotide ");
			msg.append("must be between 0 and 3.");
			msg.append("The int value is ");
			msg.append(Util::int2string((int) nucleotide));
			msg.append(" of nucleotide at index ");
			msg.append(Util::int2string(i));

			throw InvalidInputException(msg);
		}
	}

	I lastHash = hash(sequence, start);
	hashList->push_back(lastHash);

	for (int i = start + 1; i <= end; i++) {
		I s1 = 4 * (lastHash - mMinusOne[(int) sequence[i - 1]])
				+ (int) sequence[i + k - 1];
		hashList->push_back(s1);
		lastHash = s1;
	}
}

/**
 * This method put the key-value pair in the table.
 * Note: keys are unique, i.e. no duplicate keys.
 */
template<class I, class V>
void KmerHashTable<I, V>::insert(const char* key, V value) {
	insert(key, 0, value);
}

/**
 * Similar to the above method.
 * The key begins at start in seq.
 * The length of the key is k.
 */
template<class I, class V>
void KmerHashTable<I, V>::insert(const char* sequence, int keyStart, V value) {
	values[hash(sequence, keyStart)] = value;
}

template<class I, class V>
void KmerHashTable<I, V>::insert(I keyHash, V value) {
	values[keyHash] = value;
}

/**
 * Call wholesaleIncrement on the segment itself.
 * Then, call it again on the reverse complement of this segment.
 *
 * sequence: is a long sequence usually a long segment of a chromosome.
 * sFirstKmer: is the start index of the first k-mer.
 * sLastKmer: is the start index of the last k-mer.
 */
template<class I, class V>
void KmerHashTable<I, V>::wholesaleIncrement(const char* sequence,
		int firstKmerStart, int lastKmerStart) {
	// Increment k-mer's in the forward strand
	vector<I> hashList = vector<I>();
	hash(sequence, firstKmerStart, lastKmerStart, &hashList);

	int size = hashList.size();
	for (int i = 0; i < size; i++) {
		I keyHash = hashList.at(i);
		if (keyHash >= maxTableSize) {
			cerr << "array out of bounds" << endl;
			throw "";
		}
		values[keyHash]++;
	}

	// Increment k-mer's in the reverse complement
	/*
	string rc("");
	Util::revCompDig(sequence, firstKmerStart, lastKmerStart + k - 1, &rc);

	hashList.clear();
	hash(rc.c_str(), 0, rc.size() - k, &hashList);
	size = hashList.size();

	for (int i = 0; i < size; i++) {
		I keyHash = hashList.at(i);
		values[keyHash]++;
	}*/
}


/**
 * Call wholesaleIncrement on the segment itself.
 * Then, call it again on the reverse complement of this segment.
 *
 * sequence: is a long sequence usually a long segment of a chromosome.
 * sFirstKmer: is the start index of the first k-mer.
 * sLastKmer: is the start index of the last k-mer.
 */
template<class I, class V>
int  KmerHashTable<I, V>::wholesaleIncrementNoOverflow(const char* sequence,
		int firstKmerStart, int lastKmerStart) {
	// Increment k-mer's in the forward strand
	vector<I> hashList = vector<I>();
	hash(sequence, firstKmerStart, lastKmerStart, &hashList);
	int ret = 0;
	int size = hashList.size();
	for (int i = 0; i < size; i++) {
		I keyHash = hashList.at(i);
		if (keyHash >= maxTableSize) {
			cerr << "array out of bounds" << endl;
			throw "";
		}
		if (values[keyHash] < std::numeric_limits<V>::max()) {
			values[keyHash]++;
		} else {
			ret = -1;
		}
	}
	return ret;
}

/**
 * Increment the entry associated with the key by one.
 */
template<class I, class V>
void KmerHashTable<I, V>::increment(const char* key) {
	increment(key, 0);
}

/**
 * Increment the value associated with the key starting at keyStart in the
 * sequence by one. Also, this method increments the count of the reverse complement
 * of the kmer by one.
 */
template<class I, class V>
void KmerHashTable<I, V>::increment(const char* sequence, int keyStart) {
	// Increment the count of the kmer by one.
	I index = hash(sequence, keyStart);
	values[index]++;

	// Generate the reverse complement of the kmer.
	char * rcKmer = new char[k];
	for (int j = 0; j < k; j++) {
		switch (sequence[j + keyStart]) {
		case 0:
			rcKmer[k - 1 - j] = 3;
			break;
		case 1:
			rcKmer[k - 1 - j] = 2;
			break;
		case 2:
			rcKmer[k - 1 - j] = 1;
			break;
		case 3:
			rcKmer[k - 1 - j] = 0;
			break;
		default:
			string msg = string("Invalid code of a nucleotide: ");
			msg.append(1, sequence[j + keyStart]);
			msg.append(". Valid codes are 0, 1, 2, and 3.");
			throw InvalidInputException(msg);
		}
	}

	// Update the count of the reverse complement of the kmer by one.
	I rcIndex = hash(rcKmer, 0);
	values[rcIndex]++;

	// Free memory
	delete[] rcKmer;
}

/**
 * Return the value associated with the key
 */
template<class I, class V>
V KmerHashTable<I, V>::valueOf(const char* key) {
	return valueOf(key, 0);
}

/**
 * Return the value associated with the key
 * The key is a substring of length k starting at keyStart in the sequence
 */
template<class I, class V>
V KmerHashTable<I, V>::valueOf(const char* sequence, int keyStart) {
	return values[hash(sequence, keyStart)];
}

template<class I, class V>
V KmerHashTable<I, V>::valueOf(I keyHash) {
	return values[keyHash];
}

template<class I, class V>
void KmerHashTable<I, V>::wholesaleValueOf(const char * sequence,
		int firstKmerStart, int lastKmerStart, vector<V> * results) {
	wholesaleValueOf(sequence, firstKmerStart, lastKmerStart, results, 0);
}

/**
 * The values are set in the results vector starting at the resultsStart.
 * The contents of vector "results" must be initialized.
 * Otherwise, the program will crash outputting: "segmentation fault 11"
 */
template<class I, class V>
void KmerHashTable<I, V>::wholesaleValueOf(const char * sequence,
		int firstKmerStart, int lastKmerStart, vector<V> * results,
		int resultsStart) {

	int index = resultsStart;
	vector<I> hashList = vector<I>();
	hash(sequence, firstKmerStart, lastKmerStart, &hashList);
	int size = hashList.size();

	for (int i = 0; i < size; i++) {
		(*results)[index] = values[hashList.at(i)];
		index++;
	}
}

/**
 * This method returns the number of occupied entries in the table.
 * A non-occupied entry has the initial value.
 */
template<class I, class V>
I KmerHashTable<I, V>::countNonInitialEntries() {
	I count = 0;
	for (I i = 0; i < maxTableSize; i++) {
		if (values[i] != initialValue) {
			count++;
		}
	}
	return count;
}

/**
 * Make a list of the k-mers.
 */
template<class I, class V>
vector<string>* KmerHashTable<I, V>::getKeys() {
	vector<char> * alpha = new vector<char>();
	alpha->push_back((char) 0);
	alpha->push_back((char) 1);
	alpha->push_back((char) 2);
	alpha->push_back((char) 3);

	vector<string> *words = new vector<string>();
	for (int h = 0; h < alpha->size(); h++) {
		words->push_back(string(1, alpha->at(h)));
	}

	int wLen = k;
	for (int i = 1; i < wLen; i++) {
		vector<string> *wordsAtItrI = new vector<string>();
		for (I j = 0; j < words->size(); j++) {
			for (int h = 0; h < alpha->size(); h++) {
				string w = string(words->at(j));
				w.append(1, alpha->at(h));
				wordsAtItrI->push_back(w);
			}
		}
		words->clear();
		delete words;
		words = new vector<string>(*wordsAtItrI);

		// Free memory
		wordsAtItrI->clear();
		delete wordsAtItrI;
	}

	// Free memory
	alpha->clear();
	delete alpha;
	return words;
}

/**
 * Print the contents of the whole table
 */
template<class I, class V>
void KmerHashTable<I, V>::printTable(string output) {
	vector<const char *> keys;
	//getKeys(keys);

	ofstream out(output.c_str());

	for (I i = 0; i < keys.size(); i++) {
		const char * kmer = keys.at(i);
		for (int j = 0; j < k; j++) {
			out << (int) kmer[j];
		}
		cerr << "Hash: " << hash(keys.at(i), 0) << endl;

		out << " -> " << values[hash(keys.at(i), 0)] << endl;
	}

	out.close();
	keys.clear();
}

template<class I, class V>
int KmerHashTable<I, V>::getK() {
	return k;
}

template<class I, class V>
I KmerHashTable<I, V>::getMaxTableSize() {
	return maxTableSize;
}

template<class I, class V>
const V * KmerHashTable<I, V>::getValues() const {
	return values;
}

/**
 * Call after building the table.
 * A negative value is a likely indication of overflow.
 */
template<class I, class V>
void KmerHashTable<I, V>::checkOverflow() {
	for (I y = 0; y < maxTableSize; y++) {
		if (values[y] < 0) {
			string msg("A negative value is a likely indication of overflow. ");
			msg.append(
					"To the developer, consider larger data type in KmerHashTable.");
			throw InvalidStateException(msg);
		}
	}
}

template<class I, class V>
V KmerHashTable<I, V>::getMaxValue() {
	V max = 0;
	for (I y = 0; y < maxTableSize; y++) {
		if (values[y] > max) {
			max = values[y];
		}
	}
	return max;
}
```
