## Supplementary Data 1 for "FASTCAR: Rapid alignment-free prediction of sequence alignment identity scores using self-supervised general linear models": KmerHashTable.h

```
/*
 * KmerHashTable.h
 *
 *  Created on: Jul 25, 2012
 *      Author: Hani Zakaria Girgis, PhD - NCBI/NLM/NIH
 */

#ifndef KMERHASHTABLE_H_
#define KMERHASHTABLE_H_

#include <string>
#include <vector>
#include "ITableView.h"

using namespace std;
using namespace nonltr;

namespace nonltr {

template<class I, class V>
class KmerHashTable: public ITableView<I,V> {

protected:
	/* Fields */
	static const int maxKeyLength = 15;
	int k;


	I maxTableSize;

	// The hashed values, i.e. the values of the hash table.
	// The index is the 4ry representation of the key
	V * values;
	V initialValue;

private:
	// [4^0, 4^1, ... , 4^(k-1)]
	I * bases;
	I * mMinusOne;
	void initialize(int, V);

public:
	/* Methods */
	KmerHashTable(int);
	KmerHashTable(int, V);

	virtual ~KmerHashTable();

	I hash(const char *);
	I hash(const char *, int);
	void hash(const char *, int, int, vector<I> *);

	void insert(const char*, V);
	void insert(const char*, int, V);
	void insert(I, V);

	void increment(const char*);
	void increment(const char*, int);
	void wholesaleIncrement(const char*, int, int);
	int wholesaleIncrementNoOverflow(const char*, int, int);

	void addReverseComplement();
	I countNonInitialEntries();
	vector<string> *getKeys();
	void printTable(string);
	void checkOverflow();

	/*Vritual methods from ITableView*/
	virtual V valueOf(const char*);
	virtual V valueOf(const char*, int);
	virtual V valueOf(I);
	virtual void wholesaleValueOf(const char *, int, int, vector<V> *);
	virtual void wholesaleValueOf(const char *, int, int, vector<V> *, int);

	virtual int getK();
	virtual I getMaxTableSize();
	virtual V getMaxValue();
	virtual const V * getValues() const;
};
}

#include "KmerHashTable.cpp"

#endif /* KMERHASHTABLE_H_ */
```
