## Supplementary Data 1 for "FASTCAR: Rapid alignment-free prediction of sequence alignment identity scores using self-supervised general linear models": LocationList.cpp

```
/*
 * LocationList.cpp
 *
 *  Created on: Feb 19, 2015
 *      Author: Hani Zakaria Girgis, PhD
 *
 *
 * An instance of this class holds a list of merged locations.
 */

#include "LocationList.h"

namespace nonltr {

LocationList::LocationList(string chromNameIn) {
	chromName = chromNameIn;
	regionList = new vector<ILocation *>();
	merge();
}

LocationList::~LocationList() {
	Util::deleteInVector(regionList);
	delete regionList;
}

void LocationList::add(int start, int end) {
	regionList->push_back(new Location(start, end));
}

void LocationList::merge() {
	int regionCount = regionList->size();
	int gg = 0;
	while (gg < regionCount) {
		ILocation * region = regionList->at(gg);

		int regionStart = region->getStart();
		int regionEnd = region->getEnd();

		if (gg > 0) {
			ILocation * pRegion = regionList->at(gg - 1);
			int pStart = pRegion->getStart();
			int pEnd = pRegion->getEnd();

			if (Util::isOverlapping(pStart, pEnd, regionStart, regionEnd)) {
				pRegion->setEnd(regionEnd > pEnd ? regionEnd : pEnd);
				regionList->erase(regionList->begin() + gg);
				delete region;
				regionCount = regionList->size();
			} else {
				gg++;
			}
		}

		if (gg == 0) {
			gg++;
		}
	}
}

void LocationList::mergeWithAnotherList(
		const vector<ILocation *> * const otherList) {
	//A pre-condition: Ensure that the other list is sorted
	for (int h = 1; h < otherList->size(); h++) {
		if (otherList->at(h)->getStart() < otherList->at(h - 1)->getStart()) {
			throw InvalidStateException(
					string("LocationList - The other list is not sorted."));
		}
	}

	// Start
	vector<ILocation *> * mergedList = new vector<ILocation *>();

	int i = 0;
	int j = 0;
	int iLimit = regionList->size();
	int jLimit = otherList->size();

	// Continue until one list is finished
	while (i < iLimit && j < jLimit) {
		ILocation * iLoc = regionList->at(i);
		ILocation * jLoc = otherList->at(j);

		if (iLoc->getStart() < jLoc->getStart()) {
			mergedList->push_back(iLoc);
			i++;
		} else {
			mergedList->push_back(new Location(*jLoc));
			j++;
		}
	}

	// Once one list is finished, copy the rest of the other list
	if (i == iLimit) {
		for (; j < jLimit; j++) {
			mergedList->push_back(new Location(*(otherList->at(j))));
		}
	} else if (j == jLimit) {
		for (; i < iLimit; i++) {
			mergedList->push_back(regionList->at(i));
		}
	}

	// Once done
	// Util::deleteInVector(regionList);
	regionList->clear();	// Need to test this line
	delete regionList;
	regionList = mergedList;

	merge();

	//A post-condition: Ensure that the list is sorted
	for (int h = 1; h < regionList->size(); h++) {
		if (regionList->at(h)->getStart() < regionList->at(h - 1)->getStart()) {
			throw InvalidStateException(string("This list is not sorted."));
		}
	}
}

void LocationList::print() {
	cout << endl << chromName << endl;
	for (int i = 0; i < regionList->size(); i++) {
		int s = regionList->at(i)->getStart();
		int e = regionList->at(i)->getEnd();
		cout << s << "-" << e << endl;
	}
}

const vector<ILocation*> * LocationList::getList() {
	return regionList;
}

void LocationList::convertToRedFormat() {
	trim(1);
}

void LocationList::trim(int x) {
	for (int i = regionList->size() - 1; i >= 0; i--) {
		ILocation * region = regionList->at(i);
		int start = region->getStart();
		int newEnd = region->getEnd() - x;

		if (newEnd < 0 || start > newEnd) {
			regionList->erase(regionList->begin() + i);
			delete region;
		} else {
			region->setEnd(newEnd);
		}
	}
}

}

/* namespace nonltr */
```
