## Supplementary Data 1 for "FASTCAR: Rapid alignment-free prediction of sequence alignment identity scores using self-supervised general linear models": LocationList.h

```
/*
 * LocationList.h
 *
 *  Created on: Feb 19, 2015
 *      Author: Hani Z. Girgis, PhD
 */

#ifndef SRC_NONLTR_LOCATIONLIST_H_
#define SRC_NONLTR_LOCATIONLIST_H_

#include <vector>
#include "../utility/Util.h"
#include "../utility/ILocation.h"
#include "../utility/Location.h"
#include "../exception/InvalidStateException.h"

using namespace std;
using namespace utility;
using namespace exception;

namespace nonltr {

class LocationList {
private:
	string chromName;
	vector<ILocation *> * regionList;
	void merge();

public:
	LocationList(string);
	virtual ~LocationList();

	void add(int, int);

	/**
	 * Take a sorted list
	 */
	void mergeWithAnotherList(const vector<ILocation *> * const);


	/**
	 * Print locations
	 */
	void print();

	const vector<ILocation*> * getList();
	void convertToRedFormat();
	void trim(int );
};

} /* namespace nonltr */

#endif /* SRC_NONLTR_LOCATIONLIST_H_ */
```
