## Supplementary Data 1 for "FASTCAR: Rapid alignment-free prediction of sequence alignment identity scores using self-supervised general linear models": LocationListCollection.cpp

```
/*
 * LocationListCollection.cpp
 *
 *  Created on: Feb 19, 2015
 *      Author: Hani Zakaria Girgis, PhD
 */

#include "LocationListCollection.h"

namespace nonltr {

LocationListCollection::LocationListCollection(string fileNameIn) {
	fileName = fileNameIn;
	collection = new map<string, LocationList *>();
	readCoordinates();
}

LocationListCollection::~LocationListCollection() {
	collection->clear();
	delete collection;
}

void LocationListCollection::readCoordinates() {
	Util::checkFile(fileName);

	ifstream in(fileName.c_str());
	LocationList * locList;
	string previousChromName("");

	while (in.good()) {
		string line;
		getline(in, line);

		if (line.compare(string("")) != 0) {
			int colIndex = line.find_last_of(':');
			int dashIndex = line.find_last_of('-');

			string chromName = line.substr(0, colIndex);

			if (previousChromName.compare(chromName) != 0) {

				cout << "Processing regions of " << chromName << endl;

				locList = new LocationList(chromName);
				collection->insert(
						map<string, LocationList *>::value_type(chromName,
								locList));

				previousChromName = chromName;
			}

			int start =
					atoi(
							line.substr(colIndex + 1, dashIndex - colIndex - 1).c_str());
			int end = atoi(line.substr(dashIndex + 1).c_str());
			locList->add(start, end);
		}
	}

	in.close();
}

void LocationListCollection::print() {
	map<string, LocationList *>::iterator itr_s = collection->begin();
	map<string, LocationList *>::iterator itr_e = collection->end();
	while (itr_s != itr_e) {
		collection->at(itr_s->first)->print();
		++itr_s;
	}
}

LocationList * const LocationListCollection::getLocationList(string header) {
	if (collection->count(header) == 0) {
		string msg("Regions of ");
		msg.append(header);
		msg.append(" cannot be found.\n");
		throw InvalidStateException(msg);
	}

	return collection->at(header);
}

void LocationListCollection::convertToRedFormat() {
	map<string, LocationList *>::iterator itr_s = collection->begin();
	map<string, LocationList *>::iterator itr_e = collection->end();
	while (itr_s != itr_e) {
		collection->at(itr_s->first)->convertToRedFormat();
		++itr_s;
	}
}

void LocationListCollection::trim(int x) {
	map<string, LocationList *>::iterator itr_s = collection->begin();
	map<string, LocationList *>::iterator itr_e = collection->end();
	while (itr_s != itr_e) {
		collection->at(itr_s->first)->trim(x);
		++itr_s;
	}
}

} /* namespace nonltr */
```
