## Supplementary Data 1 for "FASTCAR: Rapid alignment-free prediction of sequence alignment identity scores using self-supervised general linear models": LocationListCollection.h

```
/*
 * LocationListCollection.h
 *
 *  Created on: Feb 19, 2015
 *      Author: Hani Zakaria Girgis, PhD
 */

#ifndef SRC_NONLTR_LOCATIONLISTCOLLECTION_H_
#define SRC_NONLTR_LOCATIONLISTCOLLECTION_H_

#include <fstream>
#include <map>

#include "LocationList.h"
#include "../utility/Util.h"
#include "../exception/InvalidStateException.h"

using namespace std;
using namespace utility;

namespace nonltr {

class LocationListCollection {

private:
	string fileName;
	map<string, LocationList *> * collection;
	void readCoordinates();

public:
	LocationListCollection(string);
	virtual ~LocationListCollection();
	LocationList * const getLocationList(string);
	void print();
	void convertToRedFormat();
	void trim(int );
};

} /* namespace nonltr */

#endif /* SRC_NONLTR_LOCATIONLISTCOLLECTION_H_ */
```
