## Supplementary Data 1 for "FASTCAR: Rapid alignment-free prediction of sequence alignment identity scores using self-supervised general linear models": Scanner.cpp

```
/*
 * Scanner.cpp
 *
 *  Created on: Aug 19, 2013
 *      Author: Hani Zakaria Girgis, PhD
 */
#include "Scanner.h"

namespace nonltr {

Scanner::Scanner(HMM * hmmIn, int kIn, ChromosomeOneDigit * chromIn,
		string scoresFile) {
	// ToDo: Fix this operation
	string msg("Scanning file of scores is temporarily disabled.");
	throw InvalidOperationException(msg);

	hmm = hmmIn;
	k = kIn;
	chrom = chromIn;
	segmentList = chrom->getSegment();
	scorer = NULL;
	scoreList = new vector<int>();
	ifstream in(scoresFile.c_str());
	if (in) {
		string header;
		getline(in, header);

		string score;
		while (in >> score) {
			scoreList->push_back(atoi(score.c_str()));
		}
		in.close();
	} else {
		string msg(scoresFile);
		msg.append(" does not exist.");
		throw FileDoesNotExistException(msg);
	}

	regionList = new vector<ILocation *>();

	// Start scanning
	start();
}

Scanner::Scanner(HMM * hmmIn, int kIn, ChromosomeOneDigit * chromIn,
		ITableView<unsigned long, int> * table) {
	hmm = hmmIn;
	k = kIn;

	chrom = chromIn;
	segmentList = chrom->getSegment();
	scorer = new Scorer(chrom, table);
	scorer->takeLog(hmm->getBase());
	scoreList = scorer->getScores();
	regionList = new vector<ILocation *>();

	// Start scanning
	start();
}

Scanner::~Scanner() {
	if (scorer == NULL) {
		scoreList->clear();
		delete scoreList;
	} else {
		delete scorer;
	}

	Util::deleteInVector(regionList);
	delete regionList;
}

void Scanner::start() {
	check();

	decode();

	extendByK();

	merge();
}

void Scanner::check() {
	if (chrom->size() != scoreList->size()) {
		string msg("The size of the sequence is not the same as the size of ");
		msg.append("the scores. The size of sequence is: ");
		msg.append(Util::int2string(chrom->size()));
		msg.append(". The size of the scores is: ");
		msg.append(Util::int2string(scoreList->size()));
		msg.append(".");
		throw InvalidStateException(msg);
	}
}

void Scanner::decode() {
	int segmentCount = segmentList->size();
	for (int tt = 0; tt < segmentCount; tt++) {
		vector<int> * segment = segmentList->at(tt);
		hmm->decode(segment->at(0), segment->at(1), scoreList, *regionList);
	}
}

void Scanner::extendByK() {
	int regionCount = regionList->size();
	if (regionCount > 0) {
		int firstCandIndex = 0;
		int lastCandIndex = 0;
		int segmentNumber = segmentList->size();
		for (int i = 0; i < segmentNumber; i++) {
			vector<int> * s = segmentList->at(i);
			ILocation * c = regionList->at(firstCandIndex);
			// Sometimes a segment have no repeats
			if (Util::isOverlapping(s->at(0), s->at(1), c->getStart(),
					c->getEnd())) {
				lastCandIndex = extendByKHelper(s->at(0), s->at(1),
						firstCandIndex);
				firstCandIndex = lastCandIndex + 1;
				if (firstCandIndex >= regionCount) {
					break;
				}
			}
		}
	}
}

int Scanner::extendByKHelper(int segStart, int segEnd, int firstCandIndex) {
	ILocation * cand = regionList->at(firstCandIndex);

	// Make sure that the first region is overlapping with the segment
	if (!Util::isOverlapping(segStart, segEnd, cand->getStart(),
			cand->getEnd())) {
		string msg("The first region is not overlapping with the segment.");
		msg.append(" Region: ");
		msg.append(Util::int2string(cand->getStart()));
		msg.append(":");
		msg.append(Util::int2string(cand->getEnd()));
		msg.append(" Segment: ");
		msg.append(Util::int2string(segStart));
		msg.append(":");
		msg.append(Util::int2string(segEnd));
		throw InvalidInputException(msg);
	}

	int lastCandIndex = -1;
	int candidateNumber = regionList->size();
	for (int c = firstCandIndex; c < candidateNumber; c++) {
		ILocation * cand = regionList->at(c);
		if (Util::isOverlapping(segStart, segEnd, cand->getStart(),
				cand->getEnd())) {
			int newEnd = cand->getEnd() + k - 1;
			if (newEnd > segEnd) {
				newEnd = segEnd;
			}
			cand->setEnd(newEnd);
			lastCandIndex = c;
		} else {
			break;
		}
	}

	if (lastCandIndex < 0) {
		string msg("The index of the last region cannot be negative.");
		throw InvalidStateException(msg);
	}

	return lastCandIndex;
}

void Scanner::merge() {
	int regionCount = regionList->size();
	int gg = 0;
	while (gg < regionCount) {
		ILocation * region = regionList->at(gg);

		int regionStart = region->getStart();
		int regionEnd = region->getEnd();

		if (gg > 0) {
			ILocation * pRegion = regionList->at(gg - 1);
			int pStart = pRegion->getStart();
			int pEnd = pRegion->getEnd();

			if (Util::isOverlapping(pStart, pEnd, regionStart, regionEnd)) {
				pRegion->setEnd(regionEnd > pEnd ? regionEnd : pEnd);
				regionList->erase(regionList->begin() + gg);
				delete region;
				regionCount = regionList->size();
			} else {
				gg++;
			}
		}

		if (gg == 0) {
			gg++;
		}
	}
}

void Scanner::mergeWithOtherRegions(const vector<ILocation *> * otherList) {
	vector<ILocation *> * mergedList = new vector<ILocation *>();

	int i = 0;
	int j = 0;
	int iLimit = regionList->size();
	int jLimit = otherList->size();

	// Continue until one list is finished
	while (i < iLimit && j < jLimit) {
		ILocation * iLoc = regionList->at(i);
		ILocation * jLoc = otherList->at(j);

		if (iLoc->getStart() < jLoc->getStart()) {
			mergedList->push_back(iLoc);
			i++;
		} else {
			mergedList->push_back(new Location(*jLoc));
			j++;
		}
	}

	// Once one list is finished, copy the rest of the other list
	if (i == iLimit) {
		for (; j < jLimit; j++) {
			mergedList->push_back(new Location(*(otherList->at(j))));
		}
	} else if (j == jLimit) {
		for (; i < iLimit; i++) {
			mergedList->push_back(regionList->at(i));
		}
	}

	// Once done
	// Util::deleteInVector(regionList);
	// @@ Need to be tested
	regionList->clear();
	delete regionList;
	regionList = mergedList;

	merge();

	//Ensure that the list is sorted
	for (int h = 1; h < regionList->size(); h++) {
		if (regionList->at(h)->getStart() < regionList->at(h - 1)->getStart()) {
			throw InvalidStateException(string("This list is not sorted."));
		}
	}
}

void Scanner::makeForwardCoordinates() {
	int regionNum = regionList->size();
	int lastBase = chrom->size() - 1;

	// Calculate the coordinate on the main strand
	for (int i = 0; i < regionNum; i++) {
		ILocation * oldLoc = regionList->at(i);
		regionList->at(i) = new Location(lastBase - oldLoc->getEnd(),
				lastBase - oldLoc->getStart());
		delete oldLoc;
	}

	// Reverse the regions within the list
	int lastRegion = regionNum - 1;
	int middle = regionNum / 2;
	for (int i = 0; i < middle; i++) {
		ILocation * temp = regionList->at(lastRegion - i);
		regionList->at(lastRegion - i) = regionList->at(i);
		regionList->at(i) = temp;
	}

}

/**
 * Warning: this method prints the logarithm values of the scores
 */
void Scanner::printScores(string outputFile, bool canAppend) {
	cout << "Printing the logarithmic values of the scores ";
	cout << "NOT the original scores." << endl;

	ofstream outScores;
	if (canAppend) {
		outScores.open(outputFile.c_str(), ios::out | ios::app);
	} else {
		outScores.open(outputFile.c_str(), ios::out);
	}

	int step = 50;
	outScores << chrom->getHeader() << endl;
	int len = scoreList->size();
	for (int i = 0; i < len; i = i + step) {
		int e = (i + step - 1 > len - 1) ? len - 1 : i + step - 1;
		for (int k = i; k <= e; k++) {
			outScores << scoreList->at(k) << " ";
		}
		outScores << endl;
	}
	outScores << endl;
	outScores.close();
}

void Scanner::printIndex(string outputFile, bool canAppend, int frmt) {

	if(frmt != FRMT_POS && frmt != FRMT_BED){
		string msg("Unknown output format: ");
		msg.append(Util::int2string(frmt));
		msg.append(". The known formats are: ");
		msg.append(Util::int2string(FRMT_POS));
		msg.append(" and ");
		msg.append(Util::int2string(FRMT_BED));
		msg.append(".");
		throw InvalidInputException(msg);
	}

	ofstream outIndex;
	if (canAppend) {
		outIndex.open(outputFile.c_str(), ios::out | ios::app);
	} else {
		outIndex.open(outputFile.c_str(), ios::out);
	}

	// Write the index of the repeat segment [x,y[
	string header = chrom->getHeader();

	if(frmt == FRMT_POS){
		for (int j = 0; j < regionList->size(); j++) {
			outIndex << header << ":";
			outIndex << ((int) (regionList->at(j)->getStart())) << "-";
			outIndex << ((int) (regionList->at(j)->getEnd() + 1));
			outIndex << endl;
		}
	}else if(frmt == FRMT_BED){
		for (int j = 0; j < regionList->size(); j++) {
			outIndex << header << "\t";
			outIndex << ((int) (regionList->at(j)->getStart())) << "\t";
			outIndex << ((int) (regionList->at(j)->getEnd() + 1));
			outIndex << endl;
		}
	}

	outIndex.close();
}

void Scanner::printMasked(string outputFile, Chromosome& oChrom,
		bool canAppend) {

	string baseCopy = *(oChrom.getBase());
	int regionCount = regionList->size();
	for (int j = 0; j < regionCount; j++) {
		for (int h = regionList->at(j)->getStart();
				h <= regionList->at(j)->getEnd(); h++) {
			baseCopy[h] = tolower(baseCopy[h]);
		}
	}

	ofstream outMask;

	if (canAppend) {
		outMask.open(outputFile.c_str(), ios::out | ios::app);
	} else {
		outMask.open(outputFile.c_str(), ios::out);
	}

	outMask << oChrom.getHeader() << endl;
	int step = 50;
	int len = baseCopy.size();
	for (int i = 0; i < len; i = i + step) {
		int e = (i + step - 1 > len - 1) ? len - 1 : i + step - 1;
		for (int k = i; k <= e; k++) {
			outMask << baseCopy[k];
		}
		outMask << endl;
	}
	outMask.close();
}

const vector<ILocation*>* Scanner::getRegionList() {
	return regionList;
}

} /* namespace nonltr */
```
