## Supplementary Data 1 for "FASTCAR: Rapid alignment-free prediction of sequence alignment identity scores using self-supervised general linear models": Scanner.h

```
/*
 * Scanner.h
 *
 *  Created on: Aug 19, 2013
 *      Author: Hani Zakaria Girgis, PhD
 */

#ifndef SCANNER_H_
#define SCANNER_H_

#include <vector>
#include <iostream>
#include <fstream>

#include "Chromosome.h"
#include "ChromosomeOneDigit.h"
#include "HMM.h"
#include "ITableView.h"
#include "Scorer.h"
#include "../utility/Util.h"
#include "../utility/ILocation.h"
#include "../utility/Location.h"
#include "../exception/InvalidInputException.h"
#include "../exception/InvalidStateException.h"
#include "../exception/FileDoesNotExistException.h"
#include "../exception/InvalidOperationException.h"

using namespace std;
using namespace utility;
using namespace exception;

namespace nonltr {

class Scanner {
private:
	//string chromFile;
	ChromosomeOneDigit * chrom;
	const vector<vector<int> *> * segmentList;
	Scorer * scorer;
	vector<int> * scoreList;
	vector<ILocation *> * regionList;
	int k;
	HMM * hmm;
	// bool isTrainMode;

	// Methods
	void start();
	void check();
	void decode();
	void extendByK();
	int extendByKHelper(int, int, int);
	void merge();

public:
	static const int FRMT_POS = 1;
	static const int FRMT_BED = 2;

	Scanner(HMM *, int, ChromosomeOneDigit *, string);
	Scanner(HMM *, int, ChromosomeOneDigit *, ITableView<unsigned long, int> *);
	virtual ~Scanner();
	void makeForwardCoordinates();

	void printScores(string, bool);
	void printIndex(string, bool, int);
	void printMasked(string, Chromosome&, bool);
	void mergeWithOtherRegions(const vector<ILocation *> *);
	const vector<ILocation*>* getRegionList();
};

} /* namespace nonltr */
#endif /* SCANNER_H_ */
```
