## Supplementary Data 1 for "FASTCAR: Rapid alignment-free prediction of sequence alignment identity scores using self-supervised general linear models": Scorer.cpp

```
/*
 * Scorer.cpp
 *
 *  Created on: Aug 3, 2012
 *      Author: Hani Zakaria Girgis, PhD
 */
#include "Scorer.h"

Scorer::Scorer(ChromosomeOneDigit * chromIn,
		ITableView<unsigned long, int> * const table) {
	chrom = chromIn;
	kmerTable = table;
	scores = new vector<int>(chrom->getBase()->size(), 0);
	k = kmerTable->getK();
	max = -1;
	score();
	calculateMax();
}

Scorer::~Scorer() {
	scores->clear();
	delete scores;
}

/**
 * This method scores each nucleotide in the chromosome.
 * The nucleotides represented by 'N' are assigned zero.
 */
void Scorer::score() {
	const vector<vector<int> *> * segment = chrom->getSegment();
	const char * segBases = chrom->getBase()->c_str();

	for (int s = 0; s < segment->size(); s++) {
		int start = segment->at(s)->at(0);
		int end = segment->at(s)->at(1);
		kmerTable->wholesaleValueOf(segBases, start, end - k + 1, scores,
				start);

		// Handle the last word from end - k + 2 till the end, inclusive.
		for (int i = end - k + 2; i <= end; i++) {
			(*scores)[i] = scores->at(i - 1);
		}
	}
}

/**
 * This method takes the logarithm of the scores according to the base.
 * If the score equals zero, it is left the same.
 */
void Scorer::takeLog(double base) {
	// Handle the case where base is one
	bool isOne = false;
	if (fabs(base - 1.0) < std::numeric_limits<double>::epsilon()) {
		isOne = true;
	}
	double logBase = isOne ? log(1.5) : log(base);

	const vector<vector<int> *> * segment = chrom->getSegment();
	for (int s = 0; s < segment->size(); s++) {
		int start = segment->at(s)->at(0);
		int end = segment->at(s)->at(1);
		for (int h = start; h <= end; h++) {
			int score = scores->at(h);

			if (score != 0) {
				if (!isOne || (isOne && score > 1)) {
					(*scores)[h] = ceil(log(score) / logBase);
				}
			}
		}
	}
}

int Scorer::getK() {
	return k;
}

vector<int>* Scorer::getScores() {
	return scores;
}

void Scorer::printScores(string outputFile, bool canAppend) {
	ofstream outScores;
	if (canAppend) {
		outScores.open(outputFile.c_str(), ios::out | ios::app);
	} else {
		outScores.open(outputFile.c_str(), ios::out);
	}

	int step = 50;
	outScores << chrom->getHeader() << endl;
	int len = scores->size();
	for (int i = 0; i < len; i = i + step) {
		int e = (i + step - 1 > len - 1) ? len - 1 : i + step - 1;
		for (int k = i; k <= e; k++) {
			outScores << scores->at(k) << " ";
		}
		outScores << endl;
	}
	outScores << endl;

	outScores.close();
}

int Scorer::countLessOrEqual(int thr) {
	int count = 0;
	const vector<vector<int> *> * segment = chrom->getSegment();
	for (int s = 0; s < segment->size(); s++) {
		int start = segment->at(s)->at(0);
		int end = segment->at(s)->at(1);
		for (int h = start; h <= end; h++) {
			if (scores->at(h) <= thr) {
				count++;
			}
		}
	}
	return count;
}

void Scorer::calculateMax() {
	const vector<vector<int> *> * segmentList = chrom->getSegment();
	int segmentCount = segmentList->size();
	for (int jj = 0; jj < segmentCount; jj++) {
		vector<int> * segment = segmentList->at(jj);
		int start = segment->at(0);
		int end = segment->at(1);
		for (int ss = start; ss <= end; ss++) {
			int score = scores->at(ss);
			if (score > max) {
				max = score;
			}
		}
	}

	if (max == -1) {
		string msg("Error occurred while finding the maximum score.");
		throw InvalidStateException(msg);
	}
}

int Scorer::getMax() {
	return max;
}
```
