## Supplementary Data 1 for "FASTCAR: Rapid alignment-free prediction of sequence alignment identity scores using self-supervised general linear models": Scorer.h

```
/*
 * Scorer.h
 *
 *  Created on: Aug 3, 2012
 *      Author: Hani Zakaria Girgis, PhD
 */

#ifndef SCORER_H_
#define SCORER_H_

#include <vector>
#include <fstream>
#include <iostream>
#include <math.h>
#include <limits>

#include "ITableView.h"
#include "ChromosomeOneDigit.h"
#include "../utility/Util.h"
#include "../exception/InvalidStateException.h"

using namespace std;
using namespace nonltr;
using namespace utility;
using namespace exception;

namespace nonltr {
class Scorer {
private:
	/* Fields */
	ChromosomeOneDigit * chrom;
	ITableView<unsigned long, int> * kmerTable;
	vector<int> * scores;
	int k;
	int max;

	/* Methods */
	void score();
	void calculateMax();

public:
	/* Methods */
	Scorer(ChromosomeOneDigit *, ITableView<unsigned long, int> *);
	virtual ~Scorer();
	void printScores(string, bool);
	vector<int>* getScores();
	int getK();
	void takeLog(double);
	int countLessOrEqual(int);
	int getMax();
};
}

#endif /* Scorer_H_ */
```
