## Supplementary Data 1 for "FASTCAR: Rapid alignment-free prediction of sequence alignment identity scores using self-supervised general linear models": TableBuilder.cpp

```
/*
 * TableBuilder.cpp
 *
 *  Created on: Jul 31, 2012
 *      Author: Hani Zakaria Girgis, PhD
 */

#include "TableBuilder.h"

TableBuilder::TableBuilder(string dir, int motifSize, int order, int minObs) {
	genomeDir = dir;
	k = motifSize;
	genomeLength = 0;
	// kmerTable = new KmerHashTable(k);
	// kmerTable = new EnrichmentView(k);

	// Whenever you change the template, modify line 50 and 70 and the header file line 35
	kmerTable = new EnrichmentMarkovView<unsigned long, int>(k, order, minObs);

	buildTable();
}

TableBuilder::~TableBuilder() {
	delete kmerTable;
}

void TableBuilder::buildTable() {
	vector<string> * fileList = new vector<string>();
	Util::readChromList(genomeDir, fileList, "fa");

	for (int i = 0; i < fileList->size(); i++) {
		cout << "Counting k-mers in " << fileList->at(i) << " ..." << endl;
		ChromListMaker * maker = new ChromListMaker(fileList->at(i));
		const vector<Chromosome *> * chromList = maker->makeChromOneDigitDnaList();

		for (int h = 0; h < chromList->size(); h++) {
			ChromosomeOneDigit * chrom =
					dynamic_cast<ChromosomeOneDigit *>(chromList->at(h));
			if (chrom) {
				genomeLength += chrom->getEffectiveSize();
				updateTable(chrom);
			} else {
				throw InvalidStateException(string("Dynamic cast failed."));
			}
		}

		delete maker;
	}
	// Check if overflow has occurred
	kmerTable->checkOverflow();

	// View
	// EnrichmentView * view = dynamic_cast<EnrichmentView *>(kmerTable);
	EnrichmentMarkovView<unsigned long, int> * view =
			dynamic_cast<EnrichmentMarkovView<unsigned long, int> *>(kmerTable);

	if (view) {
		view->generateProbapilities();
		view->processTable();
		maxValue = view->getMaxValue();
	} else {
		throw InvalidStateException(string("Dynamic cast failed."));
	}
	cout << "Enrichment view is ready." << endl;

	fileList->clear();
	delete fileList;

	/* If you would like to see the contents of the table.*/
	// kmerTable-> printTable();
}

void TableBuilder::updateTable(ChromosomeOneDigit * chrom) {
	// EnrichmentView * view = dynamic_cast<EnrichmentView *>(kmerTable);
	EnrichmentMarkovView<unsigned long, int> * view =
			dynamic_cast<EnrichmentMarkovView<unsigned long, int> *>(kmerTable);

	const vector<vector<int> *> * segment = chrom->getSegment();
	const char * segBases = chrom->getBase()->c_str();

	for (int s = 0; s < segment->size(); s++) {
		int start = segment->at(s)->at(0);
		int end = segment->at(s)->at(1);
		// cerr << "The segment length is: " << (end-start+1) << endl;

		// Fast, but require some memory proportional to the segment length.
		kmerTable->wholesaleIncrement(segBases, start, end - k + 1);
		if (view) {
			view->count(segBases, start, end);
		} else {
			throw InvalidStateException(string("Dynamic cast failed."));
		}

		// Slow, but memory efficient
		/*
		 vector<int> hashList = vector<int>();
		 kmerTable->hash(segBases, start, end - k + 1, &hashList);

		 for (int i = start; i <= end - k + 1; i++) {
		 kmerTable->increment(segBases, i);
		 }
		 */
	}
}

KmerHashTable<unsigned long, int> * const TableBuilder::getKmerTable() {
	return kmerTable;
}

long TableBuilder::getGenomeLength() {
	if (genomeLength < 0) {
		string msg("The length of the genome cannot be negative.");
		throw InvalidStateException(msg);
	}

	return genomeLength;
}

int TableBuilder::getMaxValue() {
	return maxValue;
}
```
