## Supplementary Data 1 for "FASTCAR: Rapid alignment-free prediction of sequence alignment identity scores using self-supervised general linear models": TableBuilder.h

```
/*
 * TableBuilder.h
 *
 *  Created on: Jul 31, 2012
 *      Author: Hani Zakaria Girgis, PhD - NCBI/NLM/NIH
 */

#ifndef TABLEBUILDER_H_
#define TABLEBUILDER_H_

#include "KmerHashTable.h"
#include "EnrichmentMarkovView.h"
#include "ChromosomeOneDigit.h"
#include "ChromListMaker.h"
#include "IChromosome.h"

#include "../utility/Util.h"
#include "../exception/InvalidStateException.h"

#include <iostream>

using namespace std;
using namespace nonltr;
using namespace utility;
using namespace exception;

namespace nonltr {
class TableBuilder {
private:
	/**
	 * k-mer table
	 */
	KmerHashTable<unsigned long,int> * kmerTable;
	int maxValue;

	/**
	 * Directory including the FASTA files comprising the genome.
	 * These files must have the
	 */
	string genomeDir;

	/**
	 * The size of the motif
	 */
	int k;

	/**
	 * The total length of the whole genome
	 */
	long genomeLength;

	/**
	 * Methods
	 */
	void buildTable();
	void updateTable(ChromosomeOneDigit *);

public:
	TableBuilder(string, int, int, int);
	virtual ~TableBuilder();
	KmerHashTable<unsigned long,int> * const getKmerTable();
	void printTable();
	long getGenomeLength();
	int getMaxValue();
};
}

#endif /* TABLEBUILDER_H_ */
```
