## Supplementary Data 1 for "FASTCAR: Rapid alignment-free prediction of sequence alignment identity scores using self-supervised general linear models": Trainer.cpp

```
/*
 * Trainer.cpp
 *
 *  Created on: Aug 20, 2013
 *      Author: Hani Zakaria Girgis, PhD
 */

#include "Trainer.h"

namespace nonltr {

// Pass the isCND and the isCON parameters

Trainer::Trainer(string genomeDirIn, int orderIn, int kIn, double sIn,
		double tIn, string candidateDirIn, int m) : minObs(m) {
	candidateDir = candidateDirIn;
	canPrintCandidates = true;
	isCND = true;
	isCON = false;
	initialize(genomeDirIn, orderIn, kIn, sIn, tIn);
}

Trainer::Trainer(string genomeDirIn, int orderIn, int kIn, double sIn,
		double tIn, string candidateDirIn, bool isCNDIn, string otherDirIn, int m) : minObs(m) {
	candidateDir = candidateDirIn;
	canPrintCandidates = true;
	isCND = isCNDIn;
	isCON = true;
	otherDir = otherDirIn;
	initialize(genomeDirIn, orderIn, kIn, sIn, tIn);
}

Trainer::Trainer(string genomeDirIn, int orderIn, int kIn, double sIn,
		double tIn, int m) : minObs(m) {
	canPrintCandidates = false;
	isCND = true;
	isCON = false;
	initialize(genomeDirIn, orderIn, kIn, sIn, tIn);
}

Trainer::Trainer(string genomeDirIn, int orderIn, int kIn, double sIn,
		double tIn, bool isCNDIn, string otherDirIn, int m) : minObs(m) {
	canPrintCandidates = false;
	isCND = isCNDIn;
	isCON = true;
	otherDir = otherDirIn;
	initialize(genomeDirIn, orderIn, kIn, sIn, tIn);
}

void Trainer::initialize(string genomeDirIn, int orderIn, int kIn, double sIn,
		double tIn) {

	if (isCND == false && isCON == false) {
		string msg("Training using the candidates or the other repeats is required. ");
		msg.append("Please specify which regions to be used for training. ");
		msg.append("Any of the two sets or a combination of both can be used.");
		throw InvalidStateException(msg);
	}

	genomeDir = genomeDirIn;
	fileList = new vector<string>();
	Util::readChromList(genomeDir, fileList, string("fa"));
	chromCount = fileList->size();
	order = orderIn;
	k = kIn;
	s = sIn;
	t = tIn;
	p = 0.0;
	tDetector = tIn + 0.1;
	max = -1;

	stage1();

	if (isCND) {
		stage2();
	}
	stage3();
}

Trainer::~Trainer() {
	fileList->clear();
	delete fileList;
	delete builder;
	delete hmm;
}

/**
 * Stage 1: Building the table
 */
void Trainer::stage1() {
	cout << endl << endl;
	cout << "Stage 1: Building the table ..." << endl;
	builder = new TableBuilder(genomeDir, k, order, minObs);
	table = builder->getKmerTable();
	genomeLength = builder->getGenomeLength();
	max = builder->getMaxValue();
}

void Trainer::stage2() {
	cout << endl << endl;
	cout << "Stage 2: Calculating the percentage ..." << endl;

	double effectiveSize = 0.0;
	double countLessOrEqual = 0.0;
	for (int i = 0; i < chromCount; i++) {
		cout << "Calculating the percentage in: " << fileList->at(i) << " ...";
		cout << endl;
		ChromListMaker * maker = new ChromListMaker(fileList->at(i));
		const vector<Chromosome *> * chromList = maker->makeChromOneDigitDnaList();

		for (int h = 0; h < chromList->size(); h++) {
			ChromosomeOneDigit * chrom =
					dynamic_cast<ChromosomeOneDigit *>(chromList->at(h));
			Scorer * scorer = new Scorer(chrom, table);

			effectiveSize += chrom->getEffectiveSize();
			countLessOrEqual += scorer->countLessOrEqual(t);

			delete scorer;
		}
		delete maker;
	}

	if (effectiveSize == 0) {
		string msg("The size of the genome cannot be zero.");
		throw InvalidStateException(msg);
	} else {
		p = 100.00 * countLessOrEqual / effectiveSize;
		cout << "The percentage is " << p << endl;
		if (p < 52.5) {
			p = 52.5;
			cout << "The percentage is increased to " << p << endl;
		}
	}
}

/**
 * Stage 3: Training
 */
void Trainer::stage3() {
	cout << endl << endl;
	cout << "Stage 3: Training ..." << endl;

	// Handle the case when the threshold is one.
	bool isOne = false;
	if (fabs(t - 1.0) < std::numeric_limits<double>::epsilon()) {
		isOne = true;
	}
	double hmmBase = isOne ? 1.5 : t;

	// Make a list of candidate HMM
	int stateCount = 2 * (ceil(log(max) / log(hmmBase)) + 1);

	// Initialize the HMM
	hmm = new HMM(hmmBase, stateCount);

	// Start training the models
	for (int i = 0; i < chromCount; i++) {
		cout << "Training on: " << fileList->at(i) << endl;
		// Name of candidates file
		string path(fileList->at(i));
		int slashLastIndex = path.find_last_of(Util::fileSeparator);
		int dotLastIndex = path.find_last_of(".");
		string nickName = path.substr(slashLastIndex + 1, dotLastIndex - slashLastIndex - 1);

		// May or may not be used
		string cndFile = candidateDir + Util::fileSeparator + nickName + ".cnd";

		// Work on the other repeats if desired
		LocationListCollection * otherRegionListCollection;
		bool isConRepAvailable = false;
		if (isCON) {
			string otherFile = otherDir + Util::fileSeparator + nickName + ".rpt";
			ifstream f1(otherFile.c_str());
			if (!f1) {
				string message = string("Warning: ");
				message.append(otherFile);
				message.append(" does not exist. ");
				message.append("Repeats of this sequence will not used for training the HMM.");
				cout << message << endl;
			} else {
				otherRegionListCollection = new LocationListCollection(otherFile);
				otherRegionListCollection->convertToRedFormat();
				otherRegionListCollection->trim(k - 1);

				isConRepAvailable = true;
			}
			f1.close();
		}

		// Read sequences in the file
		ChromListMaker * maker = new ChromListMaker(fileList->at(i));
		const vector<Chromosome *> * chromList = maker->makeChromOneDigitDnaList();

		for (int h = 0; h < chromList->size(); h++) {
			ChromosomeOneDigit * chrom = dynamic_cast<ChromosomeOneDigit *>(chromList->at(h));
			Scorer * scorer = new Scorer(chrom, table);
			vector<int> * scoreList = scorer->getScores();

			// Detect candidates if desired
			ChromDetectorMaxima * detector;
			const vector<ILocation*> * trainingRegionList;
			bool canDeleteDetector = true;
			if (isCND) {
				if (canPrintCandidates) {
					detector = new ChromDetectorMaxima(s, 10, 0, tDetector, p,s, scoreList, chrom);
					if (h > 0) {
						bool canAppend = true;
						detector->printIndex(cndFile, canAppend);
					} else {
						cout << "Printing candidates to: " << cndFile << endl;
						detector->printIndex(cndFile);
					}
				} else {
					detector = new ChromDetectorMaxima(s, 10, 0, tDetector, p, s, scoreList, chrom->getSegment());
				}
				trainingRegionList = detector->getRegionList();


			}

			if (isCON && isConRepAvailable) {
				LocationList * const locList = otherRegionListCollection->getLocationList(chrom->getHeader());
				if (isCND) {
					locList->mergeWithAnotherList(detector->getRegionList());
				}
				trainingRegionList = locList->getList();

			}

			// The candidate regions are already copied to the location list
			if (isCND && isCON && isConRepAvailable) {
				delete detector;
				canDeleteDetector = false;
			}

			// Train the HMM
			if(isCND || (isCON && isConRepAvailable)){

				scorer->takeLog(t);
				scoreList = scorer->getScores();
				hmm->train(scoreList, chrom->getSegment(), trainingRegionList);
			}

			// Free more memory
			if (isCND && canDeleteDetector) {
				delete detector;
			}
			delete scorer;
		}

		if (isCON && isConRepAvailable) {
			delete otherRegionListCollection;
		}
		delete maker;
	}

	// Normalize HMM's once training is finished
	hmm->normalize();
}

void Trainer::printTable(string fileName) {
	table->printTable(fileName);
}

HMM*& Trainer::getHmm() {
	return hmm;
}

KmerHashTable<unsigned long, int> * Trainer::getTable() {
	return table;
}

void Trainer::printHmm(string fileName) {
	hmm->print(fileName);
}

} /* namespace nonltr */
```
