## Supplementary Data 1 for "FASTCAR: Rapid alignment-free prediction of sequence alignment identity scores using self-supervised general linear models": Trainer.h

```
/*
 * Trainer.h
 *
 *  Created on: Aug 20, 2013
 *      Author: Hani Zakaria Girgis, PhD
 */

#ifndef TRAINER_H_
#define TRAINER_H_

#include <string>
#include <vector>
#include <iostream>
#include <limits>
#include <algorithm>

#include "TableBuilder.h"
#include "KmerHashTable.h"
#include "HMM.h"
#include "ChromDetectorMaxima.h"
#include "Scorer.h"
#include "ChromListMaker.h"
#include "LocationListCollection.h"
#include "../utility/Util.h"
#include "../exception/InvalidStateException.h"

using namespace std;
using namespace utility;
using namespace exception;

namespace nonltr {

class Trainer {
private:
	string genomeDir;
	string candidateDir;
	string otherDir;
	bool canPrintCandidates;
	vector<string> * fileList;
	int chromCount;
	int order;
	int k;
	int max; // Maximum score in the entire genome
	double t; // Score threshold
	double tDetector; // threshold for the detector because it uses < not <=;
	double p; // Percentage of scores below the threshold, t, in non-repeats
	//double r;
	double s; // Half width of the mask
	long genomeLength;
	//vector<int> * sampleList;
	TableBuilder * builder;
	KmerHashTable<unsigned long, int> * table;
	HMM * hmm;
	int isCND;
	int isCON;
	// The minimum number of the observed k-mers
	const int minObs;

	void stage1();
	void stage2();
	void stage3();
	//void stage4();

public:
	Trainer(string, int, int, double, double, string, int);
	Trainer(string, int, int, double, double, string, bool, string, int);
	Trainer(string, int, int, double, double, int);
	Trainer(string, int, int, double, double, bool, string, int);

	void initialize(string, int, int, double, double);
	virtual ~Trainer();
	void printTable(string);
	void printHmm(string);
	HMM*& getHmm();
	KmerHashTable<unsigned long, int> * getTable();

};

} /* namespace nonltr */
#endif /* TRAINER_H_ */
```
