## Supplementary Data 1 for "FASTCAR: Rapid alignment-free prediction of sequence alignment identity scores using self-supervised general linear models": RepeatsDetector.cpp

```
//============================================================================
// Name        : RepeatsDetector.cpp
// Author      : Hani Zakaria Girgis, PhD
// Version     :
// Description : Red (RepeatsDetector)
//============================================================================
#include <stdio.h>
#include <stdlib.h>
#include <cstdlib>
#include <iostream>
#include <limits>
#include <cmath>
#include <vector>

#include "nonltr/Trainer.h"
#include "nonltr/KmerHashTable.h"
#include "nonltr/TableBuilder.h"
#include "nonltr/HMM.h"
#include "nonltr/Scanner.h"
#include "nonltr/ChromListMaker.h"
#include "utility/Util.h"

using namespace std;
using namespace nonltr;
using namespace utility;
using namespace exception;

/**
 * Parameters
 */
// Required parameters
const static string LEN_PRM = string("-len"); // k - length of the motif.

// Train and Scan the whole genome
const static string GNM_PRM = string("-gnm"); // Train and scan.
const static string ORD_PRM = string("-ord"); // order of background markov chain.
const static string GAU_PRM = string("-gau"); // Half width of the Gaussian mask.
const static string THR_PRM = string("-thr"); // The threshold part of the definition of non-repeats
const static string MIN_PRM = string("-min"); // The minimum number of observations

// Scan using pre-calculated scores and a trained HMM
const static string HMI_PRM = string("-hmi"); // File including the trained model
const static string SEQ_PRM = string("-seq"); // File including the sequence
const static string SCI_PRM = string("-sci"); // File including the scores of the sequence

// Output options with -gnm only
const static string TBL_PRM = string("-tbl"); // Write the k-mer to the provided file
const static string SCO_PRM = string("-sco"); // Write the scores to the
const static string HMO_PRM = string("-hmo"); // The Markov model is writen to this file.
const static string CND_PRM = string("-cnd"); // Write candidate region to a directory

// Output options with -gnm and -hmm
const static string MSK_PRM = string("-msk"); // Write masked sequence(s) to file or directory
const static string RPT_PRM = string("-rpt"); // Write coordinates to file or directory
const static string DIR_PRM = string("-dir"); // Read additional sequences(.fa) or scores (.sc) under directory
const static string FRM_PRM = string("-frm"); // Format of the ouput

void drive(map<string, string> * const param) {
  // Delete old output files
  if (param->count(MSK_PRM) > 0) {
    if (param->count(GNM_PRM) > 0) {
      cout << "Deleting pre-existing files under " << param->at(MSK_PRM);
      cout << endl;
      Util::deleteFilesUnderDirectory(param->at(MSK_PRM));
    } else if (param->count(HMI_PRM) > 0) {
      cout << "Deleting pre-existing " << param->at(MSK_PRM) << endl;
      Util::deleteFile(param->at(MSK_PRM));
    }
  }

  if (param->count(RPT_PRM) > 0) {
    if (param->count(GNM_PRM) > 0) {
      cout << "Deleting pre-existing files under " << param->at(RPT_PRM);
      cout << endl;
      Util::deleteFilesUnderDirectory(param->at(RPT_PRM));
    } else if (param->count(HMI_PRM) > 0) {
      cout << "Deleting pre-existing " << param->at(RPT_PRM) << endl;
      Util::deleteFile(param->at(RPT_PRM));
    }
  }

  if (param->count(SCO_PRM) > 0 && param->count(GNM_PRM) > 0) {
    cout << "Deleting pre-existing files under " << param->at(SCO_PRM);
    cout << endl;
    Util::deleteFilesUnderDirectory(param->at(SCO_PRM));
  }

  if (param->count(HMO_PRM) > 0 && param->count(GNM_PRM) > 0) {
    cout << "Deleting pre-existing " << param->at(HMO_PRM) << endl;
    Util::deleteFile(param->at(HMO_PRM));
  }

  if (param->count(TBL_PRM) > 0 && param->count(GNM_PRM) > 0) {
    cout << "Deleting pre-existing " << param->at(TBL_PRM) << endl;
    Util::deleteFile(param->at(TBL_PRM));
  }

  // Process the input
  int k = atoi(param->at(LEN_PRM).c_str());

  if (param->count(GNM_PRM) > 0) {
    string genomeDir = param->at(GNM_PRM);
    int order = atoi(param->at(ORD_PRM).c_str());
    double s = atoi(param->at(GAU_PRM).c_str());
    double t = atoi(param->at(THR_PRM).c_str());
    int minObs = atoi(param->at(MIN_PRM).c_str());

    // Adjust the threshold when it is one because of the log base.
    if (((int) t) == 1) {
      t = 1.5;
      cout << "The base of the logarithmic function is adjusted." << endl;
    }


    // This part or the next
    Trainer * trainer;
    if (param->count(CND_PRM) > 0) {
      trainer = new Trainer(genomeDir, order, k, s, t, param->at(CND_PRM), minObs);
    } else {
      trainer = new Trainer(genomeDir, order, k, s, t, minObs);
    }


    if (param->count(TBL_PRM)) {
      cout << "Printing the count of the kmer's to: ";
      cout << param->at(TBL_PRM) << endl;
      trainer->printTable(param->at(TBL_PRM));
    }

    if (param->count(HMO_PRM) > 0) {
      cout << "Printing the HMM to: " << endl;
      cout << param->at(HMO_PRM) << endl;
      trainer->printHmm(param->at(HMO_PRM));
    }

    // Stage 3: Scan
    cout << endl << endl;
    cout << "Stage 4: Scanning ..." << endl;
    vector<string> * fileList = new vector<string>();
    Util::readChromList(genomeDir, fileList, string("fa"));
    if (param->count(DIR_PRM) > 0) {
      Util::readChromList(param->at(DIR_PRM), fileList, string("fa"));
    }

    int chromCount = fileList->size();
    for (int i = 0; i < chromCount; i++) {
      cout << "Scanning: " << fileList->at(i) << endl;

      // Output file name
      string path(fileList->at(i));
      int slashLastIndex = path.find_last_of(Util::fileSeparator);
      int dotLastIndex = path.find_last_of(".");
      string nickName = path.substr(slashLastIndex + 1, dotLastIndex - slashLastIndex - 1);

      // Process each sequence with the ith file
      ChromListMaker * maker = new ChromListMaker(fileList->at(i));
      const vector<Chromosome *> * chromList = maker->makeChromOneDigitDnaList();

      ChromListMaker * oMaker = new ChromListMaker(fileList->at(i));
      const vector<Chromosome *> * oChromList;
      if (param->count(MSK_PRM) > 0) {
	oChromList = oMaker->makeChromList();
      }

      for (int h = 0; h < chromList->size(); h++) {
	ChromosomeOneDigitDna * chrom = dynamic_cast<ChromosomeOneDigitDna *>(chromList->at(h));

	// Scan the forward strand
	Scanner * scanner = new Scanner(trainer->getHmm(), k, chrom,trainer->getTable());

	// Scan the reverse complement
	chrom->makeRC();
	Scanner * scannerRC = new Scanner(trainer->getHmm(), k, chrom, trainer->getTable());
	scannerRC->makeForwardCoordinates();
	scanner->mergeWithOtherRegions(scannerRC->getRegionList());
	delete scannerRC;
	chrom->makeRC();


	// Scan the reverse
	chrom->makeR();
	Scanner * scannerR = new Scanner(trainer->getHmm(), k, chrom, trainer->getTable());
	scannerR->makeForwardCoordinates();
	scanner->mergeWithOtherRegions(scannerR->getRegionList());
	delete scannerR;

	//@@ The chromosome now has the sequence of the reverse strand
	// The actual strand is calculated if the user requested the scores.

	// Print according to the user's requests
	bool canAppend = (h == 0) ? false : true;

	if (param->count(SCO_PRM) > 0) {
	  // Calculate the forward strand from the reverse
	  chrom->makeR();

	  string scoFile = param->at(SCO_PRM) + Util::fileSeparator + nickName + ".scr";
	  if (!canAppend) {
	    cout << "Printing scores to: " << scoFile << endl;
	  }
	  // Make sure to print the original E-values not their logarithm
	  Scorer * scorer = new Scorer(chrom, trainer->getTable());
	  scorer->printScores(scoFile, canAppend);
	  delete scorer;
	}

	if (param->count(RPT_PRM) > 0) {
	  string rptFile = param->at(RPT_PRM) + Util::fileSeparator + nickName + ".rpt";
	  if (!canAppend) {
	    cout << "Printing locations to: " << rptFile << endl;
	  }
	  scanner->printIndex(rptFile, canAppend, atoi(param->at(FRM_PRM).c_str()));
	}

	if (param->count(MSK_PRM) > 0) {
	  string mskFile = param->at(MSK_PRM) + Util::fileSeparator + nickName + ".msk";
	  if (!canAppend) {
	    cout << "Printing masked sequence to: " << mskFile << endl;
	  }
	  Chromosome * oChrom = oChromList->at(h);
	  scanner->printMasked(mskFile, *oChrom, canAppend);
	}

	// Free memory
	delete scanner;
      }

      delete maker;
      delete oMaker;
    }

    // Free memory
    fileList->clear();
    delete fileList;
    delete trainer;
  } else if (param->count(HMI_PRM) > 0) {
    HMM * hmm = new HMM(param->at(HMI_PRM));

    string chromFile = param->at(SEQ_PRM);
    string scoresFile = param->at(SCI_PRM);

    ChromosomeOneDigitDna * chrom = new ChromosomeOneDigitDna(chromFile);
    Scanner * scanner = new Scanner(hmm, k, chrom, scoresFile);

    if (param->count(RPT_PRM) > 0) {
      string rptFile = param->at(RPT_PRM);
      cout << "Printing locations to: " << rptFile << endl;
      scanner->printIndex(rptFile, false, atoi(param->at(FRM_PRM).c_str()));
    }

    if (param->count(MSK_PRM) > 0) {
      string mskFile = param->at(MSK_PRM);
      cout << "Printing masked sequence to: " << mskFile << endl;
      Chromosome oChrom(chromFile);
      scanner->printMasked(mskFile, oChrom, false);
    }

    // Free memory
    delete scanner;
    delete chrom;
    delete hmm;
  }
}

int main(int argc, char * argv[]) {
  cout << endl << endl;
  cout << "This is Red (REpeat Detector) designed and developed by ";
  cout << "Hani Zakaria Girgis, PhD." << endl << endl;

  cout << "Version: 05/22/2015" << endl << endl;

  string message = string("Valid argument pairs:\n");

  message.append("\t-gnm input genome directory, required.\n");
  message.append("\t\tFiles with \".fa\" extension in this directory are used for completing the table of the adjusted counts.\n");
  message.append("\t\tThese Files are scanned for repeats.\n");
  message.append("\t-dir directory including additional input sequences, optional.\n");
  message.append("\t\tFiles with \".fa\" extension in this directory are NOT used for completing the table.\n");
  message.append("\t\tThese Files MUST have different names from those in the genome directory.\n");
  message.append("\t\tThese Files are scanned for repeats.\n");


  message.append("\t-len word length equals k defining the k-mer. The default is floor(log_4(genome size)).\n");
  message.append("\t-ord order of the background Markov chain. The default is floor(k/2)-1.\n");
  message.append("\t-gau half width of the mask. The default is based on the GC content.\n");
  message.append("\t\t20 if the GC content > 33% and < 67%, 40 otherwise.\n");

  message.append("\t-thr the threshold score of the low adjusted scores of non-repeats. The default is 2.\n");
  message.append("\t-min the minimum number of the observed k-mers. The default is 3.\n");
  message.append("\t-tbl file where the table of the adjusted counts is written, optional.\n");
  message.append("\t-sco directory where scores are saved, optional.\n");
  message.append("\t\tScore files have the \".scr\" extension.\n");

  message.append("\t-cnd directory where candidate regions are saved, optional.\n");
  message.append("\t\tCandidates files have the \".cnd\" extension.\n");
  message.append("\t-rpt directory where repeats locations are saved, optional.\n");
  message.append("\t\tRepeats files have the \".rpt\" extension.\n");
  message.append("\t-msk directory where masked sequences are saved, optional.\n");
  message.append("\t\tMasked sequences files have the \".msk\" extension.\n");

  message.append("\t-frm the format of the output: 1 (chrName:start-end) or 2 (chrName\tstart\tend).\n");
  message.append("\t\tThe output format are zero based and the end is exclusive.\n");

  message.append("\t-hmo file where the HMM is saved, optional.\n\n");

  message.append("Examples:\n");
  message.append("\tThe following command runs Red with the defaults and generates the masked sequences.\n");
  message.append("\tRed -gnm genome_directory -msk output_directory\n\n");
  message.append("\tThe following command runs Red with the defaults and generates the masked sequences and the locations of repeats.\n");
  message.append("\tRed -gnm genome_directory -msk output_directory -rpt output_directory\n\n");

  // Table of valid argument pairs
  map<string, string> * validParam = new map<string, string>();
  validParam->insert(map<string, string>::value_type(LEN_PRM, "DUMMY"));
  validParam->insert(map<string, string>::value_type(GNM_PRM, "DUMMY"));
  validParam->insert(map<string, string>::value_type(ORD_PRM, "DUMMY"));
  validParam->insert(map<string, string>::value_type(GAU_PRM, "DUMMY"));
  validParam->insert(map<string, string>::value_type(THR_PRM, "DUMMY"));
  validParam->insert(map<string, string>::value_type(HMI_PRM, "DUMMY"));
  validParam->insert(map<string, string>::value_type(SEQ_PRM, "DUMMY"));
  validParam->insert(map<string, string>::value_type(SCI_PRM, "DUMMY"));
  validParam->insert(map<string, string>::value_type(TBL_PRM, "DUMMY"));
  validParam->insert(map<string, string>::value_type(SCO_PRM, "DUMMY"));
  validParam->insert(map<string, string>::value_type(HMO_PRM, "DUMMY"));
  validParam->insert(map<string, string>::value_type(MSK_PRM, "DUMMY"));
  validParam->insert(map<string, string>::value_type(RPT_PRM, "DUMMY"));
  validParam->insert(map<string, string>::value_type(CND_PRM, "DUMMY"));
  validParam->insert(map<string, string>::value_type(DIR_PRM, "DUMMY"));
  validParam->insert(map<string, string>::value_type(MIN_PRM, "DUMMY"));
  validParam->insert(map<string, string>::value_type(FRM_PRM, "DUMMY"));

  // Make a table of the user provided arguments
  map<string, string> * param = new map<string, string>();
  if (argc > 1 && argc % 2 == 1) {
    for (int i = 1; i < argc - 1; i += 2) {
      if (validParam->count(argv[i]) > 0) {
	param->insert(map<string, string>::value_type(argv[i], argv[i + 1]));
      } else {
	cerr << "Invalid argument: " << argv[i] << " " << argv[i + 1];
	cerr << endl;
	cerr << message << endl;
	return 1;
      }
    }


    // Check if the user provided the essential arguments


    if (param->count(LEN_PRM) == 0) {
      if (param->count(GNM_PRM) > 0) {
	// Calculate the size of the genome
	long genomeLength = 0;
	vector<string> * fileList = new vector<string>();
	Util::readChromList(param->at(GNM_PRM), fileList, "fa");
	cout << "Calculating the length, k, of the k-mer ";
	cout << "based on the input genome ... " << endl;
	for (int i = 0; i < fileList->size(); i++) {
	  ChromListMaker * maker = new ChromListMaker(fileList->at(i));
	  const vector<Chromosome *> * chromList = maker->makeChromList();
	  for (int h = 0; h < chromList->size(); h++) {
	    genomeLength += chromList->at(h)->getEffectiveSize();
	  }
	  delete maker;
	}
	fileList->clear();
	delete fileList;

	double temp = log(genomeLength) / log(4.0);

	int k = floor(temp);
	cout << "The recommended k is " << k << "." << endl;
	if (k > 15) {
	  cout << "Due to a memory constraint, k is set to 15.";
	  cout << endl;
	  k = 15;
	}

	if (k < 12) {
	  cout<< "Due to a statistical consideration, k is set to 12.";
	  cout << endl;
	  k = 12;
	}
	cout << endl;

	string kString = Util::int2string(k);
	param->insert(map<string, string>::value_type(LEN_PRM, kString));

      } else {
	cerr << "The word length is required." << endl;
	cerr << message << endl;
	return 1;
      }
    }

    if(param->count(FRM_PRM) == 0){
      cout << "Using the default output format chrName:start-end" << endl;
      param->insert(map<string, string>::value_type(FRM_PRM, Util::int2string(Scanner::FRMT_POS)));
    } else {
      if (atoi(param->at(FRM_PRM).c_str()) !=  Scanner::FRMT_POS && atoi(param->at(FRM_PRM).c_str()) !=  Scanner::FRMT_BED) {
	cerr << "The output format must be " << Scanner::FRMT_POS << " or ";
	cerr << Scanner::FRMT_BED << ". The format received is " ;
	cerr << param->at(FRM_PRM) << "." << endl;
	return 1;
      }
    }

    if (param->count(GNM_PRM) > 0) {
      Util::checkFile(param->at(GNM_PRM));

      if (param->count(ORD_PRM) == 0) {
	double k = atoi(param->at(LEN_PRM).c_str());
	int o = floor(k / 2.0) - 1;

	cout << "Using the default background order: " << o << ".";
	cout << endl;

	string oString = Util::int2string(o);
	param->insert(map<string, string>::value_type(ORD_PRM, oString));
      }

      if (param->count(THR_PRM) == 0) {
	cout << "Using the default threshold: 2." << endl;
	param->insert(map<string, string>::value_type(THR_PRM, string("2")));
      } else {
	if (atoi(param->at(THR_PRM).c_str()) < 1) {
	  cerr << "The threshold cannot be less than 1.";
	  cerr << endl;
	  cerr << message << endl;
	  return 1;
	}
      }

      if (param->count(MIN_PRM) == 0) {
	cout << "Using the default minimum of the observed count of k-mers: 3." << endl;
	param->insert(map<string, string>::value_type(MIN_PRM, string("3")));
      } else {
	if (atoi(param->at(MIN_PRM).c_str()) < 0) {
	  cerr << "The minimum of the observed count of k-mers cannot be less than 0.";
	  cerr << endl;
	  cerr << message << endl;
	  return 1;
	}
      }

      if (param->count(GAU_PRM) == 0) {
	cout << "Calculating GC content ..." << endl;

	// 1: Count the gc content of the input genome
	long genomeLength = 0;
	long genomeGc = 0;
	vector<string> * fileList = new vector<string>();
	Util::readChromList(param->at(GNM_PRM), fileList, "fa");
	for (int i = 0; i < fileList->size(); i++) {
	  ChromListMaker * maker = new ChromListMaker(fileList->at(i));
	  const vector<Chromosome *> * chromList = maker->makeChromList();

	  for (int h = 0; h < chromList->size(); h++) {
	    genomeGc += chromList->at(h)->getGcContent();
	    genomeLength += chromList->at(h)->getEffectiveSize();
	  }
	  delete maker;
	}
	fileList->clear();
	delete fileList;

	// 2: Calculate the gc content of the input genome
	double gc = 100.00 * genomeGc / genomeLength;
	int w = 20;
	if (gc < 33 || gc > 67) {
	  w = 40;
	}
	cout << "Using the default half width: " << w;
	cout << " based on the GC content of " << gc << endl;
	string wString = Util::int2string(w);
	param->insert(map<string, string>::value_type(GAU_PRM, wString));
      }
    } else if (param->count(HMI_PRM) > 0) {
      Util::checkFile(param->at(HMI_PRM));

      if (param->count(SEQ_PRM) == 0) {
	cerr << "The sequence file is required.";
	cerr << endl;
	cerr << message << endl;
	return 1;
      } else {
	Util::checkFile(param->at(SEQ_PRM));
      }

      if (param->count(SCI_PRM) == 0) {
	cerr << "The scores file is required.";
	cerr << endl;
	cerr << message << endl;
	return 1;
      } else {
	Util::checkFile(param->at(SCI_PRM));
      }

    } else {
      cerr << "A mode is required: training and scanning (-gnm) or ";
      cerr << "scanning only (-hmi)." << endl;
      cerr << message << endl;
      return 1;
    }

    // Check optional parameters
    if (param->count(TBL_PRM) > 0 && param->count(GNM_PRM) == 0) {
      cerr << "Printing the k-mer table is optional with -gnm only.";
      cerr << endl;
      cerr << message << endl;
      return 1;
    }

    if (param->count(HMO_PRM) > 0 && param->count(GNM_PRM) == 0) {
      cerr << "Printing the HMM is optional with -gnm only.";
      cerr << endl;
      cerr << message << endl;
      return 1;
    }

    if (param->count(SCO_PRM) > 0 && param->count(GNM_PRM) == 0) {
      cerr << "Printing the scores is optional with -gnm only.";
      cerr << endl;
      cerr << message << endl;
      return 1;
    } else if (param->count(SCO_PRM) > 0 && param->count(GNM_PRM) > 0) {
      Util::checkFile(param->at(SCO_PRM));
    }


    if (param->count(CND_PRM) > 0 && param->count(GNM_PRM) == 0) {
      cerr << "Printing candidate regions is optional with -gnm only.";
      cerr << endl;
      cerr << message << endl;
      return 1;
    } else if (param->count(CND_PRM) > 0 && param->count(GNM_PRM) > 0) {
      Util::checkFile(param->at(CND_PRM));
    }


    if (param->count(DIR_PRM) > 0 && param->count(GNM_PRM) == 0) {
      cerr << "Processing additional sequences is optional with -gnm only.";
      cerr << endl;
      cerr << message << endl;
      return 1;
    } else if (param->count(DIR_PRM) > 0 && param->count(GNM_PRM) > 0) {
      Util::checkFile(param->at(DIR_PRM));
    }

    if (param->count(MSK_PRM) > 0 && param->count(GNM_PRM) > 0) {
      Util::checkFile(param->at(MSK_PRM));
    }

    if (param->count(RPT_PRM) > 0 && param->count(GNM_PRM) > 0) {
      Util::checkFile(param->at(RPT_PRM));
    }

    // Print out the parameters table
    typedef map<string, string> myMap;
    myMap::iterator sIter = param->begin();
    myMap::iterator eIter = param->end();
    cout << endl << "List of final parameters: " << endl;
    while (sIter != eIter) {
      cout << (*sIter).first << ": " << (*sIter).second << endl;
      sIter++;
    }
    cout << endl;

    // Start!
    drive(param);

    // Clear parameters when done.
    param->clear();
    delete param;
  } else {
    cerr << "Argument pairs of the form: -flag value are required.";
    cerr << endl;
    cerr << message << endl;
  }

  //return EXIT_SUCCESS;
  return 0;
}
```
