## Supplementary Data 1 for "FASTCAR: Rapid alignment-free prediction of sequence alignment identity scores using self-supervised general linear models": Driver.cpp

```
/** * Author: Alex Baumgartner
 * The Bioinformatics Toolsmith Laboratory, the University of Tulsa
 * 5/15/2018
 *
 * Purpose:
 *	The pupose of this module is to serve as an interface with the other
 *	classes in this file
 *
 * Error Codes:
 *	1: Incorrect #of arguments
 *	0: Everything workes fine
 */

#include <iostream>
#include <vector>
#include <string>

#include "HandleSeq.h"

using namespace std;

vector<string> muteSequences;

int main(int numOfArgs, char *args[]) {
	//If the user
	if (numOfArgs != 7) {
		cout << "Error: incorrect number of arguments" << endl
				<< "Should be: Sequence File, Base Percent Mutation, Delta, Total Sequences to Return, Mode, File to Output To"
				<< endl
				<< "Mode: 1 for single only, 2 for Non-single only, 3 for both"
				<< endl;
		exit(1);
	} else {
		int num = 1;
		int baseMute = atoi(args[2]);
		int delta = atoi(args[3]);
		int totalSeq = atoi(args[4]);
		//Used to make results more random
		srand(time(NULL));
		string currSeq;
		ofstream fileOut;
		HandleSeq h(atoi(args[5]), 1);
		pair<vector<string>, vector<string>> myPair = h.parseFile(args[1]);
		vector<string> names = myPair.first;
		vector<string> sequences = myPair.second;
		//File which all synthetic sequences will be written to
		int perDelta = totalSeq / (((2 * delta) + 1) * sequences.size());
		string line = "";
		bool perDeltaZero = false;
		if(perDelta == 0){
			perDeltaZero = true;
		}
		//For every sequence parsed from the inputted file
		fileOut.open(args[6], ofstream::out);
		for (int i = 0; i < sequences.size(); i++) {
			//Display which number sequence is about to be mutated
			cout << num << "/" << sequences.size() << flush << "\r\033[K";
			//This loop will run on each sequence the numbe of times the user
			//defines (number of seq. to generate from each seq.)
			for(int mute = baseMute + delta; mute >= baseMute - delta && mute > 0; mute--){
				if(perDeltaZero && mute == baseMute){
					perDelta = totalSeq;
				}
				for(int n = 0; n < perDelta; n++){
					
					
					auto myPair = h.mutate(sequences.at(i), mute);
					line = ">";
					line.append(to_string(myPair.first));
					line.append("_");
					line.append(to_string(mute));
					line.append("_");
					line.append(names.at(i));
					fileOut << line << endl;
					bool start = false;
					int mutationNum = 0;
					fileOut << myPair.second;
					fileOut << endl;
				//	cout << line << endl;
			//		cout << "Orig. seq length = " << sequences.at(i).length() << endl << mute << endl << mutationNum << endl;
				}
				if(perDeltaZero && mute == baseMute){
					perDelta = 0;
				}
			}
				num++;
		}
		if(!perDeltaZero){
		if(totalSeq % (((2 * delta) + 1) * sequences.size()) != 0){
			for(int n = totalSeq % (((2 * delta) + 1) * sequences.size()); n > 0; n--){
				int index = rand() % sequences.size();
				auto myPair = h.mutate(sequences.at(index), baseMute);
					line = ">";
					line.append(to_string(myPair.first));
					line.append("_");
					line.append(to_string(baseMute));
					line.append("_");
					line.append(names.at(index));
					fileOut << line << endl;
					bool start = false;
					int mutationNum = 0;
					fileOut << myPair.second;
					fileOut << endl;
	//								cout << line << endl;
			//	cout << "Orig. seq length extra= " << sequences.at(index).length() << endl << baseMute << endl << mutationNum << endl;
			}
		}
	}
		fileOut.close();
		std::cout << "Finished " << args[6] << endl;
		exit(0);
	}
}
```
