## Supplementary Data 1 for "FASTCAR: Rapid alignment-free prediction of sequence alignment identity scores using self-supervised general linear models": HandleSeq.cpp

```
/**
 * Author: Alex Baumgartner
 * The Bioinformatics Toolsmith Laboratory, the University of Tulsa
 * 5/15/2018
 *
 * Purpose:
 *	The pupose of this module is to take a sequence and mutate it to returns
 			It also serves as a way to parse a file for all sequences
 */

#include "HandleSeq.h"
#include <omp.h>
// d
HandleSeq::HandleSeq(int m, std::random_device::result_type rnd) {

	mode = m & HandleSeq::BOTH;
	enableTrans = m & HandleSeq::TRANSLOCATION;
	enableRev = m & HandleSeq::REVERSION;
	random = new LCG(rnd);
	// disable = (m & HandleSeq::ATYPICAL) > 0 ? 0 : 1;
}

pair<vector<string>, vector<string>> HandleSeq::parseFile(string fileName) {
	ifstream fileIn;
	//Uses the file the user supplies to take in sequences
	fileIn.open(fileName, ifstream::in);
	if(fileIn.is_open()){
	vector<string> sequences;
	vector<string> names;
	string inString;
	//Boolean to make sure that the first sequence
	//has already been found, prevents a null string being written
	bool foundFirst = false;
	string currentLine;
	while (!fileIn.eof()) {
		getline(fileIn, currentLine);
		//Skip the line if nothing is on it
		if (currentLine.length() == 0) {
			continue;
		}
		//If the line has a '>' symbol, the start of a new sequence
		else if (currentLine.at(0) == '>' && foundFirst) {
			//Push the current saved sequene onto the vector,
			//then reset the strings value
			sequences.push_back(inString);
			names.push_back(currentLine.substr(1, currentLine.find_first_of(' ')));
			inString = "";
		}
		else if(currentLine.at(0) == '>' && !foundFirst){
			foundFirst = true;
			names.push_back(currentLine.substr(1, currentLine.find_first_of(' ')));
		}
		//If this is the first >, set found first to true
		else if (!foundFirst) {
			foundFirst = true;
		}
		//Otherwise, add the current Line to
		//the string of current lines
		else {
			inString = inString + currentLine;
		}
	}
	//Push the last found string on
	//(There is no > at the end of a .fa file)
	sequences.push_back(inString);
	fileIn.close();
	return {names, sequences};
	}
	else{
		cout << "Could not find File" << endl;
		exit(2);
	}
}

pair<float, string> HandleSeq::mutate(string sequence, int muteRate, int split) {
	percMute = muteRate;
	if (muteRate == 0) {
		return std::make_pair(1, sequence);
	}
	auto nucls = countNucl(sequence);
	//Assing the percent of each nucleotide in the sequence
	int percAs = (nucls.at(0) * 100) / sequence.length();
	int percCs = (nucls.at(1) * 100) / sequence.length();
	int percGs = (nucls.at(2) * 100) / sequence.length();
	int percTs = (nucls.at(3) * 100) / sequence.length();
	int percMulti, percSing;
	string * seq = new string(sequence);
	int length = sequence.length();
	//If the user only wants single
	if (mode == 1) {
		percMulti = 0;
		//Allocate all mutations to single
		percSing = percMute;
	}
	//Or if the user only wants non single
	else if (mode == 2) {
		//Allocate all mutations to non-single
		percSing = 0;
		percMulti = percMute;
	}
	//Otherwise, assing a random percentage to both
	else {
		percMulti = split;
//		percMulti = random.randMod<int>(percMute);
		percSing = percMute - percMulti;
	}
	//Define a new multiple mutation
	MultiMute multi(percAs, percCs, percGs, percTs,
			percMulti, enableTrans, enableRev, random->nextRandSeed());
	//Run the multiple mutations,
	//get back its vector of what is valid to mutate and what isn't
	vector<bool> mutes = multi.genMulti(seq);
	uint64_t cnt = 0;
	for (bool b : mutes) {
		cnt += b ? 1 : 0;
	}
	if (mutes.size() != seq->length()) {
		cerr << "mutation size is not matching the multi-sequence" << endl;
		throw 100;
	}
	SingMute sing(percAs, percCs, percGs, percTs,
		      percSing, seq, mutes, random->nextRandSeed());
	float alignmentLength = multi.getAlignmentLength() + sing.getAlignmentLength() + length;
//	cout << "alignLength: " << alignmentLength << endl;
	float IBP = length - multi.getIBP() - sing.getIBP();
//	cout << "ibp: " << IBP << endl;
	float alignment = IBP / alignmentLength;
//	cout << "ratio: size: " << mutes.size() << " expected: " << (float)cnt / mutes.size() << " found: " << ((float)length - multi.getIBP()) / ((float)multi.getAlignmentLength() + length) << " align: " << alignment << endl;
	//assign the sequence to the
	//value that the seq pointer stores to
	//clear the heap
	delete seq;
	//Return the now mutated sequence
	std::string outseq = sing.getSeq();
	return make_pair(alignment, outseq);
}

vector<int> HandleSeq::countNucl(string sequence) {
	int a = 0;
	int c = 0;
	int g = 0;
	int t = 0;
	for (int i = 0; i < sequence.length(); i++) {
		if (sequence.at(i) == 'A') {
			a++;
		} else if (sequence.at(i) == 'C') {
			c++;
		} else if (sequence.at(i) == 'G') {
			g++;
		} else if (sequence.at(i) == 'T') {
			t++;
		}
	}
	vector<int> values;
	values.push_back(a);
	values.push_back(c);
	values.push_back(g);
	values.push_back(t);
	return values;
}
```
