## Supplementary Data 1 for "FASTCAR: Rapid alignment-free prediction of sequence alignment identity scores using self-supervised general linear models": HandleSeq.h

```
/**
 * Author: Alex Baumgartner
 * The Bioinformatics Toolsmith Laboratory, the University of Tulsa
 * 5/15/2018
 *
 * Purpose:
 *	The pupose of this module is to take a sequence and mutate it to returns
 			It also serves as a way to parse a file for all sequences
 */
#ifndef HANDLESEQ_H
#define  HANDLESEQ_H

#include <iostream>
#include <vector>
#include <fstream>
#include <string>
#include "LCG.h"
#include "MultiMute.h"
#include "SingMute.h"

using namespace std;

class HandleSeq {
public:
	// Single — point — mutations only
	static const int SINGLE = (1 << 0);
	// Non-single point mutations only
	static const int NON_SINGLE = (1 << 1);
	// Single and non-single mutations
	static const int BOTH = SINGLE | NON_SINGLE;
// translocations and reversions
	static const int TRANSLOCATION = (1 << 2);
	static const int REVERSION = (1 << 3);
	static const int ATYPICAL = TRANSLOCATION | REVERSION;
	static const int ALL = ATYPICAL | BOTH;
	/*
	 constructor

	 @param:
	 int: the mode of the program
	 				(Single only = 1, nonsingle only = 2, both = 3)
	 */
	HandleSeq(int, std::random_device::result_type seed);
	~HandleSeq() { if (random != NULL) { delete random; }}
	/*
	 returns a vector of all sequences in a file inputted

	 @param:
	 std::string: file name
	 int: the mutation rate

	 @return:
	 std::vector<std::string>: Vector of all found sequences
	 */
	pair<vector<string>, vector<string>> parseFile(string);
	/*
	 Mutates a sequence based on parameters inputted in constructor,
	 	and returns the mutated sequence
	 */
	pair<float, string> mutate(string, int, int);

	uint32_t getSeed() const { return seed; }
private:
	uint32_t seed;
	int mode;
	int percMute;
	bool enableTrans, enableRev;
        LCG *random = NULL;
	/*
	 Counts the nucleotides in a file,
	 	and returns a vector corresponding to their values {A, C, G, T}

	 @param:
	 std::string: the sequences

	 @return:
	 std::vector<int>: vector containing ints of each nucleotide count
	 */
	vector<int> countNucl(string);

};

#endif
```
