## Supplementary Data 1 for "FASTCAR: Rapid alignment-free prediction of sequence alignment identity scores using self-supervised general linear models": LCG.h

```
// -*- C++ -*_
#ifndef LCG_H
#define LCG_H

#include <limits>
#include <algorithm>
#include <stdint.h>
#include <iostream>

class LCG {
public:
	LCG(uint64_t seed_) {
		seed = seed_;
	}

	template<class T>
        T randMod(T max) {
		if (max == 0) {
			return 0;
		} else {
			uint64_t x = random() % max;
			return (T)x;
		}
	}

	uint64_t nextRandSeed() {
		return random();
	}
	double rand_between(double id, double range, double low, double high) {
		uint64_t rnd = random();
		double res = (double)rnd / std::numeric_limits<uint64_t>::max();
		double mn = std::max(id - range, low);
		double mx = std::min(id + range, high);
		return mn + (mx - mn) * res;
	}
	uint64_t random() {
		// MMIX random, from https://en.wikipedia.org/wiki/Linear_congruential_generator#Parameters_in_common_use
		// Should be thread safe
		seed = seed * 6364136223846793005 + 1442695040888963407;
		return seed;
	}
private:
	uint64_t seed;
};
#endif
```
