## Supplementary Data 1 for "FASTCAR: Rapid alignment-free prediction of sequence alignment identity scores using self-supervised general linear models": MultiMute.cpp

```
/**
 * Author: Alex Baumgartner
 * The Bioinformatics Toolsmith Laboratory, the University of Tulsa
 * 5/15/2018
 *
 * Purpose:
 *	The pupose of this module is to perform non single mutations on sequences
 */

#include "MultiMute.h"
#include <sstream>
#include "Random.h"

MultiMute::MultiMute(int a, int c, int g, int t, int alloc, bool enableTrans, bool enableRev, std::random_device::result_type seed) : rng(seed)
{
	percAs = a;
	percCs = c;
	percGs = g;
	percTs = t;
	//Set all sub allocations to 0 if the total allocation is 0
	maxTrans = 0;
	maxInsert = 0;
	maxReverse = 0;
	maxDup = 0;
	maxDel = 0;
	total_alloc = alloc;
//	max_block_size = std::max(2, std::min(50, alloc / 10)); // Max mutation block size
//	cout << "max block size: " << max_block_size << endl;
	if (alloc == 0) {
		return;
	}
	//Arbitrary, if only 1 percent is allocated overall, it is allocated to Insert
	else if (alloc == 1) {
		maxTrans = 0;
		maxReverse = 0;
		maxInsert = 1;
		maxDup = 0;
		maxDel = 0;
	} else if (enableTrans) {
		if (alloc > 1) {
			maxTrans = rng.randMod<int64_t>(alloc);
//			maxTrans = rand() % alloc;
			alloc -= maxTrans;
		}
	} else if (enableRev) {
		if (alloc > 1) {
			maxReverse = rng.randMod<int64_t>(alloc);
//			maxReverse = rand() % alloc;
			alloc -= maxReverse;
		}
	}

	if (alloc > 1) {
		maxDel = rng.randMod<int64_t>(alloc);
//		maxDel = (rand() % alloc);
		alloc -= maxDel;
	}
	if (alloc > 0) {
		maxDup = rng.randMod<int64_t>(alloc);
//		maxDup = rand() % alloc;
		alloc -= maxDup;
	} else {
		maxDup = 0;
		if (alloc == 1) {
			alloc--;
			maxDel++;
		}
	}
	maxInsert = alloc;

//	cout << "Max Rev " << maxReverse << " maxDel " << maxDel << " maxTrans " << maxTrans << " maxInsert " << maxInsert << " maxDup " << maxDup << endl;
}

int MultiMute::getAlignmentLength() {
	return alignmentLength;
}

int MultiMute::getIBP() {
	return IBP;
}

std::string abbreviated_vec(const vector<string>& v)
{
	std::ostringstream oss;
	char last = ' ';
	int block_num = 1;
	for (auto s : v) {
		for (auto c : s) {
			if (c == last) {
				block_num++;
			} else if (last != ' ') {
				oss << block_num << last;
				block_num = 1;
				last = c;
			} else {
				last = c;
			}
		}
	}
	return oss.str();
}

int get_num_blocks(const vector<string>& v)
{
	cout << "Abbrev: " << abbreviated_vec(v) << endl;
	char last = '\0';
	int n_blocks = 0;
	for (auto s : v) {
		for (char c : s) {
			if (c == 'S') {
				last = '\0';
			} else if (c != last) {
				last = c;
				n_blocks++;
			}
		}
	}
	return n_blocks;
}
vector<bool> MultiMute::genMulti(string * sequence)
{
	seq = sequence;
//
	double mut_bp = total_alloc * seq->length() / 100.0;
	max_block_size = std::max((int64_t)2, std::min((int64_t)50, (int64_t)round(mut_bp / 10))); // Max mutation block size
	//Calculate the number of nucleotides allocated to each type of mutation
	maxNonMutations = (int64_t) ((float) ((100 - maxReverse - maxTrans - maxInsert - maxDup - maxDel) / 100.0) * seq->length());
	maxReverse = (int64_t) ((float) (maxReverse / 100.0) * seq->length());
 	maxTrans = (int64_t) ((float) (maxTrans / 100.0) * seq->length());
 	maxInsert = (int64_t) ((float) (maxInsert / 100.0) * seq->length());
 	maxDel = (int64_t) ((float) (maxDel / 100.0) * seq->length());
 	maxDup = (int64_t) ((float) (maxDup / 100.0) * seq->length());

//	cout << "maxNonMutations: " << maxNonMutations << " maxReverse: " << maxReverse << " maxTrans: " << maxTrans << " maxInsert: " << maxInsert << " maxDel: " << maxDel << " maxDup: " << maxDup << endl;
 	//calculate alignment length and identical base pairs
 	alignmentLength = maxInsert + maxDup;
 	IBP = maxDel;
 	//Initialize and size vectors
 	int total = maxNonMutations + (2 * maxReverse) + maxTrans + maxInsert + maxTrans + maxDel + maxDup;
 	insertions = new vector<string>();
 	insertions->reserve(maxTrans + maxInsert);
 	mutationStrings = new vector<string>();
 	mutationStrings->reserve(total);
 	//Push 'S', which means that that is an index that wont be mutated, onto the vector
 	for (int i = 0; i < maxNonMutations; i++) {
 		mutationStrings->push_back("S");
 	}

 	reverse(mutationStrings);
//#pragma omp critical
	{
		insert(mutationStrings);
		translocate(mutationStrings);
		duplicate(mutationStrings);
		deleteNucl(mutationStrings);
	}
    //Make sure no palindromes exist
 	checkForAllPalindromes(mutationStrings);

 	//Generate a char vector from the now shuffled mutations vector
 	auto mutationChars = genCharVector(mutationStrings);
 	getTranslocations(mutationChars);
 	//Performs all mutations on the sequence
 	auto ret = formatString(seq->length() + maxTrans + maxInsert + maxDup, mutationChars);
 	delete mutationStrings;
 	delete mutationChars;
 	delete insertions;
 	return ret;
}

void MultiMute::reverse(vector<string> * toAddTo)
{
	//Keep forming strings until the allocation of reverse is used up
	int64_t size;
	//cout << "maxReverse: " << maxReverse << endl;
	while (maxReverse > 0) {
		//Automatically make it 2 to avoid modulus error
		if (maxReverse == 2) {
			size = 2;
		} else {
			size = rng.randMod<int64_t>(maxReverse - 2) + 2;
//			size = (rand() % (maxReverse - 2)) + 2;
			//Add 1 to size if the remaining reverse allocation would be 1
			if (maxReverse - size == 1) {
				size++;
			}
		}
		//cout << "Reverse size: " << size << endl;
		//Add a string of the randomized size to the vector
		string toAdd(size, 'R');
		toAddTo->push_back(toAdd);
		maxReverse -= size;
	}
}

void MultiMute::translocate(vector<string> * toAddTo)
{
	int size;
	//Keep forming strings until the allocation of Translocate is used up
	while (maxTrans > 0) {
		//Automatically make it 2 to avoid modulus error
		if (maxTrans == 2) {
			size = 2;
		} else {
			size =  rng.randMod<int>(std::min(max_block_size, maxTrans - 2)) + 2;
			//Add 1 to size if the remaining reverse allocation would be 1
			if (maxTrans - size == 1) {
				size++;
			}
		}
		//Add a string of the randomized size to the vector, and an I for where to translocate to
		//cout << "maxTrans=" << maxTrans << " Translocate: " << size << endl;
		string toAdd(size, 'T');
		toAddTo->push_back(toAdd);
		toAddTo->push_back("I");
		maxTrans -= size;
	}
}

void MultiMute::insert(vector<string> * toAddTo)
{
	int size;
	//Keep forming strings until the allocation of insert is used up
	while (maxInsert > 0) {
		//Automatically make it 2 to avoid modulus error
		if (maxInsert == 2) {
			size = 2;
		} else {
//			size = (rand() % (maxInsert - 2)) + 2;
			size = rng.randMod<int>(std::min(max_block_size, maxInsert - 2)) + 2;
			//Add 1 to size if the remaining reverse allocation would be 1
			if (maxInsert - size == 1) {
				size++;
			}
		}
		//	cout << "maxInsert=" << maxInsert << " insert " << size << endl;
		//Add an I for where to insert, and add a generated string to the insetions vector
		toAddTo->push_back("I");
		insertions->push_back(genInsert(size));
		maxInsert -= size;
	}
}

void MultiMute::deleteNucl(vector<string> * toAddTo)
{
	int size;
	//Keep forming strings until the allocation of deletion is used up
	while (maxDel > 0) {
		//Automatically make it 2 to avoid modulus error
		if (maxDel == 2) {
			size = 2;
		} else {
			size = rng.randMod<int>(std::min(max_block_size, maxDel - 2)) + 2;
			//size = (rand() % (maxDel - 2)) + 2;
			//Add 1 to size if the remaining reverse allocation would be 1
			if (maxDel - size == 1) {
				size++;
			}
		}
		//Add a string of X's to show what nucleotides will be deleted
//		cout << "maxDelete=" << maxDel << " delete " << size << endl;
		string toAdd(size, 'X');
		toAddTo->push_back(toAdd);
		maxDel -= size;
	}
}

void MultiMute::duplicate(vector<string> * toAddTo)
{
	int size;
	//Keep forming strings until the allocation of duplicate is used up
	while (maxDup > 0) {
		//Automatically make it 2 to avoid modulus error
		if (maxDup == 2) {
			size = 2;
		} else {
			size = rng.randMod<int>(std::min(max_block_size, maxDup - 2)) + 2;
//			size = (rand() % (maxDup - 2)) + 2;
			//Add 1 to size if the remaining reverse allocation would be 1
			if (maxDup - size == 1) {
				size++;
			}
		}
//		cout << "maxDup=" << maxDup << " duplicate " << size << endl;
		//Add a string of D's for duplicate to the vector
		string toAdd(size, 'D');
		toAddTo->push_back(toAdd);
		maxDup -= size;
	}
}

bool MultiMute::checkPalindrome(int start, int end)
{
	bool equal = false;
	for (; start < end; start++, end--) {
		if (seq->at(start) != seq->at(end)) {
			equal = true;
		}
	}
	return equal;
}

string MultiMute::genInsert(int size)
{
	string toInsert;
	toInsert.reserve(size);
	int value;
	//Keep adding characters based on the original distribution of nucleotides
	for (int i = 0; i < size; i++) {
		value = rng.randMod<int>(percAs + percCs + percGs + percTs);
//		value = rand() % (percAs + percCs + percGs + percTs);
		if (value < percAs) {
			toInsert.push_back('A');
		} else if (value < percAs + percCs) {
			toInsert.push_back('C');
		} else if (value < percAs + percCs + percGs) {
			toInsert.push_back('G');
		} else {
			toInsert.push_back('T');
		}
	}
	return toInsert;
}

vector<bool> MultiMute::formatString(int maxSize, vector<char> * mutationsChars)
{
	string temp;
	temp.reserve(maxSize);
	//vector that stores what indexes have/have not been mutated
	vector<bool> validCharacters;
	validCharacters.reserve(mutationsChars->size() * 2);
	unsigned seed = 0;
    // Use of shuffle to randomize the order
	shuffle(insertions->begin(), insertions->end(), default_random_engine(seed));
	int j = 0;
	int i = 0;
	//Goes through until the end of the sequence or the end of the chars vector is reached (should always be seq first)
	for (; i < seq->length() && j < mutationsChars->size();) {
		//If it is a non-mutation character, simply add the current character, increment both positions
		if (mutationsChars->at(j) == 'S') {
			temp.push_back(seq->at(i));
			i++;
			j++;
			validCharacters.push_back(true);
		}
		//If it is an I, get the next insertion string and append it to the back of the mutaton string, as long as the insertion vector still has stuffing
		else if (mutationsChars->at(j) == 'I') {
			if (insertions->size() > 0) {
				std::string ins = insertions->back();
				temp.append(ins);
				for (char c : ins) {
					validCharacters.push_back(false);
				}
				insertions->pop_back();
			}
			//Increment only the char vector
			j++;
		}
		//For duplications, it will add each charceter, and then read a string of the added characters in the same order
		else if (mutationsChars->at(j) == 'D') {
			string temp2;
			temp2.reserve(seq->length() - i);
			for (; j < mutationsChars->size() && mutationsChars->at(j) == 'D' && i < seq->length(); j++, i++) {
				temp2.push_back(seq->at(i));
				temp.push_back(seq->at(i));
				validCharacters.push_back(false);
				validCharacters.push_back(false);
			}
			//I and J are not incremented because they are incremented in the loop
			temp.append(temp2);
		// } else if (mutationChars->at(j) == 'X') {
		// 	i++;
		// 	j++;
		// }
		//Otherwise, skip over the nuleotide
		} else {
			i++;
			j++;
		}
	}
	//Add any extra insertions of there are any
	if (insertions->size() > 0) {
		for (int k = 0; k < insertions->size(); k++) {
			temp.append(insertions->at(k));
		}
	}
	//Reassign the string pointer
	seq->erase();
	seq->reserve(temp.length());
	seq->append(temp);
	return validCharacters;
}


void MultiMute::getTranslocations(vector<char> * toParseFrom) {
	for (int i = 0, j = 0; i < seq->length() && j < toParseFrom->size();) {
		//If a T is found, the string of nucleotides with corresponding T's is copied and added to the insertion vector
		if (toParseFrom->at(j) == 'T') {
			string temp;
			temp.reserve(seq->length() - i);
			for (;j < toParseFrom->size() && toParseFrom->at(j) == 'T' && i < seq->length(); i++, j++) {
				temp.push_back(seq->at(i));
			}
			insertions->push_back(temp);
		}
		//Skip over the I's
		else if (toParseFrom->at(j) == 'I') {
			j++;
		}
		//Otherwise, increment both
		else {
			j++;
			i++;
		}
	}
}

vector<char> * MultiMute::genCharVector(vector<string> * toParseFrom) {
	vector<char> * charVector = new vector<char>();
	charVector->reserve(seq->length());
	string temp;
	//For every index
	for (int i = 0; i < toParseFrom->size(); i++) {
		temp = toParseFrom->at(i);
		//Add each character in the string at the index, add it to the new character vector
		for (int j = 0; j < temp.length(); j++) {
			charVector->push_back(temp.at(j));
		}
	}
	return charVector;
}

void MultiMute::checkForAllPalindromes(vector<string> * toParseFrom) {
	int insertionChanges = 0;
	for (int i = 0, j = 0; i < seq->length() && j < toParseFrom->size();) {
		//If it is not a reversal
		if (toParseFrom->at(j).at(0) != 'R') {
			//If it is an insertion character, only increment the vector integer
			if (toParseFrom->at(j).at(0) == 'I') {
				j++;
			}
			//Otherwise, increment the string iterator by the length of the current string in the vector,
			//then increment the vector integer
			else {
				i += toParseFrom->at(j).length();
				j++;

			}
		} else {
			//If it is not a palindrome, incremtn as in the if statement
			if (checkPalindrome(i, i + toParseFrom->at(j).length() - 1)) {
				i += toParseFrom->at(j).length();
				j++;
			}
			//Otherwise, replace the reverse with a transversal
			else {
				string temp(toParseFrom->at(j).length(), 'T');
				toParseFrom->at(j) = temp;
				insertionChanges++;
			}
		}
	}
	//Insert enough I's randomly for the amount of transversals that replaced reversals
	for (int i = 0; i < insertionChanges; i++) {
		int index = rng.randMod(toParseFrom->size());
//		int index = rand() % toParseFrom->size();
		toParseFrom->insert(toParseFrom->begin() + index, "I");
	}
}
```
