## Supplementary Data 1 for "FASTCAR: Rapid alignment-free prediction of sequence alignment identity scores using self-supervised general linear models": MultiMute.h

```
/**
 * Author: Alex Baumgartner
 * The Bioinformatics Toolsmith Laboratory, the University of Tulsa
 * 5/15/2018
 *
 * Purpose:
 *	The pupose of this module is to perform non single mutations on sequences
 */

#ifndef MULTIMUTE_H
#define  MULTIMUTE_H

#include <iostream>
#include <vector>
#include <string>
#include <algorithm>
#include <random>
#include "Random.h"
#include "LCG.h"
using namespace std;

class MultiMute {
public:
	/*
	 Constructor, creates values
	 	and assignes allocations based on inputted data

	 @param:
	 int: percentage of A's
	 int: percentage of C's
	 int: percentage of G's
	 int: percentage of T's
	 int: The total allocation for non-single mutations
	 int: bool to exclude Translocate and reverse, 1 for disable, any other umber for include
	 */
	MultiMute(int, int, int, int, int, bool, bool, std::random_device::result_type);
	/*
	 Takes in a string pointer,
	 	and mutates it based on the allocation given to the constructor.
	 	Returns a vector of all valid and invalid indexes

	 @param:
	 std::string *: pointer to the string to be mutated

	 @return:
	 std::vector<bool>: vector of mutations,
	 											false means that index has been mutated
	 */
	std::vector<bool> genMulti(std::string *);
	int getAlignmentLength();
	int getIBP();

	private:
	int percAs;
	int percCs;
	int percGs;
	int percTs;
	int64_t maxReverse;
	int64_t maxInsert;
	int64_t maxTrans;
	int64_t maxDel;
	int64_t maxDup;
	int64_t maxNonMutations;
	int64_t alignmentLength;
	int64_t IBP;
	int64_t total_alloc;
	LCG rng;

	int64_t max_block_size;
	std::vector<std::string> * insertions;
	std::vector<string> * mutationStrings;
	std::string * seq;
	/*
	 Takes in a vector
	 */
	void reverse(vector<string> *);
	/*
	 Translocates a random, nonmutaded part of the sequence,
	 	no larger than its max allocation
	 */
	void translocate(vector<string> *);
	/*
	 Inserts at random, nonmutaded part of the sequence,
	 	no larger than its max allocation
	 */
	void insert(vector<string> *);
	/*
	 Deletes a random, nonmutaded part of the sequence,
	 	no larger than its max allocation
	 */
	void deleteNucl(vector<string> *);
	/*
	 Duplicates a random, nonmutaded part of the sequence,
	 	no larger than its max allocation
	 	to an index directly after the duplicated string
	 */
	void duplicate(vector<string> *);
	/*
	 Checks inclusively, [first, last], if a portion is valid

	 @param:
	 int: The starting index (first)
	 int: The ending index (last)

	 @return:
	 bool: true if all indexes in range are valid
	 */
	bool checkPalindrome(int, int);
	void checkForAllPalindromes(vector<string> *);
	/*
	 Marks all indexes in the range as invalid

	 @param:
	 int: first index to be marked false
	 int: last index tobe marked false
	 */
	vector<bool> formatString(int, vector<char> *);

	/*
		Generates a randomized string based on the inputed size
		@param:
		int: size of string to generate
		@return
		string: randomized string
	*/
	std::string genInsert(int);
	/*
		Adds all translocations to the insertions array
		@param:
		vector<char> *: pointer to a char vector with mutation characters
	*/
	void getTranslocations(vector<char> *);
	/*
		converts a vector of strings into a vector of chars
		@param:
		vector<string> *: the vector to be converted
		@return:
		vector<char> *: the vector of characters
	*/
	vector<char> * genCharVector(vector<string> *);
};
#endif
```
