## Supplementary Data 1 for "FASTCAR: Rapid alignment-free prediction of sequence alignment identity scores using self-supervised general linear models": Predictor.cpp

```
/* -*- C++ -*-
 *
 * Predictor.cpp
 *
 * Author: Benjamin T James
 *
 * Predictor implementation class
 * train(vector<>...) is entry point, generates "semi-synthetic" sequences
 * train() actually trains applicable GLM's.
 * close() and similarity() are callable once trained
 */
#include "Predictor.h"
#include "../cluster/src/Loader.h"
#include "../cluster/src/Matrix.h"
#include "HandleSeq.h"
#include "../cluster/src/Progress.h"
#include "Random.h"
#include <algorithm>
#include <iomanip>

template<class T>
void Predictor<T>::save(std::string file, std::string datatype)
{
	std::ofstream out(file);
	out << "k: " << k << endl;
	out << "mode: " << (unsigned int)mode << endl;
	out << "max_features: " << max_num_feat << endl;
	out << "ID: " << id << endl;
	out << "Datatype: " << datatype << endl;
	out << "feature_set: " << feats64 << endl;
	if (mode & PRED_MODE_CLASS) {
		write_to(out, feat_c, c_glm);
	}
	if (mode & PRED_MODE_REGR) {
		write_to(out, feat_r, r_glm);
	}

}

template<class T>
Predictor<T>::Predictor(const std::string filename)
{
	std::ifstream in(filename);
	std::string buf;
	unsigned mode_ = 0;
	in >> buf >> k;
	//cout << buf << k << endl;
	in >> buf >> mode_;
	mode = mode_;
//	cout << buf << mode << endl;
	in >> buf >> max_num_feat;
//	cout << buf << max_num_feat << endl;
	in >> buf >> id;
//	cout << buf << id << endl;
	in >> buf >> datatype;
//	cout << buf << datatype << endl;
	in >> buf >> feats64;
//	cout << buf << feats64 << endl;

	is_trained = true;
	is_training = false;
	if (mode & PRED_MODE_CLASS) {
		auto pr = read_from(in, k);
		c_glm = pr.first;
		feat_c = pr.second;
	}
	if (mode & PRED_MODE_REGR) {
		auto pr = read_from(in, k);
		r_glm = pr.first;
		feat_r = pr.second;
	}
}

template<class T>
void Predictor<T>::write_to(std::ofstream &out, Feature<T>* feat, matrix::GLM glm)
{
	auto combos = feat->get_combos();
	auto lookup = feat->get_lookup();
	auto mins = feat->get_mins();
	auto maxs = feat->get_maxs();
	out << std::endl << "n_combos: " << combos.size() << std::endl;
	out << std::setprecision(std::numeric_limits<double>::digits10) << glm.get_weights().get(0, 0) << endl;
	for (int j = 0; j < combos.size(); j++) {
		auto cmb = combos[j];
		unsigned int val = 0;
		uint64_t flags = 0;
		for (auto i : cmb.second) {
			flags |= lookup[i];
		}
		switch (cmb.first) {
		case Combo::xy:
			val = 0;
			break;
		case Combo::xy2:
			val = 1;
			break;
		case Combo::x2y:
			val = 2;
			break;
		case Combo::x2y2:
			val = 3;
			break;
		}
		out << val << " ";
		out << flags << " ";
		out << std::setprecision(std::numeric_limits<double>::digits10) << glm.get_weights().get(j+1, 0) << std::endl;
	}
	out << std::endl << "n_singles: " << lookup.size() << std::endl;
	for (int j = 0; j < lookup.size(); j++) {
		out << lookup[j] << " ";
		out << std::setprecision(std::numeric_limits<double>::digits10) << mins[j] << " ";
		out << std::setprecision(std::numeric_limits<double>::digits10) << maxs[j] << std::endl;
	}
}


template<class T>
pair<matrix::GLM, Feature<T>*> Predictor<T>::read_from(std::ifstream& in, int k_)
{
	matrix::GLM glm;
	int c_num_raw_feat, c_num_combos;
	Feature<T> *feat = new Feature<T>(k_);
	std::string buf;
	in >> buf >> c_num_combos;
//	cout << buf << "\"" << c_num_combos << "\"" << endl;
	matrix::Matrix weights(c_num_combos+1, 1);
	double d_;
	in >> d_;
	weights.set(0, 0, d_);
	for (int i = 0; i < c_num_combos; i++) {
		int cmb;
		in >> cmb;
		//	cout << (int)cmb << endl;
		uint64_t flags;
		in >> flags;
//		cout << flags << endl;
		double d;
		in >> d;
//		cout << "[" << 0 << "," << i << "] " << d << endl;
		weights.set(i+1, 0, d);//push_back(d);
		Combo cmb_ = Combo::xy;
		switch (cmb) {
		case 0:
			cmb_ = Combo::xy;
			break;
		case 1:
			cmb_ = Combo::xy2;
			break;
		case 2:
			cmb_ = Combo::x2y;
			break;
		case 3:
			cmb_ = Combo::x2y2;
			break;
		default:
			cerr << "error reading weights file" << endl;
			break;
		}
		feat->add_feature(flags, cmb_);
	}

	in >> buf >> c_num_raw_feat;
//	cout << buf << "\"" << c_num_raw_feat << "\"" << endl;
	for (int i = 0; i < c_num_raw_feat; i++) {
		uint64_t single_flag;
		double min_, max_;
		in >> single_flag;
//		cout << single_flag << endl;
		in >> min_;
//		cout << min_ << endl;
		in >> max_;
//		cout << max_ << endl;
		feat->set_normal(single_flag, min_, max_);
	}
	feat->finalize();
	glm.load(weights);
	return {glm, feat};
}

template<class T>
void Predictor<T>::add_feats(std::vector<std::pair<uint64_t, Combo> >& vec, uint64_t feat_flags)
{
	for (uint64_t i = 1; i <= feat_flags; i *= 2) {
		if ((i & feat_flags) == 0) {
			continue;
		}
		for (uint64_t j = 1; j <= i; j *= 2) {
			if ((j & feat_flags) == 0) {
				continue;
			}
			vec.emplace_back(i | j, Combo::xy);
			vec.emplace_back(i | j, Combo::x2y2);
			if (i != j) {
				vec.emplace_back(i | j, Combo::x2y);
				vec.emplace_back(i | j, Combo::xy2);
			}
		}
	}
}
template<class T>
void Predictor<T>::check()
{
	// if (!is_trained && training.size() >= threshold && !is_training) {
	// 	omp_set_lock(&lock);
	// 	is_training = true;
	// 	train();
	// 	is_training = false;
	// 	omp_unset_lock(&lock);
	// }
}
template<class T>
double Predictor<T>::similarity(Point<T>* a, Point<T>* b)
{
	if (!is_trained) {
//		double d = Selector<T>::align(a, b);
		cerr << "alignment: we don't do that here" << endl;
		throw "Bad";
		//		return d;
		// if (!is_training) {
		// 	omp_set_lock(&lock);
		// 	if (training.size() < testing.size() && training.size() < threshold) {
		// 		training.push_back(pra<T>(a, b, d));
		// 	} else if (training.size() >= testing.size() && testing.size() < threshold) {
		// 		testing.push_back(pra<T>(a, b, d));
		// 	}
		// 	omp_unset_lock(&lock);
		// }
		return 0;

	} else {
		return predict(a, b);
	}
}

template<class T>
bool Predictor<T>::close(Point<T> *a, Point<T> *b)
{
	if (!is_trained) {
//		double d = Selector<T>::align(a, b);
		cerr << "alignment shouldn't be used here" << endl;
		throw "bad";
		// if (!is_training) {
		// 	omp_set_lock(&lock);
		// 	if (training.size() < testing.size() && training.size() < threshold) {
		// 		training.push_back(pra<T>(a, b, d));
		// 	} else if (training.size() >= testing.size() && testing.size() < threshold) {
		// 		testing.push_back(pra<T>(a, b, d));
		// 	}
		// 	omp_unset_lock(&lock);
		// }
//		return d > id;
		return false;
	}
	bool val = p_close(a, b);
	if ((mode & PRED_MODE_REGR) && val) {
		// val = p_predict(a, b) > id;
		// if (!val) {
		// 	cout << "FIXED" << endl;
		// }
	}
	return val;
}

template<class T>
double Predictor<T>::p_predict(Point<T>* a, Point<T>* b)
{
	auto cache = feat_r->compute(*a, *b);
	auto weights = r_glm.get_weights();
	double sum = weights.get(0, 0);
	for (int col = 0; col < feat_r->size(); col++) {
		double val = (*feat_r)(col, cache);
		sum += weights.get(col+1, 0) * val;
	}
//	sum = scale_min + (scale_max - scale_min) * sum;
	if (sum < 0) {
		sum = 0;
	} else if (sum > 1) {
		sum = 1;
	}
	return sum;
}
template<class T>
double Predictor<T>::predict(Point<T>* a, Point<T>* b)
{
	return p_predict(a, b);
}

template<class T>
bool Predictor<T>::p_close(Point<T>* a, Point<T>* b)
{
	auto weights = c_glm.get_weights();
	double sum = weights.get(0, 0);
	auto cache = feat_c->compute(*a, *b);
	for (int col = 1; col < weights.getNumRow(); col++) {
		double d = (*feat_c)(col-1, cache);
		sum += weights.get(col, 0) * d;
	}
	return round(c_glm.logistic(sum) - 0.10) > 0;
}


template<class T>
std::pair<matrix::Matrix,matrix::Matrix> generate_feat_mat(const vector<pra<T> > &data, Feature<T>& feat, double cutoff, bool do_print=false)//bool classify, double cutoff, double smin, double smax)
{
	bool classify = (cutoff > 0);
	int nrows = data.size();
	int ncols = feat.size()+1;
	matrix::Matrix feat_mat(nrows, ncols);
	matrix::Matrix labels(nrows, 1);
	#pragma omp parallel for
	for (int row = 0; row < data.size(); row++) {
		auto kv = data.at(row);
		vector<double> cache;
 		// #pragma omp critical
		// {
			cache = feat.compute(*kv.first, *kv.second);
		// }
		feat_mat.set(row, 0, 1);
		if (classify) {
			labels.set(row, 0, kv.val >= cutoff ? 1 : -1);
		} else {
			labels.set(row, 0, kv.val);
			//	labels.set(row, 0, (kv.val - smin) / (smax - smin));
		}
		for (int col = 1; col < ncols; col++) {
			double val = feat(col-1, cache);
			feat_mat.set(row, col, val);
		}
	}
	if (do_print) {
		for (int row = 0; row < data.size(); row++) {
			cout << "FM " << labels.get(row, 0) << " ";
			for (int col = 0; col < ncols; col++) {
				auto val = feat_mat.get(row, col);
				cout << val << " ";
			}
			cout << endl;
		}
		cout << endl;
	}
	return std::make_pair(feat_mat, labels);
}

std::string bin2acgt(const std::string& input)
{
	std::string out = "";
	for (char c : input) {
		switch (c) {
		case 0:
			out += 'A';
			break;
		case 1:
			out += 'C';
			break;
		case 2:
			out += 'G';
			break;
		case 3:
			out += 'T';
			break;
		default:
			out += "ERR";
		}
	}
	return out;
}

std::string uniqheader(std::string hdr)
{
	std::string out = "";
	bool reached_space = false;
	for (char c : hdr) {
		if (c == ' ') {
			break;
		}
		out += c;
	}
	auto ptr = hdr.find("_mut");
	if (ptr != std::string::npos) {
		return out + hdr.substr(ptr);
	} else {
		return out;
	}

}
template<class T>
void Predictor<T>::train(const vector<Point<T> *> &points, uintmax_t &_id, size_t num_sample)
{
	if (is_trained) { return; }

	// for (auto p : points) {
	// 	cout << "H: " << p->get_header() << endl;
	// }
	num_sample = min(num_sample, points.size());

	vector<Point<T>*> f_points_tr, f_points_test;
	size_t total_size = points.size();// + queries.size();
	for (int i = 0; i < num_sample; i++) {
		int i1 = floor((double)i * total_size / (2 * num_sample));
		int i2 = floor((i + 1) * (double)total_size / (2 * num_sample));
		f_points_tr.push_back(points.at(i1));
		f_points_test.push_back(points.at(i2));
	}
	// size_t q_sample = min(num_sample / 10, queries.size());
	// while (10 * f_points_tr.size() <= 11 * num_sample) {
	// 	for (int i = 0; i < q_sample; i++) {
	// 		int i1 = floor((double)i * queries.size() / (2 * q_sample));
	// 		int i2 = floor((i + 1) * (double)queries.size() / (2 * q_sample));
	// 		f_points_tr.push_back(queries.at(i1));
	// 		f_points_test.push_back(queries.at(i2));
	// 	}
	// }
	training.clear();
	testing.clear();
	if (mode & PRED_MODE_CLASS) {
		vector<std::random_device::result_type> train_seeds, test_seeds;
		for (size_t i = 0; i < f_points_tr.size(); i++) {
			train_seeds.push_back(random.nextRandSeed());
		}
		for (size_t i = 0; i < f_points_test.size(); i++) {
			test_seeds.push_back(random.nextRandSeed());
		}
		std::vector<pra<T> > pos_buf, neg_buf;
		cout << "mutating sequences" << endl;
		size_t counter = 0;
		// struct timespec start, stop;
		// clock_gettime(CLOCK_MONOTONIC, &start);
		Progress prog1(f_points_tr.size(), "Generating training");
//		#pragma omp parallel for
		for (size_t i = 0; i < f_points_tr.size(); i++) {
			auto p = f_points_tr[i];
			mutate_seqs(p, 5, pos_buf, neg_buf, 100 * id, 100, _id, train_seeds[i]);
			mutate_seqs(p, 10, pos_buf, neg_buf, min_id, 100 * id, _id, train_seeds[i]);
			#pragma omp critical
			prog1++;
		}
		prog1.end();

		// clock_gettime(CLOCK_MONOTONIC, &stop);
		// printf("took %lu\n", stop.tv_sec - start.tv_sec);

		counter = 0;
		size_t buf_size = std::min(pos_buf.size(), neg_buf.size());
		cout << "training +: " << pos_buf.size() << endl;
		cout << "training -: " << neg_buf.size() << endl;
		auto pra_cmp = [](const pra<T> &a, const pra<T> &b) {
			int fc = a.first->get_header().compare(b.first->get_header());
			int sc = a.second->get_header().compare(b.second->get_header());
			return fc < 0 || (fc == 0 && sc < 0);
		};
		std::sort(pos_buf.begin(), pos_buf.end(), pra_cmp);
		std::sort(neg_buf.begin(), neg_buf.end(), pra_cmp);

		std::shuffle(pos_buf.begin(), pos_buf.end(), random.gen());
		std::shuffle(neg_buf.begin(), neg_buf.end(), random.gen());
		// for (auto p : pos_buf) {
		// 	cout << "PB: " << p.first->get_header() << " " << p.second->get_header() << endl;
		// }
		// for (auto p : neg_buf) {
		// 	cout << "NB: " << p.first->get_header() << " " << p.second->get_header() << endl;
		// }

		for (size_t i = 0; i < buf_size; i++) {
			training.push_back(pos_buf[i].deep_clone());

		}
		for (size_t i = 0; i < 2 * buf_size && i < neg_buf.size(); i++) {
			training.push_back(neg_buf[i].deep_clone());
		}
		for (auto p : pos_buf) {
			delete p.first;
			delete p.second;
		}
		for (auto p : neg_buf) {
			delete p.first;
			delete p.second;
		}
		pos_buf.clear();
		neg_buf.clear();
		Progress prog2(f_points_test.size(), "Generating testing");
//		#pragma omp parallel for
		for (size_t i = 0; i < f_points_test.size(); i++) {
			auto p = f_points_test[i];
			mutate_seqs(p, 5, pos_buf, neg_buf, 100 * id, 100, _id, test_seeds[i]);
			mutate_seqs(p, 10, pos_buf, neg_buf, min_id, 100 * id, _id, test_seeds[i]);
#pragma omp critical
			prog2++;
		}
		prog2.end();
		buf_size = std::min(pos_buf.size(), neg_buf.size());
		cout << "testing +: " << pos_buf.size() << endl;
		cout << "testing -: " << neg_buf.size() << endl;
		std::sort(pos_buf.begin(), pos_buf.end(), pra_cmp);
		std::sort(neg_buf.begin(), neg_buf.end(), pra_cmp);

		std::shuffle(pos_buf.begin(), pos_buf.end(), random.gen());
		std::shuffle(neg_buf.begin(), neg_buf.end(), random.gen());
		for (size_t i = 0; i < buf_size; i++) {
			testing.push_back(pos_buf[i].deep_clone());
		}
		for (size_t i = 0; i < 2 * buf_size && i < neg_buf.size(); i++) {
			testing.push_back(neg_buf[i].deep_clone());
		}
		for (auto p : pos_buf) {
			delete p.first;
			delete p.second;
		}
		for (auto p : neg_buf) {
			delete p.first;
			delete p.second;
		}
		// for (auto p : training) {
		// 	std::string h1 = uniqheader(p.first->get_header());
		// 	std::string h2 = uniqheader(p.second->get_header());
		// 	cout << "%" << h1 << endl;
		// 	cout << "%" << bin2acgt(p.first->get_data_str()) << endl;
		// 	cout << "!" << h1 << "!" << h2 << "!" << p.val << endl;
		// 	cout << "@" << h2 << endl;
		// 	cout << "@" << p.second->get_data_str() << endl;
		// }

		// for (auto p : testing) {
		// 	std::string h1 = uniqheader(p.first->get_header());
		// 	std::string h2 = uniqheader(p.second->get_header());
		// 	cout << "%" << h1 << endl;
		// 	cout << "%" << bin2acgt(p.first->get_data_str()) << endl;
		// 	cout << "!" << h1 << "!" << h2 << "!" << p.val << endl;
		// 	cout << "@" << h2 << endl;
		// 	cout << "@" << p.second->get_data_str() << endl;
		// }
//		exit(100);
	} else {
		for (auto p : f_points_tr) {
			mutate_seqs(p, 5, training, training, min_id, 100, _id, random.nextRandSeed());
		}
		for (auto p : f_points_test) {
			mutate_seqs(p, 5, testing, testing, min_id, 100, _id, random.nextRandSeed());
		}
	}


	train();
}
template<class T>
std::pair<double, matrix::GLM> regression_train(const vector<pra<T> > &data, Feature<T>& feat)
{
	auto pr = generate_feat_mat(data, feat, -1);
	matrix::GLM glm;
	glm.train(pr.first, pr.second);
	auto result1 = pr.first * glm.get_weights();
	auto diff1 = result1 - pr.second;
	double sum = 0;
	for (int i = 0; i < diff1.getNumRow(); i++) {
		sum += fabs(diff1.get(i, 0));
	}
	sum /= diff1.getNumRow();
	return {sum, glm};
}

template<class T>
std::pair<double, matrix::GLM> class_train(vector<pra<T> > &data, Feature<T>& feat, double cutoff)
{
	// vector<pra<T> > above, below;

	// for (auto d : data) {
	// 	if (d.val > cutoff) {
	// 		above.push_back(d);
	// 	} else {
	// 		below.push_back(d);
	// 	}
	// }
	// size_t sz = std::min(above.size(), below.size());
	// data.clear();
	// for (size_t i = 0; i < sz; i++) {
	// 	data.push_back(above[i]);
	// 	data.push_back(below[i]);
	// }
	auto pr = generate_feat_mat(data, feat, cutoff);
	matrix::GLM glm;
	glm.train(pr.first, pr.second);
	matrix::Matrix p = glm.predict(pr.first);
	for (int row = 0; row < p.getNumRow(); row++) {
		if (p.get(row, 0) == 0) {
			p.set(row, 0, -1);
		}
	}
	auto tup = glm.accuracy(pr.second, p);
	double acc = get<0>(tup);
	double sens = get<1>(tup);
	double spec = get<2>(tup);
	return {acc, glm};
}

template<class T>
double regression_test(const vector<pra<T> >& data, Feature<T>& feat, const matrix::GLM& glm, std::string prefix="")
{
	auto pr = generate_feat_mat(data, feat, -1);
	auto result1 = pr.first * glm.get_weights();
	auto diff1 = result1 - pr.second;
	double sum = 0;
	for (int i = 0; i < diff1.getNumRow(); i++) {
		sum += fabs(diff1.get(i, 0));
	}
	if (prefix != "") {
		for (int row = 0; row < result1.getNumRow(); row++) {
			cout << prefix << ";" << data[row].first->get_header() << ";" << data[row].second->get_header() << ";" << result1.get(row, 0) << ";" << pr.second.get(row, 0) << ";" << diff1.get(row, 0) << endl;
		}
	}
	sum /= diff1.getNumRow();
	return sum;
}

template<class T>
void print_wrong(matrix::Matrix oLabels, matrix::Matrix pLabels)
{
	for(int i = 0; i < oLabels.getNumRow(); i++){
	        if(oLabels.get(i,0) == pLabels.get(i, 0)){
			cout << "";
		}
	}
}

template<class T>
tuple<double,double,double> class_test(const vector<pra<T> >& data, Feature<T>& feat, const matrix::GLM& glm, double cutoff, std::string prefix="")
{
	auto pr = generate_feat_mat(data, feat, cutoff);
	matrix::Matrix p = glm.predict(pr.first);
	for (int row = 0; row < p.getNumRow(); row++) {
		if (p.get(row, 0) == 0) {
			p.set(row, 0, -1);
		}
		if (prefix != "") {
			cout << prefix << ";" << data[row].first->get_header() << ";" << data[row].second->get_header() << ";" << data[row].val << ";" << p.get(row, 0) << ";" << pr.second.get(row, 0) << endl;
		}
	}
//	print_wrong(pr.second, p);
	auto tup = glm.accuracy(pr.second, p);
	return tup;
	// return std::make_tuple(sqrt(get<1>(tup) * get<2>(tup)),
	// 		       get<1>(tup),
	// 		       get<2>(tup));

}

template<class T>
void Predictor<T>::filter(std::vector<pra<T> > &vec, std::string prefix)
{
	std::vector<std::vector<pra<T> > > bins;
	std::vector<double> limits;
	size_t num_bins = 10;
	size_t smallest_bin_size = vec.size();
	for (size_t i = 0; i < num_bins; i++) {
		limits.push_back(id + i * (1 - id) / num_bins);
		bins.push_back(std::vector<pra<T> >());
	}
	limits.push_back(1);
	for (auto p : vec) {
		for (size_t i = 1; i < limits.size(); i++) {
			if (p.val <= limits[i] && p.val > limits[i-1]) {
				bins[i-1].push_back(p);

				break;
			}
		}
	}
	size_t bin_size = 0;
	for (auto &v : bins) {
		bin_size += v.size();
		// smallest_bin_size = std::min(smallest_bin_size, v.size());
		std::shuffle(v.begin(), v.end(), random.gen());
	}
	smallest_bin_size = bin_size / bins.size();
	vec.clear();

	for (auto &v : bins) {
		for (size_t i = 0; i < std::min(v.size(), smallest_bin_size); i++) {
			vec.push_back(v[i]);
			if (prefix != "") {
				cout << prefix << " bin " << i - 1 << " " << v[i].val << endl;
			}
		}
	}
	cout << "new vector size: " << vec.size() << " divided into " << bins.size() << " equal parts" << endl;
}
template<class T>
void Predictor<T>::mutate_seqs(Point<T>* p, size_t num_seq, vector<pra<T> > &pos_buf, vector<pra<T> > &neg_buf, double id_begin, double id_end, uintmax_t& _id, std::random_device::result_type seed)
{

	LCG newRand(seed);
	HandleSeq h(mut_type, newRand.nextRandSeed());

	std::string bin_seq = p->get_data_str();
	std::string seq;
	for (auto c : bin_seq) {
		switch (c) {
		case 0:
			seq += 'A';
			break;
		case 1:
			seq += 'C';
			break;
		case 2:
			seq += 'G';
			break;
		case 3:
			seq += 'T';
			break;
		case 'N':
			seq += 'C';
			break;
		default:
			cout << "Invalid character " << c << endl;
			cout << "from sequence " << bin_seq << endl;
			throw 3;
		}
	}

	double inc = (id_end - id_begin) / num_seq;
	for (size_t i = 0; i < num_seq; i++) {
		double iter_id = id_begin + inc * (i + 0.5);
		double actual_id = newRand.rand_between(iter_id, inc, id_begin, id_end);
//		double actual_id = rand_between(iter_id, inc, id_begin, id_end);
		int mut = round(100 - actual_id);
		mut = (mut == 0) ? 1 : mut;
		int spt = newRand.randMod<int>(mut);
		auto newseq = h.mutate(seq, mut, spt);
		std::string chrom;
		std::ostringstream oss;
		oss << p->get_header() << "_mut" << mut << "_" << spt << "_" << i;
		std::string header = oss.str();
		Point<T>* new_pt = Loader<T>::get_point(header, newseq.second, _id, k);
		pra<T> pr;
		pr.first = p->clone();
		pr.first->set_data_str(bin_seq);
		pr.second = new_pt;
		pr.second->set_data_str(newseq.second);
		pr.val = newseq.first;
#pragma omp critical
		{
			if (pr.val > id) {
				pos_buf.push_back(pr);
			} else {
				neg_buf.push_back(pr);
			}
		}
	}
}
template<class T>
void Predictor<T>::train()
{
	Feature<T> feat(k);
	feat.set_save(true);

	uint64_t max_feat = 0;
	for (uint64_t i = 0; i < possible_feats.size(); i++) {
		if (possible_feats.at(i).first > max_feat) {
			max_feat |= possible_feats.at(i).first;
		}
	}
	for (uint64_t i = 1; i <= max_feat; i *= 2) {
		if (i & max_feat) {
			feat.add_feature(i, Combo::xy);
		}
	}
	feat.normalize(training);
	feat.normalize(testing);
	feat.finalize();


	// cout << "Class Training:" << endl;
	// for (auto p : training) {
	// 	cout << p.val << " ";
	// }
	// cout << "Class Testing:" << endl;
	// for (auto p : testing) {
	// 	cout << p.val << " ";
	// }
	if (mode & PRED_MODE_CLASS) {
		train_class(&feat);
		if (mode & PRED_MODE_REGR) {
			// vector<Point<T>*> f_points_tr, f_points_test;
			// for (int i = 0; i < 10; i++) {
			// 	f_points_tr.push_back(training[rand()%training.size()].first);
			// 	f_points_test.push_back(training[rand()%training.size()].first);
			// }
			// training.clear();
			// testing.clear();
			// for (auto p : f_points_tr) {
			// 	mutate_seqs(p, 50, training, 100 * id, 100);
			// 	mutate_seqs(p, 50, training, 60, 100 * id);
			// }
			// for (auto p : f_points_test) {
			// 	mutate_seqs(p, 50, testing, 100 * id, 100);
			// 	mutate_seqs(p, 50, testing, 60, 100 * id);
			// }
			// filter();
			auto func = [&](pra<T> pr) {
				return pr.val <= id;
			};
			training.erase(std::remove_if(training.begin(), training.end(), func), training.end());
			testing.erase(std::remove_if(testing.begin(), testing.end(), func), testing.end());
			filter(training);//, "training");
			filter(testing);//, "testing");

		}
	}
	if (mode & PRED_MODE_REGR) {
		train_regr(&feat);
	}
	cout << "Training size: " << training.size() << endl;
	cout << "Testing size: " << testing.size() << endl;
	for (auto p : training) {
		delete p.first;
		delete p.second;
	}
	for (auto p : testing) {
		delete p.first;
		delete p.second;
	}
	cout << endl;
	feat.set_save(false);
	training.clear();
	testing.clear();
	possible_feats.clear();
	is_trained = true;
	// save("weights.txt");
	// exit(100);
}

template<class T>
void Predictor<T>::train_class(Feature<T>* feat)
{
	auto c_size = feat->get_combos().size();
	for (int i = 0; i < c_size; i++) {
		feat->remove_feature();
	}
	vector<uintmax_t> used_list;
	double abs_best_acc = 0;
//	cout << "possible feats at one step: " << possible_feats.size() << endl;
	Progress prog(possible_feats.size() * max_num_feat, "Feature selection:");

	std::ostringstream oss;
	for (auto num_feat = 1; num_feat <= max_num_feat; num_feat++) {
		double best_class_acc = abs_best_acc;
		uintmax_t best_idx = -1, cur_idx = 1;
		auto best_class_feat = possible_feats.front();
		for (uint64_t i = 0; i < possible_feats.size(); i++) {
			if (std::find(used_list.begin(), used_list.end(), i) != used_list.end()) {
				continue;
			}
			auto rfeat = possible_feats[i];
		        feat->add_feature(rfeat.first, rfeat.second);
			feat->normalize(training);
			feat->finalize();
			auto name = feat->feat_names().back();
			auto pr = class_train(training, *feat, id);
			auto class_ac = class_test(testing, *feat, pr.second, id);
			double class_accuracy = get<0>(class_ac);//sqrt(get<1>(class_ac) * get<2>(class_ac));
			feat->remove_feature();
			prog++;
//			cout << "Feature: " << cur_idx++ << "/" << possible_feats.size() - used_list.size() << " " << num_feat << "/" << max_num_feat << " " << name  << " acc: " << get<0>(class_ac) << " sens: " << get<1>(class_ac) << " spec: " << get<2>(class_ac) << endl;
			if (class_accuracy > best_class_acc) {
				best_class_acc = class_accuracy;
				best_class_feat = rfeat;
				best_idx = i;
			}
		}
		/* accept the feature if either 1. we don't have enough features
		 * or 2. it improves accuracy by over 0.5%
		 */
		if (best_class_acc > abs_best_acc || num_feat <= min_num_feat) {
			feat->add_feature(best_class_feat.first, best_class_feat.second);
			feat->normalize(training);
			feat->finalize();
			abs_best_acc = best_class_acc;
			used_list.push_back(best_idx);
			oss << "Feature added: " << best_class_feat.first << " " << (int)best_class_feat.second << endl;
			oss << "Accuracy: " << best_class_acc << endl;
			possible_feats.erase(std::remove(possible_feats.begin(), possible_feats.end(), best_class_feat), possible_feats.end());
		}
	}
	prog.end();
	cout << oss.str();
	feat_c = new Feature<T>(*feat);
	feat_c->set_save(false);
	auto pr = class_train(training, *feat_c, id);
	c_glm = pr.second;
	auto train_results = class_test(training, *feat_c, c_glm, id);//, "train");
	cout << "Training ACC: " << get<0>(train_results) << " " << get<1>(train_results) << " " << get<2>(train_results) << endl;
	auto test_results = class_test(testing, *feat_c, c_glm, id);//, "test");
	double class_acc = get<0>(test_results);
	cout << "Testing ACC: " << class_acc << " " << get<1>(test_results) << " " << get<2>(test_results) << endl;

	cout << "Features: "<< endl;
	for (auto line : feat_c->feat_names()) {
		cout << "\t" << line << endl;
	}
}
template<class T>
void Predictor<T>::train_regr(Feature<T>* feat)
{
	auto c_size = feat->get_combos().size();
	for (int i = 0; i < c_size; i++) {
		feat->remove_feature();
	}
	vector<uintmax_t> used_list;
	double abs_best_regr = 1000000;
//	Progress prog(possible_feats.size() * max_num_feat, "Feature selection:");
	for (auto num_feat = 1; num_feat <= max_num_feat; num_feat++) {
		double best_regr_err = abs_best_regr;
		uintmax_t best_idx = -1, cur_idx = 1;
		auto best_regr_feat = possible_feats.front();
		for (uint64_t i = 0; i < possible_feats.size(); i++) {
			if (std::find(used_list.begin(), used_list.end(), i) != used_list.end()) {
				continue;
			}
			auto rfeat = possible_feats[i];
		        feat->add_feature(rfeat.first, rfeat.second);
			feat->normalize(training);
			feat->finalize();
			auto pr = regression_train(training, *feat);
			auto name = feat->feat_names().back();
			double regr_mse = regression_test(testing, *feat, pr.second);
			feat->remove_feature();
			//	prog++;
			//cout << "Feature: " << cur_idx++ << "/" << possible_feats.size() - used_list.size() << " " << num_feat << "/" << max_num_feat << " " << name << " err: " << regr_mse << endl;
			if (regr_mse < best_regr_err) {
				best_regr_err = regr_mse;
				best_regr_feat = rfeat;
				best_idx = i;
			}
		}
		if (best_regr_err < abs_best_regr) {
			feat->add_feature(best_regr_feat.first, best_regr_feat.second);
			feat->normalize(training);
			feat->finalize();
			abs_best_regr = best_regr_err;
			used_list.push_back(best_idx);
			//possible_feats.erase(std::remove(possible_feats.begin(), possible_feats.end(), best_regr_feat), possible_feats.end());
		}
	}
//	prog.end();

	feat_r = new Feature<T>(*feat);
	feat_r->set_save(false);
	auto pr = regression_train(training, *feat_r);
	r_glm = pr.second;
	double tr_regr_mse = regression_test(testing, *feat_r, r_glm); // "training"
	cout << "Training Mean Error: " << pr.first << endl;
	double regr_mse = regression_test(testing, *feat_r, r_glm);//, "testing");
	cout << "Testing Mean Error: " << regr_mse << endl;
	cout << "Features: "<< endl;
	for (auto line : feat_r->feat_names()) {
		cout << "\t" << line << endl;
	}
	auto w = r_glm.get_weights();
	for (int r = 0; r < w.getNumRow(); r++) {
		cout << "weight: ";
		for (int c = 0; c < w.getNumCol(); c++) {
			cout << w.get(r, c) << " ";
		}
		cout << endl;
	}
	// for (auto combo : feat.get_combos()) {
	// 	cout << combo.first << " " <<
	// }

}

template class Predictor<uint8_t>;
template class Predictor<uint16_t>;
template class Predictor<uint32_t>;
template class Predictor<uint64_t>;
template class Predictor<int>;
template class Predictor<double>;
```
