## Supplementary Data 1 for "FASTCAR: Rapid alignment-free prediction of sequence alignment identity scores using self-supervised general linear models": Predictor.h

```
/* -*- C++ -*-
 *
 * Predictor.h
 *
 * Author: Benjamin T James
 *
 * Main class for training and prediction
 * Does bulk training, but can be adapted for on-line training
 */

#ifndef PREDICTOR_H
#define PREDICTOR_H

#include "../cluster/src/GLM.h"
#include "../cluster/src/Point.h"
#include "../cluster/src/Feature.h"
#include <set>
#include "Random.h"
#include <omp.h>
#define PRED_MODE_CLASS 1
#define PRED_MODE_REGR  2

#define PRED_FEAT_FAST (FEAT_EUCLIDEAN | FEAT_MANHATTAN | FEAT_INTERSECTION | FEAT_KULCZYNSKI2 | FEAT_SIMRATIO | FEAT_NORMALIZED_VECTORS | FEAT_PEARSON_COEFF | FEAT_EMD | FEAT_LENGTHD )
#define PRED_FEAT_DIV (FEAT_JEFFEREY_DIV | FEAT_JENSEN_SHANNON)
#define PRED_FEAT_ALL (FEAT_HELLINGER|FEAT_MANHATTAN|FEAT_EUCLIDEAN|FEAT_CHI_SQUARED|FEAT_NORMALIZED_VECTORS|FEAT_HARMONIC_MEAN|FEAT_JEFFEREY_DIV|FEAT_K_DIV|FEAT_PEARSON_COEFF|FEAT_SQCHORD|FEAT_KL_COND|FEAT_MARKOV|FEAT_INTERSECTION|FEAT_RRE_K_R|FEAT_D2z|FEAT_SIM_MM|FEAT_EUCLIDEAN_Z|FEAT_EMD|FEAT_SPEARMAN|FEAT_JACCARD|FEAT_LENGTHD|FEAT_D2s|FEAT_AFD|FEAT_MISMATCH|FEAT_CANBERRA|FEAT_KULCZYNSKI1|FEAT_KULCZYNSKI2|FEAT_SIMRATIO|FEAT_JENSEN_SHANNON|FEAT_D2_star|FEAT_N2R|FEAT_N2RC|FEAT_N2RRC)

template<class T>
class Predictor {
public:
	Predictor(int k_, double id_, uint8_t mode_, uint64_t feats, int mut_type_, int min_num_feat_=3, int max_num_feat_=5, double min_id_=0.35) : k(k_), id(id_), is_trained(false), is_training(false), mode(mode_), max_num_feat(max_num_feat_), mut_type(mut_type_), min_num_feat(min_num_feat_), min_id(min_id_ * 100), feats64(feats) {
		add_feats(possible_feats, feats);
		feat_c = NULL;
		feat_r = NULL;
		omp_init_lock(&lock);
	};
	Predictor(const std::string filename);
	~Predictor() {
		possible_feats.clear();
		omp_destroy_lock(&lock);
		if (feat_c) {
			delete feat_c;
		}
		if (feat_r) {
			delete feat_r;
		}
		training.clear();
		testing.clear();
	}
	void train(const std::vector<Point<T>* >& vec, uintmax_t& _id, size_t num_sample);
	double similarity(Point<T>* a, Point<T>* b);
	bool close(Point<T>* a, Point<T>* b);
	void save(std::string file, std::string datatype);
	void check();
	uint8_t get_mode() const { return mode; }
	pair<Feature<T>*, matrix::GLM> get_class() { return std::make_pair(new Feature<T>(*feat_c), c_glm); }
	void mutate_seqs(Point<T>* p, size_t num_seq, vector<pra<T> > &,vector<pra<T> > & , double id_begin, double id_end, uintmax_t& _id, std::random_device::result_type seed);

	std::string get_datatype() const { return datatype; }
	int get_k() const { return k; }
	double get_id() const { return id; }
private:
	static void add_feats(std::vector<std::pair<uint64_t, Combo> >& vec, uint64_t flags);
	static pair<matrix::GLM, Feature<T>*> read_from(std::ifstream &in, int k_);
	static void write_to(std::ofstream &out, Feature<T>* f, matrix::GLM glm);
	void filter(std::vector<pra<T> > &s, std::string prefix="");
	void train();
	void train_class(Feature<T>* feat);
	void train_regr(Feature<T>* feat);
	void train_class_regr(Feature<T>* feat);
	double predict(Point<T>* a, Point<T>* b);
	bool p_close(Point<T>* a, Point<T>* b);
	double p_predict(Point<T>* a, Point<T>* b);

	Feature<T> *feat_c, *feat_r;
	matrix::GLM c_glm, r_glm;
        vector<pra<T> > training, testing;
	bool is_trained, is_training;
	int min_num_feat, max_num_feat, k, mut_type;
	uint8_t mode;
	double id, min_id;
	vector<std::pair<uint64_t, Combo> > possible_feats;
	omp_lock_t lock;
	Random random;
	uint64_t feats64;
	std::string datatype;
	double scale_min = 1000;
	double scale_max = -1000;
};
#endif
```
