## Supplementary Data 1 for "FASTCAR: Rapid alignment-free prediction of sequence alignment identity scores using self-supervised general linear models": Random.h

```
// -*- C++ -*-
#ifndef RANDOM_H
#define RANDOM_H
#include <random>
#include <iostream>
class Random {
public:
	Random(std::random_device::result_type seed=0xAA) : mt(seed) {}

	template<class T>
	T randMod(T max) {
		T res;
#pragma omp critical
		{
			if (max == 0) {
				res = 0;
			} else {
				std::uniform_int_distribution<T> distribution(0, max-1);
				res = distribution(mt);
			}
		}
		return res;
	}

	double random() {
		double res = 0;
		#pragma omp critical
		{
		std::uniform_real_distribution<double> distribution(0.0, 1.0);
		res = distribution(mt);
		}
		return res;
	}
	double rand_between(double id, double range, double low, double high) {
		double res = 0;
		#pragma omp critical
		{
		double mn = std::max(id - range, low);
		double mx = std::min(id + range, high);
		std::uniform_real_distribution<double> distribution(mn, mx);

		res = distribution(mt);
		}
		return res;
	}
	std::random_device::result_type nextRandSeed() {
		using rt = std::random_device::result_type;
		return randMod<rt>(std::numeric_limits<rt>::max());
	}
	std::mt19937& gen() { return mt; }
private:
        std::mt19937 mt;

};
#endif
```
