## Supplementary Data 1 for "FASTCAR: Rapid alignment-free prediction of sequence alignment identity scores using self-supervised general linear models": SingleMute.cpp

```
/**
 * Author: Alex Baumgartner
 * The Bioinformatics Toolsmith Laboratory, the University of Tulsa
 * 5/15/2018
 *
 * Purpose:
 *	The pupose of this module is to perform single mutations on sequences
 */

#include "SingleMute.h"

SingleMute::SingleMute(int a, int c, int g, int t, int alloc) {
	percAs = a;
	percCs = c;
	percGs = g;
	percTs = t;
	//If allocation is 0, all sub allocations are 0
	if (alloc == 0) {
		maxDel = 0;
		maxInsert = 0;
		maxSwitch = 0;
	}
	//Arbitrary, if only 1 percent is allocated, then only insert gets an allocation
	else if (alloc == 1) {
		maxSwitch = 0;
		maxDel = 0;
		maxInsert = 1;
	}
	//Otherwise, allocations are assigned randomly
	else {
		//Max switch gets a random allocation,
		//but allocation has to be even
		//(don't want to switch something with itself)
		maxSwitch = rand() % alloc;
		alloc -= maxSwitch;
		//If alloc is odd,
		//and there is still percent that can be allocated
		if (maxSwitch % 2 == 1 && alloc >= 1) {
			//Make allocation 1 less,
			//and switch allocation one more (now even)
			maxSwitch++;
			alloc--;
		}
		//Otherwise, make allocation one larger,
		//switch allocation one less (even)
		else if (alloc == 0) {
			maxSwitch--;
			alloc++;
		}
		//If alloc is greater than 1 (must be for % purposes),
		//calculate random value for inerst allocation
		if (alloc > 1) {
			maxInsert = rand() % alloc;
			alloc -= maxInsert;
		} else {
			maxInsert = alloc;
			alloc -= maxInsert;
		}
		//Max delete is assigned whatever is left
		maxDel = alloc;
	}
}

int SingleMute::getAlignmentLength(){
	return alignmentLength;
}

int SingleMute::getIBP(){
	return IBP;
}

void SingleMute::genSing(string * sequence, vector<bool> mutes) {
	seq = sequence;
	//Assign vector of mutes to inputted vector
	validIndexes = new vector<int>();
	validIndexes->reserve(mutes.size());
	//Adds all valid indexes to the validIndexes vector
	for(int i = 0; i < mutes.size(); i++){
		if(mutes.at(i)){
			validIndexes->push_back(i);
		}
	}
	float tempFloat;
	//Calculate number of characters each mutation can mutate
	tempFloat = maxDel / 100.0;
	maxDel = (int) (tempFloat * seq->length());
	tempFloat = maxInsert / 100.0;
	maxInsert = (tempFloat * seq->length());
	tempFloat = maxSwitch / 100.0;
	maxSwitch = (tempFloat * seq->length());
	//Calculates Alignment length and identical base pairs
	alignmentLength = maxInsert;
	IBP = maxDel + maxSwitch;
	//Vectors to keep track of where insertions and deletions need to be made
	insertions = new vector<int>();
	insertions->reserve(maxInsert);
	deletions = new vector<int>();
	deletions->reserve(maxDel);
	//Since switch makes 2 invalid,
	//switchNucl is run maxSwitch/2 times
	for (int i = 0; i < maxSwitch; i++) {
		switchNucl();
	}
	//Insert maxInsert times
	for (int i = 0; i < maxInsert; i++) {
		insert();
	}
	//Delete maxDel nucleotides
	for (int i = 0; i < maxDel; i++) {
		deleteNucl();
	}
	//perfroms deletions and insertions
	performInsertAndDelete();
}

void SingleMute::insert() {
	//Calculate the index to insert at
	int index = rand() % validIndexes->size();
	insertions->push_back(validIndexes->at(index));
	//Remove that as a valid index
	validIndexes->erase(validIndexes->begin() + index, validIndexes->begin() + index + 1);
}

void SingleMute::deleteNucl() {
	//Choose a valid index to delete
	int index = rand() % validIndexes->size();
	deletions->push_back(validIndexes->at(index));
	//Remove from the 
	validIndexes->erase(validIndexes->begin() + index, validIndexes->begin() + index + 1);
}

void SingleMute::switchNucl() {
	//Pick a random valid index
	int index = rand() % validIndexes->size();
	char character = seq->at(validIndexes->at(index));
	int value;
	//Keep generating characters until one different than the one we are trying to switch is found
	while(character == seq->at(validIndexes->at(index))){
		value = rand() % (percAs + percCs + percGs + percTs);
		if (value < percAs) {
			character = 'A';
		} else if (value < percAs + percCs) {
			character = 'C';
		} else if (value < percAs + percCs + percGs) {
			character = 'G';
		} else {
			character = 'T';
		}
	}
	//Switch that character
	seq->at(validIndexes->at(index)) = character;
	//Remove the chosen index as a valid index
	validIndexes->erase(validIndexes->begin() + index, validIndexes->begin() + index + 1);
}

void SingleMute::performInsertAndDelete(){
	//sorts the vectors based
	std::sort(insertions->begin(), insertions->end());
	std::sort(deletions->begin(), deletions->end());
	//Goes through both vectors untill all have been processed
	for(int i = insertions->size() - 1, j = deletions->size() - 1; i >= 0 && j >= 0;){
		//If i is -1, all insertions have been processed
		if(i == -1){
			removeNucl(deletions->at(j));
			j--;
		}
		//If i is -1, all deletions have been processed
		else if(j == -1){
			insertNucl(insertions->at(i));
			i--;
		}
		else{
			//If the index of the current next insertion is higher than the next deletion, insert, else delete
			if(insertions->at(i) > deletions->at(j)){
				insertNucl(insertions->at(i));
				i--;
			}
			else{
				removeNucl(deletions->at(j));
				j--;
			}
		}
	}
}

void SingleMute::removeNucl(int index){
	seq->erase(index, 1);
}

void SingleMute::insertNucl(int index){
	string character;
	//Use a weighted die to
	//calculate which character to insert
	int value = rand() % (percAs + percCs + percGs + percTs);
	if (value < percAs) {
		character = "A";
	} else if (value < percAs + percCs) {
		character = "C";
	} else if (value < percAs + percCs + percGs) {
		character = "G";
	} else {
		character = "T";
	}
	//insert at that index
	seq->insert(index, character);
}
```
