## Supplementary Data 1 for "FASTCAR: Rapid alignment-free prediction of sequence alignment identity scores using self-supervised general linear models": SingleMute.h

```
/**
 * Author: Alex Baumgartner
 * The Bioinformatics Toolsmith Laboratory, the University of Tulsa
 * 5/15/2018
 *
 * Purpose:
 *	The pupose of this module is to perform single mutations on sequences
 */

#ifndef SINGLEMUTE_H
#define  SINGLEMUTE_H

#include <iostream>
#include <vector>
#include <string>
#include <algorithm>

using namespace std;

class SingleMute {
public:
	/*
	 Constructor, creates values
	 	and assignes allocations based on inputted data

	 @param:
	 int: percentage of A's
	 int: percentage of C's
	 int: percentage of G's
	 int: percentage of T's
	 int: The total allocation for non-single mutations
	 */
	SingleMute(int, int, int, int, int);
	/*
	 Takes a string and mutates it based
	 	on the allocation given in the constructor

	 @param:
	 std::string *: pointer to the sequence to be mutated
	 std::vector<bool> : boolean vector of valid and invalid indexes
	 */
	void genSing(std::string *, std::vector<bool>);

	int getAlignmentLength();

	int getIBP();
	
	~SingleMute(){delete validIndexes; delete insertions; delete deletions;};

  private:
	int percAs;
	int percCs;
	int percGs;
	int percTs;
	int maxDel;
	int maxInsert;
	int maxSwitch;
	int alignmentLength;
	int IBP;
	std::vector<int> * validIndexes;
	std::vector<int> * deletions;
	std::vector<int> * insertions;
	std::string * seq;
	/*
	 Inserts a sequence randomly in the list
	 	at a valid index
	 */
	void insert();
	/*
	 Deletes a random nucleotide
	 	that has not been previously mutated
	 */
	void deleteNucl();
	/*
	 Switches two random nucleotides
	 	that have not been mutated previously
	 */
	void switchNucl();
	/*
	Performs necessary insertions and deletions in the string based on the insertion and deletion vectors
	*/
	void performInsertAndDelete();

	void removeNucl(int);

	void insertNucl(int);
};
#endif
```
