## Supplementary Data 1 for "FASTCAR: Rapid alignment-free prediction of sequence alignment identity scores using self-supervised general linear models": SingMute.cpp

```
#include "SingMute.h"
#include <set>
#include <random>
#include <algorithm>
#include <iostream>

#ifdef MUTDEBUG
static const std::string INSERT_BEGIN = "[";
static const std::string INSERT_END = "]";
static const std::string SWITCH_BEGIN = "(";
static const std::string SWITCH_END = ")";
static const std::string DEL = "-";
#else
static const std::string INSERT_BEGIN = "";
static const std::string INSERT_END = "";
static const std::string SWITCH_BEGIN = "";
static const std::string SWITCH_END = "";
static const std::string DEL = "";
#endif


char SingMute::randNucl()
{
	char character;
	int value = rng.randMod<int>(percAs + percCs + percGs + percTs);
//	int value = 40436 % (percAs + percCs + percGs + percTs);
	if (value < percAs) {
		character = 'A';
	} else if (value < percAs + percCs) {
		character = 'C';
	} else if (value < percAs + percCs + percGs) {
		character = 'G';
	} else {
		character = 'T';
	}
	return character;
}
void SingMute::init(const std::vector<bool> &valid)
{
	maxInsert = 0;
	maxDel = 0;
	maxSwitch = 0;
	if (num_mut == 0) {
		out_seq = std::string(*seq);
		IBP = 0;
		alignmentLength = 0;
		return;
	} else if (num_mut == 1) {
		maxInsert = 1;
		maxDel = 0;
		maxSwitch = 0;
	} else {
		maxSwitch = rng.randMod<long>(num_mut);
		num_mut -= maxSwitch;

		if (maxSwitch % 2 == 1 && num_mut >= 1) {
			maxSwitch++;
			num_mut--;
		} else if (num_mut == 0) {
			maxSwitch--;
			num_mut++;
		}
		if (num_mut > 1) {
			maxInsert = rng.randMod<long>(num_mut);
			num_mut -= maxInsert;
		} else {
			maxInsert = num_mut;
			num_mut -= maxInsert;
		}
		maxDel = num_mut;
	}
	size_t seq_len = seq->length();

	maxDel *= seq_len / 100.0;
	maxInsert *= seq_len / 100.0;
	maxSwitch *= seq_len / 100.0;
	alignmentLength = maxInsert;
	IBP = maxDel + maxSwitch;


	std::vector<char> command_str(seq_len, 'S');
	long idx = 0;
	long nons_len = maxInsert + maxDel + maxSwitch;
	for (long i = 0; i < maxInsert; i++) {
		command_str[idx++] = 'I';
	}
	for (long i = 0; i < maxDel; i++) {
		command_str[idx++] = 'D';
	}
	for (long i = 0; i < maxSwitch; i++) {
		command_str[idx++] = 'W';
	}
	//std::shuffle(command_str.begin(), command_str.end(), rng.gen());
	std::shuffle(command_str.begin(), command_str.end(), std::minstd_rand0(rng.nextRandSeed()));
	std::vector<long> valid_indices;
	long repl = command_str.size() - 1;
	for (long i = 0; i < command_str.size(); i++) {
		if (command_str[i] != 'S' && !valid[i]) {
			if (!valid_indices.empty()) {
				repl = valid_indices.back();
				valid_indices.pop_back();
			} else {
				for (; repl > 0; repl--) {
					if (valid[repl]) {
						break;
					}
				}
			}
			std::swap(command_str[i], command_str[repl]);
		} else if (command_str[i] == 'S'
			   && valid[i]
			   && valid_indices.size() < nons_len) {

			valid_indices.push_back(i);
		}
	}
	// std::set<long> s_ins, s_del, s_switch;
	// generate_unique_set(command_str.size(), s_ins, maxInsert, s_del, s_switch, valid);
	// generate_unique_set(command_str.size(), s_del, maxDel, s_ins, s_switch, valid);
	// generate_unique_set(command_str.size(), s_switch, maxSwitch, s_ins, s_del, valid);
	// for (auto idx : s_ins) {
	// 	command_str[idx] = 'I';
	// }
	// for (auto idx : s_del) {
	// 	command_str[idx] = 'D';
	// }
	// for (auto idx : s_switch) {
	// 	command_str[idx] = 'W';
	// }
	out_seq = "";
	out_seq.reserve(maxInsert + seq_len - maxDel + 1);

	for (long i = 0; i < seq_len; i++) {
		auto cmd = command_str.at(i);
		switch (cmd) {
		case 'I': {
			out_seq += INSERT_BEGIN + randNucl() + INSERT_END;
			out_seq += seq->at(i);
			break;
		}
		case 'S': {
			out_seq += seq->at(i);
			break;
		}
		case 'D': {
			out_seq += DEL;
			break;
		}
		case 'W': {
			out_seq += SWITCH_BEGIN + randNucl() + SWITCH_END;
			break;
		}
		}
	}
}
```
