## Supplementary Data 1 for "FASTCAR: Rapid alignment-free prediction of sequence alignment identity scores using self-supervised general linear models": SingMute.h

```
#ifndef SINGMUTE_H
#define SINGMUTE_H

#include <vector>
#include <string>
#include "Random.h"
#include "LCG.h"

class SingMute {
public:
	/*
	 Constructor, creates values
	 	and assignes allocations based on inputted data

	 @param:
	 int: percentage of A's
	 int: percentage of C's
	 int: percentage of G's
	 int: percentage of T's
	 int: The total allocation for non-single mutations
	 */
	SingMute(int pa, int pc, int pg, int pt, uintmax_t tt, const std::string* s, const std::vector<bool> &valid_, std::random_device::result_type seed) : percAs(pa),
																	percCs(pc), percGs(pg), percTs(pt), num_mut(tt), seq(s), rng(seed) {
		init(valid_);
	}
	long getAlignmentLength() { return alignmentLength; }
	long getIBP() { return IBP; }
	void init(const std::vector<bool> &valid);
	std::string& getSeq() { return out_seq; };
  private:
	uintmax_t num_mut;
	int percAs;
	int percCs;
	int percGs;
	int percTs;

	long maxDel;
	long maxInsert;
	long maxSwitch;

	long alignmentLength;
	long IBP;
	const std::string * seq;
	std::string out_seq;
	char randNucl();
	LCG rng;
};
#endif
```
