## Supplementary Data 1 for "FASTCAR: Rapid alignment-free prediction of sequence alignment identity scores using self-supervised general linear models": Trainer.cpp

```
/* -*- C++ -*-
 *
 * Trainer.cpp
 *
 * Author: Benjamin T James
 *
 * Artifact from MeShClust
 */
#include "Trainer.h"

#include <algorithm>
#include <set>
#include <map>
#include <cmath>
#include "../cluster/src/Point.h"
#include "../cluster/src/GLM.h"
#include "../cluster/src/Feature.h"
#include "../cluster/src/Progress.h"

#include "../cluster/src/DivergencePoint.h"
#include <random>


template<class T>
std::tuple<Point<T>*,double,size_t,size_t> Trainer<T>::get_close(Point<T> *p, bvec_iterator<T> istart, bvec_iterator<T> iend, bool &is_min_r) const
{
	int ncols = weights.getNumRow();
#pragma omp declare reduction(pmax:std::tuple<Point<T>*,double,size_t,size_t>: \
			      omp_out = get<1>(omp_in) > get<1>(omp_out) ? omp_in : omp_out ) \
	initializer (omp_priv=std::make_tuple((Point<T>*)NULL,-1,0,0))

	std::tuple<Point<T>*,
		   double,
		   size_t,
		   size_t> result = {NULL,
				     -1,
				     0,
				     0};
	bool has_found = false;

	#ifdef DEBUG
	cout << "begin " << istart.r << " " << istart.c << " end " << iend.r << " " << iend.c << endl;
	for (auto data : *istart.col) {
		cout << "\t" << data.size() << endl;
	}
	#endif

	bool is_min = true;
#pragma omp parallel for reduction(pmax:result), reduction(&&:is_min)
	for (bvec_iterator<T> i = istart; i <= iend; ++i) {
		Point<T>* pt = (*i).first;
		double sum = weights.get(0, 0);
		double dist = 0;
		auto cache = feat->compute(*pt, *p);
		for (int col = 1; col < ncols; col++) {
			if (col == 1) {
				dist = (*feat)(col-1, cache);
				sum += weights.get(col, 0) * dist;
			} else {
				sum += weights.get(col, 0) * (*feat)(col-1, cache);
			}
		}
		double res = round(1.0 / (1 + exp(-sum)));
		//cout << "res: " << res << " " << dist << endl;
// set second to true if result is not 1.0
		// which means it will be removed
		result = (dist > std::get<1>(result)) ? std::make_tuple(pt, dist, i.r, i.c) : result;
		is_min = is_min && (res != 1.0);
//		has_found = has_found || (res != 1.0);
		if (res == 1.0) {
			*i = std::make_pair(pt, true);
//			(*i).second = true;
		}
	}

//	is_min = !has_found;
	is_min_r = is_min;
//	return get<0>(result);
	return result;

}

template<class T>
long Trainer<T>::merge(vector<Center<T> > &centers, long current, long begin, long last) const
{
#pragma omp declare reduction(ldpmax:std::pair<long,double>:			\
			      omp_out = omp_in.second > omp_out.second ? omp_in : omp_out ) \
	initializer (omp_priv=std::make_pair(0, std::numeric_limits<double>::min()))
	std::pair<long,double> best = std::make_pair(0, std::numeric_limits<double>::min());
	Point<T>* p = centers[current].getCenter();
#pragma omp parallel for reduction(ldpmax:best)
	for (long i = begin; i <= last; i++) {
		double sum = weights.get(0, 0);
		double dist = 0;
		Point<T>* cen = centers[i].getCenter();
		auto cache = feat->compute(*cen, *p);
		for (int col = 1; col < weights.getNumRow(); col++) {
			double d = (*feat)(col-1, cache);
			if (col == 1) {
				dist = d;
			}
			sum += weights.get(col, 0) * d;
		}
		double res = round(1.0 / (1 + exp(-sum)));

		if (res == 1) {
			best = best.second > dist ? best : std::make_pair(i, dist);
		}
	}
	return best.first;
}


template<class T>
void Trainer<T>::filter(Point<T> *p, vector<pair<Point<T> *, bool> > &vec) const
{
	for (auto& pt : vec) {
		double sum = weights.get(0, 0);
		auto cache = feat->compute(*pt.first, *p);
		for (int col = 1; col < weights.getNumRow(); col++) {
			sum += weights.get(col, 0) * (*feat)(col-1, cache);
		}
		double res = round(1.0 / (1 + exp(-sum)));
		pt.second = (res != 1);
	}
	vec.erase(std::remove_if(vec.begin(), vec.end(), [](pair<Point<T>*, bool> p) {
				return p.second;
			}), vec.end());
}

template<class T>
Point<T>* Trainer<T>::closest(Point<double> *p, vector<pair<Point<T> *, bool> > &vec) const
{
	Point<T>* best_pt = NULL;
	double best_dist = 0;
	for (auto& pt : vec) {
		double sum = weights.get(0, 0);
		double dist = pt.first->distance_d(*p);
		if (best_pt == NULL || dist < best_dist) {
			best_dist = dist;
			best_pt = pt.first;
		}
	}
	return best_pt;
}

template<class T>
std::pair<matrix::Matrix,matrix::Matrix> Trainer<T>::generate_feat_mat(pair<vector<pra<T> >, vector<pra<T> > > &data, int ncols, bool do_set_to_1)
{
	int nrows = data.first.size() + data.second.size();
	matrix::Matrix feat_mat(nrows, ncols);
	matrix::Matrix labels(nrows, 1);
#pragma omp parallel for
	for (int i = 0; i < data.first.size(); i++) {
		auto kv = data.first[i];
		int row = i;
		auto cache = feat->compute(*kv.first, *kv.second);
		for (int col = 0; col < ncols; col++) {

			if (col == 0) {
				feat_mat.set(row, col, 1);//do_set_to_1 ? 1 : kv.val);
			} else {
//				double val = ff[col-1](kv.first, kv.second);
				////#pragma omp critical
				double val = (*feat)(col-1, cache);
				feat_mat.set(row, col, val);
			}

		}
		////#pragma omp critical
		labels.set(row, 0, do_set_to_1 ? 1 : kv.val);
	}
	// if (!do_set_to_1) {
	// 	return std::make_pair(feat_mat, labels);
	// }
#pragma omp parallel for
	for (int i = 0; i < data.second.size(); i++) {
		auto kv = data.second[i];
		int row = data.first.size() + i;
		auto cache = feat->compute(*kv.first, *kv.second);
		for (int col = 0; col < ncols; col++) {

			if (col == 0) {
				feat_mat.set(row, col, 1);//do_set_to_1 ? 1 : kv.val);
			} else {
//				double val = ff[col-1](kv.first, kv.second);
				////#pragma omp critical
				double val = (*feat)(col-1, cache);
				feat_mat.set(row, col, val);
			}

		}
		////#pragma omp critical
		labels.set(row, 0, do_set_to_1 ? -1 : kv.val);
	}
	return std::make_pair(feat_mat, labels);
}
template<class T>
double Trainer<T>::train_n(pair<vector<pair<Point<T> *, Point<T> *> >, vector<pair<Point<T> *, Point<T> *> > > &data, int ncols)
{
	std::cout << "done" << endl;
	cout << "Training on " << ncols << " columns" << endl;
	int nrows = data.first.size() + data.second.size();

	matrix::Matrix feat_mat(nrows, ncols);
	matrix::Matrix labels(nrows, 1);
	double avg_label = 0;
#pragma omp parallel for
	for (int i = 0; i < data.first.size(); i++) {
		auto kv = data.first[i];
		int row = i;
		auto cache = feat->compute(*kv.first, *kv.second);
		for (int col = 0; col < ncols; col++) {

			if (col == 0) {
				feat_mat.set(row, col, 1);
			} else {
//				double val = ff[col-1](kv.first, kv.second);
				////#pragma omp critical
				double val = (*feat)(col-1, cache);
				feat_mat.set(row, col, val);
			}

		}
		////#pragma omp critical
		labels.set(row, 0, 1);
	}
#pragma omp parallel for
	for (int i = 0; i < data.second.size(); i++) {
		auto kv = data.second[i];
		int row = data.first.size() + i;
		auto cache = feat->compute(*kv.first, *kv.second);
		for (int col = 0; col < ncols; col++) {

			if (col == 0) {
				feat_mat.set(row, col, 1);
			} else {
//				double val = ff[col-1](kv.first, kv.second);
				////#pragma omp critical
				double val = (*feat)(col-1, cache);
				feat_mat.set(row, col, val);
			}

		}
		////#pragma omp critical
		labels.set(row, 0, -1);
	}
	for (int row = 0; row < nrows; row++) {
		for (int col = 0; col < ncols; col++) {
			double val = feat_mat.get(row, col);
			std::cout << val << "\t";
		}
		std::cout << endl;
	}
	glm.train(feat_mat, labels);
	weights = glm.get_weights();
	#ifdef DEBUG
	for (int i = 0; i < ncols; i++) {
		cout << "weight: " << weights.get(i, 0) << endl;

	}
	#endif
	matrix::Matrix p = glm.predict(feat_mat);
	for (int row = 0; row < nrows; row++) {
		if (p.get(row, 0) == 0) {
			p.set(row, 0, -1);
		}
	}
	auto tup = glm.accuracy(labels, p);
	return get<0>(tup);
}

template<class T>
Point<T>* seq_by_name(std::string name, vector<Point<T>*> points)
{
	for (auto p : points) {
		if (p->get_header() == name) {
			return p;
		}
	}
	return NULL;
}
template<class T>
void Trainer<T>::train(pair<vector<pra<T> >,
		       vector<pra<T> > > training,
		       pair<vector<pra<T> >,
		       vector<pra<T> > >testing,
		       double acc_cutoff)
{
	vector<std::pair<uint64_t, Combo> > bit_feats;
	//bit_feats.push_back(std::make_pair(FEAT_ALIGN, COMBO_SELF));
	Feature<T> allfeat(k);
	Feature<T> best_class(k);
	Feature<T> best_regr(k);
	auto filter_feat_good = [](uint64_t flag) {
		if ((flag & FEAT_SIM_MM) | (flag & FEAT_MARKOV)) {
			return false;
		}
		if (flag & FEAT_SPEARMAN) {
			return false;
		}
		if ((flag & FEAT_N2R) | (flag & FEAT_N2RC) | (flag & FEAT_N2RRC)) {
			return false;
		}
		if ((flag & FEAT_KL_COND) | (flag & FEAT_AFD)) {
			return false;
		}
		return true;
	};
	auto class_train = [&]() {
		auto mtraining = generate_feat_mat(training, feat->size()+1);
		auto mtesting = generate_feat_mat(testing, feat->size()+1);
		glm.train(mtraining.first, mtraining.second);


		weights = glm.get_weights();
		matrix::Matrix p = glm.predict(mtesting.first);
		for (int row = 0; row < testing.first.size() + testing.second.size(); row++) {
			if (p.get(row, 0) == 0) {
				p.set(row, 0, -1);
			}
		}
		double acc = get<0>(glm.accuracy(mtesting.second, p));
//		glm.accuracy(mtraining.second, q);
		return acc;
	};

	auto regr_train = [&]() {
		auto rtraining = generate_feat_mat(training, feat->size()+1, false);
		auto rtesting = generate_feat_mat(testing, feat->size()+1, false);
		regr.train(rtraining.first, rtraining.second);
		auto result1 = rtraining.first * regr.get_weights();
		auto diff1 = result1 - rtraining.second;
		double sum = 0;
		for (int i = 0; i < diff1.getNumRow(); i++) {
			sum += fabs(diff1.get(i, 0));
		}
		sum /= diff1.getNumRow();
		//cout << allfeat.size() << " [" << diff1.getNumRow() << "] Training Mean Error: " << sum << endl;
		auto result2 = rtesting.first * regr.get_weights();
		auto diff2 = result2 - rtesting.second;

		sum = 0;
		for (int i = 0; i < diff2.getNumRow(); i++) {
			sum += fabs(diff2.get(i, 0));
		}
		sum /= diff2.getNumRow();
//		cout << allfeat.size() << " [" << diff2.getNumRow() << "] Testing Mean Error: " << sum << endl;

		return sum;
	};

	vector<pair<uint64_t, Combo> > possible_feats, regr_feats, class_feats;
	for (uint64_t ia = 1; ia <= 33; ia++) {
		if (!filter_feat_good(1UL << ia)) {
			continue;
		}
		for (uint64_t ib = 1; ib <= ia; ib++) {
			if (!filter_feat_good(1UL << ib)) {
				continue;
			}
			possible_feats.emplace_back((1UL << ia) | (1UL << ib), Combo::xy);
			possible_feats.emplace_back((1UL << ia) | (1UL << ib), Combo::x2y);
			possible_feats.emplace_back((1UL << ia) | (1UL << ib), Combo::xy2);
			possible_feats.emplace_back((1UL << ia) | (1UL << ib), Combo::x2y2);
		}
	}
	double abs_best_regr = 100000000;
	double abs_best_class = 0;
	for (int num_feat = 1; num_feat <= 8; num_feat++) {
		double best_class_acc = 0;
		double best_regr_err = 100000000;
		auto backup_class = best_class;
		auto backup_regr = best_regr;
		auto best_class_feat = possible_feats.front();
		auto best_regr_feat = possible_feats.front();
		for (auto rfeat : possible_feats) {
			//cout << "Trying feat " << rfeat.first << " " << (rfeat.second == Combo::Self ? "self" : "squared") << endl;
			best_class.add_feature(rfeat.first, rfeat.second);
			best_class.normalize(training.first);
			best_class.normalize(training.second);
			best_class.finalize();
			feat = &best_class;
			double class_acc = class_train();
			cout << "class acc: " << class_acc << endl;
			if (class_acc > best_class_acc) {
				best_class_feat = rfeat;
				best_class_acc = class_acc;
			}
			best_regr.add_feature(rfeat.first, rfeat.second);
			best_regr.normalize(training.first);
			best_regr.normalize(training.second);
			best_regr.finalize();
			feat = &best_regr;
			double regr_mse = regr_train();
			cout << "regr mse: " << regr_mse << endl;
			if (regr_mse < best_regr_err) {
				best_regr_err = regr_mse;
				best_regr_feat = rfeat;
			}
			best_class = backup_class;
			best_regr = backup_regr;
		}
		if (best_class_acc > abs_best_class) {
			auto mvec = Feature<T>::multi_to_log(best_class_feat.first);
			cout << "CLASS " << num_feat << ": " << best_class_acc << " -> " << (best_class_feat.second == Combo::xy ? "self" : "squared") << " ";
			for (auto i : mvec) {
				cout << i << " ";
			}
			cout << endl;
			class_feats.push_back(best_class_feat);
			best_class.add_feature(best_class_feat.first, best_class_feat.second);
			best_class.normalize(training.first);
			best_class.normalize(training.second);
			best_class.finalize();
			abs_best_class = best_class_acc;

		}
		if (best_regr_err < abs_best_regr) {
			auto mvec = Feature<T>::multi_to_log(best_regr_feat.first);
			cout << "REGR " << num_feat << ": " << best_regr_err << " -> " << (best_regr_feat.second == Combo::xy ? "self" : "squared") << " ";
			for (auto i : mvec) {
				cout << i << " ";
			}
			cout << endl;

			regr_feats.push_back(best_regr_feat);
			best_regr.add_feature(best_regr_feat.first, best_regr_feat.second);
			best_regr.normalize(training.first);
			best_regr.normalize(training.second);
			best_regr.finalize();
			abs_best_regr = best_regr_err;

		}

	}
	cout << "Done with feature training" << endl;
	exit(0);

	int feat_set = 1;
	if (k == 0) {
		feat->add_feature(FEAT_ALIGN, Combo::xy);
		feat->normalize(training.first);
		feat->finalize();
		weights = matrix::Matrix(2, 1);
		weights.set(0, 0, -1 * cutoff);
		weights.set(1, 0, 1);
		return;
	} else if (feat_set == 0) {
		bit_feats.push_back(std::make_pair(FEAT_LENGTHD | FEAT_INTERSECTION, Combo::xy));
		bit_feats.push_back(std::make_pair(FEAT_LENGTHD | FEAT_JENSEN_SHANNON, Combo::xy));
		bit_feats.push_back(std::make_pair(FEAT_SIMRATIO, Combo::xy));
		bit_feats.push_back(std::make_pair(FEAT_SQCHORD, Combo::xy));
	} else {
		bit_feats.push_back(std::make_pair(FEAT_INTERSECTION | FEAT_LENGTHD, Combo::xy));
		bit_feats.push_back(std::make_pair(FEAT_MANHATTAN | FEAT_LENGTHD, Combo::x2y2));
		bit_feats.push_back(std::make_pair(FEAT_PEARSON_COEFF, Combo::xy));
		bit_feats.push_back(std::make_pair(FEAT_KULCZYNSKI2 | FEAT_LENGTHD, Combo::x2y2));
	}
	bool do_regr = true;
	double prev_acc = -10000;
	vector<matrix::Matrix> matvec;
	vector<Feature<T> > features;
	const size_t min_no_features = std::max(1, (int)bit_feats.size()-1);
	for (size_t num_features = min_no_features; num_features <= bit_feats.size(); num_features++) {
		for (size_t j = feat->size(); j < num_features && j < bit_feats.size(); j++) {
			feat->add_feature(bit_feats[j].first, bit_feats[j].second);
		}
		feat->normalize(training.first);
		feat->normalize(training.second);
		feat->finalize();
		feat->print_bounds();
		auto mtraining = generate_feat_mat(training, num_features+1);
		auto mtesting = generate_feat_mat(testing, num_features+1);
		glm.train(mtraining.first, mtraining.second);
		if (do_regr) {
			auto rtraining = generate_feat_mat(training, num_features+1, false);
			regr.train(rtraining.first, rtraining.second);
			auto result1 = rtraining.first * regr.get_weights();
			auto diff1 = result1 - rtraining.second;
			double sum = 0;
			for (int i = 0; i < diff1.getNumRow(); i++) {
				sum += fabs(diff1.get(i, 0));
			}
			sum /= diff1.getNumRow();
			cout << "[" << diff1.getNumRow() << "] Training Mean Error: " << sum << endl;

			auto rtesting = generate_feat_mat(testing, num_features+1, false);
			auto result2 = rtesting.first * regr.get_weights();
			auto diff2 = result2 - rtesting.second;

			sum = 0;
			for (int i = 0; i < diff2.getNumRow(); i++) {
				sum += fabs(diff2.get(i, 0));
			}
			sum /= diff2.getNumRow();
			cout << "[" << diff2.getNumRow() << "] Testing Mean Error: " << sum << endl;
		}
		weights = glm.get_weights();
		matrix::Matrix p = glm.predict(mtesting.first);
		for (int row = 0; row < testing.first.size() + testing.second.size(); row++) {
			if (p.get(row, 0) == 0) {
				p.set(row, 0, -1);
			}
		}
		double acc = get<0>(glm.accuracy(mtesting.second, p));
		matrix::Matrix q = glm.predict(mtraining.first);
		for (int row = 0; row < training.first.size() + training.second.size(); row++) {
			if (q.get(row, 0) == 0) {
				q.set(row, 0, -1);
			}
		}
		glm.accuracy(mtraining.second, q);
		if (acc - prev_acc <= 1 && acc >= 90.0) {
			weights = matvec.back();
			*feat = features.back();
			cout << "feat size is " << feat->size() << endl;
			break;
		}
		matvec.push_back(weights);
		features.push_back(*feat);
		prev_acc = acc;
		if (acc >= acc_cutoff) {
			cout << "breaking from acc cutoff" << endl;
			break;
		}
	}
	cout << "Final: feat size is " << feat->size() << endl;
	cout << "Using " << weights.getNumRow()-1 << " features " << __DATE__ << endl;
}


template class Trainer<uint8_t>;
template class Trainer<uint16_t>;
template class Trainer<uint32_t>;
template class Trainer<uint64_t>;
template class Trainer<int>;
template class Trainer<double>;
```
