## Supplementary Data 1 for "FASTCAR: Rapid alignment-free prediction of sequence alignment identity scores using self-supervised general linear models": Trainer.h

```
/* -*- C++ -*-
 *
 * Trainer.h
 *
 * Author: Benjamin T James
 */
#ifndef TRAINER_H
#define TRAINER_H

#include "../cluster/src/Point.h"
#include "../cluster/src/GLM.h"
#include "../cluster/src/Feature.h"
#include "../cluster/src/bvec.h"
#include "../cluster/src/Center.h"
#include "Predictor.h"
#include <set>
template<class T>
class Trainer {
public:
	Trainer(double cutoff_, int ksize=0) : cutoff(cutoff_), k(ksize) {
		uintmax_t size = 1000 * 1000 * 10;
		log_table = new double[size];
		log_coeff = size / 2;
		double lsize = log(size);
		log_table[0] = 0;
		for (uintmax_t i = 1; i < size; i++) {
			log_table[i] = log(2 * i) - lsize;
		}
		feat = new Feature<T>(k);
	};
	~Trainer() { delete feat_mat; delete feat; delete[] log_table;}

	double train_n(pair<vector<pair<Point<T>*,
			 Point<T>*
			 > >,
	     vector<pair<Point<T>*,
		       Point<T>*> > > &data, int ncols);
	void train(pair<vector<pra<T> >,
		        vector<pra<T> > > training,
		   pair<vector<pra<T> >,
		        vector<pra<T> > > testing,
		   double acc_cutoff=97.5);
	std::tuple<Point<T>*,double,size_t,size_t> get_close(Point<T>*, bvec_iterator<T> istart, bvec_iterator<T> iend,  bool& is_min) const;
	void filter(Point<T>*, vector<pair<Point<T>*,bool> >&) const;
	Point<T>* closest(Point<double>*, vector<pair<Point<T>*,bool> >&) const;
	long merge(vector<Center<T> > &centers, long current, long begin, long end) const;
private:
	matrix::GLM glm, regr;
	matrix::Matrix weights;
	std::pair<matrix::Matrix,matrix::Matrix> generate_feat_mat(pair<vector<pra<T> >,
								   vector<pra<T> > > &data, int ncols, bool do_set_to_1=true);

	Feature<T> *feat;
	double *log_table;
//	std::vector<Point<T>*> points;
	matrix::Matrix *feat_mat = NULL;
	double cutoff, log_coeff;
	int k;
};
#endif
```
