## Supplementary Data 1 for "FASTCAR: Rapid alignment-free prediction of sequence alignment identity scores using self-supervised general linear models": AffineId.cpp

```
/*
 * AffineId.cpp
 *
 *  Created on: Dec 6, 2012
 *  Modified on: Nov 6, 2017
 *      Author: Hani Zakaria Girgis, PhD
 */

// ToDo:
// 1. Add pre-conditions after testing
#include "AffineId.h"

#include "Util.h"
#include "../exception/InvalidInputException.h"

#include <iostream>
#include <cstring>
using namespace std;
//using namespace exception;

namespace utility {

AffineId::AffineId(const char * seq1In, int start1In, int end1In,
		const char * seq2In, int start2In, int end2In) {

	// The shorter of the two sequences is seq2
	seq1 = seq1In;
	start1 = start1In;
	end1 = end1In;

	seq2 = seq2In;
	start2 = start2In;
	end2 = end2In;

	if (end1 - start1 < end2 - start2) {
		seq1 = seq2In;
		start1 = start2In;
		end1 = end2In;

		seq2 = seq1In;
		start2 = start1In;
		end2 = end1In;
	}

	/*	if (start1 < 0 || end1 < 0 || start1 > end1) {
	 string msg("Invalid Input. Start1 is ");
	 msg.append(Util::int2string(start1));
	 msg.append(". End 1 is ");
	 msg.append(Util::int2string(end1));
	 msg.append(".");
	 //throw InvalidInputException(msg);

	 cerr << msg << endl;
	 throw exception();
	 }

	 if (start2 < 0 || end2 < 0 || start2 > end2) {
	 string msg("Invalid Input. Start2 is ");
	 msg.append(Util::int2string(start2));
	 msg.append(". End2 is ");
	 msg.append(Util::int2string(end2));
	 msg.append(".");
	 //throw InvalidInputException(msg);

	 cerr << msg << endl;
	 throw exception();
	 }*/

	// Validate input
	// cout << start1 << " " << end1 << endl;
	// cout << start2 << " " << end2 << endl;

	len1 = end1 - start1 + 2;
	len2 = end2 - start2 + 2;

	align();
}

AffineId::~AffineId() {
}

void AffineId::align() {
	// Initialize needed arrays
	auto m = new int[len2][2](); // Middle level array
	auto u = new int[len2][2](); // Upper level array
	auto mId = new int[len2][2](); // Array storing number of matches in the middle array
	auto uId = new int[len2][2](); // Array storing number of matches in the upper array
	auto mPath = new int[len2][2](); // Array storing number of steps in the middle array
	auto uPath = new int[len2][2](); // Array storing number of steps in the upper array

	// Apply the DP
	// The i index is only used to get a character from the first sequence
	// It is not used for filling the DP matrix
	for (int i = 1; i < len1; i++) {
		char base1 = seq1[start1 + i - 1];
		int lower = 0;
		int lowerId = 0;
		int lowerPath = 0;

		// j is the row. There are only two columns 0 and 1
		for (int j = 1; j < len2; j++) {
			// Update the lower value
			int extLower = lower + EXT;
			int openLower = m[j - 1][0] + OPEN;
			if (extLower > openLower) {
				lower = extLower;
				lowerPath++;
			} else {
				lower = openLower;
				lowerId = mId[j - 1][0];
				lowerPath = mPath[j - 1][0] + 1;
			}

			// Fill the array of the upper level
			int extUpper = u[j][0] + EXT;
			int openUpper = m[j][0] + OPEN;
			if (extUpper > openUpper) {
				u[j][1] = extUpper;
				uId[j][1] = uId[j][0];
				uPath[j][1] = uPath[j][0] + 1;
			} else {
				u[j][1] = openUpper;
				uId[j][1] = mId[j][0];
				uPath[j][1] = mPath[j][0] + 1;
			}

			// Fill the array of the middle level
			int matchOrMis;
			if (base1 == seq2[start2 + j - 1]) {
				matchOrMis = m[j - 1][0] + MATCH;
			} else {
				matchOrMis = m[j - 1][0] + MIS;
			}

			int lowerOrUpper;
			if (lower > u[j][1]) {
				lowerOrUpper = lower;
			} else {
				lowerOrUpper = u[j][1];
			}

			if (matchOrMis > lowerOrUpper) {
				m[j][1] = matchOrMis;
				mPath[j][1] = mPath[j - 1][0] + 1;
				if (base1 == seq2[start2 + j - 1]) {
					mId[j][1] = mId[j - 1][0] + 1;
				} else {
					mId[j][1] = mId[j - 1][0];
				}
			} else {
				m[j][1] = lowerOrUpper;
				if (lower > u[j][1]) {
					mId[j][1] = lowerId;
					mPath[j][1] = lowerPath;
				} else {
					mId[j][1] = uId[j][1];
					mPath[j][1] = uPath[j][1];
				}
			}
		}

		// // Test
		// for (int h = 0; h < len2; h++) {
		// 	cout << m[h][0] << "\t" << m[h][1] << "----" << mId[h][0] << "\t"
		// 			<< mId[h][1] << endl;
		// }
		// cout << "---------------------------------------------------" << endl;
		// // End of test

		// Copy the second column to the first one
		if (i != len1 - 1) {
			for (int h = 0; h < len2; h++) {
				m[h][0] = m[h][1];
				u[h][0] = u[h][1];
				mId[h][0] = mId[h][1];
				uId[h][0] = uId[h][1];
				mPath[h][0] = mPath[h][1];
				uPath[h][0] = uPath[h][1];
			}
		}
	}

	lenCS = mId[len2 - 1][1];
	lenPath = mPath[len2 - 1][1];
	//cout << "Alignment length = " << lenPath << endl;
	delete[] u;
	delete[] m;
	delete[] mId;
	delete[] uId;
	delete[] mPath;
	delete[] uPath;
}

double AffineId::getAlign() {
	double amt = lenCS;
	return amt / (double)lenPath;
}

}
/* namespace utility */

// // Testing code
// int main() {
// 	string s1("GATCTCAG");
// 	string s2("GACAG");

// 	utility::AffineId id(s1.c_str(), 0, s1.length() - 1, s2.c_str(), 0,
// 			s2.length() - 1);
// 	cout << "Length = " << id.getLenCS() << endl;

// 	return 0;
// }
```
