## Supplementary Data 1 for "FASTCAR: Rapid alignment-free prediction of sequence alignment identity scores using self-supervised general linear models": AffineId.h

```
/*
 * AffineId.h
 *
 *  Created on: Dec 6, 2012
 *  Modified on: Nov 6, 2017
 *      Author: Hani Zakaria Girgis, PhD
 */

#ifndef AFFINEID_H_
#define AFFINEID_H_

namespace utility {

class AffineId {
private:
	const char * seq1;
	int start1;
	int end1;
	const char * seq2;
	int start2;
	int end2;

	int len1;
	int len2;
	//int lenTotal;
	int lenCS;
	int lenPath;
	int * m; // Middle level
	//int * l; // Lower level
	int * u; // Upper level

	// const int MATCH = 4; // Score of a match
	// const int MIS = -4; // Score of a mismatch
	// const int OPEN = -2; // Score of a gap opening
	// const int EXT = -1; // Score of a gap extension

	const int MATCH = 1;
	const int MIS = -1;
	const int OPEN = -2;
	const int EXT = -1;
	void align();

public:
	AffineId(const char *, int, int, const char *, int, int);
	virtual ~AffineId();
        double getAlign();
};

} /* namespace utility */
#endif /* AFFINEID_H_ */
```
