## Supplementary Data 1 for "FASTCAR: Rapid alignment-free prediction of sequence alignment identity scores using self-supervised general linear models": EmptyLocation.cpp

```
/*
 * EmptyLocation.cpp
 *
 *  Created on: Dec 28, 2012
 *      Author: Hani Zakaria Girgis, PhD
 */

#include "EmptyLocation.h"
#include "../exception/InvalidOperationException.h"

using namespace exception;

namespace utility {

EmptyLocation * EmptyLocation::INSTANCE = new EmptyLocation();

EmptyLocation * EmptyLocation::getInstance(){
	return INSTANCE;
}

EmptyLocation::EmptyLocation() {
	msg = new string("Empty location does not allow this operation.");
}

EmptyLocation::~EmptyLocation() {
	delete msg;
}

string EmptyLocation::toString() {
	return string("Empty");
}

int EmptyLocation::getEnd() const {
	throw InvalidOperationException(*msg);
}

int EmptyLocation::getStart() const {
	throw InvalidOperationException(*msg);
}

void EmptyLocation::setEnd(int int1) {
	throw InvalidOperationException(*msg);
}

void EmptyLocation::setStart(int int1) {
	throw InvalidOperationException(*msg);
}

int EmptyLocation::getLength() {
	throw InvalidOperationException(*msg);
}

} /* namespace tr */
```
