## Supplementary Data 1 for "FASTCAR: Rapid alignment-free prediction of sequence alignment identity scores using self-supervised general linear models": EmptyLocation.h

```
/*
 * EmptyLocation.h
 *
 *  Created on: Dec 28, 2012
 *      Author: Hani Zakaria Girgis, PhD
 */

#ifndef EMPTYLOCATION_H_
#define EMPTYLOCATION_H_

#include "ILocation.h"

namespace utility {

class EmptyLocation: public ILocation {
private:
	string * msg;
	static EmptyLocation * INSTANCE;
	EmptyLocation();
	virtual ~EmptyLocation();

public:
	virtual int getEnd() const;
	virtual int getStart() const;
	virtual void setEnd(int);
	virtual void setStart(int);
	virtual int getLength();
	virtual string toString();

	static EmptyLocation * getInstance();

};

} /* namespace tr */
#endif /* EMPTYLOCATION_H_ */
```
