## Supplementary Data 1 for "FASTCAR: Rapid alignment-free prediction of sequence alignment identity scores using self-supervised general linear models": GlobAlignE.cpp

```
/**
 * Author: Joseph Valencia
 * Modified by Benjamin James
 * Date: 12/14/17
 * Bioinformatics Toolsmith Laboratory, University of Tulsa
 * */
#include <string>
#include "../exception/InvalidStateException.h"
#include <algorithm>
#include <vector>
#include <iostream>
#include <fstream>
#include <limits.h>
#include <string.h>
#include <cmath>
#include "GlobAlignE.h"

using namespace std;
using namespace utility;
using namespace exception;

GlobAlignE::GlobAlignE(const char * seq1In, int start1In, int end1In, const char * seq2In,
        int start2In, int end2In, int matchIn, int mismatchIn, int gapOpenIn, int gapContinueIn){

    seq1 = seq1In;
	start1 = start1In;
	end1 = end1In;

    seq2 = seq2In;
	start2 = start2In;
	end2 = end2In;

	len1 = end1 - start1 + 2;
    len2 = end2 - start2 + 2;

    //Incremental score storage
    matches = new int[len1];
    upperGap = new int[len1];
    lowerGap = new int[len1];


    //Incremental length storage
    matchLen = new int[len1];
    upperLen = new int[len1];
    lowerLen = new int[len1];

    //Incremental identity storage
    matchId = new int[len1];
    upperId = new int[len1];
    lowerId = new int[len1];

    match = matchIn;
    mismatch = mismatchIn;
    gapOpen = gapOpenIn;
    gapContinue = gapContinueIn;
    findAlignment();

}
/*
GlobAlignE::GlobAlignE(string filename1,string filename2, int matchIn, int mismatchIn, int gapOpenIn, int gapContinueIn):GlobAlignE(string1.c_str(),0,string.size(),string2.c_str(),0,string2.size(),matchIn,mismatchIn,gapOpenIn,gapContinueIn){

    ifstream ifs;

    ifs.open (filename1, ifstream::in);
    cout<<"FILE OPENED"<<endl;
    char c = ifs.get();

    if(c == '>'){

        while(c!='\n'){
            c = ifs.get();

        }
    }

     string string1  ="";

      while (ifs.good()) {


        if (c!='\n'){
        string1+=c;
        }
        c = ifs.get();
      }

      ifs.close();


     ifstream ifs2;

     ifs2.open (filename2, ifstream::in);

     c = ifs2.get();

     if(c == '>'){

         while(c!='\n'){
              c = ifs2.get();
         }
     }

     string string2  ="";

     while (ifs2.good()) {

         if(c!='\n'){
         string2+=c;
         }
         c = ifs2.get();
     }

     ifs2.close();

     std::transform(string1.begin(),string1.end(),string1.begin(),::toupper);
     std::transform(string2.begin(),string2.end(),string2.begin(),::toupper);

   // return GlobAlignE(string1.c_str(),0,string.size(),string2.c_str(),0,string2.size(),matchIn,mismatchIn,gapOpenIn,gapContinueIn);

}
*/
void GlobAlignE::findAlignment(){

    int shorter = min(len2,len1)-1;
    int lenDiff = abs(len2-len1);
    int maxDiff=0;

    if (lenDiff >=1){
        maxDiff += -gapOpen- (lenDiff*gapContinue);
    }

    maxDiff+= (mismatch* shorter)-1;

    const int negativeInf = maxDiff;

    matches[0]= 0;
    upperGap[0] = negativeInf;
    lowerGap[0] = negativeInf;

    matchLen[0] =0;
    upperLen[0] =0;
    lowerLen[0] =0;

    matchId[0] =0;
    upperId[0] = 0;
    lowerId[0] =0;

    //initial values
    for (int i = 1; i<len1;i++){
        upperGap[i] = negativeInf;
        matches[i] = negativeInf;
        lowerGap[i] = (-gapOpen)- (i*gapContinue);
        matchLen[i]=i;
        upperLen[i]=i;
        lowerLen[i]=i;
        matchId[i] =0;
        upperId[i] =0;
        lowerId[i] =0;
    }

    for( int j = 1;j<len2;j++){

        int matchLag = matches[0]; //used for calculation of matches
        int matchLenLag = matchLen[0];
        int matchIdLag = matchId[0];

        int upperGapLag = (-gapOpen)-((j-1)*gapContinue);
        int upperLenLag = j-1;
        int upperIdLag =0;

        for(int i =1;i<len1;i++){

            //compute values for upperGap
            int ygapBegin = matches[i]-(gapOpen+gapContinue);
            int ygapCont = upperGap[i]-gapContinue;

            int ans = max(ygapBegin,ygapCont);

            int store1 = upperGap[i];
            int store2 = upperLen[i];
            int store3 = upperId[i];

            upperGap[i] = ans;

            if( ans == ygapBegin){
                upperLen[i] = matchLen[i]+1;
                upperId[i] = matchId[i];
            }
            else if(ans == ygapCont){
                upperLen[i] = upperLen[i]+1;
                upperId[i] = upperId[i];
            }


            // compute values for match/mismatch
            char a= seq1[start1+i-1];
            char b = seq2[start2+j-1];
            int matchScore = (a == b) ? match : mismatch;

            int matched = matchLag + matchScore;

            int xgapEnd = lowerGap[i-1] + matchScore;

            int ygapEnd = upperGapLag+ matchScore;

            ans = max(max(matched,xgapEnd),ygapEnd);

            matchLag = matches[i]; //store current val matches in lag
            matches[i] =ans;

            int temp = matchLen[i];
            int save = matchId[i];

            if(ans == matched){
                matchLen[i] = matchLenLag+1;
                if(matchScore == match){
                    matchId[i] = matchIdLag+1;
                }
                else{
                    matchId[i] = matchIdLag;
                }
            }
            else if (ans == xgapEnd){
                matchLen[i] = lowerLen[i-1]+1;
                if(matchScore ==match){
                    matchId[i] = lowerId[i-1]+1;
                }
                else{
                    matchId[i] = lowerId[i-1];
                }
            }
            else{
                matchLen[i] = upperLenLag+1;
                if(matchScore ==match){
                matchId[i] = upperIdLag+1;
                }
                else{
                    matchId[i] = upperIdLag;
                }
            }
            matchLenLag = temp;
            matchIdLag = save;
            upperGapLag= store1;
            upperLenLag = store2;
            upperIdLag = store3;

        }

        matches[0] = negativeInf;
        matchLen[0] = j;
        matchId[0] =0;

        lowerGap[0]= negativeInf;
        lowerLen[0] = j;
        lowerId[0] =0;

        for(int i = 1;i<len1;i++){

                int xgapBegin = matches[i-1] -(gapOpen+gapContinue);
                int xgapCont = lowerGap[i-1]- gapContinue;
                int ans = max(xgapBegin,xgapCont);
                lowerGap[i]=ans;
                if(ans ==xgapBegin){
                    lowerLen[i] = matchLen[i-1]+1;
                    lowerId[i] = matchId[i-1];
                }
                else{
                    lowerLen[i] = lowerLen[i-1]+1;
                    lowerId[i] = lowerId[i-1];
                }

        }


    }

   alignmentScore= max(max(matches[len1-1], lowerGap[len1-1]), upperGap[len1-1]);

   if(alignmentScore == matches[len1-1]){
           alignmentLength = matchLen[len1-1];
           totalMatches = matchId[len1-1];
    }
    else if(alignmentScore == lowerGap[len1-1]){
            alignmentLength = lowerLen[len1-1];
            totalMatches= lowerId[len1-1];
    }
    else{
            alignmentLength = upperLen[len1-1];
            totalMatches = upperId[len1-1];
    }
}

int GlobAlignE::getScore(){
    return alignmentScore;
}
int GlobAlignE::getLength(){
    return alignmentLength;
}

double GlobAlignE::getIdentity(){
   double totalMatch = (double) totalMatches;

    return totalMatch/alignmentLength;
}
GlobAlignE::~GlobAlignE(){
    delete [] matches;
    delete [] upperGap;
    delete [] lowerGap;
    delete [] matchLen;
    delete [] upperLen;
    delete [] lowerLen;
    delete [] matchId;
    delete [] upperId;
    delete [] lowerId;

}
```
