## Supplementary Data 1 for "FASTCAR: Rapid alignment-free prediction of sequence alignment identity scores using self-supervised general linear models": GlobAlignE.h

```
/**
 * Author: Joseph Valencia
 * Modified by Benjamin James
 * Date: 12/14/17
 * Bioinformatics Toolsmith Laboratory, University of Tulsa
 * */
#ifndef Glob_AlignE_H_
#include <string>

using namespace std;

namespace utility{

class GlobAlignE{

private:
    const char * seq1; //first sequence to be aligned
    int start1;
    int end1;
    const char * seq2;//second sequence to be aligned
    int start2;
    int end2;
    int len1;
    int len2;
    int lenTotal;
    int match; //score for base pair match
    int mismatch;//score for base pair mismatch
    int gapOpen; //cost to open a gap
    int gapContinue; //cost to continue a gap
    int * matches;
    int * upperGap;
    int * lowerGap;
    int * matchLen;
    int * upperLen;
    int * lowerLen;
    int * matchId;
    int * upperId;
    int * lowerId;
    int alignmentScore;
    int alignmentLength;
    int totalMatches;
    string topString;
    string bottomString;
public:
    GlobAlignE(const char*,int,int,const char *,int,int, int,int,int,int);
    GlobAlignE(string,string,int,int,int,int);
    virtual ~GlobAlignE();
    void findAlignment();
    double getIdentity();
    int getLength();
    void printAlignment();
    int getScore();
    int getLengthAlignment();

};
}
#endif
```
