## Supplementary Data 1 for "FASTCAR: Rapid alignment-free prediction of sequence alignment identity scores using self-supervised general linear models": ILocation.h

```
/*
 * ILocation.h
 *
 *  Created on: Dec 20, 2012
 *      Author: Hani Zakaria Girgis, PhD
 */

#ifndef ILOCATION_H_
#define ILOCATION_H_

#include <string>

using namespace std;

namespace utility {

class ILocation {
public:
	virtual int getEnd() const = 0;
	virtual int getStart() const = 0;
	virtual void setEnd(int) = 0;
	virtual void setStart(int) = 0;
	virtual int getLength() = 0;
	virtual string toString() = 0;
};

}

#endif /* ILOCATION_H_ */
```
