## Supplementary Data 1 for "FASTCAR: Rapid alignment-free prediction of sequence alignment identity scores using self-supervised general linear models": LCSLen.cpp

```
/*
 * LCSLen.cpp
 *
 *  Created on: Dec 6, 2012
 *      Author: Hani Zakaria Girgis, PhD
 */

#include "LCSLen.h"
#include "../utility/Util.h"
#include "../exception/InvalidInputException.h"

#include <iostream>

using namespace std;
using namespace exception;

namespace utility {

LCSLen::LCSLen(const char * seq1In, int start1In, int end1In,
		const char * seq2In, int start2In, int end2In) {
	seq1 = seq1In;
	start1 = start1In;
	end1 = end1In;

	seq2 = seq2In;
	start2 = start2In;
	end2 = end2In;

	if(start1 < 0 || end1 < 0 || start1 > end1){
		string msg("Invalid Input. Start1 is ");
		msg.append(Util::int2string(start1));
		msg.append(". End 1 is ");
		msg.append(Util::int2string(end1));
		msg.append(".");
		throw InvalidInputException(msg);
	}

	if(start2 < 0 || end2 < 0 || start2 > end2){
		string msg("Invalid Input. Start2 is ");
		msg.append(Util::int2string(start2));
		msg.append(". End2 is ");
		msg.append(Util::int2string(end2));
		msg.append(".");
		throw InvalidInputException(msg);
	}

	// Validate input
	// cout << start1 << " " << end1 << endl;
	// cout << start2 << " " << end2 << endl;


	len1 = end1 - start1 + 2;
	len2 = end2 - start2 + 2;

	lenTotal = 2 * len2;
	cTable = new int[lenTotal];

	for (int i = 0; i < lenTotal; i++) {
		cTable[i] = 0;
	}

	findLcs();
}

LCSLen::~LCSLen() {
	delete[] cTable;
}

void LCSLen::findLcs() {
	int iM1Index = 0;
	int iIndex = len2;

	for (int i = 1; i < len1; i++) {
		char base1 = seq1[start1 + i - 1];

		for (int j = 1; j < len2; j++) {
			int ijIndex = iIndex + j;
			if (base1 == seq2[start2 + j - 1]) {
				cTable[ijIndex] = cTable[iM1Index + j - 1] + 1;
			} else {
				if (cTable[iM1Index + j] > cTable[iIndex + j - 1]) {
					cTable[ijIndex] = cTable[iM1Index + j];
				} else {
					cTable[ijIndex] = cTable[iIndex + j - 1];
				}
			}
		}

		if(i != len1-1){
			for(int h = 0; h < len2; h++){
				cTable[h] = cTable[len2+h];
			}
		}
	}
	lenCS =  cTable[lenTotal-1];
}

int LCSLen::getLenCS(){
	return lenCS;
}

}
/* namespace utility */
```
