## Supplementary Data 1 for "FASTCAR: Rapid alignment-free prediction of sequence alignment identity scores using self-supervised general linear models": LCSLen.h

```
/*
 * LCSLen.h
 *
 *  Created on: Dec 6, 2012
 *      Author: Hani Zakaria Girgis, PhD
 */

#ifndef LCSLEN_H_
#define LCSLEN_H_

namespace utility {

class LCSLen {
private:
	const char * seq1;
	int start1;
	int end1;
	const char * seq2;
	int start2;
	int end2;

	int len1;
	int len2;
	int lenTotal;
	int lenCS;

	int * cTable;
	void findLcs();

public:
	LCSLen(const char *, int, int, const char *, int, int);
	virtual ~LCSLen();
	int getLenCS();
};

} /* namespace utility */
#endif /* LCSLEN_H_ */
```
