## Supplementary Data 1 for "FASTCAR: Rapid alignment-free prediction of sequence alignment identity scores using self-supervised general linear models": Location.cpp

```
/*
 * Location.cpp
 *
 *  Created on: Dec 19, 2012
 *      Author: Hani Zakaria Girgis, PhD
 */

#include "Location.h"
#include "Util.h"
#include "../exception/InvalidInputException.h"

using namespace exception;

namespace utility {

Location::Location(int startIn, int endIn) {
	initialize(startIn, endIn);
}

Location::Location(ILocation& cp) {
	initialize(cp.getStart(), cp.getEnd());
}

void Location::initialize(int startIn, int endIn) {
	start = startIn;
	end = endIn;
	check();

}

void Location::check() {
	if (start < 0 || end < 0 || start > end) {
		string msg("Invalid Input. Start is ");
		msg.append(Util::int2string(start));
		msg.append(". End is ");
		msg.append(Util::int2string(end));
		msg.append(".");
		throw InvalidInputException(msg);
	}
}

Location::~Location() {
}

int Location::getEnd() const {
	return end;
}

int Location::getStart() const {
	return start;
}

void Location::setEnd(int endIn) {
	end = endIn;
	check();
}

void Location::setStart(int startIn) {
	start = startIn;
	check();
}

int Location::getLength() {
	return end - start + 1;
}

string Location::toString() {
	string msg = (Util::int2string(start));
	msg.append("-");
	msg.append(Util::int2string(end));

	return msg;
}
}
```
