## Supplementary Data 1 for "FASTCAR: Rapid alignment-free prediction of sequence alignment identity scores using self-supervised general linear models": Location.h

```
/*
 * Location.h
 *
 *  Created on: Dec 19, 2012
 *      Author: Hani Zakaria Girgis, PhD
 */

#ifndef LOCATION_H_
#define LOCATION_H_

#include "ILocation.h"

#include <string>

using namespace std;

namespace utility {

class Location : public ILocation{
private:
	int start;
	int end;
	void initialize(int, int);
	void check();

public:
	Location(int, int);
	Location(ILocation&);
	virtual ~Location();

	int getEnd() const;
	int getStart() const;
	void setEnd(int);
	void setStart(int);
	int getLength();
	string toString();
};

}

#endif /* LOCATION_H_ */
```
