## Supplementary Data 1 for "FASTCAR: Rapid alignment-free prediction of sequence alignment identity scores using self-supervised general linear models": Util.cpp

```
/*
 * Util.cpp
 *
 *  Created on: Apr 24, 2012
 *      Author: Hani Zakaria Girgis, PhD
 *      This class has a collection of utilities.
 */
#include "Util.h"

Util::Util() {
	// TODO Auto-generated constructor stub

}

Util::~Util() {
	// TODO Auto-generated destructor stub
}

string Util::fileSeparator("/");

string * Util::emptyString = new string("");

bool Util::isDna = true;

const int Util::getAlphabetSize(){
	return Util::isDna? 4 : 22;
}

void Util::readFasta(string seqFile, vector<string> * infoList,
		vector<string> * seqList, bool canCheckFormat) {
	ifstream in(seqFile.c_str());
	string info;

	bool isFirst = true;
	string basePtr("");

	while (in.good()) {
		string line;
		getline(in, line);
		if (line[0] == '>') {
			if (canCheckFormat) {
				int colIndex = line.find_first_of(':');
				int dashIndex = line.find_first_of('-');
				if (colIndex < 0 || dashIndex < 0) {
					string msg =
							"The header must be in the following format: chromosome:start-end\n";
					msg += "The current input: " + line;
					throw InvalidInputException(msg);
				}
			}

			infoList->push_back(line);
			if (!isFirst) {
				seqList->push_back(basePtr);
				basePtr = string("");
			} else {
				isFirst = false;
			}
		} else {
			basePtr.append(line);
		}
	}
	seqList->push_back(basePtr);
	in.close();

	// cout << "The system read " << infoList->size() << " sequences." << endl;

	// Post condition
	if (infoList->size() != seqList->size()) {
		cerr << "Error while reading the fasta input file. "
				<< "Header count = " << infoList->size() << " "
				<< "Sequence count = " << seqList->size() << endl;
		exit(1);
	}
}

void Util::readFasta(string seqFile, vector<string> * infoList,
		vector<string> * seqList) {
	ifstream in(seqFile.c_str());
	string info;

	bool isFirst = true;
	string * basePtr = new string("");
	while (in.good()) {
		string line;
		getline(in, line);
		if (line[0] == '>') {
			infoList->push_back(line);
			if (!isFirst) {
				seqList->push_back(*basePtr);
				basePtr = new string("");
			} else {
				isFirst = false;
			}
		} else {
			basePtr->append(line);
		}
	}
	seqList->push_back(*basePtr);
	in.close();

	// Post condition
	if (infoList->size() != seqList->size()) {
		cerr << "Error while reading the fasta input file. "
				<< "Header count = " << infoList->size() << " "
				<< "Sequence count = " << seqList->size() << endl;
		exit(1);
	}
}

void Util::readCoordinates(string fileName, vector<Location *> * coor) {
	checkFile(fileName);

	ifstream in(fileName.c_str());
	string line;

	while (in >> line) {
		int colIndex = line.find_first_of(':');
		int dashIndex = line.find_first_of('-');

		int start = atoi(line.substr(colIndex + 1, dashIndex - colIndex - 1).c_str());
		int end = atoi(line.substr(dashIndex + 1).c_str());
		Location * loc = new Location(start, end);
		coor->push_back(loc);
	}

	//cout << "Read ";
	//cout << coor->size() << endl;

	in.close();
}

void Util::readChromList(string genomeDir, vector<string> * chromList,
		string ext) {
	// This function may not be platform-independent
	// Credit: http://www.cplusplus.com/forum/beginner/9173/
	DIR * dirPtr = opendir(genomeDir.c_str());

	struct dirent * entry;
	entry = readdir(dirPtr);
	while (entry) {
		string file(entry->d_name);
		// Credit: http://stackoverflow.com/questions/51949/how-to-get-file-extension-from-string-in-c
		if (file.substr(file.find_last_of(".") + 1) == ext) {
			chromList->push_back(genomeDir + fileSeparator + entry->d_name);
		}
		entry = readdir(dirPtr);
	}

	closedir(dirPtr);
}

// This method will modify the contents of its parameter basePtr!
void Util::toUpperCase(string * basePtr) {
	string base = *basePtr;
	// Convert alphabet to upper case
	for (int i = 0; i < base.length(); i++) {
		base[i] = toupper(base[i]);
	}
}

void Util::toUpperCase(string& base) {
	// Convert alphabet to upper case
	for (int i = 0; i < base.length(); i++) {
		base[i] = toupper(base[i]);
	}
}

// credit: http://stackoverflow.com/questions/228005/alternative-to-itoa-for-converting-integer-to-string-c
string Util::int2string(int i) {
	string s;
	stringstream out;
	out << i;
	s = out.str();
	return s;
}

// Need to use templates
string Util::double2string(double i) {
	string s;
	stringstream out;
	out << i;
	s = out.str();
	return s;
}

string Util::long2string(long i) {
	string s;
	stringstream out;
	out << i;
	s = out.str();
	return s;
}

void Util::checkFile(string fileName) {
	ifstream f1(fileName.c_str());
	if (!f1) {
		string message = string("ERROR: ");
		message.append(fileName);
		message.append(" does not exist.\n");
		throw FileDoesNotExistException(message);
	}
	f1.close();
}

void Util::deleteFile(string fileName) {
	ifstream f1(fileName.c_str());
	if (f1) {
		if (remove(fileName.c_str()) != 0) {
			cerr << "Could not remove: " << fileName << endl;
		} else {
			cout << "Deleting: " << fileName << endl;
		}
	} else {
		cerr << "Warning! This file does not exist: " << fileName << endl;
	}
	f1.close();
}

void Util::deleteFilesUnderDirectory(string dirName) {
	// This function may not be platform-independent
	// Credit: http://www.cplusplus.com/forum/beginner/9173/
	DIR * dirPtr = opendir(dirName.c_str());
	struct dirent * entry;
	entry = readdir(dirPtr);
	while (entry) {
		string file(entry->d_name);
		if (file.compare(string(".")) == 0 || file.compare(string("..")) == 0) {
			// Skip current and parent directories
		} else {
			string url = dirName;
			url.append(fileSeparator);
			url.append(file);
			deleteFile(url);
			// cerr << "Deleting " << file << endl;
		}
		entry = readdir(dirPtr);
	}
	closedir(dirPtr);
}

bool Util::isOverlapping(int s1, int e1, int s2, int e2) {
	if (s1 > e1) {
		string msg("Util::isOverlapping. Invalid Input. s1 is ");
		msg.append(Util::int2string(s1));
		msg.append(". e1 is ");
		msg.append(Util::int2string(e1));
		msg.append(".");
		throw InvalidInputException(msg);
	}

	if (s2 > e2) {
		string msg("Util::isOverlapping. Invalid Input. s2 is ");
		msg.append(Util::int2string(s2));
		msg.append(". e2 is ");
		msg.append(Util::int2string(e2));
		msg.append(".");
		throw InvalidInputException(msg);
	}

	bool isStartWithin = s2 >= s1 && s2 <= e1;
	bool isEndWithin = e2 >= s1 && e2 <= e1;
	bool isIncluding = s2 >= s1 && e2 <= e1;
	bool isIncluded = s1 >= s2 && e1 <= e2;
	bool isAdjacent = (e1 == (s2 + 1)) || (e2 == (s1 + 1));

	return (isStartWithin || isEndWithin || isIncluding || isIncluded
			|| isAdjacent);
}

/**
 * The input string is s.
 * The reverse complement is rc.
 * The start, and the end are inclusive.
 */
void Util::revCompDig(const char * s, int start, int end, string * rc) {
	for (int i = end; i >= start; i--) {
		char b = s[i];
		switch (b) {
		case 0:
			rc->append(1, 3);
			break;
		case 3:
			rc->append(1, 0);
			break;
		case 1:
			rc->append(1, 2);
			break;
		case 2:
			rc->append(1, 1);
			break;
		default:
			string msg("Valid codes are 0-3. The invalid code is ");
			msg.append(1, b);
			throw InvalidInputException(msg);
		}
	}
}

void Util::revCompDig(string * s, string * rc) {
	revCompDig(s->c_str(), 0, s->size() - 1, rc);

	/*
	 int len = s->size();
	 for (int i = len - 1; i >= 0; i--) {
	 char b = s->at(i);
	 switch (b) {
	 case 0:
	 rc->append(1, 3);
	 break;
	 case 3:
	 rc->append(1, 0);
	 break;
	 case 1:
	 rc->append(1, 2);
	 break;
	 case 2:
	 rc->append(1, 1);
	 break;
	 default:
	 string msg("Valid codes are 0-3. The invalid code is ");
	 msg.append(1, b);
	 throw InvalidInputException(msg);
	 }
	 }
	 */
}

void Util::writeFasta(const string& sequence, const string& header,
		const string& outputFile) {
	ofstream outMask;
	outMask.open(outputFile.c_str(), ios::out);
	outMask << header << endl;
	int step = 50;
	int len = sequence.size();
	for (int i = 0; i < len; i = i + step) {
		int e = (i + step - 1 > len - 1) ? len - 1 : i + step - 1;
		for (int k = i; k <= e; k++) {
			outMask << sequence[k];
		}
		outMask << endl;
	}
	outMask.close();
}

int Util::sumTotalLength(const vector<ILocation *> * list) {
	int size = list->size();
	int sum = 0;
	for (int i = 0; i < size; i++) {
		sum += list->at(i)->getLength();
	}
	return sum;
}
```
