## Supplementary Data 1 for "FASTCAR: Rapid alignment-free prediction of sequence alignment identity scores using self-supervised general linear models": Util.h

```
/*
 * Util.h
 *
 *  Created on: Apr 24, 2012
 *      Author: Hani Zakaria Girgis, PhD
 */

#ifndef UTIL_H_
#define UTIL_H_

#include "Location.h"
#include "../exception/FileDoesNotExistException.h"
#include "../exception/InvalidInputException.h"

#include <vector>
#include <string>
#include <iostream>
#include <fstream>
#include <sstream>
#include <stdlib.h>
#include <dirent.h>

using namespace std;
using namespace utility;
using namespace exception;

namespace utility {
class Util {
private:
	Util();
	~Util();

public:
	static string * emptyString;
	static string fileSeparator;
	static bool isDna;
	static void readFasta(string, vector<string> *, vector<string> *, bool);
	static void readFasta(string, vector<string> *, vector<string> *);
	static void readCoordinates(string, vector<Location *> *);
	static void readChromList(string, vector<string> *, string);
	static void toUpperCase(string*);
	static void toUpperCase(string&);
	static string int2string(int);
	static string double2string(double);
	static string long2string(long);
	static void deleteFile(string);
	static void deleteFilesUnderDirectory(string);
	static void checkFile(string);
	static bool isOverlapping(int, int, int, int);
	static void revCompDig(string *, string *);
	static void revCompDig(const char* sequence, int, int, string *);

	static void writeFasta(const string&, const string&, const string&);

	static int sumTotalLength(const vector<ILocation *> *);

	// Added on Oct 6 2018
	static const int getAlphabetSize();

	/**
	 * Delete the objects pointed to by pointers in a vector.
	 * It does not delete the vector itself.
	 *
	 * Credit: http://stackoverflow.com/questions/594089/does-stdvector-clear-do-delete-free-memory-on-each-element
	 */
	template<class T>
	static void deleteInVector(vector<T*> * deleteMe) {
		while (!deleteMe->empty()) {
			delete deleteMe->back();
			deleteMe->pop_back();
		}

		// Set the size to zero
		deleteMe->clear();

		// Set the capacity to zero
		vector<T*> empty;
		deleteMe->swap(empty);
	}
};
}

#endif /* UTIL_H_ */
```
