## Supplementary Data 2 for "FASTCAR: Rapid alignment-free prediction of sequence alignment identity scores using self-supervised general linear models": draw_newick.py

```
#!/usr/bin/env python3

from ete3 import Tree
import sys
import argparse
import subprocess as sp
def draw(in_f, out_f):
        t = Tree(in_f)
        t.render(out_f)


if __name__ == "__main__":
        parser = argparse.ArgumentParser(description="Use all-vs-all to draw hierarchical clustering")
        parser.add_argument('in_dir', type=str, help="the input directory")
        parser.add_argument('method', type=str, help="the hierarchical methods passed to R's hclust, namely, one of ward.D, ward.D2, single, complete, average, mcquitty, median, centroid")
        parser.add_argument('tree', type=str, help="the Newick format tree to save to")
        parser.add_argument('out', type=str, help="the rendered output file")

        args = parser.parse_args()
        sp.run(["Rscript", "phylo.R", args.in_dir, args.method, args.tree])
        draw(args.tree, args.out)
```
