## Supplementary Data 2 for "FASTCAR: Rapid alignment-free prediction of sequence alignment identity scores using self-supervised general linear models": fastcar_all_vs_all.pl

```
#!/usr/bin/env perl
use warnings;
use strict;
use Bio::SeqIO;
use Data::Dumper;

sub main {

	my %options = parse(@_);
        my $fastcar_binary = $options{"fastcar"};
        my $fastcar_options = $options{"options"};
	my @db_file_list = @{$options{"dataset"}};
	my $weights_file = "weight.txt";
	if ($options{"weight"} eq "") {
		# no weights so run on db to get weights
		my $run_str = $fastcar_binary . " " . $fastcar_options . " " . join(" ", @db_file_list) . " --dump " . $weights_file;
		print("$run_str\n");
		qx($run_str);
	} else {
		# precomputed weights - use them
		$weights_file = $options{"weight"};
	}
	my @single_files;
	if ($options{"splitfiles"}) {
		# files are already split so don't re-split them
		@single_files = @db_file_list;
	} else {
		# split the files in db_file_list to single files in fa_dir
		@single_files = fasplit($options{"fa_dir"}, $options{"fa_per_file"}, \@db_file_list);
	}
	open(NOUT, ">", $options{"output_names"}) or die "could not open " . $options{"output_names"} . " $!";
	for my $f (@single_files) {
		my $line = $f;
		$line =~ s/.[^.]+$//;
		print NOUT "$line\n";
	}
	close(NOUT);
	open(OUT, ">", $options{"output_matrix"}) or die "could not open " . $options{"output_matrix"} . ": $!";
	for (my $i = 0; $i <= $#single_files; $i++) {
		print "$i\n";
		for (my $j = 0; $j < $i; $j++) {
			print OUT "0 ";
		}
		print OUT "100 ";
		# for (my $j = $i + 1; $j <= $#single_files; $j++) {
		# 	my $output = $i . "_" . $j . "_out";
		# 	my $fc_run = $fastcar_binary . " " . $fastcar_options . " -q " . $single_files[$i]  . " --recover " . $weights_file . " " . $single_files[$j] . " --output $output";
		# 	print($fc_run . "\n");
		# 	qx($fc_run);
		# 	print OUT postprocess($i, $j, $output);
		# }
		open(my $f1, ">", "sf_list.txt");
		for (my $k = $i + 1; $k <= $#single_files; $k++) {
		    print $f1 $single_files[$k] . "\n";
		}
		close($f1);
		my $run_str = "< sf_list.txt parallel \"" . $fastcar_binary . " " . $fastcar_options . " -t 1 -q " . $single_files[$i]  . " --recover " . $weights_file . " {} --output {.}.out\"";
		qx($run_str);

		for (my $j = $i + 1; $j <= $#single_files; $j++) {
			my $output = $single_files[$j];
			$output =~ s/\.[^.]*$/.out/;
			print OUT postprocess($i, $j, $output);
		}
		print OUT "\n";
	}

	close(OUT);

}

sub postprocess {
	my ($i, $j, $input) = @_;
	my $out = "";
	for my $fname (glob $input . "*") {
		open(my $in, '<', $fname) or die "could not open $fname: $!";
		my $found = 0;
		while (<$in>) {
			chomp;
			if (/\s+(\S+)$/) {
				$found = 1;
				$out .= $1 . " ";
			}
		}
		if ($found == 0) {
			$out .= "0 ";
		}
		close($in);
		system("rm $fname");
#		unlink($fname);
	}
	return $out;
}
sub fasplit {
	my ($fa_dir, $fa_per_file, $files_ref) = @_;
	my @ret_list = ();
	my $total_idx = 0;
	for my $file (@{$files_ref}) {
		my $seqio = Bio::SeqIO->new(-file => $file, '-format' => 'Fasta');
		while (my $seq = $seqio->next_seq) {
			my $outfile = $fa_dir . "/" . $total_idx . ".fa";
			my $newseq = Bio::SeqIO->new(-file => ">" . $outfile,
						     -format => 'fasta');
			# for (my $f_idx = 0; $f_idx < $fa_per_file; $f_idx++) {
			# 	if ($seq) {
					$newseq->write_seq($seq);
			# 		$seq = $seqio->next_seq;
			# 	}
			# }
			$total_idx++;
			push @ret_list, $outfile;
		}
	}
	return @ret_list;
}
sub parse {
	my @arg_list = @_;
	my %table = (
		     "fastcar" => "fastcar",
		     "options" => "",
		     "dataset" => [],
		     "fa_per_file" => 1,
		     "weight" => "",
		     "fa_dir" => ".",
		     "output_matrix" => "matrix.txt",
		     "output_names" => "names.txt"
		    );
	my $l1 = $#arg_list - 1;
	for (my $i = 0; $i <= $#arg_list; $i++) {
	#	print "$i => $arg_list[$i]\n";
		my $a = $arg_list[$i];

		if ($a eq "--fastcar" && $i <= $l1) {
			$i++;
			$table{"fastcar"} = $arg_list[$i];
		} elsif ($a eq "--output-matrix" && $i <= $l1) {
			$i++;
			$table{"output_matrix"} = $arg_list[$i];
		} elsif ($a eq "--output-names" && $i <= $l1) {
			$i++;
			$table{"output_names"} = $arg_list[$i];
		} elsif ($a eq "--weight" && $i <= $l1) {
			$i++;
			$table{"weight"} = $arg_list[$i];
		} elsif ($a eq "--files-are-split") {
			$table{"splitfiles"} = 1;
		} elsif ($a eq "--fa-per-file" && $i <= $l1) {
			$i++;
			$table{"fa_per_file"} = $arg_list[$i];
		} elsif ($a eq "--fa-dir" && $i <= $l1) {
			$i++;
			$table{"fa_dir"} = $arg_list[$i];
		} elsif ($a eq "--help") {
			die "Options: $0 --fastcar /path/to/fastcar database_files [--output-matrix /path/to/matrix.txt] [--output-names /path/to/names.txt]\n" . " [--weight precomputed_weights.txt] [--fc-opt1 --fc-opt2] [--files-are-split] [--fa-dir .]\n";
		} elsif ($a =~ /^-/ && $i <= $l1) {
			$i++;
			$table{"options"} .= $a . " " . $arg_list[$i] . " ";
		} else {
			push @{$table{"dataset"}}, $a;
		}
	}
	print Dumper(\%table);
	if (scalar(@{$table{"dataset"}}) == 0) {
		print "Please supply a dataset.\n";
		die "Options: $0 --fastcar /path/to/fastcar database_files [--output-matrix /path/to/matrix.txt] [--output-names /path/to/names.txt]\n" . " [--weight precomputed_weights.txt] [--fc-opt1 --fc-opt2] [--files-are-split] [--fa-dir .]\n";

	}
	return %table;
}
main(@ARGV);
```
