## Supplementary Data 2 for "FASTCAR: Rapid alignment-free prediction of sequence alignment identity scores using self-supervised general linear models": phylo.R

```
#!/usr/bin/env Rscript

library(phangorn)
library(Matrix)
hierarchical = function(d, mode="single", out="tree.newick") {
  d = paste(d,"/",sep="")
  mfile = paste(d,"matrix.txt", sep="")
  lfile = paste(d,"names.txt", sep="")
  mat = as.matrix(read.csv(mfile, sep=" ", header=FALSE))
  mat <- mat[,1:(ncol(mat)-1)]
  names = readLines(lfile)
  sim_mat = 100 - (as.matrix(t(triu(mat)) + mat - diag(diag(mat))))
  colnames(sim_mat) = names
  rownames(sim_mat) = names
  hc = hclust(as.dist(sim_mat), mode)
  write.tree(phy=as.phylo(hc), file=out)
  return(hc)
}

args = commandArgs(trailingOnly=TRUE)
if (length(args) != 3) {
  print("usage: phylo.R directory mode output_file")
} else {
  hierarchical(args[1], args[2], args[3])
}
```
