## Supplementary Data 2 for "FASTCAR: Rapid alignment-free prediction of sequence alignment identity scores using self-supervised general linear models": README

FASTCAR ALL VS ALL SCRIPT (proof of concept)

The fastcar_all_vs_all.pl file will do an upper triangular search on all vs all.
The arguments have to be provided correctly.

If fastcar is not in the user's PATH variable, the --fastcar parameter will allow a specific binary to be used.

A dataset must be provided. These files are provided as arguments without a parameter (just like FASTCAR).

Then, the script breaks up every sequence into its own FASTA file in the --fa-dir (parameter) directory.
If the files provided are already single files, the --files-are-split parameter doesn't re-split them.

The output matrix is output using the --output-matrix parameter, and the output names are output to --output-names.
Both of those files are required to draw the tree.

If weights have already been calculated for the dataset, the --weight parameter can provide FASTCAR with the weight file.

Other options not matched are then passed to every FASTCAR instance, such as identity percentage, mutation model, etc.


TREE DRAWING SCRIPTS
The draw_newick.py and phylo.R are two scripts that use the matrix and names files output by the above all versus all script.
The ./draw_newick.py --help gives all the options and their descriptions:

usage: draw_newick.py [-h] in_dir method tree out

Use all-vs-all to draw hierarchical clustering

positional arguments:
  in_dir      the input directory
  method      the hierarchical methods passed to R's hclust, namely, one of
              ward.D, ward.D2, single, complete, average, mcquitty, median,
              centroid
  tree        the Newick format tree to save to
  out         the rendered output file

optional arguments:
  -h, --help  show this help message and exit
